# An activity-guided map of electrophile-cysteine interactions in primary human immune cells

**DOI:** 10.1101/808113

**Authors:** Ekaterina V. Vinogradova, Daniel C. Lazar, Radu M. Suciu, Yujia Wang, Giulia Bianco, Yu Yamashita, Vincent M. Crowley, Dave Remillard, Kenneth M. Lum, Gabriel M. Simon, Esther K. Kemper, Michael R. Lazear, Sifei Yin, Megan M. Blewett, Melissa M. Dix, Nhan Nguyen, Maxim N. Shokhirev, Emily Chin, Luke Lairson, Stefano Forli, John R. Teijaro, Benjamin F. Cravatt

## Abstract

Electrophilic compounds originating from nature or chemical synthesis have profound effects on immune cells. These compounds are thought to act by cysteine modification to alter the functions of immune-relevant proteins; however, our understanding of electrophile-sensitive cysteines in the human immune proteome remains limited. Here, we present a global map of cysteines in primary human T cells that are susceptible to covalent modification by electrophilic small molecules. More than 3000 covalently liganded cysteines were found on functionally and structurally diverse proteins, including many that play fundamental roles in immunology. We further show that electrophilic compounds can impair T cell activation by distinct mechanisms involving direct functional perturbation and/or ligand-induced degradation of proteins. Our findings reveal a rich content of ligandable cysteines in human T cells, underscoring the potential of electrophilic small molecules as a fertile source for chemical probes and ultimately therapeutics that modulate immunological processes and their associated disorders.

---

Modern human genetics has emphasized the central role that the immune system plays as both a causal basis for and guardian against a broad range of diseases (Hansen et al., 2018; Lucherini et al., 2018; Sims et al., 2017). This knowledge has, in specific instances, been translated into compelling new medicines, including drugs that stimulate immune responses to cancer (Ribas and Wolchok, 2018; Wei et al., 2018) and suppress autoimmunity in diseases like rheumatoid arthritis (Pesu et al., 2005; Telliez et al., 2016). Advanced profiling technologies, such as RNA-sequencing (Chen et al., 2019; Monaco et al., 2019; Papalexi and Satija, 2018; Stubbington et al., 2017) and mass spectrometry (MS)-based proteomics (Dybas et al., 2019; Hubel et al., 2019; Myers et al., 2019; Rebsamen et al., 2013; Rieckmann et al., 2017; Tan et al., 2017), have also improved our understanding of the molecular and cellular composition of the human immune system, leading to the discovery of many genes and proteins with enriched expression in specific subtypes of immune cells. Despite these advances, most immunologically relevant proteins lack chemical probes, and this gap hinders the pharmacological study of immune system function for basic and translational research purposes.

Among the various categories of chemical probes, electrophilic small molecules may offer a particularly attractive class for studying the functions of proteins involved in innate and adaptive immune responses. Multiple immunomodulatory drugs, either approved (Saidu et al., 2019) or in clinical development (Rip et al., 2018; Telliez et al., 2016), covalently modify cysteine residues in key immune-relevant proteins. Cysteines appear more generally to serve as important electrophilic and oxidative stress sensors in immunological signaling pathways, possibly providing a means for the immune system to swiftly respond to diverse environmental insults (Belikov et al., 2015; Franchina et al., 2018; Morgan and Liu, 2011; Nathan and Cunningham-Bussel, 2013). From a chemical biology perspective, electrophilic compounds, by combining features of molecular recognition and reactivity, can target proteins, and sites on proteins, that have proven challenging to address with reversibly binding small molecules (Backus et al., 2016; Maurais and Weerapana, 2019; Roberts et al., 2017). Finally, the durability of irreversible binding afforded by electrophilic compounds provides a direct means, when coupled with modern proteomic methods, to broadly and deeply map small molecule-protein interactions in native biological systems (Backus et al., 2016; Bar-Peled et al., 2017; Grossman et al., 2017).

In previous studies, we used chemical proteomics to identify a discrete set of cysteines targeted by the immunosuppressive drug dimethyl fumarate (DMF) in primary human T cells (Blewett et al., 2016). DMF-sensitive sites included cysteines at protein-protein interaction sites in interleukin-1 receptor-associated kinase 4 (IRAK4) (Zaro et al., 2019) and protein kinase C theta (PRKCQ) (Blewett et al., 2016), underscoring the potential for electrophilic compounds to engage atypical (i.e., non-active) functional sites on proteins. Inspired by these initial findings, we sought herein to generate a global portrait of the reactivity and ligandability (i.e., covalent small-molecule interactions) of cysteine residues in primary human T cells and to integrate these results with human genetic and expression profiling databases to furnish a resource of immune-relevant proteins that can be targeted by electrophilic small molecules. Leveraging a site-specific chemical proteomic platform and broadly reactive electrophilic fragments, we report the discovery of 3466 liganded cysteines in 2283 proteins (from a total of 18077 quantified cysteines in 5090 proteins) and show that these cysteines populate diverse adaptive and innate immune signaling networks, as well as pathways involved in T cell differentiation and lineage commitment. We acquired further knowledge on the pharmacological effects and tractability of small molecule-cysteine interactions by deploying chemically elaborated electrophiles combined with a functional screen of T cell activation, which uncovered immunomodulatory compounds that act by diverse mechanisms, including the direct inhibition of protein activity and induction of protein degradation.

## Results

### Chemical proteomic map of cysteine reactivity in activated T cells

Upon activation, T cells enter a growth phase associated with a number of biochemical changes that include alterations in cellular redox state (Belikov et al., 2015; Franchina et al., 2018; Simeoni and Bogeski, 2015), cytoskeletal rearrangements (Acuto and Cantrell, 2000), and increased glycolytic and mitochondrial metabolism (Almeida et al., 2016; Geltink et al., 2018). The molecular pathways that both execute and are influenced by these changes have been studied by global gene and protein expression (Hiemer et al., 2019; Rieckmann et al., 2017), as well as phosphoproteomic (Chylek et al., 2014; Tan et al., 2017) and metabolomic (Hiemer et al., 2019), analyses that compare resting versus activated T cells. Some of the discovered changes in activated T cells occur in general biochemical pathways associated with, for instance, cell proliferation, while others reflect immune-restricted processes. The extent to which these types of activation state-dependent changes in the biochemistry of T cells might also create a landscape of new targets for chemical probes that regulate T cell function remains largely unexplored. Here, we set out to address this question on a global scale by using activity-based protein profiling (ABPP) methods to quantify cysteine reactivity (Weerapana et al., 2010) and electrophilic small-molecule interactions (Backus et al., 2016; Wang et al., 2014) in primary human T cells activated by T cell receptor (TCR) stimulation.

T cells were isolated from human blood using a Lymphoprep gradient followed by a negative selection (EasySep human T Cell isolation) kit and activated by exposure to anti-CD3 and anti-CD28 antibodies for three days (activated T cells, Figure 1A). Control T cells were obtained by transient activation with anti-CD3 and anti-CD28 antibodies for three days followed by expansion in culture in the presence of IL2 for an additional 10-12 days to return the cells to a quiescent state. Cysteine reactivity changes were measured by treating proteomic lysates from activated or expanded control T cells with a broad-spectrum, cysteine-directed iodoacetamide-desthiobiotin probe (IA-DTB) (Patricelli et al., 2016), protease digestion of the IA-DTB-treated proteomes, streptavidin enrichment of IA-DTB-labeled cysteine-containing peptides, and quantitative, multiplexed LC-MS-based proteomic analysis using tandem mass tags (TMT, 10-plex experiments; (Rauniyar and Yates, 2014)(Figures 1A). The cysteine reactivity profiles were then integrated with data from complementary proteomic experiments measuring protein expression changes in control versus activated T cells (Figures 1A). In total, more than 4800 proteins were quantified in expression-based (TMT-exp) proteomic experiments, the vast majority of which (~80%) were also quantified by cysteine reactivity profiling (TMT-ABPP) (**Table S1, S2**), and nearly a quarter of these proteins qualified as “immune-relevant” (Figure 1B, **Table S3**), based on immune cell-enriched expression profiles derived from public databases and/or human genetic evidence of causing, upon mutation, immune-related disorders (acquired from the OMIM (Online Mendelian Inheritance in Man) database) (see **Supplementary Methods**). We considered a protein to show altered expression if its abundance was elevated or reduced by greater than two-fold in activated T cells, and ~1100 proteins satisfied this requirement (Figure 1C), including several immune-relevant proteins (e.g., IL2RA, TNFAIP3) (Figure 1D), proteins involved in glycolysis, and proteins regulated by mTORC1 and MYC pathways that are known markers of T-cell activation and proliferation (Tan et al., 2017).

**Figure 1.**
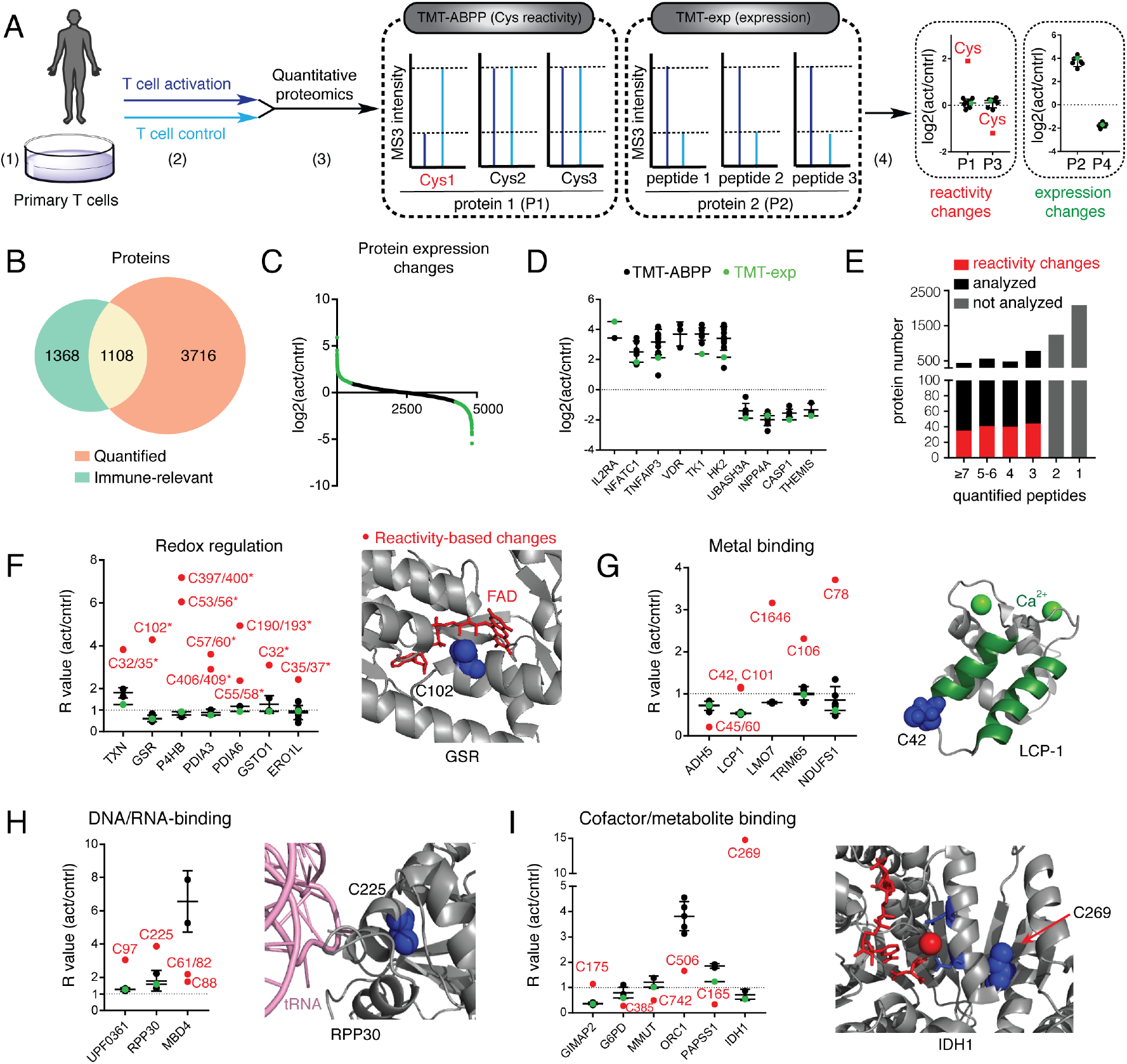
Chemical proteomic map of cysteine reactivity changes in activated T cells. (A) Experimental workflow for proteomic experiments measuring cysteine reactivity (TMT-ABPP) and protein expression (TMT-exp) in primary human T cells: 1) T cells were isolated from peripheral blood mononuclear cells (PBMCs) using a negative selection EasySep Human T Cell Isolation Kit; 2) T cells were exposed to αCD3 and αCD28 pre-coated 6-well plates (3 days) to furnish “activated T cells” (act); a subset of cells was transiently activated with lower amounts of antibodies (see **Supplementary Methods** for details) and then expanded by growing in RPMI media containing recombinant IL2 (10 U/mL) for 10-12 days, splitting the cells every 3-4 days, to return cells to a quiescent state, which represented “control (expanded, but not activated) T cells” (cntrl); 3) activated and control T cells were processed for TMT-ABPP and TMT-exp analysis (see Experimental Methods); and 4) cysteine reactivity and protein expression changes were distinguished by integrating TMT-ABPP and TMT-exp data (see Experimental Methods). (B) Overlap between proteins quantified by TMT-exp and immune-relevant proteins (see Experimental Methods for description of protocol used to designate proteins as being immune-relevant). Results are derived from two independent TMT-exp experiments (four biological replicates total), where a protein was required to have two unique peptides per experiment. (C) Protein expression differences between control and activated T cells. Results represent mean values from two independent TMT-exp experiments (four biological replicates). (D) Representative protein expression differences between control and activated T cells, where results from both TMT-ABPP (black dots) and TMT-exp (green dots) concordantly support the expression changes. Dashed line marks unchanged R value for act vs cntrl cells, and horizontal black lines for each protein measurement mark average R value for the quantified peptides from that protein. (E) Bar graph representation of the fraction of proteins with cysteine reactivity changes observed for proteins with the indicated numbers of quantified peptides in TMT-ABPP experiments. A cysteine was considered to show a reactivity change if the R value for its parent tryptic peptide differed more than two-fold from the protein expression value measured by TMT-exp (if quantified) and/or from other quantified cysteines on that same protein measured by TMT-ABPP (for proteins with ≥5 quantified cysteines). Proteins with only 1 or 2 quantified cysteines in TMT-ABPP experiments were not interpreted for reactivity changes (gray bars). (F-I) Representative cysteine reactivity changes in activated human T cells organized by functional categories. Dashed line marks unchanged R value for act vs cntrl cells, and horizontal black lines for each protein measurement mark average R value for the quantified peptides (excluding the reactivity-changing cysteine(s)) from that protein. (F) Active-site cysteines in redox-related proteins; x-ray crystal structure of FAD bound to the active site of GSR (PDB: 1GRF) with the reactivity-changing cysteine C102 highlighted in blue. (G) Metal-binding cysteines; solution NMR structure of the EF-hand domain of LCP-1 (PDB: 5JOJ) bound to calcium ions with the reactivity-changing cysteine C42 highlighted in blue. LCP-1 α-helices undergoing most significant rearrangement upon calcium binding are highlighted in green. (H) Cysteines at DNA/RNA-binding sites; cryo-EM structure of human ribonuclease P RPP30 (PDB: 6AHU) bound to mature tRNA with the reactivity-changing cysteine C225 highlighted in blue. (I) Cysteines at cofactor/metabolite-binding sites; x-ray crystal structure of the human NADP(+)-dependent isocitrate dehydrogenase 1 (IDH1) in complex with NADP, isocitrate, and calcium (PDB: 1T0L) with the reactivity-changing cysteine C269 highlighted in blue.

A distinct set of proteins (160 in total) harbored cysteine reactivity changes that differed substantially (>two-fold) from the corresponding expression profiles for these proteins in activated T cells (Figures 1E and **S1A, B** and **Table S4**). These cysteine reactivity changes were found in immune-relevant proteins (**Figure S1A**) and featured functional sub-groups that may reflect the diverse modulation of cellular biochemistry in activated T cells (**Figure S1B**). For example, a number of catalytic and active-site cysteines in proteins involved in redox regulation showed greater reactivity in activated T cells (Figure 1F), which is possibly a consequence of the higher intracellular reducing potential of these cells generated, at least in part, by increases in glutathione production (Lian et al., 2018; Mak et al., 2017). Reactivity changes were also found for cysteines in the metal-binding domains of proteins (Figure 1G), with one prominent example being the immune-relevant protein L-plastin (LCP-1), which is a calcium-regulated actin-binding protein that participates in remodeling of the actin cytoskeleton during T cell activation (Ishida et al., 2017). Calcium decreases the actin bundling efficiency of LCP-1 by inducing structural changes to the EF-hand motif of LCP-1, including α-helices 2 and 3 surrounding C42 (Figure 1G), which showed increased reactivity in activated T cells (Figure 1G). Additional reactivity changes were observed for cysteines at sites of DNA/RNA-protein (Figure 1H) and protein-protein (**Figure S1C**) interactions, as well as for cysteines proximal to cofactor- and metabolite-binding sites (Figures 1I and **S1D**). These cysteine reactivity changes may reflect a landscape of remodeled intermolecular interactions in activated T cells. As one example, C269 in isocitrate dehydrogenase 1 (IDH1) shows a striking increase in reactivity in activated T cells (Figure 1I), which could reflect changes in cofactor (NADP) and/or substrate (isocitrate) binding that have been shown to promote a structural rearrangement in residues 271-286 (Xu et al., 2004), which may, in turn, alter the reactivity of C269.

Having established quantitative proteome profiles that uncovered numerous cysteine reactivity and protein expression changes in activated T cells, we next turned our attention to mapping electrophilic small-molecule-cysteine interactions in this immunological cell state.

### Chemical proteomic map of cysteine ligandability in human T cells

We recently described an efficient strategy to globally assess the ligandability of cysteines in native biological systems that leverages broadly reactive, electrophilic small-molecule fragments referred to as “scouts” (Backus et al., 2016; Bar-Peled et al., 2017). Two scout fragments bearing an α-chloroacetamide (KB02) or acrylamide (KB05) (Figure 2A, 2B) – reactive groups frequently found in covalent chemical probes and drugs (Baillie, 2016; Chen et al., 2017; Honigberg et al., 2010; Ostrem et al., 2013; Xu et al., 2019; Yang et al., 2014; Zeng et al., 2017) – were used to construct in-depth cysteine ligandability maps across primary human T cells in both control and activated states. We analyzed scout fragment-cysteine interactions using two complementary chemical proteomic methods that provided a balance of confidence in quantitative accuracy (isoTOP-ABPP) with greater multiplexing capacity (TMT-ABPP) (Figures 2A and **S2A, B**). Both proteomic methods provided similar ratio (R) values (DMSO/scout fragment (500 μM, 1 h)) for cysteines in T cell proteomes, with the MS3-based quantification used in TMT-ABPP resulting in mild ratio compression (**Figure S2C, D**), which was countered by a substantial increase in the number of quantified cysteines compared to isoTOP-ABPP (**Figure S2E**). We accordingly designated cysteines as liganded if they showed an R value of ≥ 5 as measured by either isoTOP-ABPP or TMT-ABPP. From a total of 18077 cysteines in 5990 proteins quantified in human T cells, we identified 3466 liganded cysteines in 2283 proteins (Figure 2C, 2D and **Table S5**). These ligandability events were broadly distributed across cysteines with diverse intrinsic reactivities (Weerapana et al., 2010) (Figure 2E and **Table S6, S7**), indicating contributions from both the binding and electrophilic groups of scout fragments in conferring strong engagement of cysteines in the T cell proteome.

**Figure 2.**
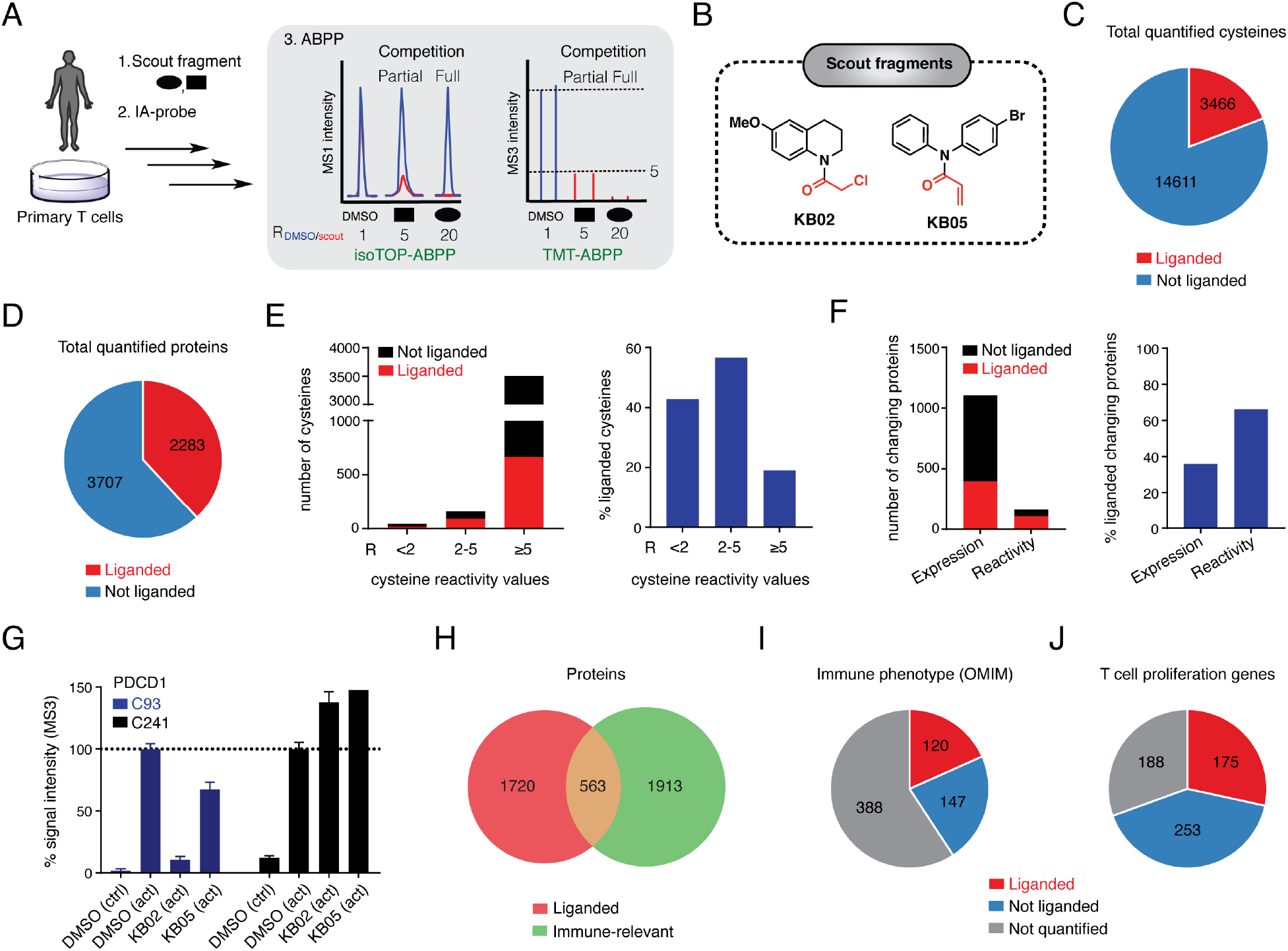
Chemical proteomic map of fragment electrophile-cysteine interactions in human T cells. (A) Experimental workflow for chemical proteomic experiments measuring scout fragment electophile effects on cysteine reactivity in primary human T cells: 1) T cells isolated from human blood were lysed by probe sonication (2 x 8 pulses) and the soluble and particulate fractions separated by ultracentrifugation (100,000 *g*, 45 min) and treated with DMSO or scout fragments (KB02, KB05; 500 μM, 1 h); 2) fractions were then treated with a broadly cysteine-reactive iodoacetamide (IA) probe (IA-alkyne or IA-desthiobiotin (DTB) (100 μM, 1 h) for isoTOP-ABPP and TMT-ABPP experiments, respectively); 3) DMSO- and fragment-treated T cell proteomes were analyzed by ABPP, where a cysteine was considered liganded if it displayed an R value (DMSO/scout fragment) of ≥5 (see Experimental Methods for more information). This protocol was applied to both control and activated T cells. (B) Structures of scout fragments KB02 and KB05. (C, D) Pie chart representations of cysteines (C) and proteins (D) liganded with scout fragments. Results are obtained by combining soluble and particulate proteomic data for KB02 and KB05 treatments (500 μM, 1 h) of both control and activated T cells. R-values within each experimental treatment group were derived from 3-5 independent isoTOP-ABPP experiments and 4 independent TMT-ABPP experiments (6 TMT channels per experiment). A cysteine was required to be quantified in at least two experiments for each compound treatment or proteomic fraction group to be reported. (E) Bar graphs showing the total number (left) and percentage (right) of liganded cysteines per total number of cysteines quantified across the indicated reactivity ranges, where cysteine reactivity was determined by isoTOP-ABPP experiments performed with different concentrations of the IA-alkyne probe (10 and 100 μM; R values = 100 μM/10 μM), as described previously (Weerapana et al., 2010). (F) Bar graphs showing the total number (left) and percentage (right) of liganded proteins with expression or reactivity changes in activated T cells. (G) Bar graphs showing percent MS3-signal intensity for quantified peptides from PDCD1, revealing elevated expression of this protein in activated T cells and KB02-sensitivity for C93 in these cells. Results represent mean R-values derived from TMT-ABPP experiments. Error bars represent SD or SE from 2-4 independent experiments. (H) Overlap of liganded proteins with immune-relevant proteins. (I) Fraction of liganded proteins from total proteins with human genetics-based immune phenotypes (reference list generated from the Online Mendelian Inheritance in Man (OMIM) database, see Supplementary Methods for details). (J) Fraction of liganded proteins from total proteins encoded by T cell proliferation genes (Shifrut et al., 2018).

Among the liganded cysteines were several targeted by existing covalent probes and drug candidates, including those being pursued for immunological disorders (e.g., C909 in JAK3 (Xu et al., 2019); C528 in XPO1 (Haines et al., 2015); **Table S8**). These results confirm that ABPP can “rediscover” established druggable sites on immune-relevant proteins. Liganded cysteines were also well-represented within the subset of proteins showing expression and/or cysteine reactivity changes in activated T cells, where cysteines with altered reactivity showed a greater propensity for liganding by scout fragments (Figure 2F). We identified, for instance, a liganded cysteine (C93) in the programmed cell death protein 1 (PDCD1 or PD-1), which was only observed in activated T cells (Figure 2G), likely reflecting the induced expression of this key immune checkpoint protein following T cell stimulation (Agata et al., 1996). Liganded cysteines showing reactivity-based changes included the catalytic cysteine in the deubiquitinase USP16 (C205) (**Figure S2F**), which has been shown to regulate hematopoietic stem cell differentiation (Gu et al., 2016).

Cross-referencing our ligandability map with the aforementioned set of immune-relevant proteins furnished >500 shared targets (Figure 2H and **Table S5**), of which 120 produce, upon mutation in humans, monogenic diseases with a strong immune phenotype (Figure 2I). We emphasize this point because such Mendelian genetic relationships to immune disorders can be used to prioritize proteins for drug development programs aimed at treating autoimmune or autoinflammatory disorders (e.g., JAK3 (Xu et al., 2019)) as well as promoting immune responses to cancer (e.g., CTLA4 (Wei et al., 2018)). Integrating our ligandability map with additional genomic and proteomic studies identified electrophile-sensitive cysteines in proteins: 1) found by genome-wide CRISPR screening to regulate T cell proliferation following stimulation (Shifrut et al., 2018) (Figure 2J and **Table S9**); and 2) that are part of immune-enriched modules, including those relevant for T cell activation (module 17, **Figure S2G**), established in expression-based proteomics studies (Rieckmann et al., 2017) (**Table S10**).

A more focused analysis revealed the striking breadth of liganded cysteines in key immune signaling pathways, including those mediating T cell- and TNF-receptor signaling and NF-κB activation (Figures 3A and **S3**). The proteins in these pathways harboring liganded cysteines derived from diverse structural and functional classes, including enzymes (e.g., DGKA/Z, IKBKB), adaptor proteins (e.g., MYD88) and transcription factors (e.g., NFKB1) (Figure 3A). Even for the more classically druggable proteins like kinases, we observed sites of cysteine ligandability that were far removed from the ATP-binding pocket (e.g., C464 in IKBKB, C406 in CHUK; Figure 3B), underscoring the potential for covalent ligands to engage non-canonical sites on proteins. Also supporting this conclusion was the large number of liganded cysteines identified in historically challenging to drug proteins like transcription factors and adaptor proteins, including a subset that have Mendelian links to immunological disorders (Figure 3C and **Table S11**). Some of these cysteines reside in proximity to DNA-protein (Figure 3D) and protein-protein (Figure 3E) interfaces important for the assembly and function of immune-relevant protein complexes.

**Figure 3.**
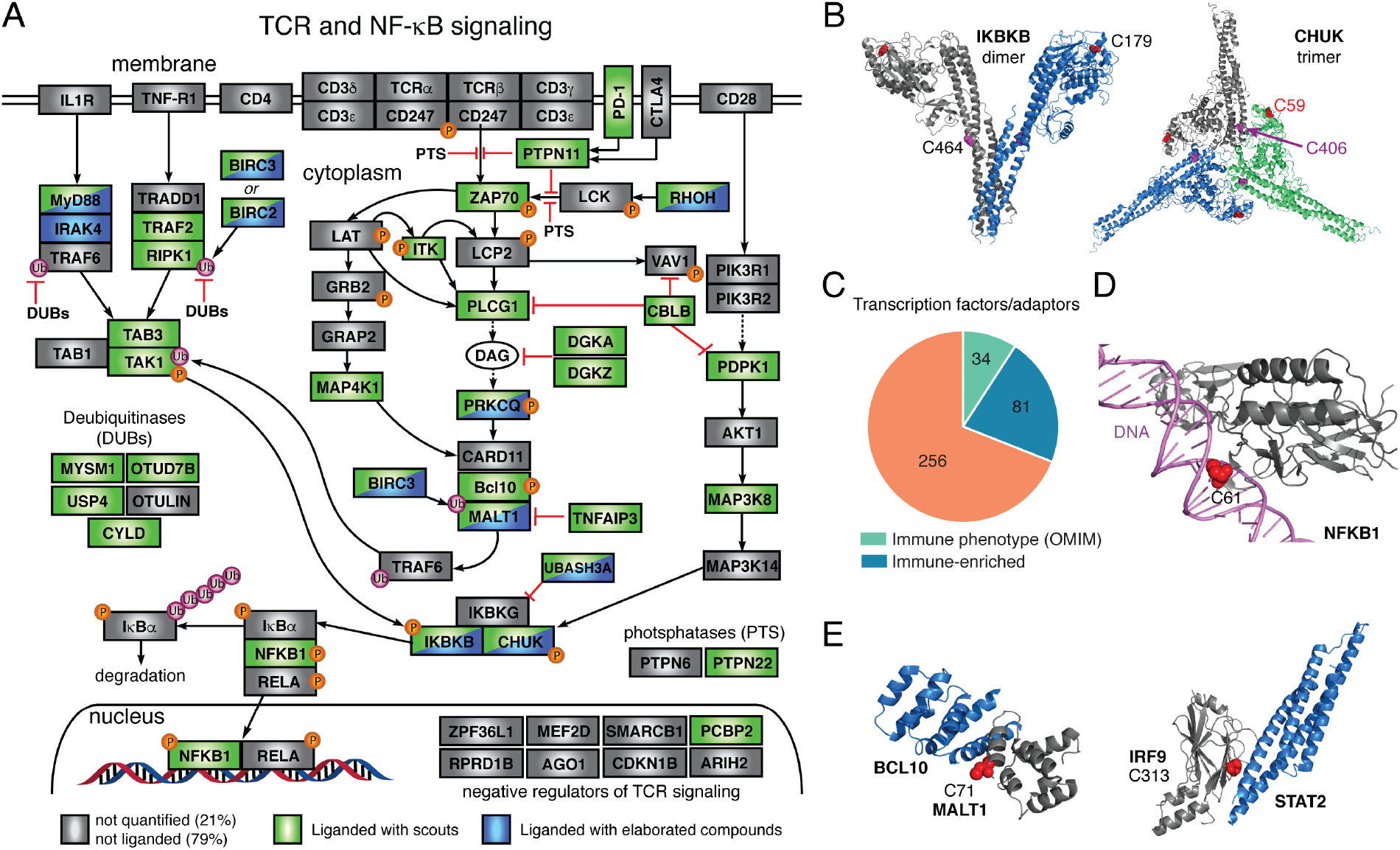
Liganded cysteines in immune-relevant proteins. (A) Diagram of TCR, TNF-alpha and NF-κB signaling pathways marking proteins that possess cysteines liganded by scout fragments (green) or elaborated electrophilic compounds (blue). (B) Location of liganded cysteines in three-dimensional structures of immune-relevant kinases IKBKB (PDB: 4E3C) and CHUK (PDB: 5EBZ), including non-active site cysteines (C464 in IKBKB, C406 in CHUK). (C) Location of the liganded cysteine at the site of NFKB1-DNA interaction (C61; PDB: 2O61). (D) Fractions of liganded transcription factors and adaptor proteins that are also immune-relevant (immune-enriched (blue) and/or have human genetics-based immune phenotypes (green, listed on the right)). Asterisks mark kinases that also possess adaptor functions and where non-active sites cysteines were liganded. (E) Location of liganded cysteines at sites of protein-protein interactions for MALT1 (C71; interaction partner BCL10; PDB: 6GK2) and IRF9 (C313 (mouse orthologue to human C319); interaction partner STAT2; PDB: 5OEN).

Our scout fragment profiling results, taken together, present an extensive landscape of ligandable cysteines in immune-relevant proteins, pointing to a potentially broad opportunity to discover and develop covalent chemical probes that modulate T cell function. We next aimed to establish an experimental workflow that would illuminate the functional effects and tractability of electrophilic small-molecule-cysteine interactions, while also preserving the globality and biological integrity afforded by profiling these interactions in primary human T cells.

### A functional screen of elaborated electrophilic compounds in T cells

Fragment-based screening, whether performed on a single protein of interest, or more broadly across the proteome as described herein, offers advantages for discovering hit compounds for challenging protein classes (Scott et al., 2012). Nonetheless, progressing fragments to more advanced chemical probes and drugs can be confounded by a variety of factors, including the low-affinity and promiscuity of initial hits and the tractability of fragment-binding sites on proteins (Murray et al., 2012; Scott et al., 2012). These problems have been historically addressed by labor-intensive, structure-guided protocols that require purified protein and have limited throughput. We aspired instead to create an *in cellulo* strategy that integrates phenotypic screening with our chemical proteomic methods to furnish structure-activity relationships (SARs) on many electrophilic small-molecule-cysteine interactions in parallel, such that the tractability and potential functional effects of these interactions could be comparatively evaluated.

We performed a multidimensional screen of a focused library of structurally elaborated electrophilic small molecules to identify compounds that suppress T cell activation at low-μM concentrations without causing cytotoxicity (Figure 4A). Primary human T cells isolated from the blood of healthy donors were activated with anti-CD3 and anti-CD28 antibodies in the presence of an in-house collection of electrophilic compounds (10 μM each; ~130 compounds tested in total; average compound MW = 400 Da; **Table S12**), where suppression of T cell activation was measured 24 h later by reductions in secreted IL2 and IFN-γ and in the expression of the cell surface markers CD69 and CD25 (Figure 4A). The viability of T cells was monitored by flow cytometry using a near-IR live-dead stain. 17 compounds, along with the immunosuppressive drug DMF, which was used as a positive control, were found to substantially suppress T cell activation (>65%) without causing cytotoxicity (viability > 85%) (Figure 4B–D, **Figure S4A**, and **Table S13**). The active compounds included different classes of electrophiles, of which representative acrylamides (BPK-21, BPK-25) and α-chloroacetamides (EV-3, EV-93) (Figure 4C) were selected for further characterization based on their concentration-dependent profiles, which revealed near-complete blockade of T cell activation at < 20 μM with negligible cytotoxicity (Figure 4D).

**Figure 4.**
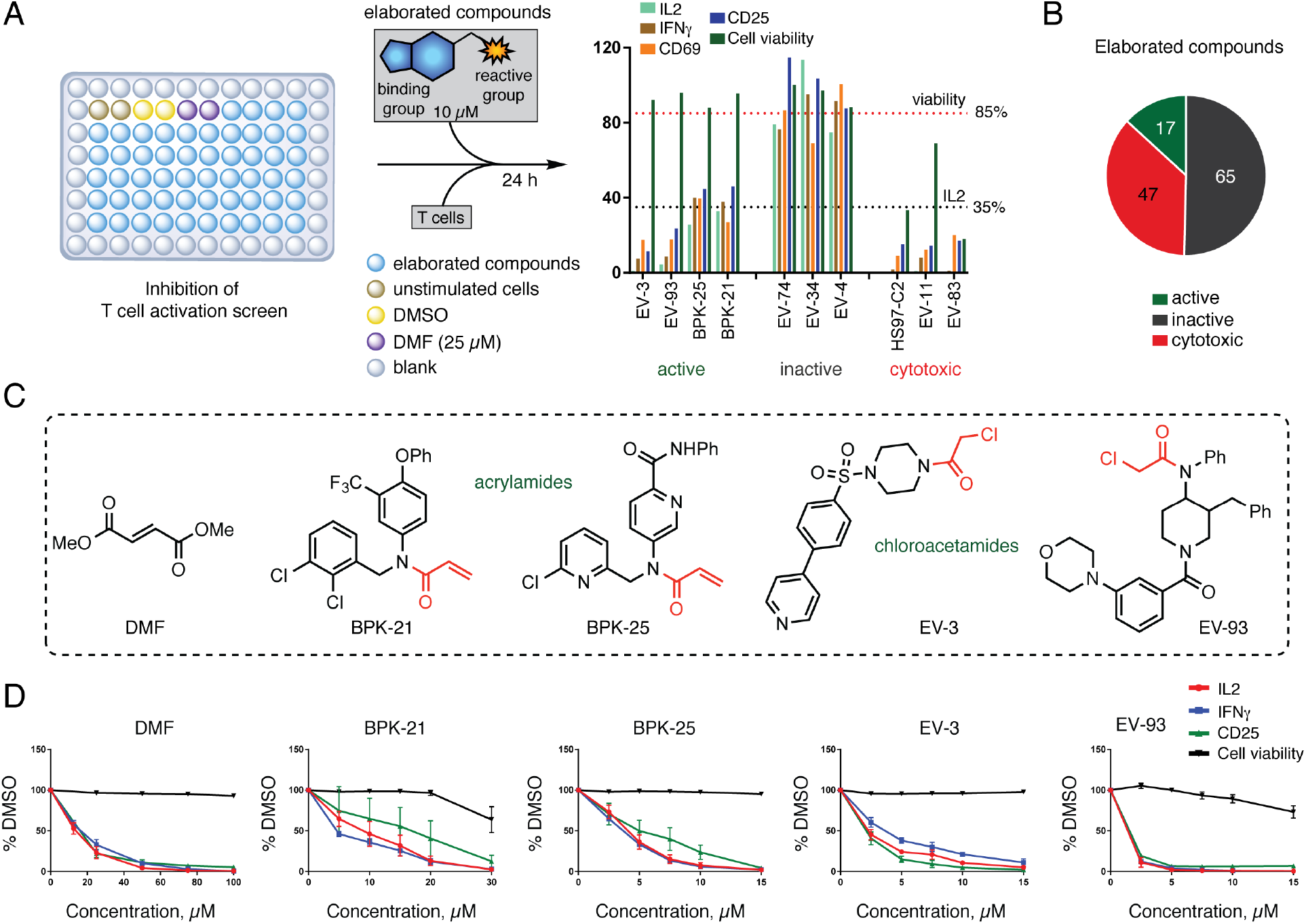
Multidimensional screen identifies elaborated electrophilic compounds that suppress T cell activation. (A) Workflow for T cell activation screen. Primary human T cells were treated with a focused library of elaborated electrophilic compounds (10 *μ*M; see **Table S12** for structures of compounds), a positive control immunosuppressive compound (DMF, 50 *μ*M), or DMSO under TCR-stimulating conditions in 96-well plates pre-coated with 5 *μ*g/mL αCD3 and 2 *μ*g/mL αCD28 for 24 h. T cell activation was measured using a combination of markers, including IL2 and IFNγ secretion, as well as surface expression of CD25 and CD69. T cell viability was measured by flow cytometry using a Fixable Near-IR LIVE/DEAD™ Dead Cell Stain. Compounds were considered as active hits if they reduced IL2 cytokine production by >65% with <15% reduction in T cell viability in the cytotoxicity assay compared to DMSO control. Representative primary screen results are shown for active hit compounds, inactive compounds, and cytotoxic compounds. (B) Screening results for elaborated electrophilic compounds. (C) Structures of active hit compounds selected for follow-up studies, including two acrylamides (BPK-21, BPK-25), two α-chloroacetamides (EV-3, EV-93), and DMF as a positive control. (D) T cell activation and cytotoxicity profiles for selected active hit compounds. Data are presented as the mean percentage of DMSO-treated control cells ± SEM; n = 3/group.

### Chemical proteomic analysis of electrophilic compounds that suppress T cell activation

We next mapped the protein targets of active compounds in primary human T cells by ABPP. Cysteine reactivity profiles were acquired for T cells treated with active compounds (10-20 μM for BKP-21/25 and EV-3/93; 50 μM for DMF) for 3 h. On average, 10,000-12,000 cysteines were quantified for each compound (aggregated across at least 6 biological replicates, where we required that a cysteine was quantified in at least two replicates for interpretation), of which only a modest fraction (0.2-1.0%) was substantially altered in reactivity by compound treatment s(R_DMSO/compound_ ≥ 4) (Figures 5A and **S4B** and **Table S14**). Each active compound engaged a markedly distinct set of cysteines in largely non-overlapping proteins (Figure 5A, B) that originated from diverse structural and functional classes (Figure 5C and **Table S15**) and included several immune-relevant proteins (Figure 5B). Most of the protein targets were engaged site-specifically in that only one of multiple quantified cysteines in these proteins was sensitive to the active compounds (**Figure S4C**).

**Figure 5.**
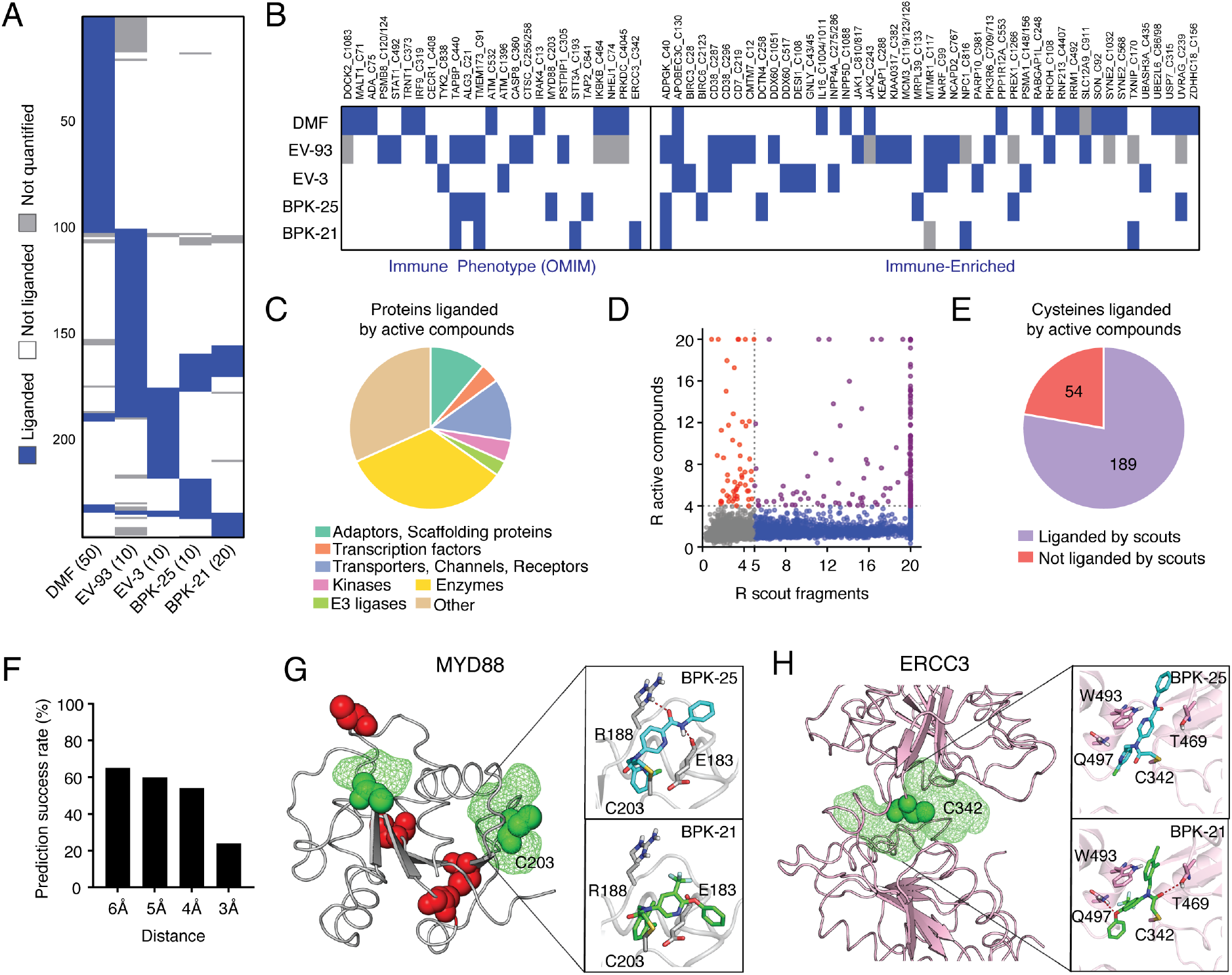
Cysteines liganded by active compounds in human T cells. (A) Heatmap showing liganded cysteine profiles for active compounds in primary human T cells (treated with the indicated concentrations of compounds (μM) for 3 h followed by ABPP). Cysteines quantified for at least two active compounds with R values ≥4 (DMSO/compound) for at least one of them are shown. Results are obtained by combining isoTOP-ABPP and TMT-ABPP data for both soluble and particulate proteomic fractions. R values within each experimental treatment group were derived from 3-6 independent isoTOP-ABPP experiments and 2-3 independent TMT-ABPP experiments (6 TMT channels). A cysteine was required to be quantified in at least two experiments for each proteomic fraction to be reported (See Supplementary Methods for details). (B) Heatmap showing cysteines liganded by active compounds that are found in immune-relevant proteins. (C) Distribution of protein classes containing cysteines liganded by active compounds. (D, E) Comparison of cysteines liganded by active compounds versus scout fragments in human T cells, as displayed in correlation plot (D) and pie chart (E) analyses. In (D), cysteines liganded by both active compounds and scout fragments, only by active compounds, and only by scout fragments are showing in purple, red, and blue, respectively. In (E), cysteines liganded by active compounds and scout fragments are shown in purple and cysteines only liganded by active compounds are shown in red. (F) Prediction success rate querying for pockets within the indicated distances from cysteines liganded by active compounds. (G) Modeling of active compound interactions with C203 in the TLR domain of MYD88 (4DOM). Predicted pockets highlighted as green mesh. Docking of BPK-21 and BPK-25 interactions with C203 showing preferential liganding with BPK-25 due to predicted hydrogen bonds with E183 and R188 (top), which are not accessible in docked structure of BPK-21 (bottom). (H) Modeling of active compound interactions with C342 in the helicase domain of ERCC3 (low resolution Cryo-EM structure of the protein, PDB 5OF4, 4.4 Å resolution). Docking of BPK-25 (top) and BPK-21 (bottom) interactions with C342 showing preferential liganding with BPK-21 due to predicted hydrogen bonds with T469 and Q497 and π-π interaction with W493, which are less accessible in the docked structure of BPK-25.

The vast majority of cysteines liganded by active compounds (~80%) were also engaged by scout fragments (Figure 5D, E and **Table S16**), underscoring the potential for fragment profiling to discover tractable sites of ligandability across the human proteome that can also be targeted by more elaborated electrophiles with improved potency (low-μM) and credible SARs. In general support of this conclusion, we performed molecular modeling on the structures of proteins containing cysteines targeted by active compounds (**Figure S5A** and **Table S17**), which revealed, in ~60% of the cases, predicted binding pockets within 5 Å of the liganded cysteines (Figures 5F). Docking studies on representative protein targets of the structurally related active compounds BPK-21 and BPK-25 further supported the observed SAR profiles (e.g., selectivity of C203 in MYD88 for BPK-25, C342 in ERCC3 for BPK-21, and lack of observed selectivity for C91 in TMEM173; Figure 5G, H and **S5B**). The liganded cysteine residue C91 in TMEM173, or STING, has been shown to be palmitoylated (Mukai et al., 2016) and targeted by other covalent ligands that antagonize TMEM173-dependent inflammatory cytokine production (Haag et al., 2018). Consistent with this past work, we found that BPK-25 inhibited TMEM173 activation by the cyclic dinucleotide ligand cGAMP in both the human monocyte cell line THP1 and in primary human PBMCs (**Figure S5C-E**). More global analysis of our scout fragment data sets revealed additional liganded cysteines subject to palmitoylation (**Table S18**), suggesting the broader potential to pharmacologically target these dynamic lipid modification sites on proteins by electrophilic compounds.

While we were initially drawn to the limited number of shared sites of engagement as potentially representing a common mechanism of action for active compounds, this model was undermined by several factors. First, the shared cysteines mostly reflected highly ligandable sites that have been observed in previous studies to interact broadly with diverse electrophilic compounds (Backus et al., 2016; Bar-Peled et al., 2017). We therefore questioned whether such cysteines would show selective interaction with active compounds over inactive compounds in the screened library. Second, very few of the shared targets had evidence of immune relevance (**Table S14**) and were instead mostly ubiquitously expressed proteins. The active compounds also variably depleted glutathione (GSH) from T cells, but we excluded GSH reductions as a candidate mechanism because buthionine sulphoximine (BSO), an inhibitor of the GSH biosynthetic enzyme gamma-glutamyl cysteine ligase (GCLC), did not affect T cell activation despite depleting GSH content (Figure 6A), a result that is consistent with previous work showing that genetic deletion of GCLC does not impair T cell activation (Mak et al., 2017). We alternatively hypothesized, based on the diverse target profiles of the active compounds, combined with the greater body of literature documenting various ways that electrophiles can affect immune cell function (Backus et al., 2019; Groeger and Freeman, 2010), that the active compounds acted by distinct mechanisms. In support of this hypothesis, we found that active compounds differentially impacted key transcriptional pathways involved in T cell activation, with DMF, EV-3, and BPK-25 suppressing NF-κB activation, as measured by p65 phosphorylation (Figure 6B), but not affecting NFAT transcriptional activity (Figure 6C), while BPK-21 showed the opposite profile (Figure 6B, C). As noted above, a unique target of BPK-21 was C342 of ERCC3, a cysteine in the active site of this helicase that is also targeted by the electrophilic immunosuppressive natural product triptolide (Titov et al., 2011). Like BPK-21, triptolide has been shown to impair T cell activation (Chang et al., 2001) and block NFAT transcriptional activity, but not NF-κB DNA binding (Qiu et al., 1999). We confirmed using CRISPR/Cas9 technology that disruption of the *ERCC3* gene, but not other representative targets of BPK-21, significantly impaired T cell activation to a similar degree as BPK-21 treatment (Figures 6D, E and **S5F**). These data indicate that BPK-21 likely suppresses T cell activation through blockade of ERCC3 function.

**Figure 6.**
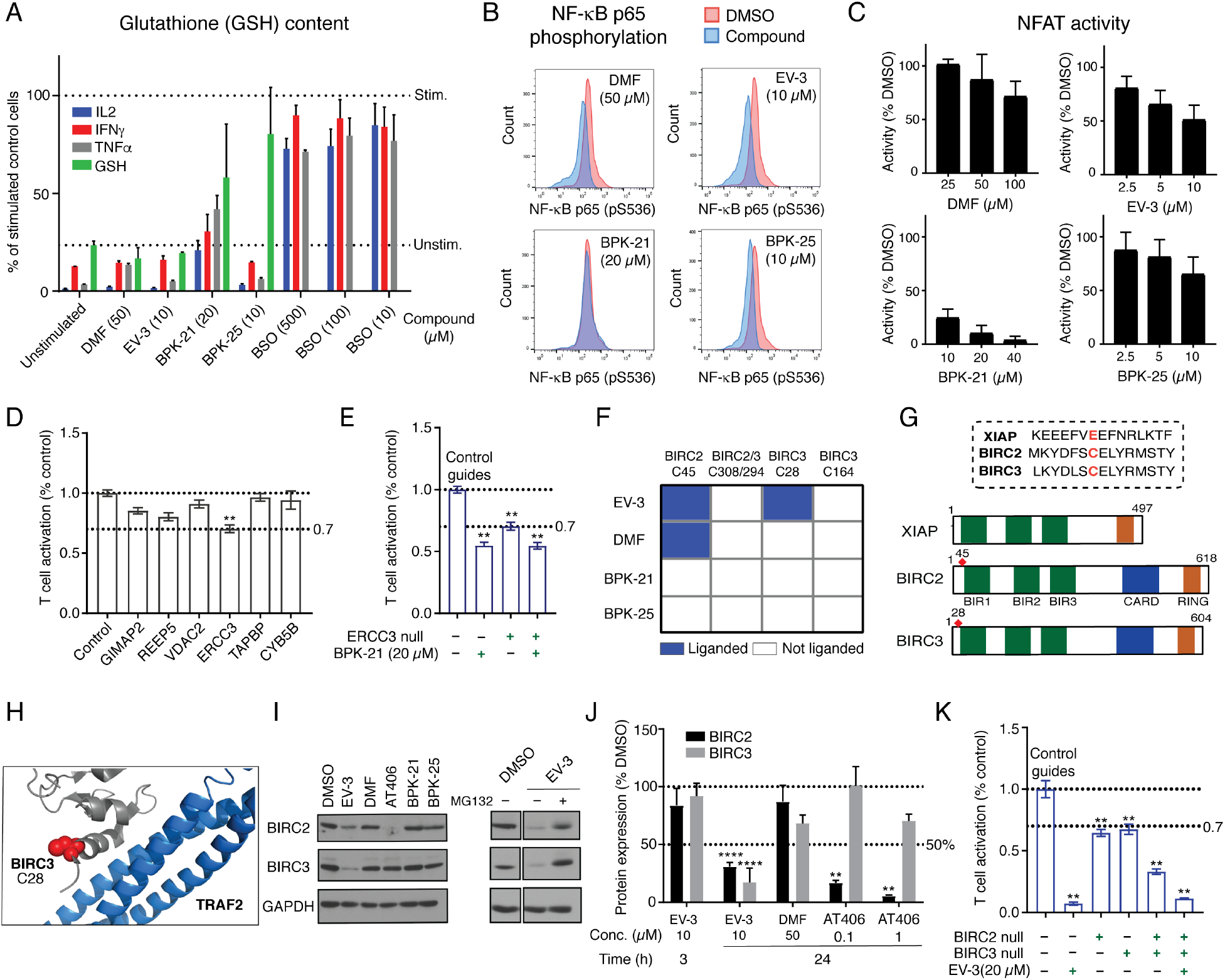
Functional analysis of protein targets of active compounds in human T cells. (A) Effects of active compounds versus the GCLC inhibitor BSO on glutathione (GSH) content and T cell activation parameters. Data are presented as the mean percentage of DMSO-treated control cells ± SEM; n = 3/group. (B) Effects of active compounds on NF-κB activity. Flow cytometry analysis of phosphorylation (S536) of p65 (or RELA) in T cells treated with active compounds at the indicated concentrations (24 h). Plots show p65 (pS536) content in DMSO versus compound-treated cells. Data are from a single experiment representative of at least two independent biological experiments. (C) Effects of active compounds on NFAT activity. Jurkat-Lucia™ NFAT cells were stimulated with PMA (50 ng/mL) and ionomycin (3 μg/mL) in the presence of the indicated concentrations of active compounds, and NFAT transcriptional activity was determined by the levels of Lucia luciferase measured with QUANTI-Luc™ detection reagent in comparison to DMSO-treated control cells. Data are presented as the mean percentage of DMSO-treated control cells ± SEM; n = 2-5/group. (D) Effect of genetic disruption of representative targets of active compound BPK-21 by CRISPR/Cas9 genome editing on T cell activation. Target disruption was considered to have an effect if T cell activation was suppressed >33% with a p value < 0.01. Data are presented as the mean percentage of control guide sgRNA-treated control cells ± SEM; n = 6/group. **, p < 0.01 by two-tailed unpaired *t* test with Welch’s correction compared to control guides. (E) Genetic disruption of ERCC3 and BPK-21 treatment produce similar degrees of blockade of T cell activation. Data are presented as the mean percentage of control sgRNA-treated control cells ± SEM; n = 6/group. **, p < 0.01 by two-tailed unpaired *t* test with Welch’s correction compared to control guides. (F) Active compound EV-3 engages C45 and C28 of BIRC2 and BIRC3, respectively. Heat map showing cysteines liganded by active compounds in BIRC2 and BIRC3. (G) Domain maps for BIRC2, BIRC3, and related protein XIAP, highlighting location of EV-3-sensitive cysteines in BIRC2/3. (H) Location of EV-3-sensitive cysteine C28 in structure of a BIRC3-TRAF2 protein complex (PDB: 3M0A). (I, J) EV-3 causes loss of BIRC2 and BIRC3 in human T cells. (I) Left panels, western blots showing reductions in BIRC2 and BIRC3 content in human T cells treated with EV-3 (10 *μ*M), but not other active compounds (DMF (50 *μ*M), BPK-21 (20 *μ*M), and BPK-25 (10 *μ*M)). The BIR3 domain ligand AT406 (1 *μ*M) was also included for comparison and found to cause loss of BIRC2, but not BIRC3. Right panels, western blots showing that the proteasome inhibitor MG132 (10 *μ*M) blocks EV-3-induced loss of BIRC2 and BIRC3. All treatments were for 24 h. (J) Bar graph representation of western blot data shown in (I), where protein band intensity calculations were made using ImageJ software and represent the mean percentage of DMSO treated control ± SEM; n = 2-5 independent experiments per group. **p < 0.01; ****, p < 0.0001 by two-tailed unpaired *t* test with Welch’s correction compared to DMSO-treated control cells. (K) Effect of genetic disruption of BIRC2, BIRC3, or BIRC2 and BIRC3, by CRISPR/Cas9 genome editing on T cell activation. Target disruption was considered to have an effect if T cell activation was suppressed >33% with a p value < 0.01. EV-3 treatment in BIRC2/BIRC3-disrupted T cells is shown for comparison. Data are presented as the mean percentage of control guide sgRNA-treated control cells ± SEM; n = 6/group. **, p < 0.01 by two-tailed unpaired *t* test with Welch’s correction compared to control guides.

### Electrophilic compound-dependent degradation of BIRC2 and BIRC3

The NF-κB pathway is known to be regulated, both positively and negatively, by reactive oxygen species (ROS) (Morgan and Liu, 2011) and electrophilic compounds (Blewett et al., 2016; Kastrati et al., 2016; Khoo et al., 2018), and consistent with this past work, we discovered cysteines throughout this pathway that showed sensitivity to scout fragments and/or elaborated active compounds (Figure 3A). In reviewing these cysteines, C28 in BIRC3 was noted as a unique target of EV-3 compared to other active compounds (Figures 5B and 6F), and the corresponding cysteine (C45) in BIRC2 was also engaged by EV-3, as well as by DMF, but not other active compounds (Figures 5B, 6F, and **S6A**). Other quantified cysteines in BIRC2 and BIRC3 were not affected by EV-3 treatment (Figure 6F). These proteins, also referred to as cellular inhibitor of apoptosis proteins C-IAP1 and C-IAP2, respectively, regulate both canonical and non-canonical NF-κB activation (Gyrd-Hansen and Meier, 2010) through ubiquitination of diverse substrates (Figure 3A) (Bertrand et al., 2011; Hu et al., 2006; Samuel et al., 2006; Yang et al., 2016; Zarnegar et al., 2008). C28 of BIRC3 (and C45 of BIRC2) is located in close proximity to the BIR1 domain, which interacts with TRAF2 (Figure 6G, H) to facilitate recruitment to the TNF receptor. This interaction has been suggested to stabilize BIRC2, preventing its autoubiquitination and subsequent degradation (Csomos et al., 2009), and mutations in the BIR1 domain of BIRC2/3 (Samuel et al., 2006; Zheng et al., 2010) have been found to abolish or significantly diminish interactions with TRAF2, but, to our knowledge, chemical probes targeting this region of BIRC2/3 have not yet been described.

We found that treatment of human T cells with EV-3, but not other active compounds, including DMF, led to the concentration-dependent (**Figure S6B**), time-dependent (**Figure S6C**), and proteasome-sensitive (Figure 6I) degradation of both BIRC2 and BIRC3. This profile differed from the described Smac mimetic inhibitor AT406, which targets the BIR3 domain (Cai et al., 2011) and selectively promoted the degradation of BIRC2, but not BIRC3 (Figure 6I, J). EV-3 caused minimal changes in mRNA content for BIRC2 or BIRC3 (**Figure S6D**), supporting a direct effect on protein stability in T cells. Finally, genetic disruption of BIRC2 and BIRC3, either independently or in combination (BIRC2/BIRC3-null), using CRISPR/Cas-9 technology impaired T cell activation, with the combined disruption producing a more substantial effect (Figure 6K and **Figure S6E**). Treatment with EV-3 further decreased T cell activation in BIRC2/BIRC3-null cells, indicating that the genetic inactivation of BIRC2 and BIRC3 by CRISPR/Cas-9 may be incomplete in the T cell population (e.g., heterozygotic inactivation in a subset of cells) or additional targets of EV-3 contribute to its full suppressive effects in T cells. We evaluated several additional targets of EV-3 by CRISPR/Cas-9 disruption, but none were found to substantially impair T cell activation (**Figure S6F**).

### The acrylamide BPK-25 promotes degradation of the NuRD complex

As noted previously, the structurally related acrylamides BPK-21 and BPK-25 both suppress T cell activation, but by apparently different mechanisms involving impairment of NFAT transcriptional activity and NF-κB signaling respectively (Figure 6B, C). A survey of the cysteines preferentially engaged by BPK-25 over BPK-21 did not reveal obvious candidate proteins within the NF-κB pathway as potential targets of relevance for the former compound (**Table S13**). Motivated by the finding that EV-3 promoted the degradation of immunologically relevant BIRC proteins (Figure 6I, J), we performed an expression-based proteomic analysis of primary human T cells treated with BPK-25 or other active compounds (Figure 7A). This study revealed that BPK-25, but not other active compounds, including BPK-21, led to the striking and selective reduction of several proteins in the Nucleasome Remodeling and Deacetylation Complex (NuRD), including a >50% decrease in GATAD2B, MBD2, MBD3, and MTA2, as well as directionally similar losses in GATAD2A, HDAC1, HDAC2, MTA1, and RBBP4 (Figure 7B, C and **Tables S18** and **S19**). Only two other proteins across the >3000 quantified proteins in our proteomic experiments showed substantial (>50%) reductions in BPK-25-treated cells (FAM213B and HLA-F), and these proteins were also affected by other active compounds (Figure 7C). BPK-25-mediated reductions in NuRD complex proteins were time-dependent (Figure 7D) and blocked by treatment with the proteasome inhibitor MG132 (Figure 7C). The reductions in NuRD complex proteins were not accompanied by corresponding changes in mRNA expression for the proteins (Figure 7E), supporting that BPK-25 lowers NuRD complex proteins by a post-transcriptional mechanism. While our cysteine reactivity profiles indicated an ~50% reduction in C10/11 and C359 in MBD2, C266 in MBD3, C308 and C423 of GATAD2B in BPK-25-treated T cells (but not T cells treated with BPK-21 or EV-3 (Figure 7F); 3 h treatment), these partial changes were difficult to assign with confidence as being decreases in cysteine reactivity versus protein expression. We also note that DMF decreased the apparent reactivity of C308 of GATAD2B (Figure 7F), despite not leading to NuRD complex degradation (Figure 7C). Whether one or more of the apparently BPK-25-sensitive cysteines in NuRD complex proteins is responsible for mediating the loss of this complex in T cells remains to be determined. Despite this mechanistic uncertainty, our studies have uncovered a remarkably selective, electrophile-mediated process for degradation of a key transcriptional regulatory complex in primary human T cells. Considering that HDAC inhibitors have been previously shown to block T cell activation (Takahashi et al., 1996), and HDACs can also support NF-κB function (Jung et al., 2009; Kumar et al., 2017; Wagner et al., 2015), it is possible that the BPK-25-mediated loss of the NuRD complex is relevant to the T cell-suppressive activity of this compound.

**Figure 7.**
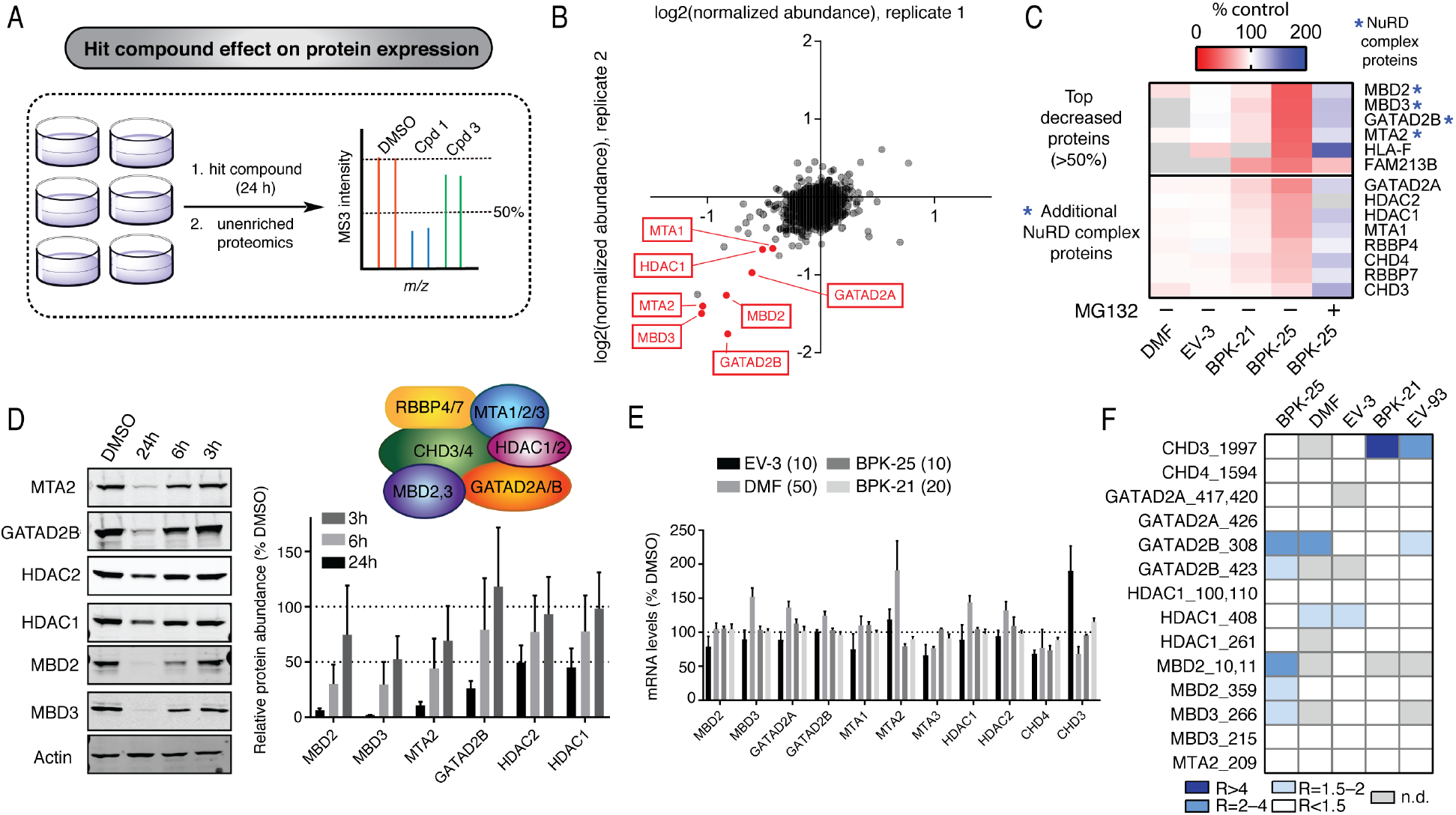
Active compound BPK-25 promotes degradation of NuRD complex proteins in human T cells. (A) Experimental workflow for quantitative proteomic experiments evaluating protein abundance changes caused by active compound treatment in primary human T cells: 1) T cells were treated with hit compounds (DMF (50 *μ*M), EV-3 (10 *μ*M), BPK-21 (20 *μ*M), BPK-25 (10 *μ*M)) or DMSO control for 24 h; 2) cells were then processed and analyzed by TMT-based quantitative proteomics where a 50% reduction in average peptide signals for a protein were interpreted as a reduction in the quantity of that protein. (B) Scatter plot representation of protein abundance changes caused by BPK-25 (10 *μ*M, 24 h) in two independent replicate experiments, with decreases in NuRD complex proteins highlighted in red. (C) Heatmap of top proteins with decreased abundance in BPK-25-treated T cells showing that the subset of these proteins in the NuRD complex (asterisks) were largely unaltered by other active compounds and blocked in their degradation by co-treatment with the proteasome inhibitor MG132. Additional NuRD complex proteins are also displayed, and most of these proteins also showed evidence of reduced abundance (25–50%) in T cells treated with BPK-25. (D) Time-dependent reductions in NuRD complex proteins in human T cells treated with BPK-25 (10 *μ*M). Left, representative western blots. Right, quantification of changes in protein expression. Western blot protein band intensity calculations were made using ImageJ software and represent the mean percentage of DMSO treated control ± SD; n = 2. (E) Active compound treatment did not, in general, reduce the mRNA content for NuRD complex proteins. mRNA quantities were measured by RNAseq. Data are presented as the mean percentage of DMSO-treated control cells ± SEM; n = 3/group. (F) Changes in cysteine reactivity for NuRD complex proteins in human T cells treated with BPK-25 and other active compounds.

## Discussion

In this manuscript, we describe a chemical proteomic strategy aimed at addressing a paradoxical challenge at the interface of immunology and pharmacology, namely that, despite major advances in our understanding of the genetic basis of human immune disorders, the majority of immune-relevant proteins lack chemical probes to facilitate their functional characterization and streamline future drug development programs. By generating and integrating global maps of cysteine reactivity and ligandability in primary human T cells, studied in both quiescent and activated states, we provide a resource that greatly expands the landscape of electrophilic compound interactions for diverse immune-relevant proteins. We show how cysteine ligandability maps can be further refined by phenotypic screening to illuminate tractable and functional sites of electrophilic compound action on immune-relevant proteins, including those historically considered challenging to target with small molecules (e.g., adaptor proteins, transcription factors; Figure 5C). We note that, even for traditionally druggable protein classes like kinases, covalent ligand engagement frequently occurred at sites that were structurally distinct from the ATP-binding pocket, underscoring the potential for electrophilic small molecules to target non-canonical sites on proteins. The discovery of non-active site ligands may prove useful for parsing the catalytic versus non-catalytic (e.g., scaffolding) functions of proteins in immune cells.

That some of the active compound-cysteine interactions led to protein degradation without requiring a separate E3 ligase-directing ligand (e.g., EV-3-mediated degradation of BIRC2/3) underscores the potential for covalent modification by small molecules to directly affect protein stability in cells (Foulkes et al., 2018; Jones, 2018; Schreiber, 2019; Yang et al., 2019). We do not yet understand how EV-3 promotes BIRC2 and BIRC3 degradation, but this could involve disruption of protein-protein interactions proximal to the compound-modified cysteines in these proteins. In this model, electrophilic compounds engaging the same cysteine on a protein target may produce different functional outcomes depending on whether they perturb the interactions with partner proteins. This may explain why a small electrophile like DMF failed to promote BIRC2 degradation, despite engaging C45 in this protein. It is also possible that E3 ligases like BIRC2 and BIRC3, which can undergo auto-ubiquitination (Dueber et al., 2011; Varfolomeev et al., 2007), are especially susceptible to direct compound-mediated degradation. Arguing for the broader potential of this pharmacological effect, however, we also observed electrophilic compound-induced degradation of the transcriptional regulatory NuRD complex in T cells, albeit by a yet undefined mechanism. More generally, we should emphasize that some of the cysteine ligandability events mapped in this study, including those showing good tractability, may fail to produce direct functional effects on proteins. Such so-called “silent” compounds still have the potential to be converted into heterobifunctional small-molecule degraders of proteins, an approach that has been successfully used with other covalent ligands (Buckley et al., 2015; Burslem et al., 2018; Tovell et al., 2019).

One of the goals of this study was to assess the potential of electrophilic fragment screening to identify novel sites of druggability in primary human immune cells. Among the >3400 cysteines liganded by scout fragments in human T cells (from >18000 total quantified cysteines) were not only several known sites of electrophilic drug action (**Table S8**), but also cysteines in diverse protein components of key immune signaling pathways (e.g., NF-κB, TLR, TCR; Figure 3A) and in transcription factors involved in immune cell lineage commitment, including CD8^+^ cytotoxic T cell development and differentiation (e.g., RunX3, EOMES) (Wang and Bosselut, 2009), late thymocyte development (e.g., THEMIS) (Lesourne et al., 2009), and Th1 (e.g., STAT1, STAT4) (Oestreich and Weinmann, 2012), Th2 (e.g., IRF4) (Zhu, 2010), Th17 (e.g., IRF4, STAT3) (Ivanov et al., 2007), and Treg (e.g., FOXP3) (Fontenot et al., 2003) differentiation. The number of cysteines engaged in T cells by more elaborated electrophilic compounds (Figure 5) further indicates that the global ligandability maps furnished by scout fragments canvas a substantial number of tractable sites for covalent ligand development in immune-relevant proteins. We should emphasize that only ~130 elaborated electrophilic compounds were screened (and the target landscape only analyzed for four of these compounds) in human T cells, and it is therefore likely that many additional cysteines found only to engage scout fragments herein will show evidence of tractability as they are assayed against a greater number of structurally diverse electrophiles. We envision that our fragment ligandability maps may even be used to focus such future chemical proteomic screens, possibly performed in a targeted manner (Uzozie and Aebersold, 2018), on individual cysteines in proteins of high immunological relevance, such as the transcription factors noted above. We also anticipate that structure-guided approaches may facilitate optimization of hit compound-protein interactions, but caution that only a minor fraction of the ligandability events mapped herein (~25%) were found in proteins or domains in proteins for which high-resolution structural information is available (**Figure S5A**). This outcome emphasizes the large gap still remaining in our understanding of the three-dimensional structures of full-length human proteins.

Cysteine is a uniquely nucleophilic amino acid that performs diverse biochemical functions in catalysis, redox regulation, metal coordination, and signaling via post-translational modification (Giles et al., 2003a; Giles et al., 2003b; Jacob et al., 2003; Weerapana et al., 2010), each of which may be affected by changes in the redox environment, metabolism, and signaling pathways associated with T cell activation. The extent to which such state-dependent changes in cysteine reactivity may be exploited for the development of chemical probes and possibly drugs that have a more selective effect on the activity of T cells remains an open and exciting question for future inquiry. T cell-restricted pharmacology may also emerge from targeting proteins selectively expressed in this immune cell type, and we note, in this regard, the rich content of ligandable cysteines in immune-relevant proteins discovered in our studies. The presence of ligandable cysteines in proteins historically considered challenging to target with small molecules, as well as our demonstration that electrophilic compounds engaging these cysteines can directly affect the functions and/or stability of immune-relevant proteins in cells, points to the broad potential for covalent small molecules to serve as probes and future drugs for modulating diverse immunological processes.

## Supporting information

Supplementary Table - RNAseq_data

## AUTHOR CONTRIBUTIONS

Conceptualization, E.V.V., J.R.T., and B.F.C.; Methodology, E.V.V., D.C.L., M.M.B., J.R.T., and B.F.C.; Formal analysis, E.V.V., R.M.S., Y.W., and B.F.C.; Investigation, E.V.V., D.C.L., Y.Y., V.M.C., D.R., E.K.K., M.R.L., S.Y., M.M.D., N.N., E.C., and L.L.; Resources, E.V.V., G.M.S., R.M.S., and K.M.L.; Data Curation, E.V.V., R.M.S., Y.W., M.N.S., and B.F.C.; Writing – Original Draft, E.V.V.; Writing – Review and Editing, E.V.V., B.F.C.; Visualization: E.V.V., R.M.S., and B.F.C.; Supervision: E.V.V., J.R.T., and B.F.C.

## ACKNOWLEDGEMENTS

We thank D. Tallman for technical assistance, Bingwen Lu for help with Table S3, and M. Niphakis for helpful discussions. This work was supported by NIH CA231991 (B.F.C.), CA212467 (V.M.C.), CA211526 (M.M.D.), GM069832 (S.F.), a Clinical Translational Science Award (UL1 TR001114), and a Life Sciences Research Foundation Fellowship to E.V.V.. M.N.S. was supported by funding from NIH-NCI CCSG: P30 014195, R01 GM102491-07, and the Helmsley Trust.

