## Supplementary Information for "An activity-guided map of electrophile-cysteine interactions in primary human immune cells"

### (A) Supplementary Figures

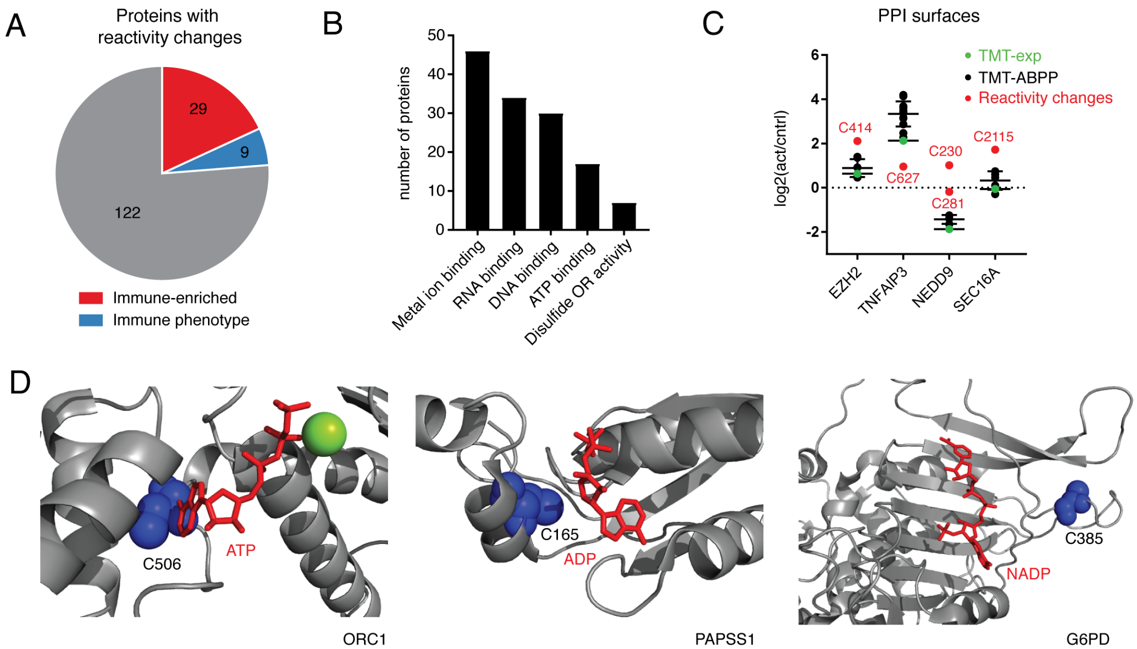

**Figure S1. Chemical proteomic map of cysteine reactivity changes in activated T cells. Related to Figure 1.**
(A) Fraction of proteins with human genetics-based immune phenotypes and immune-enriched protein expression from total proteins showing cysteine reactivity changes in activated T cells.
(B) Bar graph representation of proteins with cysteine reactivity changes organized by molecular function GO term enrichment.
(C) Representative cysteine reactivity changes in activated human T cells for cysteines at protein-protein interaction (PPI) surfaces.
(D) Representative cysteine reactivity changes in activated human T cells for cysteines in cofactor/metabolite-binding sites; x-ray crystal structure of the active form of human origin recognition complex subunit 1 (ORC1) in complex with ATP and $Mg^{2+}$ (PDB: 5UJ7) with the reactivity-changing cysteine C506 highlighted in blue (left image); x-ray crystal structure of human 3’-phosphoadenosine-5’-phosphosulfate synthetase 1 (PAPSS1) in complex with ADP (PDB: 1X6V) with the reactivity-changing cysteine C165 highlighted in blue (middle image); x-ray crystal structure of human glucose-6-phosphate dehydrogenase (G6PD) in complex with $NADP^{+}$ (PDB: 2BH9) with the reactivity-changing cysteine C385 highlighted in blue (right image).

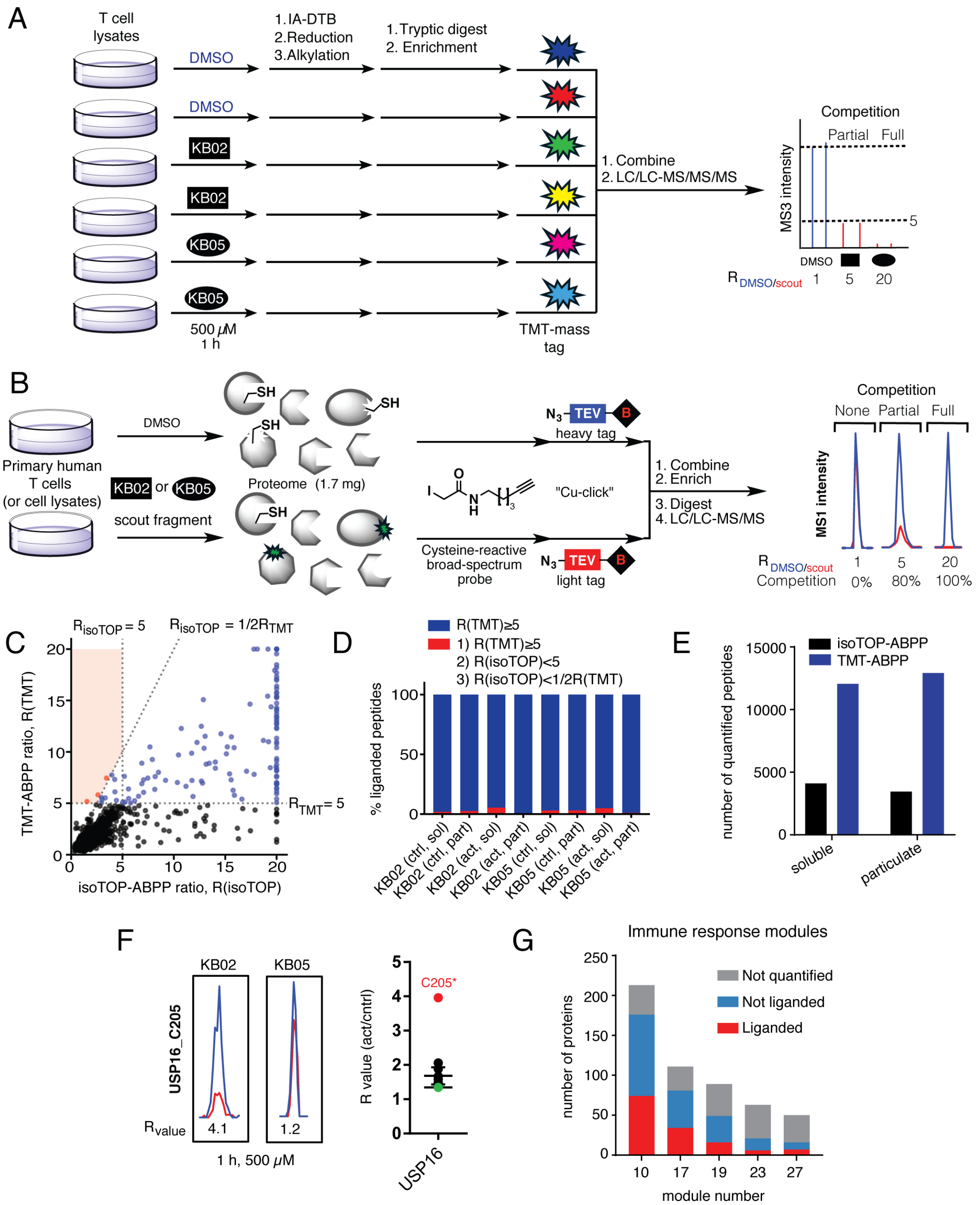

**Figure S2. Chemical proteomic map of fragment electrophile-cysteine interactions in human T cells. Related to Figure 2.**

(A) Extended experimental workflow for chemical proteomic experiments measuring scout fragment electophile effects on cysteine reactivity in primary human T cells using isobaric tandem mass tags for mass differentiation and MS3-based quantification (TMT-ABPP).

(B) Extended experimental workflow for chemical proteomic experiments measuring scout fragment electophile effects on cysteine reactivity in primary human T cells using clickable, TEV protease-sensitive, isotopically labeled tags for mass differentiation and MS1-based quantification (isoTOP-ABPP).

(C, D) Comparison of R-values from isoTOP-ABPP and TMT-ABPP experiments, as displayed in correlation plot (C) and bar graph (D) analyses. Results represent mean R-values derived from 3-5 independent isoTOP-ABPP experiments and 4 independent TMT-ABPP experiments (6 TMT channels per experiment) for each

compound treatment and proteomic fraction (2-5 biological donors for each method). A cysteine was required to be quantified in at least two experiments for each compound treatment or proteomic fraction group. KB02-treated soluble proteome samples are used as an example for the correlation plot. For (C) and (D), data points with R(TMT-ABPP) ≥5 are colored in blue with the exception of rare outlier cases that also exhibited R(isoTOP-ABPP) <5 and R(isoTOP-ABPP)<1/2R(TMT), which are colored in red.
(E) Bar graphs showing the total number of quantified peptides in isoTOP-ABPP and TMT-ABPP experiments for soluble and particulate proteomic fractions of primary human T cells. Results represent a combination of data from KB02 and KB05 experminents (500 μM, 1 h) with both control and activated T cells. R-values within each experimental treatment group were derived from 3-5 independent isoTOP-ABPP experiments and 4 independent TMT-ABPP experiments. A cysteine was required to be quantified in at least two experiments for each compound treatment or proteomic fraction group to be reported.
(F) Left: MS1 signal intensities for C205 in USP16 in KB02- and KB05-treated T cell proteome; right: C205 of USP16 reactivity change in activated human T cells.
(G) Fraction of liganded proteins from total proteins found in previously described immune-enriched modules (Rieckmann et al., 2017).

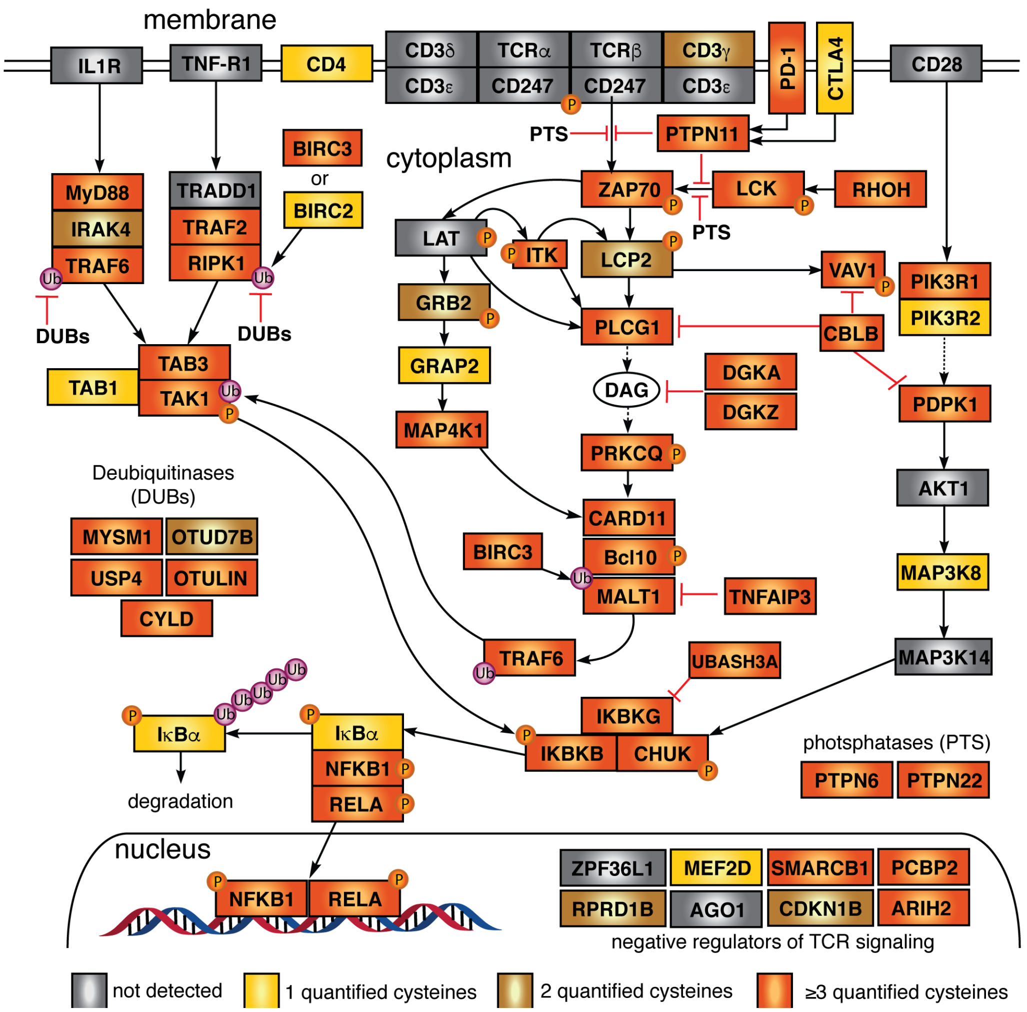

**Figure S3**. **Diagram of immune signaling pathways, related to Figure 3.** Cysteine quantification events in experiments with scout fragments and elaborated compounds with the following colors used to mark proteins with no (grey), one (yellow), two (brown), or > two (orange) quantified cysteines.

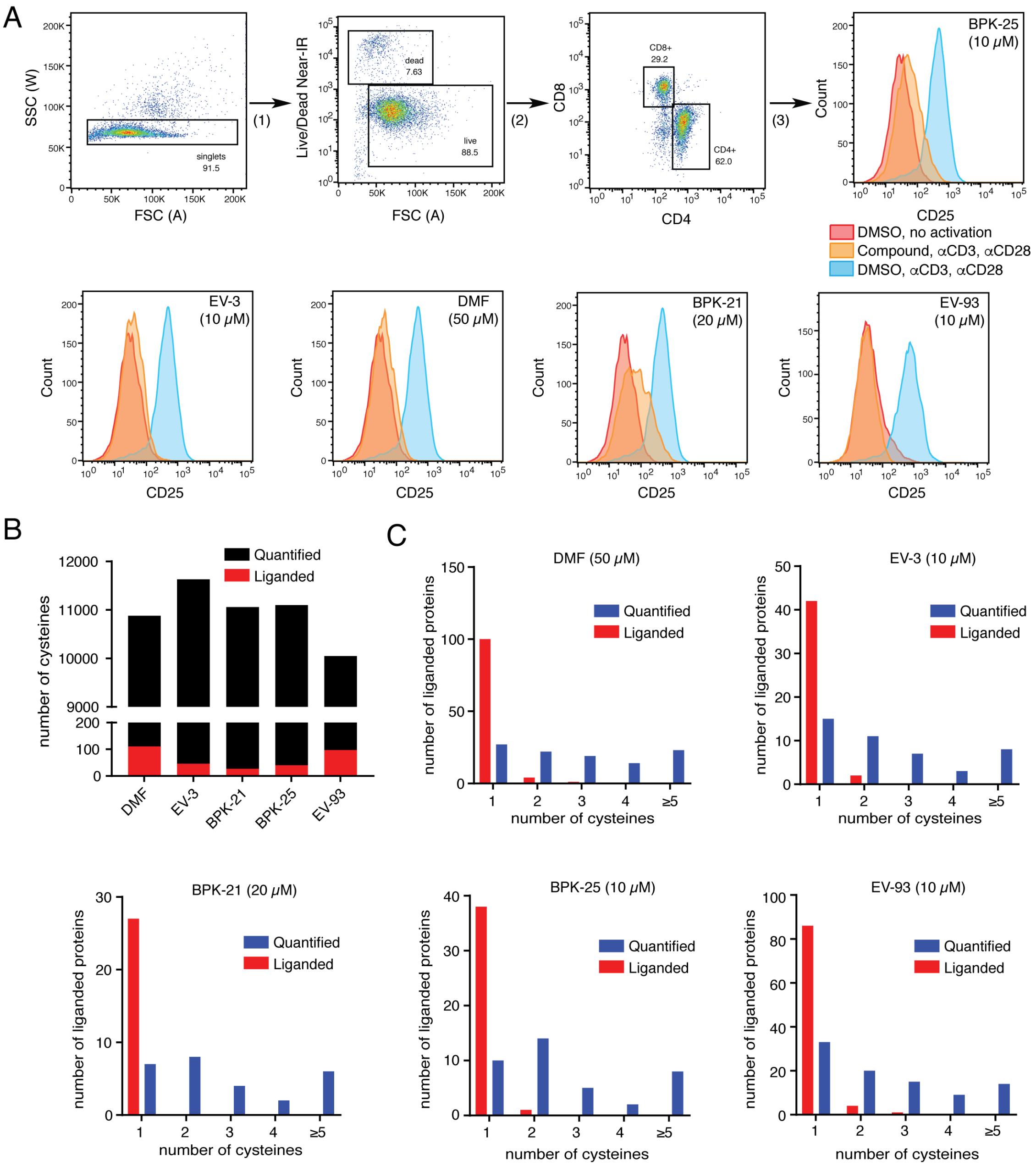

**Figure S4. Further analysis of active compound effects in human T cells. Related to Figures 4 and 5.** (A) Flow cytometry analysis of T cell activation. T cell activation was monitored by measuring CD25 expression levels after overnight stimulation with αCD3 and αCD28 antibodies in the presence of active compounds. The following FACS gating strategy was used: 1) Doublet discrimination was achieved by plotting forward scatter area (x-axis) versus side scatter width (y-axis) and selecting the area containing singlets for further analysis; 2) Live cells were selected by plotting forward scatter area (x-axis) versus Live/Dead Near-IR channel (y-axis, area, logarithmic scale) and gating on the negative cell population; 3) $CD4^+$ cells were differentiated from $CD8^+$ cells by plotting Pacific Blue channel (x-axis, area, logarithmic scale, PB-CD4 antibody) versus PerCP-Cy5.5 channel (y-axis, area, logarithmic scale, PerCP-Cy5.5-CD8a antibody). $CD4^+$ positive, $CD8^+$ negative cells were further analyzed for CD25 expression levels by measuring mean fluorescence intensity of the PE-channel (PE-CD25). Similar results were obtained by gating on the $CD8^+$ positive, $CD4^+$ negative cell population. BPK-25-treated cells are used as an example for the gating strategy. Representative histograms showing CD25 levels in unstimulated T cells (red), stimulated T cells in the

presence of DMSO (light blue), and stimulated T cells in the presence of active compounds (orange) are presented. Data is representative of three biological replicates used in **Figure 4D**.
(B) Bar graph showing the total number of quantified (black) and liganded (R≥4, red) cysteines in cells treated with active compounds. Results are obtained by combining isoTOP-ABPP and TMT-ABPP data for both soluble and particulate proteomic fractions. R-values within each experimental treatment group were derived from 3-6 independent isoTOP-ABPP experiments and 2-3 independent TMT-ABPP experiments (6 TMT channels). A cysteine was required to be quantified in at least two experiments for each proteomic fraction to be reported.
(C) Bar graphs showing the total number of liganded proteins as relates to the corresponding number of quantified (blue) and liganded (R≥4, red) cysteines per protein. Results are obtained by combining isoTOP-ABPP and TMT-ABPP data for both soluble and particulate proteomic fractions for each compound treatment. A cysteine was required to be quantified in at least two experiments for each proteomic fraction to be reported.

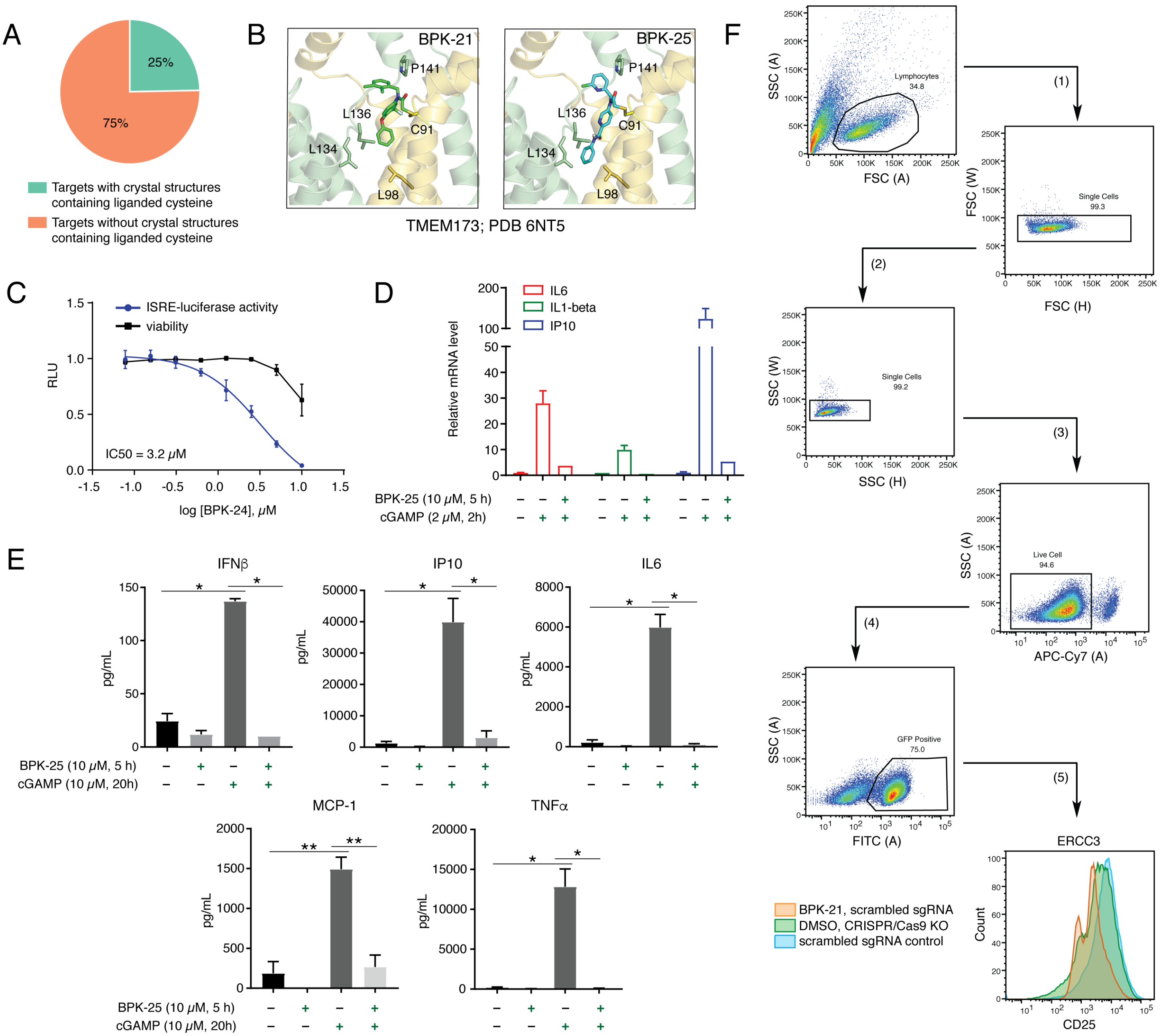

**Figure S5. Functional analysis of protein targets of active compounds in human T cells. Related to Figure 5.**

(A) Fraction of protein targets of active compounds with available crystal structures containing the corresponding liganded cysteine residues.

(B) Modeling of BPK-21 and BPK-25 interactions with C91 of stimulator of interferon genes protein (STING, or TMEM173).

(C) IRF response of THP1-Lucia™ ISG cells. THP-1 Lucia ISG cells were treated with DMSO or BPK-25 in the presence of viral dsDNA (2 µg/mL) for 24h and the levels of IRF-induced Lucia luciferase were determined using QUANTI-Luc™. Cell viability was measured using CellTiter-Glo™ assay. Luminescence signals for test samples were normalized to DMSO-treated samples and reported as relative light units (RLU) ± SEM; n = 3/group.

(D) Effect of BPK-25 treatment on gene expression related to TMEM173/STING pathway activation in PBMCs. PBMCs were treated with DMSO or BPK-25 (10 μM) for 5 h and stimulated with cGAMP (2 μM) for 2 h. Relative expression of IL-6, IL-1β, and IP10 (CXCL10) genes was measured by qPCR and normalized to actin.
(E) Effect of BPK-25 treatment on secretion of cytokines related to TMEM173/STING pathway activation in PBMCs. PBMCs were treated with DMSO or BPK-25 (10 μM) for 5 h and stimulated with cGAMP (10 μM) for 20 h. Cytokine levels (IFN-β, IP10 (CXCL10), IL-6, MCP-1, TNFα) were measured using standard ELISA and Bio-Plex protocols (see Supplementary Methods for details). Data are presented as the mean percentage of DMSO-treated control cells ± SEM; n = 3/group. *, $p < 0.05$; **, $p < 0.01$ by two-tailed unpaired *t* test with Welch's correction between indicated groups.
(F) Flow cytometry analysis of T cell activation following ERCC3 gene disruption by CRISPR/Cas9 genome editing. T cells were activated for 2 days prior to Cas9 RNP transfection and were then cultured in IL2 containing RPMI media to return the cells to a quiescent state. Seven days post-transfection, T cells were stimulated overnight with αCD3 and αCD28 antibodies in the presence of DMSO or BPK-21 with the activation monitored by measuring CD25 and CD69 expression levels. The following FACS gating strategy was used: 1) Lymphocyte gating; 2-3) Doublet discrimination by plotting forward scatter height (x-axis) versus forward scatter width (y-axis) and side scatter height (x-axis) versus side scatter width (y-axis); 4) Live cells were selected by plotting APC-Cy7 channel (x-axis, area, logarithmic scale, eBioscience™ Fixable Viability Dye eFluor™ 780) versus side scatter area (y-axis) and gating on the negative cell population; 5) Transfected cells were selected by gating on FITC-positive cell population (GFP-positive cells). GFP-positive cells were further analyzed for CD25 and CD69 expression levels by measuring mean fluorescence intensity of the PE- (PE-CD25) and APC-channels (APC-CD69). Representative histograms showing CD25 levels in stimulated T cells in the presence of scrambled sgRNA control (light blue) or ERCC3 guide RNAs (green) in the presence of DMSO or BPK-21 (orange) are presented. Data is representative of a total of six replicate treatments used in **Figure 6E**.

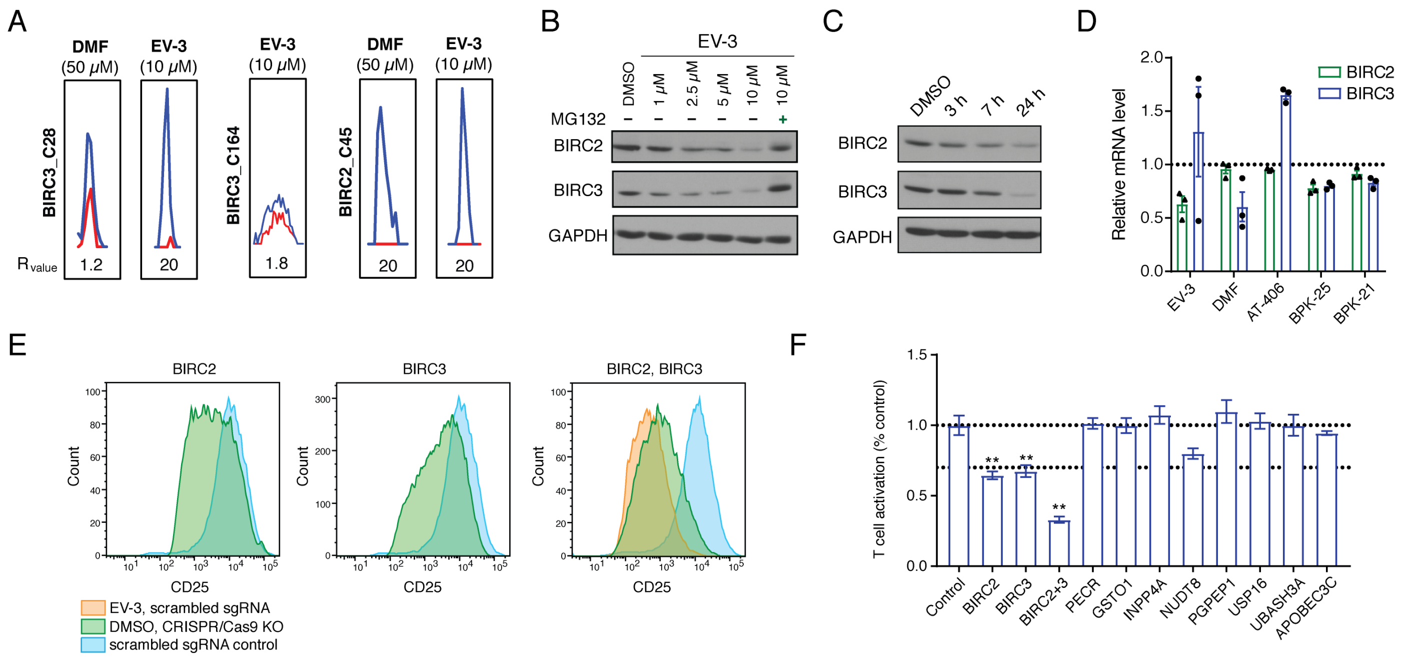

**Figure S6. Functional analysis of protein targets of active compounds in human T cells. Related to Figure 6.**

(A) MS1 signal intensities for BIRC3_C28, BIRC3_C164, and BIRC2_C45 in isoTOP-ABPP experiments of expanded T cells treated with EV-3 (10 μM, 3 h) or DMF (50 μM, 3 h).

(B, C) EV-3 causes loss of BIRC2 and BIRC3 in human T cells. (B) Full western blot from **Figure 6I** (right) showing reductions in BIRC2 and BIRC3 content in human T cells treated with EV-3 (1–10 μM, 24 h) and the blockade of EV-3-induced loss of BIRC2 and BIRC3 by co-treatment with the proteasome inhibitor MG132 (10 μM). (C) Western blot showing time-dependent reductions in BIRC2 and BIRC3 content in human T cells treated with EV-3 (10 μM).

(D) mRNA content for BIRC2 and BIRC3 quantified by RNA-sequencing of T cells treated with EV-3 (10 μM), DMF (50 μM), AT-406 (1 μM), BPK-25 (10 μM), or BPK-21 (20 μM). All treatments were done with T cells for 24 h. Data are presented as the mean percentage of DMSO-treated control cells ± SEM; n = 3/group.

(E) Flow cytometry analysis of T cell activation following BIRC2, BIRC3, or BIRC2 and BIRC3 gene disruption by CRISPR/Cas9 genome editing. T cells were activated for 2 days prior to Cas9 RNP transfection and were then cultured in IL2 containing RPMI media to return the cells to a quiescent state. Seven days post-transfection, T cells were stimulated overnight with αCD3 and αCD28 antibodies in the presence of DMSO or BPK-21 with the activation monitored by measuring CD25 and CD69 expression levels. The same FACS gating strategy as in **Figure S5F** was used. Representative histograms showing CD25 levels in stimulated T cells in the presence of scrambled sgRNA control (light blue) or BIRC2/BIRC3/BIRC2+BIRC3 guide RNAs (green) in the presence of DMSO or EV-3 (orange) are presented. Data is representative of a total of six replicate treatments used in **Figure 6K** and **S6F**.

(F) Effect of genetic disruption of representative targets of active compound EV-3 by CRISPR/Cas9 genome editing on T cell activation. Target disruption was considered to have an effect if T cell activation was suppressed >33% with a p value < 0.01. Data are presented as the mean percentage of control guide sgRNA-treated control cells ± SEM; n = 3/group. **, p < 0.01 by two-tailed unpaired *t* test with Welch's correction compared to control guides.

### (B) Supplementary Data Set Legends

**Supplementary Data Set 1**
**Tab 1.** Table of contents for **Supplementary Dataset 1**.

**Tab 2 (Table S1).** TMT-exp proteomic data showing changes in protein expression in control vs activated primary human T cells.

**Tab 3 (Table S2).** TMT-ABPP proteomic data showing changes in cysteine reactivity in control vs activated primary human T cells.

**Tab 4 (Table S3).** Resource table for determination of immune-relevant genes. Multiple Z-scores were calculated from several transcriptomic tissue profiles, and these Z-scores were summed to generate the value in column H. Genes with summed Z-scores in the top ~10% were considered "immune-enriched", which corresponded to Z scores > 11. Annotation of immune phenotype (column T) was performed by parsing data in the Online Mendelian Inheritance of Man (OMIM) database and querying phenotype titles and descriptions for immune-related substrings ('*immun*', '*inflam*', '*rheum*', '*psoria*', etc), as described in the Supplementary Methods.. In total, there were 2004 immune-enriched genes and 655 genes associated with immune phenotypes which, when combined, result in 2746 total immune-relevant genes (column U).

**Tab 5 (Table S4).** List of proteins with identified cysteine reactivity changes, along with designations of immune-relevance and GO-terms: "RNA-binding", "Metal ion binding", "DNA binding", "disulfide oxidoreductase activity", and "ATP binding". The summary of GO-term analysis is provided in **Figure S1B**.

**Tab 6 (Table S5).** Master table containing total cysteine reactivity proteomic data from combined isoTOP-ABPP and TMT-ABPP experiments with scout fragments ("broad ligandability", KB02 and KB05) and active compounds (DMF, EV-3, EV-93, BPK-21, BPK-25), as well as isoTOP-ABPP hyperreactivity experiments. Values for each treatment group are final average values resulting from at least 2 replicate experiments described in **Supplementary methods**. Maximum values from all treatment groups (Columns J and P) for active compounds and scout fragments are used to determine "ligandability" of a select residue (related to **Figure 2**).

**Tabs 7-8 (Tables S6, S7).** Comparison of isoTOP ABPP and isoTOP-ABPP hyperreactivity R values for activated and control T cells (related to **Figure 2E**).

**Tab 9 (Table S8).** List of liganded cysteines in immune-relevant proteins that are also targeted by other electrophile compounds.

**Tab 10 (Table S9).** Analysis of proteins liganded with scout fragments (**Table S5**) for assignment to T cell proliferation genes (related to **Figure 2J**) (Shifrut et al., 2018).

**Tab 11 (Table S10).** Analsyis of liganded and quantified proteins for assignment to previously described immune-enriched modules (modules 10, 17, 19, 23, 27, related to **Figure S2G**). (Rieckmann et al., 2017).

**Tab 12 (Table S11).** List of immune-enriched transcription factors and adaptor proteins liganded with scout fragments (related to **Figure 3C**). See **Supplementary Methods** for details.

**Tab 13 (Table S12).** Chemical structures of compounds used in the inhibition of T cell activation screen (related to **Figure 4**).

**Tab 14 (Table S13).** Summary of the results of the phenotypic screen for inhibition of T cell activation (related to **Figure 4**).

**Tab 15 (Table S14).** isoTOP-ABPP and TMT-ABPP proteomic data for active compounds (DMF, EV-3, EV-93, BPK-21, BPK-25; related to **Figure 5**). Values for each treatment group are final average values resulting from at least 2 replicate experiments described in **Supplementary methods**. For each compound, maximum values across both methods and proteomic fractions are reported in the Master table (**Tab 6**, **Table S5**).

**Tab 16 (Table S15).** Distribution of protein classes containing cysteines liganded by active compounds.

**Tab 17 (Table S16).** Comparison of isoTOP-ABPP and TMT-ABPP R values for cysteines liganded by active compounds versus scout fragments in human T cells, as displayed in correlation plot (**Figure 5D**) and pie chart (**Figure 5E**) analyses.

**Tab 18 (Table S17).** Prediction of pocket volumes within the indicated distances from cysteines liganded by active compounds (related to **Figure 5F**).

**Tab 19 (Table S18).** Liganded cysteine residues with scout fragments at known sites of palmitoylation (obtained by cross-referencing with SwissPalm proteins and sites).

**Tab 20 (Table S19).** Protein expression changes after BPK-25 (10 μM, 24 h) treatment as determined by TMT-exp (10 replicates from 5 donors, related to **Figure 7B**).

**Tab 21 (Table S20).** Summary of protein expression changes after DMF (50 μM, 24 h), EV-3 (10 μM, 24 h), BPK-21 (20 μM, 24 h), BPK-25 (10 μM, 24 h), and BPK-25 (10 μM, 24h) + MG132 (10 μM, 24 h) treatment as determined by TMT-exp (average values from 2-5 donors are reported, related to **Figure 7C**).

**Supplementary Dataset 2**
RNA-sequencing data for primary human T cells treated with DMF (50 μM, 24 h), EV-3 (10 μM, 24 h), BPK-21

### (C) Biological Methods

EXPERIMENTAL MODEL AND SUBJECT DETAILS

#### Isolation of peripheral blood mononuclear cells (PBMC) and T cells

All studies with primary human cells were performed with samples from human volunteers followed by protocols approved by The Scripps Research Institute Institutional Review Board. Blood from healthy donors (age 18 to 65) was obtained after informed donor consent. Peripheral blood mononuclear cells (PBMCs) were isolated over Lymphoprep (STEMCELL Technologies) gradient using slightly modified manufacturer's instructions. Briefly, 25 mL of freshly isolated blood was layered on top of 12.5 mL of Lymphoprep in a 50 mL Falcon tube minimizing mixing of blood with Lymphoprep. The tubes were centrifuged at room temperature (931 *g*, 20 min, 23 °C) with brake off and the plasma and Lymphoprep layers containing PBMCs were transferred to new 50 mL Falcon tubes with a 2:1 dilution with PBS. The cells were pelleted (524 *g*, 8 min, 4 °C) and washed with PBS (20 mL) once. T cells were isolated from fresh PBMCs using EasySep Human T Cell Isolation Kit (STEMCELL Technologies, negative selection) according to manufacturer's instructions.

METHODS DETAILS

#### T cell activation for mass-spectrometry analysis

Non-tissue culture treated 6-well plates were pre-coated with $\alpha$CD3 (5 µg/mL, BioXCell) and $\alpha$CD28 antibodies (2 µg/mL, BioXCell) in PBS (2 mL/well) and kept at 4 °C overnight. The next day, the plates were transferred to a 37 °C incubator for 1 h and washed with PBS (2 x 5 mL/well). Freshly isolated T cells were resuspended in RPMI media supplemented with 10% FBS, *L*-glutamine (2 mM), penicillin (100 U/mL), and streptomycin (100 µg/mL) at 1 x $10^6$ cells/mL, plated into the pre-coated 6-well plates (6-10 mL/well) and kept at 37 °C in a 5% $CO_2$ incubator for 3 days. Following this incubation period, the cells were combined in 50 mL Falcon tubes, pelleted (524 *g*, 5 min, 4 °C), and washed with PBS (10 mL). The cells were then transferred into Eppendorf tubes in 1 mL of PBS and pelleted. PBS was then aspirated and the cells were either re-suspended in fresh RPMI media for *in situ* treatments or flash-frozen and kept at -80 °C until further analysis (*in vitro* treatments).

#### T cell expansion for mass-spectrometry analysis (control T cells)

A non-tissue culture treated 6-well plate was pre-coated with $\alpha$CD3 (1.5 µg/mL) antibody in PBS (3 mL/well) and kept at 4 °C overnight. The next day, the plates were transferred to a 37 °C incubator for 1 h and washed with PBS (2 x 5 mL/well). Freshly isolated T cells were re-suspended in RPMI media (10% FBS, *L*-glutamine (2 mM), penicillin (100 U/mL), streptomycin (100 µg/mL)), containing $\alpha$CD28 antibody (1 µg/mL) at 1 x $10^6$ cells/mL, plated into the pre-coated 6-well plate (6-10 mL/well) and kept at 37 °C in a 5% $CO_2$ incubator for 3 days. Following this incubation period the cells were combined in 50 mL Falcon tubes, pelleted (524 *g*, 5 min, 4 °C), and washed with PBS (10 mL). The cells were then re-suspended in RPMI media containing recombinant IL2 (10 U/mL) and kept at 37 °C in a 5% $CO_2$ incubator for 10-12 days, splitting the cells every 3-4 days to keep cell density below 2 x $10^6$ cells/mL. After this time, the cells were pelleted (524 *g*, 5 min, 4 °C), washed with PBS (10 mL) and either re-suspended in fresh RPMI media for *in situ* treatments or flash-frozen and kept at -80 °C until further analysis (*in vitro* treatments).

#### *In situ* labeling with cysteine-reactive electrophiles

Activated or expanded (control) T cells were re-suspended in RPMI media supplemented with 10% FBS, *L*-glutamine (2 mM), penicillin (100 U/mL), and streptomycin (100 µg/mL) at 2 x $10^6$ cells/mL. The compounds were added to cells as 1000x DMSO stocks and mixed well with the media by pipetting up and down after addition. The cells were kept at 37 °C in 5% $CO_2$ containing incubators for 3h (or otherwise specified times), then pelleted by centrifugation (524 *g*, 5 min, 4 °C), washed with cold PBS (10 mL) and transferred to Eppendorf tubes (1 mL PBS). The cells were pelleted again (524 *g*, 5 min, 4 °C), flash-frozen, and kept at -80 °C until further analysis.

#### Multidimensional screen for inhibition of T cell activation

Non-tissue culture treated 96-well plates were pre-coated with αCD3 (5 µg/mL) and αCD28 antibodies (2 µg/mL) in PBS (100 µL/well) and left at 4 °C overnight. Freshly isolated T cells were re-suspended in RPMI media supplemented with 10% FBS, *L*-glutamine (2 mM), penicillin (100 U/mL), and streptomycin (100 µg/mL) at 2 x $10^6$ cells/mL. Compound stocks (200x) in DMSO were diluted to 2x stocks in the working RPMI media in another 96-well plate. The pre-coated 96-well treatment plates were washed with PBS (2 x 200µL), T cells (100 µL/well, 2 x $10^5$ cells/well) were then added to the wells, followed by the addition of 2x compound stocks in RPMI media (100 µL). The outer wells of the plates were filled with media without cells to avoid the edge effect in the assay. The plates were then incubated at 37 °C in a 5% $CO_2$ containing incubator for 24 h. Following the treatment, the cells were transferred to a U-bottom 96-well plate and harvested by centrifugation (600 *g*, 3 min, 4 °C). The supernatants were kept and stored at -80 °C for further cytokine analysis, while the cells were washed with PBS (2 x 150 µL) prior to staining for flow cytometry analysis.

**Flow cytometry analysis**

Following the PBS washes, the cells were stained with fixable near-IR LIVE/DEAD cell stain (Invitrogen) according to manufacturer's instructions. Briefly, one vial of near-IR LIVE/DEAD dye was resuspended in DMSO (50 µL) and diluted with PBS (1:1000). The diluted stain was added to each well (200 µL) and the cells were incubated for 30 min at room temperature in the dark. After this time, the cells were pelleted (600 *g*, 3 min, 4 °C), washed once with PBS (200 µL/well) and incubated with a freshly made cocktail of antibodies for the appropriate cell surface markers diluted in PBS containing 2% FBS (1:400 antibody dilution). The corresponding data in **Figure 4D** is presented as the mean percentage of DMSO treated control ± SEM, n = 3/group.

**Measurement of phospho-NF-□B p65 (Ser536) levels**

Freshly isolated T cells (2 x $10^5$ cells/well) were harvested and stimulated as described before in a 96-well plate in the presence of DMSO or compounds of interest. Following the 24 h treatment, the cells were pelleted in a U-bottom plate, harvested by centrifugation (600 *g*, 3 min, 4 °C), washed with PBS, and stained with near-IR LIVE/DEAD dye as described above. After the staining, intracellular phospho-NF-κB p65 (Ser536) levels were measured using PE conjugate of phospho-NF-κB p65 (Ser536) (93H1) rabbit antibody (Cell Signaling Technology) according to manufacturer's instructions. Briefly, the cells were washed with PBS and fixed with 4% PFA in PBS (100 µL, 15 min, rt). The cells were washed with PBS again (2 x 150 µL), placed on ice and permeabilized with 90% aqueous MeOH (100 µL/well, slow addition with gentle mixing by pipetting up and down). Following a 30 min incubation on ice, the plate was sealed and stored at -20 °C overnight. The following day, the cells were thawed on ice, washed with PBS (150 µL x 2), and stained with PE conjugate of phospho-NF-κB p65 (Ser536) (93H1) rabbit antibody (50 µL, 1:100 dilution in incubation buffer (1% FBS in PBS)) for 1 h at rt in the dark. The cells were then washed with incubation buffer (150 µL x 2) and resuspended in PBS for further flow cytometry analysis. Data in **Figure 6B** are from a single experiment representative of at least two independent biological experiments.

**Measurement of intracellular glutathione levels**

Intracellular glutathione levels were determined using GSH-Glo glutathione assay (Promega Corporation) according to manufacturer's instructions. Briefly, freshly isolated T cells were treated with compounds or DMSO for 24 h under TCR-stimulating conditions (96-well plate, 1 x $10^5$ cells/well) at 37 °C in 5% $CO_2$ containing incubator, then transferred to a U-shape bottom 96-well plate and pelleted (600 *g*, 3 min, 4 °C). The supernatants were kept and stored at -80 °C for cytokine analysis. The cells were washed with PBS (2 x 150 µL) and resuspended in 50 µL of PBS. An aliquot of treated cells (25 µL) was then added to an equal volume of 2x GSH reaction buffer containing Glutathione S-transferase and Luciferin-NT substrate (1:50 dilution in GSH-Glo Reaction Buffer). The reaction was incubated for 30 min at rt, after which Luciferin Detection Reagent (in reconstitution buffer with esterase, 25 µL/well) was added, and the plate was incubated for an additional 15 min and luminescence was read using a CLARIOstar (BMG Labtech) plate reader. The corresponding data in **Figure 6A** is presented as the mean percentage of DMSO-treated control ± SEM, n = 3/group.

**DuoSet ELISA quantification of secreted cytokines (IL2, IFN□, TNF□)**

The levels of secreted IL2, IFN$\gamma$□ and TNF$\alpha$ after incubating T cells in the presence of DMSO or electrophilic compounds under TCR-stimulating conditions were measured using DuoSet ELISA cytokine kits (R&D Systems) in clear microplates (R&D Systems) according to manufacturer's instructions and read using a CLARIOstar (BMG Labtech) plate reader (450 nm). All cytokine concentrations were calculated according to the standard curve generated for each experiment. The corresponding data in **Figures 4D** and **6A** is presented as the mean percentage of DMSO-treated control ± SEM, n = 3/group.

**NFAT (nuclear factor of activated T cells) luciferase reporter assay**

NFAT activity was measured using the Jurkat-Lucia NFAT reporter cell line (Invivogen) according to manufacturer's procedure. Briefly, Jurkat-Lucia NFAT cells were cultured at 37 °C in 5% $CO_2$ containing incubator in manufacturer-recommended growth medium (RPMI, 2 mM *L*-glutamine, 25 mM HEPES, 10% heat-inactivated fetal bovine serum (FBS, 30 min at 56 °C), 100 µg/mL Normocin, Pen-Strep (50 U/mL-50 µg/mL)) keeping cell density below 2 x $10^6$ cells/mL. To maintain selection pressure, Zeocin (100 µg/mL) was added to the growth medium every other passage and the cell passage number was kept less than 10. On the day of compound treatment, the cells were pelleted (300 *g*, 5 min) and resuspended at 2.2 x $10^6$ cells/mL in fresh, pre-warmed test medium (RPMI, 2 mM *L*-glutamine, 25 mM HEPES, 10% heat-inactivated FBS, Pen-Strep (100 U/mL-100 µg/mL) without Normocin). The cell suspension (180 µL, 4 x $10^5$ cells/well) was then added to the test plate containing stimulating solution (20 µL/well, PMA (50 ng/mL) and ionomycin (3 µg/mL) in growth media) and test compounds (2 µL, 100x stock in DMSO) or DMSO, and the plate was kept at 37 °C in a 5% $CO_2$ containing incubator for 24 h. To evaluate expression of the luciferase reporter, 50 µL of Quanti-luc (Invivogen) detection reagent was combined with 20 µL of cell suspension from each well in a new 96-well white (opaque) plate and the luminescence was read using a CLARIOstar microplate reader (BMG Labtech). The corresponding data in **Figure 6C** is presented as the mean percentage of DMSO treated control ± SD or SEM, n = 2–5/group.

**ISRE-luciferase and CellTiter Glo assays**

THP-1 Lucia ISG cells were resuspended in low-serum growth media (2% FBS) at a density of 5 x $10^5$ cells/mL and treated with BPK-25 or vehicle (DMSO) in the presence of viral dsDNA (2 µg/mL). 50 µL of cells/well were seeded into each well of a 384-well white greiner plates and incubated for 24 h. To evaluate expression of the luciferase reporter, 30 µL of Quanti-luc (Invivogen) detection reagent was added to each well and luminescence was read using an Envision plate reader (Perkin Elmer) set with an integration time of 0.1 seconds. To evaluate cell viability, 30 µL of CellTiter-Glo (Promega) reagent was added to each well and each plate was read using the same instrument settings utilized for the luciferase assay. For each cell type and assay, luminescence signals for test article samples were normalized to vehicle-treated samples and reported as relative light units (RLU). The corresponding data in **Figure S5C** is presented as the mean percentage of DMSO-treated control ± SEM, n = 3/group.

**Bio-Plex quantification of secreted cytokines**

Freshly isolated PBMCs (4 x $10^6$ cells/mL, 1 mL/well), were treated with BPK-25 (10 µM) or vehicle (DMSO) for 6 h in a 24-well plate, after which cGAMP (10 µM) was added to the wells and the cells were incubated for additional 20 h. Following this treatment, the cells were transferred to 1.5 mL Eppendorf tubes and harvested by centrifugation (600 *g*, 8 min, 4 °C). The supernatants were saved (-80 °C) and used for further cytokine analysis using Bio-Plex Pro Human Cytokine assay (Bio-Rad) according to manufacturer's instructions. Bio-Plex Assay is a multiplex flow immunoassay that simultaneously detects and identifies cytokines based on fluorescent dye-labeled 6.5 µm magnetic beads in a single reaction. When run on the Bioplex 200 system, 50 µL of supernatant was mixed with 50 µL of beads and quantified against human cytokines standard curves. The corresponding data in **Figure S5E** is presented as the mean percentage of DMSO-treated control ± SEM, n = 3/group.

**ELISA quantification of secreted IFN-β**

Concentrations of IFN-β were determined with VeriKine-HS human IFN-β serum ELISA kit (PBL Assay Science) according to manufacturer's instructions. All concentrations of IFN-β were calculated according to the standard curve generated for each experiment. The corresponding data in **Figure S5E** is presented as the mean percentage of DMSO-treated control ± SD, n = 2/group.

**Western blot analysis**

Western blot analysis was performed on freshly isolated or expanded T cells. For Western blot protein degradation analysis, primary human T cells (2 x $10^7$ cells/treatment) were re-suspended in RPMI media at 2 x $10^6$ cells/mL and treated with the compounds or DMSO at 37 °C in a $CO_2$ containing incubator for 24 h (or otherwise indicated times). Following this incubation period, the cells were pelleted (600 *g*, 5 min, 4 °C), washed with PBS (10 mL), transferred to 1.5 mL Eppendorf tubes, flash-frozen, and stored at -80 °C until further analysis. On the day of the analysis, the cell pellets were thawed on ice, re-suspended in cold PBS and lysed by sonication with probe sonicator (2 x 8 pulses). Protein concentrations for all the samples were adjusted to 1 mg/mL, 4x loading buffer was added (10 µL to 30 µL of proteome), and the samples were heated at 95 °C for 5 min. The proteins were resolved using SDS-PAGE (10% acrylamide gel) and transferred to 0.45 µM nitrocellulose membranes (GE Healthcare). The membrane was blocked with 5% milk in Tris-buffered saline (20 mM Tris-HCl 7.6, 150 mM NaCl) with 0.1% tween 20 (TBST) buffer at rt for 1 h (or at 4 °C overnight), washed 3 times with TBST, and incubated with primary antibodies in 5% BSA in TBST at 4 °C overnight. Following another TBST wash (3 times), the membrane was incubated with secondary antibody (1:5000 in 5% milk in TBST) at 4 °C overnight. The membrane was washed with TBST (3 times), developed with ECL western blotting detection reagent kit (Thermo Scientific) and recorded on CL-X Posure film (Thermo Scientific). Relative band intensities were quantified using ImageJ software.(Rasband, W.S., ImageJ, U. S. National Institutes of Health, Bethesda, Maryland, USA, https://imagej.nih.gov/ij/, 1997-2018.)

**Western blot analysis of chromatin-bound proteins**

BPK-25 (10 µM) treated expanded T cells were washed with PBS before permeabilization by rotation at 4 °C for 10 min with cytoplasm lysis buffer (10 mM sodium phosphate pH 7.4, 25 mM KCl, 1.5 mM $MgCl_2$, 10% glycerol, and 0.025% NP-40 supplemented with 1x HALT protease inhibitor cocktail (Thermo Scientific)). Nuclei were pelleted (500 *g*, 5 min), and washed with cytoplasm lysis buffer without detergent, before being lysed by gentle sonication (Branson Sonifier 250) in cell lysis buffer (10 mM sodium phosphate pH 7.4, 25 mM KCl, 1.5 mM $MgCl_2$, 10% glycerol, and 1% NP-40, 0.1% SDS supplemented with 1x HALT, and 1x Benzoase (Pierce)) and rotated for 2 h at 4 °C. Insoluble material was precipitated by centrifugation (12,000 *g*, 10 min) and the protein concentration of nuclear extracts was measured using standard BCA assay (Thermo Scientific) and normalized. Electrophoretic separation was performed on Novex 4-20% Tris-Glycine Mini Gels (Invitrogen) using the Novex Wedgewell system, and transferred to 0.45 µM Nitrocellulose membranes (GE Healthcare). Primary antibodies were applied overnight at 4 °C in 5% BSA/TBST. Secondary antibodies were applied for 1 h at rt in 5% milk/TBST. Blots were imaged using fluorescence-labeled secondary antibodies (LI-COR) on the Odyssey CLx Imager. Relative band intensities were quantified using ImageJ software.

**Gene expression (qPCR) analysis**

Total RNA from compound or DMSO treated T cells (1.5 x $10^7$ cells/group) was isolated using RNeasy Mini Kit (Qiagen) according to manufacturer's protocol. RNA concentration was determined using NanoDrop and adjusted to 1 µg RNA in 15 µL RNAse free water for the reverse transcription reaction. cDNA amplification was done using iScript Reverse Transcription Supermix kit (BioRad) according to manufacturer's instructions. The following PCR settings were used for the reverse transcription reaction: 5 min at 25 °C (priming), 20 min at 46 °C (Reverse transcription), 1 min at 95 °C (RT inactivation), hold at 4 °C. qPCR analysis was performed on ABI Real Time PCR System (Applied Biosystems) with the SYBR green Mastermix (Applied Biosystems). Relative gene expression was normalized to actin.

qPCR primers used (5' to 3'):

| | |
|---|---|
| actin-fwd | AGAGCTACGAGCTGCCTGAC |
| actin-rev | AGCACTGTGTTGGCGTACAG |

| | |
|---|---|
| BIRC2-fwd | AGCACGATCTTGTCAGATTGG |
| BIRC2-rev | GGCGGGGAAAGTTGAATATGTA |
| BIRC3-fwd | AAGCTACCTCTCAGCCTACTTT |
| BIRC3-rev | CCACTGTTTTCTGTACCCGGA |
| IL6-fwd | AATTCGGTACATCCTCGACGG |
| IL6-rev | GGTTGTTTTCTGCCAGTGCC |
| IL1-beta-fwd | ACAGATGAAGTGCTCCTTCCA |
| IL1-beta-rev | GTCGGAGATTCGTAGCTGGAT |
| CXCL10-fwd | CCAGAATCGAAGGCCATCAA |
| CXCL10-rev | CATTTCCTTGCTAACTGCTTTCAG |

**RNA sequencing**

Total RNA from compound or DMSO treated T cells (1.5 x $10^7$ cells/group) was isolated using RNeasy Mini Kit (Qiagen) using RNAse free DNAse set (Qiagen) for on column DNA digestion according to manufacturer's protocol and stored at -80 °C until further analysis. RNA quality was assessed using TapeStation 4200 and RNA-Seq libraries were prepared using the TruSeq stranded mRNA Sample Preparation Kit v2 according to Illumina protocols. Multiplexed libraries were validated using TapeStation 4200, normalized, pooled and quantified by qPCR for sequencing. High-throughput sequencing was performed on the NextSeq 500 system (Illumina). Image analysis and base calling were done with Illumina CASAVA-1.8.2. Sequenced reads were quality-tested using FASTQC (Andrews S. (2010). FastQC: a quality control tool for high throughput sequence data. Available online at: http://www.bioinformatics.babraham.ac.uk/projects/fastqc) and aligned to the hg19 human genome using the STAR (Dobin et al., 2013) version 2.5.3a. Mapping was carried out using default parameters (up to 10 mismatches per read, and up to 9 multi-mapping locations per read). The genome index was constructed using the gene annotation supplied with the hg19 Illumina iGenomes (iGenomes online. Illumina. 2015. http://support.illumina.com/sequencing/sequencing_software/igenome.html) collection and overhang value of 100. Gene expression counts were quantified with HOMER (Heinz et al., 2010) v4.10.4 analyzeRepeats.pl using the parameters -raw -count exons -condenseGenes -strand +. Normalized gene expression counts were calculated using HOMER v4.10.4 analyzeRepeats.pl using the parameters -fpkm -count exons -condenseGenes -strand + -normMatrix 1000000. Differential expression was carried out with HOMER v4.10.4 getDiffExpression.pl using the default DESeq2 v1.14.1 subroutine on the raw count table.

**isoTOP-ABPP sample preparation**

Activated or expanded (control) primary human T cells were re-suspended in RPMI (1 x 10^6 cells/mL) containing 10% fetal bovine serum (FBS), penicillin (100 U/mL), streptomycin (100 μg/mL), and *L*-glutamine (2 mM). The cells were treated with DMSO or compounds for 3 h, pelleted (524 *g*, 5 min), washed with PBS, and lysed by sonication (2 x 8 pulses). Soluble and particulate proteomic fractions were separated by ultracentrifugation (100,000 *g*, 45 min), and protein concentration was normalized to 1.7 mg/mL using a standard DC protein assay (Bio-Rad). The resulting proteomes were analyzed by competitive isotopic Tandem Orthogonal Proteolysis Activity-Based Protein Profiling (isoTOP-ABPP) using a protocol from Gao *et al*. (Gao et al., 2018)

**IA-alkyne labeling and click chemistry**

Samples (500 μL, 1.7 mg/mL) were treated with iodoacetamide alkyne (IA-alkyne, 5 μL of 10 mM stock in DMSO, final concentration: 100 μM) for 1 h at ambient temperature. Modified proteins were then conjugated to isotopically labeled, TEV-cleavable biotin tags (TEV-tags) using copper-catalyzed azide-alkyne cycloaddition reaction (CuAAC). Reagents for the CuAAC reaction were pre-mixed prior to their addition to the proteome samples. TEV tags (light or heavy, 10 μL of 5 mM stocks in DMSO, final concentration = 100 μM), tris(benzyltriazolylmethyl)amine ligand (TBTA; 30 μL of 1.7 mM stock in DMSO:*t*-butanol 1:4, final concentration = 100 μM), tris(2-carboxyethyl)phosphine hydrochloride (TCEP; 10 μL of fresh 50 mM stock in water, final concentration = 1 mM), and $Cu(OAc)_2$ (10 μL of 50 mM stock in water, final concentration = 1 mM) were combined in an Eppendorf tube, vortexed and added to the proteomes (55 μL/sample). The CuAAC reaction mixture containing "heavy" TEV tag was added to the DMSO-treated control samples and the CuAAC reaction mixture containing “light” TEV tag was added to compound-treated samples. The reaction was allowed to proceed at rt for 1 h, “heavy” and “light” samples were combined pairwise in 15 mL conical Falcon tubes on ice containing 4 mL of cold methanol (pre-chilled at –80 °C), 1 mL $CHCl_3$, and 1 mL $H_2O$. Eppendorf tubes from the reaction mixtures were washed with additional $H_2O$ (1mL each) and the washes were added to the same Falcon tube (final ratios MeOH : $CHCl_3$ : $H_2O$ = 4 : 1 : 4). Following centrifugation (5,000 *g*, 10 min, 4 °C), a protein disk formed at the interface of $CHCl_3$ and aqueous layers. Both layers were aspirated without perturbing the disk, which was resuspended in cold MeOH (2 mL) and $CHCl_3$ (1 mL) by vortexing. The proteins were pelleted (5,000 *g*, 10 min, 4 °C), and the resulting pellets were solubilized in 1.2% SDS in PBS (1 mL) with sonication and heating (95 °C, 5 min).

**isoTOP-ABPP sample streptavidin enrichment**

Once solubilized, the samples were diluted with PBS (4 mL) and streptavidin-agarose beads were added for the enrichment (final SDS concentration: 0.2% in PBS). The beads (100 µL of a 50% slurry per sample) were washed with PBS (2 x 10 mL) and resuspended in 1 mL of PBS per sample prior to addition. The final mixture was rotated for 3 h at rt. Following this enrichment step, the beads were pelleted by centrifugation (2,000 *g*, 2 min) and washed to remove non-specifically binding proteins (2 x 10 mL 0.2% SDS in PBS, 2 x 10 mL PBS, and 2 x 10 mL $H_2O$).

**isoTOP-ABPP sample trypsin and TEV digestion**

After the last wash, the beads were transferred to new Eppendorf tubes in water (2 x 0.5 mL), pelleted (4,000 *g*, 3 min), and resuspended in 6M urea in PBS (0.5 mL). DTT (25 µL of a fresh 200 mM stock in water, final concentration – 10 mM) was added and the beads were incubated at 65 °C for 15 min. Iodoacetamide (25 µL of a 400 mM stock in water, final concentration – 20 mM) was then added and the samples were incubated in the dark at 37 °C with shaking for 30 min. Following this incubation, the mixture was diluted with PBS (900 µL), the beads were pelleted by centrifugation and resuspended in 2M urea in PBS (200 µL). Trypsin (Promega, sequencing grade; 2 µg in 6 µL of trypsin buffer containing 1 mM $CaCl_2$) was added to the mixture and the digestion was allowed to proceed overnight at 37 °C with shaking. The beads were pelleted (2,000 *g*, 2 min) and the tryptic digest was aspirated. The beads were then washed (3 x 1 mL PBS, 3 x 1 mL $H_2O$), transferred to a new Eppendorf tube in $H_2O$ (2 x 0.5 mL), washed with TEV buffer (200 µL, 50 mM Tris, pH 8, 0.5 mM EDTA, 1 mM DTT), and resuspended in TEV buffer (140 µL). TEV protease (4 µL, 80 µM) was then added and the beads were incubated at 30 °C overnight with rotation. Following the overnight digestion, the beads were pelleted by centrifugation (2,000 *g*, 2 min) and the TEV digest was separated from the beads using Micro Bio-Spin coulumns (Bio-rad) with centrifugation (800 *g*, 0.5 min) and an additional wash (100 µL $H_2O$). The samples were then acidified by the addition of 0.1% FA (14 µL, final concentration: 5% v/v) and stored at -80 °C prior to analysis.

**isoTOP-ABPP liquid-chromatography-mass-spectrometry (LC-MS/MS) analysis**

Samples were pressure-loaded onto a 250 µm (inner diameter) fused silica capillary columns packed with C18 resin (Aqua 5 µm, Phenomenex) and analyzed by multidimensional liquid chromatography tandem mass-spectrometry (MudPIT) using an LTQ-Velos Orbitrap mass spectrometer (Thermo Scientific) coupled to an Agilent 1200-series quaternary pump. The peptides were eluted onto a biphasic column with a 5 µm tip (100 µm fused silica, packed with C18 (10 cm) and bulk strong cation exchange resin (3 cm, SCX, Phenomenex) in a 5-step MudPIT experiment, using 0%, 30%, 60%, 90%, and 100% salt bumps of 500 mM aqueous ammonium acetate and a 5%–100% gradient of buffer B in buffer A (buffer A: 95% water, 5% $CH_3CN$, 0.1% FA; buffer B: 5% water, 95% $CH_3CN$, 0.1% FA) as previously described.(Weerapana et al., 2007) Data were collected in data-dependent acquisition mode with dynamic exclusion enabled (20 s, repeat of 2). One full MS (MS1) scan (400-1800 m/z) was followed by 30 MS2 scans (ITMS) of the $n^{th}$ most abundant ions.

**isoTOP-ABPP peptide identification**

The MS2 spectra data were extracted from the raw file using RAW Converter (version 1.1.0.22; available at http://fields.scripps.edu/rawconv/), uploaded to Integrated Proteomics Pipeline (IP2), and searched using the ProLuCID algorithm (publicly available at http://fields.scripps.edu/downloads.php) using a reverse concatenated, non-redundant variant of the Human UniProt database (release-2012_11). Cysteine residues were searched with a static modification for carboxyamidomethylation (+57.02146) and up to one differential modification for either the light or heavy TEV tags (+464.28595 or +470.29976 respectively). Peptides were required to have at least one tryptic terminus and to contain the TEV modification. ProLuCID data was filtered through DTASelect (version 2.0) to achieve a peptide false-positive rate below 1%.

**isoTOP-ABPP R value calculation and data processing**

The heavy/light isoTOP-ABPP ratios (R values) for each unique peptide (DMSO/compound treated) were quantified with in-house CIMAGE software (Weerapana et al., 2010) using default parameters (3 MS1 acquisitions per peak and signal to noise threshold set to 2.5). Site-specific engagement of cysteine residues was assessed by blockade of IA-alkyne probe labeling. A maximal ratio of 20 was assigned for peptides that showed a ≥ 95% reduction in MS1 peak area in the compound treated proteome (light TEV tag) compared to the control DMSO-treated proteome (heavy TEV tag). Ratios for unique peptide sequences were calculated for each experiment; overlapping peptides with the same modified cysteine (e.g., different charge states, elution times or tryptic termini) were grouped together and the median ratio was reported as the final ratio (R). Additionally, ratios for peptide sequences containing multiple cysteines were grouped together. When aggregating data across experimental replicates, the mean of each experimental median R was reported. The peptide ratios reported by CIMAGE were further filtered to ensure the removal or correction of low-quality ratios in each individual dataset. The quality filters applied were the following: removal of half-tryptic peptides, removal of peptides with more than one tryptic miscleavage site, removal of peptides with R = 20 and only a single MS2 event triggered during the elution of the parent ion, removal of non-unique peptides. Further filtering was then performed as described below for each experiment type.

**Combining data across experimental groups**

To combine data across replicates from different experiment groups (e. g., broad ligandability and elaborated fragment data or hyper-reactivity) or different experiment types (e. g., TMT and isoTOP), we used identifiers consisting of the Uniprot accession concatenated with the tryptic sequence associated with the particular peptide. Peptides that contained the same modified cysteine or where multiple cysteines were modified on that peptide were combined. When data from an experiment group associated with a miscleaved peptide sequence was combined with data from another group which contained a non miscleaved variant of the same peptide, all data was reported under the fully tryptic identifier, unless the non miscleaved variant introduced an additional cysteine, in which case the data was not merged.

**Filtering of broad ligandability (scout fragment) data**

All peptides with R = 20 were manually reviewed. Peptides with R = 20 were discarded if the ratio set contained a single 20, and the minimum ratio in the set was less than 4. If the ratio set contained two or more 20 values and the minimum ratio in the set was less than 2, these 20 values were also discarded. This filter was applied on R values within a single experiment and when aggregating data from replicate experiments.

When aggregating data from replicate experiments, for peptides that had standard deviations greater than 60% of the mean, the lowest ratio of that set was reported, unless the minimum ratio of the set was ≥ 4, in which case the average ratio was reported. Individual peptide sequences were required to have been quantified (R ≠ 0) in at least two replicates per condition. Peptides were considered liganded if they had a final value of R ≥ 5.

**Filtering of elaborated compound data**

All peptides with R = 20 were manually reviewed. Within individual replicates, peptides with R = 20 were discarded if the ratio set contained a single 20, and the minimum ratio in the set was less than 4. When aggregating ratios across replicates, peptides with R = 20 were discarded if the ratio set contained a single 20, and the minimum ratio in the set was less than 3. Individual peptide sequences were required to have been quantified (R ≠ 0) in at least two replicates per condition, unless they had R ≥ 4 in both particulate and soluble conditions for a given compound.

During manual review of the data, some peptides were exempted from specific filters due to additional evidence of their validity. BIRC2 (C45) R = 20 and CASP2 (C366, C370) R = 20 values derived from DMF datasets values derived from DMF datasets were exempted from applied 20-filters as the same residues were convincingly liganded in the TMT datasets.
Peptides were considered liganded if they had a final value of R ≥ 4.

**Filtering and processing of hyper-reactivity data**

Peptides with R = 20 were discarded if the ratio set contained a single 20, and the minimum ratio in the set was less than 4. This filter was applied on R values within a single experiment and when aggregating data from replicate experiments.
Data from these experiments was separated according to activation state and the minimal ratio between soluble and particulate fractions for each state was reported for each peptide.

**TMT-ABPP sample preparation and IA-DTB labeling**

Samples (500µL, 1.7 mg/mL) were treated with iodoacetamide polyethyleneoxide desthiobiotin (IA-DTB (Santa Cruz), 5 µL of 10 mM stock in DMSO, final concentration: 100 µM) for 1 h at ambient temperature. Ice-cold MeOH (500 µL) and $CHCl_3$ (200 µL) were then added, the mixture was vortexed and centrifuged (10,000 *g*, 10 min, 4 °C) to afford a protein disc at the interface of $CHCl_3$ and aqueous layers. Both layers were aspirated without perturbing the disk, which was re-suspended in cold methanol (500 µL) and $CHCl_3$ (200 µL) by sonication. The proteins were pelleted (10,000 *g*, 10 min, 4 °C), and the resulting pellets were re-suspended in 90 µL of buffer containing 9M urea, 10 mM DTT and 50 mM triethylammonium bicarbonate (1/20 dilution of 1.0 M stock solution, pH 8.5) by thorough pipetting up and down. The resulting mixture was heated at 65 °C for 20 min. Sample was cooled to room temp, iodoacetamide (10 µL, 500 mM solution in $H_2O$) was added, and the samples were incubated at 37 ˚C for 30 min with shaking.

**TMT-ABPP trypsin digestion and streptavidin enrichment**

Following the labeling with iodoacetamide, samples were diluted with 305 µL of triethylammonium bicarbonate buffer (50 mM, 1/20 dilution of 1.0 M stock, pH 8.5; Final urea concentraition: 2.0 M). Trypsin (4 µL of 0.25 µg/µL trypsin in trypsin buffer, containing 25 mM $CaCl_2$) was then added and the proteins were digested at 37 ˚C overnight. The following day, samples were diluted with wash buffer (400 µL, 25 mM Tris-HCl, pH 7.5, 150 mM NaCl, 0.2% NP-40), streptavidin-agarose beads (50% slurry in wash buffer) were added to each sample (40 µL/sample) and the bead mixture was rotated for 2 h at rt. Briefly, for a 10-plex sample, streptavidin-agarose bead slurry (440µL, 50% slurry) was washed (2 x 1 mL, 25 mM Tris-HCl, pH 7.5, 150 mM NaCl, 0.1% NP-40) and brought up to the initial volume in the wash buffer prior to the addition to the sample. After incubation, the beads were pelleted by centrifugation (2,000 *g*, 1 min), transferred to a BioSpin column and washed (3 x 1 mL wash buffer, 3 x 1 mL PBS, 3 x 1 mL $H_2O$). Peptides were eluted by the addition of 300 µL of 50% aqueous $CH_3CN$ containing 0.1% FA. The eluate was then evaporated to dryness using SpeedVac vacuum concentrator.

**TMT tag labeling**

Peptides were resuspended in 100 µL EPPS buffer (200 mM, pH 8.0) with 30% dry $CH_3CN$, vortexed and spun down (2,000 *g*, 1 min). TMT tags (3 µL/channel in dry $CH_3CN$, 20 µg/µL) were added to the corresponding tubes and the reaction was allowed to proceed for 75 min. The reaction was quenched by the addition of 5% hydroxylamine (3 µL per sample), vortexed and left at room temperature for 15 min. FA (5 µL) was then added to each tube, the tubes were vortexed, spun down and combined in a low binding 1.5 mL Eppendorf tube. The final combined sample was dried in a SpeedVac vacuum concentrator and kept at -80 °C until the high pH fractionation step.

**High pH fractionation**

The spin columns for high pH fractionation were pre-equilibrated prior to use. Briefly, the columns were placed in Eppendorf tubes (2 mL), spun down to remove the storage solution (5,000 *g*, 2 min), and washed with $CH_3CN$ (2 x 300 µL, 5,000 *g*, 2 min) and buffer A (2 x 300 µL, 95% $H_2O$, 5% $CH_3CN$, 0.1% FA, 5,000 *g*, 2 min). TMT labeled peptides were re-dissolved in buffer A (300 µL, 95% $H_2O$, 5% $CH_3CN$, 0.1% FA) and loaded onto pre-equilibrated spin columns for high pH fractionation. The columns were spun down (2,000 *g*, 2 min) and the flow through was used to wash the original Eppendorf tube and passed through the spin column again (2,000 *g*, 2 min). The column was then washed with buffer A (300 µL, 2,000 *g*, 2 min) and 10 mM aqueous $NH_4HCO_3$ containing 5% $CH_3CN$ (300 µL, 2,000 *g*, 2 min), and the flow through was discarded. The peptides were eluted from the spin column into fresh Eppendorf tubes (2.0 mL) with a series of 10 mM $NH_4HCO_3$ / $CH_3CN$ buffers (2000 *g*, 2 min). The following buffers were used for peptide elution:

| Fraction Number | Acetonitrile (%) | Acetonitrile (µL) | 10 mM $NH_4HCO_3$ (µL) |
|---|---|---|---|
| **1** | 7.5 | 75 | 925 |
| **2** | 10.0 | 100 | 900 |
| **3** | 12.5 | 125 | 875 |
| **4** | 15.0 | 150 | 850 |
| **5** | 17.5 | 175 | 825 |
| **6** | 20.0 | 200 | 800 |
| **7** | 22.5 | 225 | 775 |
| **8** | 25.0 | 250 | 750 |
| **9** | 27.5 | 275 | 725 |
| **10** | 30.0 | 300 | 700 |
| **11** | 32.5 | 325 | 675 |
| **12** | 35.0 | 350 | 650 |
| **13** | 37.5 | 375 | 625 |
| **14** | 40.0 | 400 | 600 |
| **15** | 42.5 | 425 | 575 |
| **16** | 45.0 | 450 | 550 |
| **17** | 47.5 | 475 | 525 |
| **18** | 50.0 | 500 | 500 |
| **19** | 52.5 | 525 | 475 |
| **20** | 55.0 | 550 | 450 |
| **21** | 75.0 | 750 | 250 |

Every 7$^{th}$ fraction was combined into a new clean Eppendorf tube (2 mL) and the solvent was removed using SpeedVac vacuum concentrator. The resulting 7 combined fractions were re-suspended in buffer A (10 µL) and analyzed on the Orbitrap Fusion mass-spectrometer (5 µL injection volume).

**Whole proteome TMT (TMT-exp) sample preparation**

Freshly isolated T cells (1.6 x 10^7 cells, 2 x $10^6$ cells/mL in RPMI media) were treated with compound or DMSO for 24 h, pelleted (600 *g*, 5 min), and washed with PBS (1 x 10 mL). The cells were then transferred to an Eppendorf tube in additional PBS (1 mL), pelleted (600 *g*, 5 min), flash frozen, and kept at -80 °C until further analysis. Cell pellets were thawed on ice and lysed in lysis buffer (150 µL, 1 tablet of Roche complete, mini, EDTA-free Protease Inhibitor Cocktail dissolved in 10 mL of PBS) using probe sonicator (2 x 8 pulses). Protein concentration was adjusted to 2.0 mg/mL and the samples (100 µL, 200 µg protein) were transferred to new

Eppendorf tubes (1.5 mL) containing urea (48 mg/tube, final urea concentration: 8 M). DTT (5 µL, 200 mM fresh stock in $H_2O$, final DTT concentration: 10 mM) was then added to the tubes and the samples were incubated at 65 °C for 15 min. Following this incubation, iodoacetamide (5 µL, 400 mM fresh stock in $H_2O$, final IA concentration: 20 mM) was added and the samples were incubated in the dark at 37 °C with shaking for 30 min. Ice-cold MeOH (600 µL), $CHCl_3$ (200 µL), and $H_2O$ (500 µL) were then added, the mixture was vortexed and centrifuged (10,000 *g*, 10 min, 4 °C) to afford a protein disc at the interface of $CHCl_3$ and aqueous layers. The top layer was aspirated without perturbing the disk, additional MeOH (600 µL) was added and the proteins were pelleted (10,000 *g*, 10 min, 4 °C) and used in the next step or stored at -80 °C overnight.

**Whole proteome TMT LysC and trypsin digestion**

The resulting protein pellets were resuspended in EPPS buffer (160 µL, 200 mM, pH 8) using probe sonicator (2 x 6 pulses). LysC solution (4 µL/sample, 20 µg in 40 µL of HPLC grade water) was added and the samples were incubated at 37 °C with shaking for 2 h. Trypsin (10 µL, 0.5 µg/µL in trypsin buffer) and $CaCl_2$ (1.8 µL, 100 mM in $H_2O$) were then added and the samples were incubated at 37 °C with shaking overnight.

**Whole proteome TMT labeling with TMT tags**

Peptide concentration was determined using the microBCA assay (Thermo Scientific) according to manufacturer's instructions. For each sample, a volume corresponding to 25 µg of peptides was transferred to a new Eppendorf tube and the total volume was brought up to 35 µL with EPPS buffer (200 mM, pH 8). The samples were diluted with $CH_3CN$ (9 µL) and incubated with the corresponding TMT tags (3 µL/channel, 20 µg/µL) at rt for 30 min. Additional TMT tag (3 µL/channel, 20 µg/µL, 30 min) was added and the samples were incubated for another 30 min. Labeling was quenched by the addition of hydroxylamine (6 µL, 5% in $H_2O$). Following a 15 min incubation at rt, formic acid was added (2.5 µL, final FA concentration: 5%) and the samples were stored at -80 °C until further analysis.

**Whole proteome TMT ratio check and high pH fractionation**

Small aliquots (2 µL) from each channel were combined in a separate Eppendorf tube and dried using SpeedVac vacuum concentrator. The residue was re-dissolved in Buffer A (20 µL) and desalted using C18 stage tips (made in-house using 200 µL pipette tips and C18 discs (3M Empore)). Briefly, the stage-tip was activated by passing MeOH (2 x 50 µL) through the stage tip and washed with Buffer B (2 x 50 µL, 5% $H_2O$, 95% $CH_3CN$, 0.1% FA), followed by Buffer A (2 x 50 µL, 5% $CH_3CN$/95% $H_2O$, 0.1% FA). The sample was then loaded and the stage-tip was washed with Buffer A. The sample was eluted into a new Eppendorf tube with Buffer B (2 x 50 µL) and dried using SpeedVac vacuum concentrator. The residue was re-dissolved in Buffer A (10 µL) and analyzed by mass-spectrometry using the following LC-MS gradient: 5% buffer B in buffer A from 0-15 min, 5-15% buffer B from 15-17.5 min, 15-35% buffer B from 17.5-92.5 min, 35-95% buffer B from 92.5-95 min, 95% buffer B from 95-105 min, 95-5% buffer B from 105-107 min, and 5% buffer B from 107-125 min (buffer A: 95% $H_2O$, 5% $CH_3CN$, 0.1% FA; buffer B: 5% $H_2O$, 95% $CH_3CN$, 0.1% FA) and standard MS3-based quantification described below. Ratios were determined from the average peak intensities corresponding to each channel. For a ten-plex experiment, samples (20 µL/channel, final volumes adjusted based on the determined ratios) were combined in a new low binding Eppendorf tube (1.5 mL) and dried using SpeedVac. The residue was subjected to high pH fractionation as described above to yield 7 fractions which were re-suspended in buffer A (24 µL/sample) and analyzed by liquid chromatography tandem mass-spectrometry.

**TMT-ABPP and whole proteome TMT liquid chromatography-mass-spectrometry (LC-MS) analysis**

Samples were analyzed by liquid chromatography tandem mass-spectrometry using an Orbitrap Fusion mass spectrometer (Thermo Scientific) coupled to an UltiMate 3000 Series Rapid Separation LC system and autosampler (Thermo Scientific Dionex). The peptides were eluted onto a capillary column (75 µm inner diameter fused silica, packed with C18 (Waters, Acquity BEH C18, 1.7 µm, 25 cm) and separated at a flow rate of 0.25 µL/min using the following gradient: 5% buffer B in buffer A from 0-15 min, 5-35% buffer B from 15-155 min, 35-

95% buffer B from 155-160 min, 95% buffer B from 160-169 min, 95-5% buffer B from 169-170 min, and 5% buffer B from 170-200 min (buffer A: 95% $H_2O$, 5% acetonitrile, 0.1% FA; buffer B: 5% $H_2O$, 95% $CH_3CN$, 0.1% FA). The voltage applied to the nano-LC electrospray ionization source was 1.9 kV. Data was acquired using an MS3-based TMT method adapted from Wang, Y. *et al*. (Wang et al., 2019) Briefly, the scan sequence began with an MS1 master scan (Orbitrap analysis, resolution 120,000, 400−1700 m/z, RF lens 60%, automatic gain control [AGC] target 2E5, maximum injection time 50 ms, centroid mode) with dynamic exclusion enabled (repeat count 1, duration 15s). The top ten precursors were then selected for MS2/MS3 analysis. MS2 analysis consisted of: quadrupole isolation (isolation window 0.7) of precursor ion followed by collision-induced dissociation (CID) in the ion trap (AGC 1.8E4, normalized collision energy 35%, maximum injection time 120 ms). Following the acquisition of each MS2 spectrum, synchronous precursor selection (SPS) enabled the selection of up to 10 MS2 fragment ions for MS3 analysis. MS3 precursors were fragmented by HCD and analyzed using the Orbitrap (collision energy 55%, AGC 1.5E5, maximum injection time 120 ms, resolution was 50,000). For MS3 analysis, we used charge state–dependent isolation windows. For charge state z = 2, the MS isolation window was set at 1.2; for z = 3-6, the MS isolation window was set at 0.7. The MS2 and MS3 files were extracted from the raw files using RAW Converter (version 1.1.0.22; available at http://fields.scripps.edu/rawconv/), uploaded to Integrated Proteomics Pipeline (IP2), and searched using the ProLuCID algorithm (publicly available at http://fields.scripps.edu/downloads.php) using a reverse concatenated, non-redundant variant of the Human UniProt database (release-2012_11). Cysteine residues were searched with a static modification for carboxyamidomethylation (+57.02146 Da) and up to one differential modification for the desthiobiotin (DTB) tag (+398.2529 Da). N-termini and lysine residues were also searched with a static modification corresponding to the TMT tag (+229.1629 Da). Peptides were required to be at least 6 amino acids long, to have at least one tryptic terminus, and to contain the DTB modification. ProLuCID data was filtered through DTASelect (version 2.0) to achieve a peptide false-positive rate below 1%. The MS3-based peptide quantification was performed with reporter ion mass tolerance set to 20 ppm with Integrated Proteomics Pipeline (IP2).

**TMT-ABPP R value calculation for broad ligandability data**

At the individual TMT experiment level, the following filters were applied to remove low-quality peptides: removal of non-unique peptides, removal of half-tryptic peptides, removal of peptides with more than one internal missed cleavage sites, removal of peptides with low (<20,000) sum of reporter ion intensities for either expanded or activated control channels, removal of peptides with high variation between the replicate control channels (coefficient of variance >0.5), and peptides corresponding to the lower average reporter ion intensity control channels (activated vs expanded) if the difference in the average reporter ion intensity between expanded and activated control channels was more than two-fold. R-value (DMSO-treated vs. KB02/KB05-treated) for each peptide entry was calculated using the reporter ion intensities of DMSO and KB02/KB05 treated TMT channels for each treatment group with a maximum ratio cap of 20. Once the R values were calculated, two types of grouping were performed to aggregate peptide quantification data: 1) overlapping peptides with the same modified cysteine (e.g., different charge states, high pH fractionation fractions, or tryptic termini) were grouped together, then their R values were averaged, and the shortest unique tryptic peptide was reported; 2) multiple modified cysteines on a tryptic peptide were grouped together, then the averaged R values were used for further data processing. Peptides with high donor variation (R>5 for one donor, while R<2 for the other donor) were discarded (<1%), then the R values of replicate channels of the same condition were averaged to obtain the final reported data. A cysteine was required to be quantified in at least two TMT channels for each proteomic fraction to be reported.

**TMT-ABPP R value calculation for elaborated compounds dataset**

At the individual TMT experiment level, the following filters were applied to remove low-quality peptides: removal of non-unique peptides; removal of half-tryptic peptides, removal of peptides with more than one internal missed cleavage site, removal of peptides with low (<10,000) sum of reporter ion intensities for control channels, and peptides with high variation between the replicate control channels (coefficient of variance >0.5). R values (compound-treated vs. DMSO-treated) for each peptide entry were calculated using the reporter ion intensities of DMSO and compound treated TMT channels for each treatment group with a maximum ratio cap of 20. Once the R values for each peptide entry were calculated, two types of grouping were performed to aggregate peptide quantification data: 1) overlapping peptides with the same modified cysteine (e.g., different charge states, high pH fractionation fractions, or tryptic termini) were grouped together, then their R values were averaged, and the shortest unique tryptic peptide was reported; 2) multiple modified cysteines on a tryptic peptide were grouped together, then the averaged R values were used for further data processing. The R values of replicate channels of the same condition were averaged to obtain the final reported data with the requirement that all included peptides have been quantified in at least two individual experiments.

**Whole proteome protein ratios calculation for elaborated compounds dataset**

At the individual TMT experiment level, the following filters were applied to remove low-quality peptides: removal of non-unique peptides, removal of half-tryptic peptides, removal of peptides with more than one internal missed cleavage site, removal of cysteine-containing peptides, removal of peptides with low (<10,000) sum of reporter ion intensities for control channels, and peptides with high variation between the replicate control channels (coefficient of variance >0.5). R values (compound-treated vs. DMSO-treated) were calculated using the reporter ion intensities of compound and DMSO treated TMT channels for each treatment group. Then the ratios of all peptides of a protein were averaged to be reported as the final protein ratio. Proteins were required to have at least two unique quantified peptides in each experiment and were quantified in at least two independent experiments.

**Whole proteome protein ratios calculation for state-dependent reactivity dataset**

The MS3-based peptide quantification was performed with reporter ion mass tolerance set to 20 ppm with Integrated Proteomics Pipeline (IP2). At the individual TMT experiment level, the following filters were applied to remove low-quality peptides: removal of non-unique peptides, removal of half-tryptic peptides, removal of peptides with more than one internal missed cleavage sites, removal of peptides with low (<10,000) sum of reporter ion intensities (5 channels/donor), and peptides with high variation between either of the replicate channels for expanded or activated T cells (coefficient of variance >0.5). R values (activated vs. expanded) for each peptide entry were calculated using the average reporter ion intensities of activated and expanded TMT channels. Then the ratios of all quantified peptides for a protein were averaged to obtain the final protein ratio. Proteins were required to have at least two unique quantified peptides in each experiment.

**TMT-ABPP R value calculation for cysteine state-dependent reactivity dataset**

At the individual TMT experiment level, the following filters were applied to remove low-quality peptides: removal of non-unique peptides, removal of half-tryptic peptides, removal of peptides with more than one internal missed cleavage site, removal of peptides with low (<10,000) sum of reporter ion intensities in both expanded or activated channels, removal of peptides with high variation (coefficient of variance >0.5) between the replicate expanded or activated channels if their sum of reporter ion intensities is greater than 5,000. R values (activated vs. expanded) for each peptide were calculated using the average reporter ion intensities of activated and expanded TMT channels. Once the R values were calculated, two types of grouping were performed to aggregate peptide quantification data: 1) overlapping peptides with the same modified cysteine (e.g., different charge states, high pH fractionation fractions, or tryptic termini) were grouped together, then their R values were averaged, and the shortest unique tryptic peptide was reported; 2) multiple modified cysteines on a tryptic peptide were grouped together, then the averaged R values were reported for further data processing. The median value derived from at least two biological replicates was reported as the final R value for each peptide with a maximum ratio cap of 20.

**Data processing and analysis for IA-DTB reactivity dataset**

Proteins must have at least three unique quantified peptides in either particulate or soluble fraction in the TMT-ABPP experiments within the state-dependent dataset to be analyzed. The fraction with the most quantified unique peptides was selected for analysis for each protein. If a protein had an equal number of unique quantified peptides in both fractions, the peptide R ratios (activated vs. expanded) from both fractions were averaged. To account for potential donor variations in protein expression level, proteins were required to have at least one peptide R ratio within 1.5-fold of the protein expression level measured in TMT-exp experiments (if available) and were excluded from the analysis if all peptide R ratios were greater than 2.0 or less than 0.5. For proteins with 5 or more quantified peptides, a cysteine was considered for potential change in reactivity if its peptide R value differed more than two-fold from both the median R value of all quantified cysteines on the same protein and from the protein expression level measured in TMT-exp experiments (if available). For proteins with three or four quantified peptides, a cysteine was considered for potential change in reactivity if its peptide R value differed more than two-fold from the protein expression level measured by TMT-exp data, with an additional requirement that the maximum peptide R ratio differed more than 2-fold from the minimum peptide R ratio. All the cysteines that passed the initial filters described above were manually curated to remove low quality profiles.

**Generation of in vitro transcribed sgRNAs**

DNA templates consisting of a T7 RNA Polymerase promoter, the ~20nt target-specific sequence, and the chimeric sgRNA scaffold were generated for each desired target by overlapping PCR using Q5 High Fidelity Master Mix (New England Biolabs) under the following conditions: 98 °C for 2 min; 50 °C for 10 min; 72 °C for 10 min. Guide RNA templates were used to transcribe guide RNAs using the HiScribe T7 High Yield RNA Synthesis kit (New England Biolabs) according to the manufacturer's instructions. Following *in vitro* transcription, guides were purified using Monarch RNA Cleanup Kit (New England Biolabs) following manufacturer's instructions.

**Cas9 Ribonucleoprotein (RNP) Assembly and Electroporation**

The Cas9 RNPs were assembled before transfection using the ArciTect™ Cas9-eGFP Nuclease (StemCell) with the T7 transcribed RNAs at a molar ratio of 1:3 in Buffer T. For each target of interest, the genome was tiled with 3 unique guide RNAs. Before the transfection, primary T cells were preactivated on αCD3/αCD28-precoated plates in complete RPMI medium supplemented with 100 U/mL IL2 for 48 h. The T cells were then washed with PBS and resuspended in Buffer T (10 x $10^6$ cells/mL) and the Cas9 RNP transfections were performed using the Neon Transfection system (ThermoFisher). Following the Cas9 RNP transfection, T cells were cultured in RPMI supplemented with 50 U/mL IL2 for 7 days.

**FACS Analysis of T cell Activation**

Seven days post-transfection, the cells were stimulated for a second time using αCD3/αCD28-precoated plates in the presence of IL2 (100 U/mL) for 24 h. Cell surface staining for T cell activation was performed using αCD25-PE (Biolegend) and αCD69-APC (Biolegend) antibodies for 1 h at 4 °C. Viable cells gating was performed using eBioscience™ Fixable Viability Dye eFluor™ 780 (ThermoFisher).

MOLECULAR MODELING

**Description of the methods**

In order to gain structural insights on the systems considered, we applied two different docking methods based on the software Autodock (Morris et al., 2009): the reactive docking and the flexible side chain covalent docking. The reactive docking is a predictive method that allows to identify the residues most likely to be modified by covalent binding. This is accomplished in two steps: first, it performs a scanning of all solvent accessible residues of a given type (cysteines, in this case), then it applies conventional, untethered docking with a special potential to simulate the incipient reaction to identify the most likely ones to be modified by the ligands. Reactive docking was successfully applied in previous studies, where it was used to model electrophile reactions with cysteine (Backus et al., 2016), tyrosine and lysine (Mortenson et al., 2018), and serine (Qinheng et al., 2019) residues. The flexible side chain covalent docking performs simulations in which the ligand is already attached to the covalent residue (via the newly formed covalent bond) and both ligand and residue are modeled as flexible.(Bianco et al., 2016) This method is used to analyze the non-covalent interactions of the bound ligands and target residues constituting the binding site.

### Reactive docking and flexible side chain covalent docking on MYD88

isoTOP-ABPP and TMT-ABPP experiments showed that the TIR domain of MYD88 is covalently modified with different potency by BPK-25 and BPK-21 at C203, and by KB02 and KB05 at either C274 or C280 within the tryptic peptide (270-282). Consequently, we applied two different docking techniques to rationalize the different potencies of the first compounds on C203, and to attempt resolving the ambiguity between C274 and C280 modification. In the first approach, we used the reactive docking method to sort the ambiguity between the labeling of the C274 and C280, then the flexible side chain covalent docking was used to generate putative binding mode of all the compounds and provide structural insight for their different activities.

We performed reactive docking simulations on the entire domain (PDB 4DOM) with ligands KB02 and KB05 and in addition, with BPK-25 and BPK-21 as a proof of concept, since experimental studies show direct labeling of C203 with BPK-25, but not BPK-21. Reactive docking analysis on BPK-25 and BPK-21 confirmed that the most favorable residue is C203, while C274 is the predicted residue for the covalent binding of KB02 and KB05. These results show that the position of C280 on the protein surface is less likely to be modified because it is largely solvent exposed, while on the contrary, C274 is located inside a cleft of the domain. Flexible side chain covalent docking was then used to refine the binding mode of compounds BPK-25 and BPK-21 (**Figure 5G**). In particular, the predicted binding mode of BPK-25 shows that it could bind by establishing two hydrogen bonds with R188 and E183 via the amide moiety (**Figure 5G, top right**), which is missing in BPK-21 (**Figure 5G, bottom right**). The lack of these interactions justifies the lower efficacy reported for BPK-21 in modifying C203.

### Flexible side chain covalent docking on ERCC3

isoTOP-ABPP and TMT-ABPP demonstrated that compounds BPK-25 and BPK-21 bind to ERCC3 by alkylating C342. Flexible side chain docking simulations were performed to rationalize the higher efficacy of BPK-21 with respect to BPK-25, by modeling the two ligands bound to C342 on a low resolution Cryo-EM structure of the protein (PDB 5OF4, 4.4 Å resolution). Results showed that BPK-21 can form two hydrogen bonds with T469 and Q497 side chains, while its central aromatic ring establishes a π□π interaction with W493 (**Figure 5H, bottom right**). None of these interactions are possible for BPK-25 (**Figure 5H, top right**), which is reflected in a lower docking score.

### Flexible side chain covalent docking on TMEM173

In order to provide a structural insight of the direct labeling of C91 of the protein TMEM173 with ligands BPK-21 and BPK-25, we applied flexible side chain covalent docking method on a low-resolution Cryo-EM structure (PDB 6NT5, res 4.1 Å). Results show that both BPK-21 and BPK-25 engage the pocket (roughly delimited by residues L98 and P141) by placing their aromatic rings: 2,3-dichlorobenzene and chloropyridine for BPK-21 and BPK-25 (**Figure S5B**), respectively, in proximity to P141. Despite their structural differences, both ligands occupy the pocket by establishing mostly hydrophobic interactions in a very similar manner, in agreement with the comparable reported efficacy.

### General Methods

The crystal structures of the proteins were retrieved from the Protein Data Bank: TIR domain of MYD88 (PDB 4DOM), ERCC3 (PDB 5OF4), TMEM173 (PDB 6NT5). Hydrogens were added with Reduce (Word et al., 1999), then were prepared using AutoDockTools (Morris et al., 2009) following the standard AutoDock protocol.(Forli et al., 2016) Reactive docking was performed following the protocol reported previously.(Backus et al., 2016) A grid box was defined for each cysteine: C168, C192, C216, C203, C247 and C280 (size x: 60, y: 60, z: 60 points).

For the flexible side chain covalent method, ligands were modelled attached to the alkylated residue via covalent bond, then processed following the covalent docking protocol (Bianco et al., 2016) (available online at http://autodock.scripps.edu/resources/covalentdocking) to be modeled as flexible during the docking. All dockings were performed using AutoDock 4.2.6 (Morris et al., 2009), generating 100 poses using the default LGA parameters. Poses with the best energy score were selected and analyzed. Figures were generated using Pymol.(The PyMOL Molecular Graphics System, Version 2.0 Schrödinger, LLC.)

GENERATION OF REFERENCE PROTEIN TABLES

### Immune-relevant genes

**Immune-enriched genes** were identified by analyzing microarray data from BioGPS (U133A and MOE430 datasets for human and mouse, respectively) and RNASeq data from GTex (release V7). Briefly, data were first filtered to restrict analyses to microarray signals above 150 and median RPKM values above 10. Samples from each transcriptomic dataset were grouped to identify immune related cells and tissues. Within each group the highest-expressing sample was chosen and group-level values were converted to Z-scores to identify genes showing immune enrichment within each dataset. Immune-enriched Z-scores above 3 or 4 (for RNA-Seq or microarray data, respectively) were summed across all probes and datasets and the summed Z-score was used to rank-order all genes. There were ~2004 genes with a summed Z-score above 11.0 and these were defined as "immune-enriched" as these represented the approximately 10% most-immune-enriched genes in the genome.

**Genes with immune-related phenotypes** were identified by parsing data in the Online Mendelian Inheritance of Man (OMIM) database (https://www.omim.org). OMIM associations were extracted from the human UniProt database downloaded in February 2019. From the 3925 genes for which human phenotypic associations could be identified, 655 genes with immune-related phenotypes were selected by querying phenotype titles and descriptions for immune-related substrings ('*immun*', '*inflam*', '*rheum*', '*psoria*', etc).

**Immune-relevant genes** were defined as those genes that were immune-enriched and/or associated with immune-related phenotypes, as described above. In total there were 2476 genes that met this criteria (column U in table S3).

### T cell proliferation gene list (SLICE) (Shifrut et al., 2018)

Hits (genes with FDR < 0.2 and |Z score| > 2, authors' criteria) were taken from the manuscript by Shifrut et al.(Shifrut et al., 2018) and cross-referenced with **Supplementrary Table S5** based on UniProt accessions.

### Immune module lists (Rieckmann et al., 2017)

Genes corresponding to immune modules were taken from Supplemental Table S2 from the manuscript by Rieckmann et al.(Rieckmann et al., 2017) and cross-referenced with **Supplementrary Table S5** based on UniProt accessions.

### Transcription Factors

The list of putative transcription factors was adapted from the GSEA website (http://software.broadinstitute.org/gsea/msigdb/gene_families.jsp). (Messina et al., 2004)

### Adaptors and Scaffolding Proteins

To generate a list of putative adaptor and scaffolding proteins we combined data from several different sources including GO,(Ashburner et al., 2000; The Gene Ontology, 2019) Uniprot,(UniProt, 2019) the scaffold protein database ScaPD,(Han et al., 2017) manual literature review, and a reagent list from R&D Biosystems (Adaptor Proteins Research Areas: R&D Systems https://www.rndsystems.com/research-area/adaptor-proteins (accessed Sep 4, 2019)). Proteins associated with following GO terms were included: GO:0035591 (signaling adaptor activity), GO:0060090 (molecular adaptor activity), GO:0008093 (cytoskeletal adaptor activity), GO:0035615 (clathrin adaptor activity). Lists of proteins for these GO terms were downloaded from the Gene Ontology project website using the AmiGO tool (http://amigo.geneontology.org/amigo/; version 2.5.12)(Carbon et al., 2009) with filters requiring that entries were of the type "protein" belonging to the "Homo sapiens" organism. Additionally, a keyword search was performed on a downloaded copy of SwissProt human data from Uniprot. Data was queried using BioPython (Cock et al., 2009) and the following search terms were used: "adapter", "adaptor", and "scaffold" and the search was performed against the following columns: comments prefixed with

“FUNCTION”, associated GO term cross-references, keywords, and entry descriptions. The data that was used from the ScaPD database consisted of experimentally verified scaffold proteins.
The resulting list was then cross-referenced with **Supplementrary Table S5**, and the categorization of every target protein was reviewed manually.

**SwissPalm list**

SwissPalm proteins and sites (Release 2 [02/18/2018])(Blanc et al., 2015) were downloaded and cross-referenced with the **Supplementrary Table S5** based on UniProt accessions. Sites were deemed a match if any of the cysteine residues in the detected tryptic peptide matched the SwissPalm reference (**Supplementrary Table S18**). These lists are likely an underrepresentation of the full extent of palmitoylated proteins (especially in immune cells) and this remains an active area of research.

### (D) Synthetic Methods

GENERAL ANALYTICAL INFORMATION

**KB63**, **BPK-5**, **BPK-7 — BPK-11**, **BPK-16**, **BPK-18 — BPK-22**, **BPK-25**, **BPK-29 — BPK-31**, **BPK-34**, **HS58A-C2**, **HS77**, **HS81C**, **HS81E**, **HS92**, **HS95 — HS98**, **HS125**, **HS126**, **HS145**, **HS175**, **HS177**, **HS178**, and **RS004** were previously characterized (Backus et al., 2016; Bar-Peled et al., 2017; Lee et al., 2018). All novel compounds were characterized using $^1$H NMR and HRMS. $^1$H NMR spectra can be found at the end of the Supporting Information. NMR spectra were recorded on a Bruker 400 MHz instrument. All $^1$H NMR experiments are reported in δ units, parts per million (ppm) and are listed relative to residual signals for DMSO (2.50 ppm), $CHCl_3$ (7.26 ppm), MeOH (3.31 ppm), $CH_3CN$ (1.94 ppm), or $H_2O$ (4.87 ppm) in the deuterated solvent. Reactions were monitored by LCMS. HRMS analyses were performed using an Agilent ESI-TOF instrument and were required to be within 5 ppm error.

**List of Abbreviations**

| | |
|---|---|
| **TEA** | triethylamine |
| **DIPEA** | *N*,*N*-diisopropylethylamine |
| **HATU** | *N*-[(Dimethylamino)-1H-1,2,3-triazolo-[4,5-*b*]pyridin-1-ylmethylene]-*N*-methylmethanaminium hexafluorophosphate *N*-oxide |
| **HOBt** | 1-Hydroxybenzotriazole hydrate |
| **EDCI** | *N*-(3-Dimethylaminopropyl)-*N'*-ethylcarbodiimide hydrochloride |
| **BOP** | (Benzotriazol-1-yloxy)tris(dimethylamino)phosphonium hexafluorophosphate |
| **TMSCl** | chlorotrimethylsilane |
| **$Boc_2O$** | Di-*tert*-butyl dicarbonate |
| **MsCl** | methanesulfonyl chloride |
| **NBS** | *N*-bromosuccinamide |
| **DCDMH** | 1,3-Dichloro-5,5-dimethylhydantoin |
| **$Ac_2O$** | acetic anhydride |
| **DAST** | diethylaminosulfur trifluoride |
| **HOAc** | glacial acetic acid |
| **TFA** | trifluoroacetic acid |
| **HCl** | hydrochloric acid |
| **FA** | formic acid |
| **DCM** | dichloromethane |
| **DMF** | *N*,*N*-dimethylformamide |
| **EtOAc** | ethyl acetate |
| **MeCN** | acetonitrile |

| | |
|---|---|
| **MeOH** | methanol |
| **THF** | tetrahydrofuran |
| **PE** | petroleum ether |
| **EtOH** | ethanol |
| **DCE** | 1,2-dichloroethane |
| **Xantphos** | 4,5-Bis(diphenylphosphino)-9,9-dimethylxanthene |
| **$Pd_2(dba)_3$** | Tris(dibenzylideneacetone)dipalladium(0) |
| **$Pd(dppf)Cl_2$** | 1,1′-Bis(diphenylphosphino)ferrocenedichloropalladium(II) |
| **$Ni(dppp)Cl_2$** | [1,3-Bis(diphenylphosphino)propane]dichloronickel(II) |
| **TLC** | thin-layer chromatography |

**General conditions for preparative HPLC (prep-HPLC)**

**HCl conditions**
*Column*: Phenomenex Synergi C18 150x25 mm,10 μm
*Mobile Phase*: solvent A – water (0.05% HCl), solvent B – MeCN
*Gradient*: 36-56% solvent B, 7.8 min

**FA conditions**
*Column*: Phenomenex Synergi C18 150x30 mm, 4 μm
*Mobile Phase*: solvent A – water (0.225% FA), solvent B – MeCN
*Gradient*: 40-70% solvent B, 12 min

**TFA conditions**
*Column*: Phenomenex Synergi C18 150x25 mm,10 μm
*Mobile Phase*: solvent A – water (0.1% TFA), solvent B – MeCN
*Gradient*: 40-70% solvent B, 10 min

**Basic**

*Column*: Phenomenex Gemini C18 150x25 mm,10 μm
*Mobile Phase*: solvent A – water (0.05% $NH_4OH$ v/v), solvent B – MeCN
*Gradient*: 58-80% solvent B, 10 min

OR

*Column*: Agela Durashell C18 150x25 mm, 5 μm
*Mobile Phase*: solvent A – water (0.05% $NH_4OH$ v/v), solvent B – MeCN
*Gradient*: 30-60% solvent B, 10 min

SYNTHETIC PROCEDURES

**General Procedures**

**General Procedure 1:** Chloroacetyl chloride (varying eq) was added to a solution of amine (1.0 eq) and TEA (varying eq) in anhydrous DCM (0.1 M) at 0 °C. The reaction was warmed to room temperature and stirred until the starting material could not be detected via TLC, generally 2-4 h. Water was added to the reaction (2x the volume of DCM) and the reaction was extracted with DCM (2x). The combined organic layers were washed with brine (1x), dried over anhydrous $Na_2SO_4$, filtered and concentrated under reduced pressure. The resulting residue was purified as indicated.

**General Procedure 2:** Acryloyl chloride (varying eq) was added to a solution of amine (1.0 eq) and TEA (varying eq) in anhydrous DCM (0.1 M) at 0 °C. The reaction was warmed to 15 °C and stirred for 1 h. Water was added to the reaction (2x the volume of DCM) and the reaction was extracted with DCM. The combined organic layers were washed with brine (1x), dried over anhydrous $Na_2SO_4$, filtered and concentrated under reduced pressure. The resulting residue was purified as indicated.

**General Procedure 3:** HATU (1.5 *eq*) and DIPEA (3.0 *eq*) were added to a solution of the carboxylic acid (1.1 eq) in DMF (0.1 M) and the mixture was stirred for 10 min. The amine (1.0 *eq)* was added to the mixture at 0 °C. The reaction was warmed to room temperature and stirred for 3 h. The reaction was quenched with water (equal volume to DMF) and extracted with EtOAc (3x). The combined organic phases were washed with brine (2x), dried over anhydrous $Na_2SO_4$ and concentrated. The resulting residue was purified as indicated.

**General Procedure 4:** A solution of acid chloride (2.0 eq) in DCM (0.1 M) was added dropwise to a solution of amine (1.0 eq) and DIPEA (3.0 eq) in DCM (0.1 M) at 0 °C. The reaction was then stirred for 3 h at 0 °C. Water (equal volume to DCM) was added to the reaction mixture and the aqueous layer was extracted with DCM (2x). The combined organic phases were washed with brine (1x), dried over anhydrous $Na_2SO_4$, and concentrated under reduced pressure. The resulting residue was purified as indicated.

**General Procedure 5:** HOAc (2.0 eq) and $NaBH_3CN$ (4.0 eq) were added to a stirred solution of ketone or aldehyde (1.0 eq) and amine (1.0 eq) in anhydrous MeCN (0.9 M) at 15 °C. The reaction mixture was heated to 50 °C and stirred for 8 h. The reaction was then cooled, water (equal volume to MeCN) was added and the mixture was extracted with EtOAc (3x). The organic layers were washed with brine (1x), dried over anhydrous $Na_2SO_4$, filtered and concentrated under reduced pressure. The resulting residue was used in the next step or purified as indicated.

**Synthesis of SI-4 as a precursor for EV-1 and EV-2**

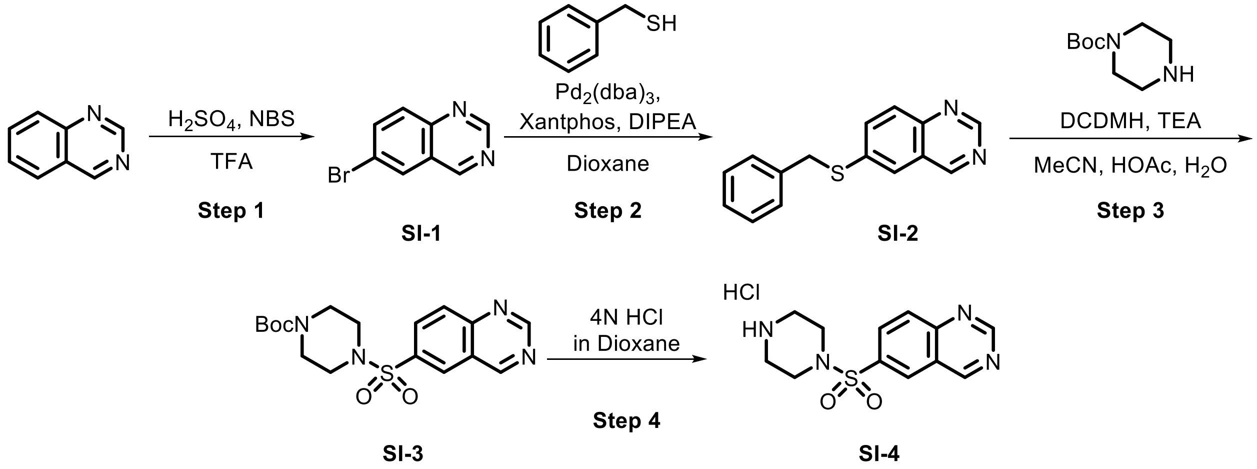

**Step 1:** Under an atmosphere of nitrogen, $H_2SO_4$ (8 mL) was added to a solution of quinazoline (5.0 g, 38.4 mmol, 1.0 eq) in anhydrous TFA (20 mL) at 15 °C. NBS (9.2 g, 51.9 mmol, 1.4 eq) was then added to the reaction mixture. The reaction was stirred at 15 °C for 8 h. The reaction mixture was poured into water (100 mL) and aqueous $Na_2CO_3$ was added until pH 8. The resulting solution was extracted with EtOAc (40 mL x 2). The combined organic extracts were washed with brine (10 mL x 2), dried over anhydrous $Na_2SO_4$, filtered and concentrated under reduced pressure. The residue was purified by preparative TLC (PE:EtOAc = 20:1) to afford **SI-1** (3.3 g, 41%) as a yellow solid.

**Step 2:** A two-neck round-bottom flask was charged with **SI-1** (2.0 g, 9.6 mmol, 1.0 eq), Xantphos (277 mg, 0.48 mmol, 0.05 eq), $Pd_2(dba)_3$ (263 mg, 0.29 mmol, 0.03 eq), DIPEA (3.3 mL, 19.1 mmol, 2.0 eq), and 1,4-dioxane (30 mL) under a nitrogen atmosphere. The flask was fitted with a reflux condenser and placed in an 80 °C heating bath. After 10 min, benzyl mercaptan (1.14 mL, 9.8 mmol, 1.0 eq.) was added dropwise to the mixture. After an additional 20 min, the mixture was cooled and the reaction mixture was concentrated under reduced pressure. The resulting residue was purified by flash chromatography (PE:EtOAc = 20:1 to 5:1) to afford **SI-2** (2.0 g, 83%) as a yellow solid.

**Step 3:** DCDMH (1.6 g, 7.9 mmol, 2.0 eq) was added to a solution of **SI-2** (1.0 g, 4.0 mmol, 1.0 eq) in $H_2O$ (33 μL), AcOH (66 μL) and MeCN (10 mL) at 0 °C. After 30 min, TEA (2.2 mL, 15.8 mmol, 4.0 eq) was added to the reaction mixture and then stirred for 30 min at 0 °C. *Tert*-butyl piperazine-1-carboxylate (738 mg, 3.96 mmol, 1.00 eq) was then added and the mixture was stirred at 15 °C for 1h. The mixture was concentrated under reduced pressure and the resulting residue was purified by flash chromatography (PE:EtOAc = 10:1 to 1:1) to afford **SI-3** (1.0 g, 66%) as a yellow solid.

**Step 4:** HCl/dioxane (4 N, 1.7 mL, 5.0 eq) was added to a solution of **SI-3** (500 mg, 1.3 mmol, 1.0 eq) in dioxane (3 mL) at 15 °C. The mixture was stirred for 1 h and then concentrated under reduced pressure to afford **SI-4** (400 mg, crude) as a yellow solid. **SI-4** was used in the next step without purification.

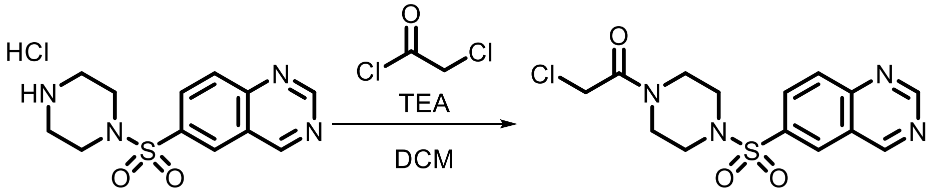

**SI-4** **EV-1**

Following **General Procedure 1**, starting from **SI-4** (200 mg, 0.64 mmol, 1.0 eq), chloroacetyl chloride (1.0 eq), and TEA (4.0 eq), **EV-1** was obtained after prep-HPLC (HCl) as an off-white solid (30 mg, 13%).

**$^1$H NMR (400 MHz, MeOD) δ** 8.73 (s, 1H), 8.00 (d, *J* = 2.0 Hz, 1H), 7.97 (dd, *J* = 8.4, 2.1 Hz, 1H), 7.53 (d, *J* = 8.4 Hz, 1H), 6.27 (s, 1H), 4.24 – 4.18 (m, 2H), 3.74-3.62 (m, 4H), 3.23 – 3.02 (m, 4H).

**HRMS ESI-TOF (*m/z*):** $[M+H]^+$ for $C_{14}H_{16}ClN_4O_3S$ 355.0626, found 355.0629.

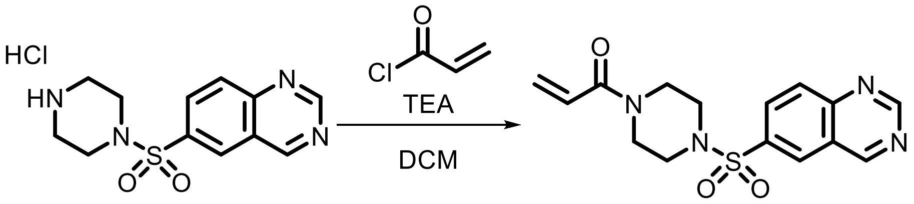

**SI-4** **EV-2**

Following **General Procedure 2**, starting from **SI-4** (200 mg, 0.64 mmol, 1.0 eq), acryloyl chloride (1.0 eq), and TEA (4.0 eq), **EV-2** was obtained after prep-HPLC (HCl) as an off-white solid (20 mg, 9%).

**$^{1}$H NMR (400 MHz, $CDCl_3$) δ** 9.56 (s, 1H), 9.49 (s, 1H), 8.44 (d, *J* = 2.0 Hz, 1H), 8.25 – 8.15 (m, 2H), 6.44 (dd, *J* = 16.8, 10.5 Hz, 1H), 6.23 (dd, *J* = 16.8, 1.8 Hz, 1H), 5.69 (dd, *J* = 10.5, 1.8 Hz, 1H), 3.74 (d, *J* = 45.3 Hz, 4H), 3.14 (t, *J* = 5.1 Hz, 4H).

**HRMS ESI-TOF (*m/z*):** $[M+H]^+$ for $C_{15}H_{17}N_4O_3S$ 333.1016, found 333.1021.

**Synthesis of SI-7 as a precursor for EV-3 and EV-4**

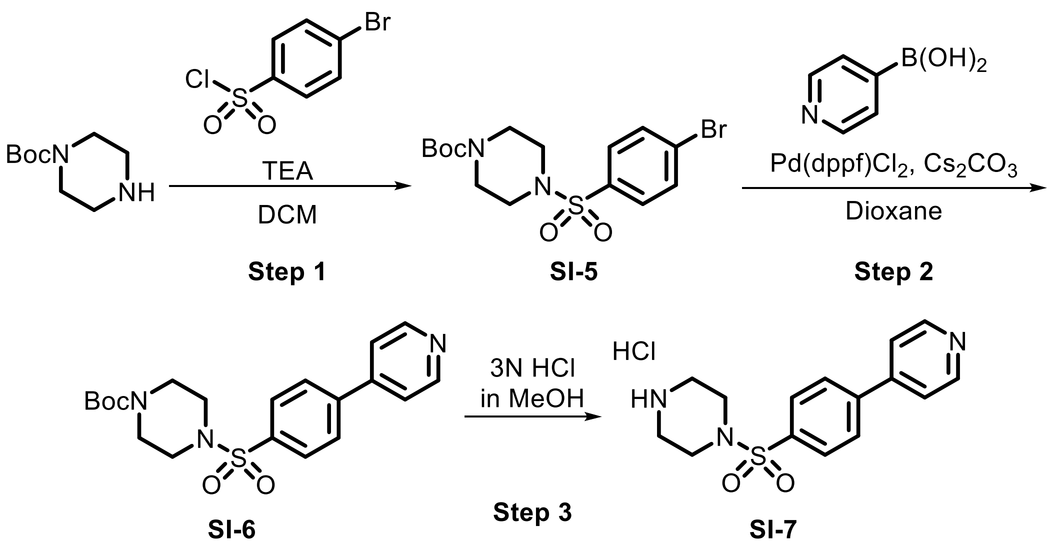

**Step 1:** TEA (1.5 mL, 10.7 mmol, 2.0 eq) was added to a mixture of *tert*-butyl piperazine-1-carboxylate (1.0 g, 5.4 mmol, 1.0 eq) and 4-bromobenzenesulfonyl chloride (1.4 g, 5.4 mmol, 1.0 eq) in DCM (30 mL) at 15 °C under a nitrogen atmosphere and stirred for 3 h. The reaction mixture was concentrated under reduced pressure to afford **SI-5** as a white solid (2.1 g, crude) which was used without purification.

**Step 2:** Under an atmosphere of nitrogen, Pd(dppf)$Cl_2$ (180 mg, 0.25 mmol, 0.1 eq) was added to a mixture of **SI-5** (1.0 g, 2.5 mmol, 1.0 eq), $Cs_2CO_3$ (1.6 g, 4.9 mmol, 2.0 eq) and 4-pyridylboronic acid (455 mg, 3.7 mmol, 1.5 eq) in dioxane (10 mL) at 15 °C. The reaction mixture was then heated to 110 °C and stirred for 4 h. After cooling, the reaction mixture was diluted with water (30 mL) and extracted with EtOAc (40 mL x 2). The combined organic phases were washed with brine (10 mL x 2), dried over anhydrous $Na_2SO_4$, filtered and concentrated under reduced pressure. The crude product was purified by flash chromatography (PE:EtOAc = 50:1 to 2:1) to afford **SI-6** (800 mg, 80%) as white solid.

**Step 3:** 3N HCl in MeOH (20 mL) was added to **SI-6** (800 mg, 2.0 mmol, 1.0 eq) and the reaction mixture was stirred at 15 °C for 2 h. The reaction mixture was concentrated under reduced pressure to provide **SI-7** (600 mg, crude) as a white solid which was used without purification.

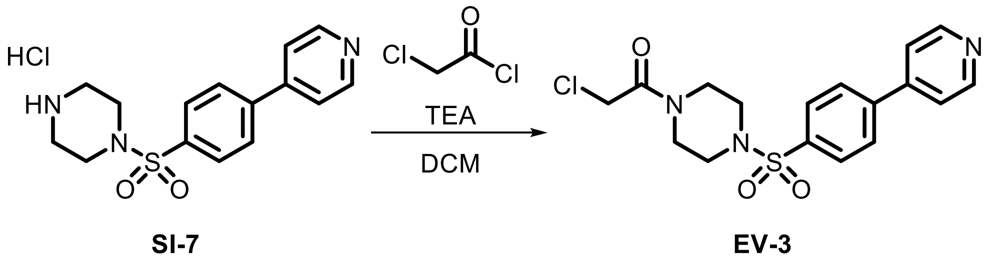

Following **General Procedure 1**, starting from **SI-7** (200 mg, 0.59 mmol, 1.0 eq), chloroacetyl chloride (3.0 eq) and TEA (2.0 eq), **EV-3** was obtained after prep-HPLC (HCl) as a white solid (43 mg, 17%).

**$^1$H NMR (400 MHz, $D_2O$) δ** 8.87 – 8.82 (m, 2H), 8.37 – 8.32 (m, 2H), 8.11 (d, *J* = 8.6 Hz, 2H), 7.99 (d, *J* = 8.6 Hz, 2H), 4.25 (s, 2H), 3.66 (t, *J* = 5.2 Hz, 4H), 3.17 (dt, *J* = 21.5, 5.2 Hz, 4H).

**HRMS ESI-TOF (*m/z*):** $[M+H]^+$ for $C_{17}H_{19}ClN_3O_3S$ 380.0830, found 380.0833.

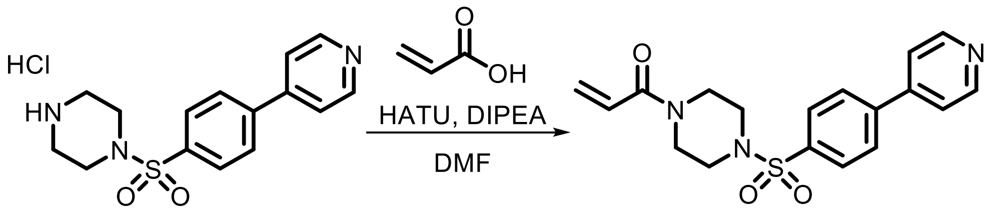

**SI-7** **EV-4**

Following **General Procedure 3**, starting with **SI-7** (200 mg, 0.66 mmol, 1.0 eq) and acrylic acid (4.0 eq), **EV-4** was obtained after prep-HPLC (basic) as a white solid (13 mg, 5%).

**$^1$H NMR (400 MHz, CDCl$_3$) δ** 8.74 (d, *J* = 5.7 Hz, 2H), 7.89 – 7.75 (m, 4H), 7.54 – 7.47 (m, 2H), 6.47 (dd, *J* = 16.7, 10.5 Hz, 1H), 6.26 (dd, *J* = 16.8, 1.8 Hz, 1H), 5.70 (dd, *J* = 10.5, 1.8 Hz, 1H), 3.74 (d, *J* = 48.4 Hz, 4H), 3.08 (t, *J* = 5.1 Hz, 4H).

**HRMS ESI-TOF (*m/z*):** $[M+H]^+$ for $C_{18}H_{20}N_3O_3S$ 358.1220, found 358.1224.

**Synthesis of SI-11 as a precursor for EV-5 and EV-6**

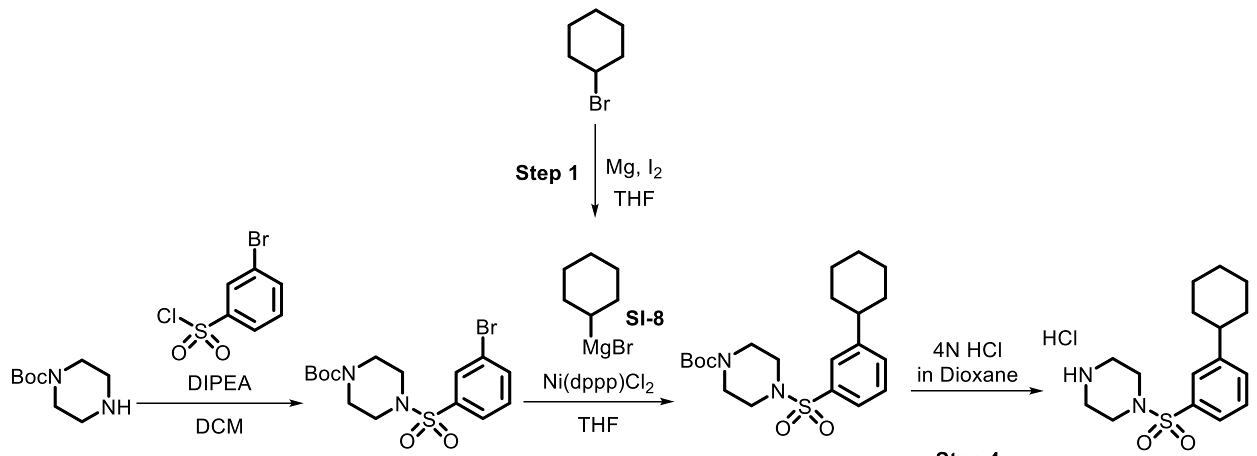

**Step 2** **SI-9** **Step 3** **SI-10** **Step 4** **SI-11**

**Step 1:** A suspension of Mg (447 mg, 18 mmol, 1.5 eq) in THF (20 mL) was treated with $I_2$ (31 mg, 0.12 mmol, 0.01 eq) and heated to 80 °C. A solution of bromocyclohexane (2.0 g, 12.3 mmol, 1.0 eq) in THF (15 mL) was added dropwise over 30 min. After addition, the resulting mixture was cooled to 20 °C and stirred for an additional 2 h. **SI-8** was used in the **Step 3** without purification.

**Step 2:** 2-bromopheylsulfonyl chloride (2.5 g, 9.8 mmol, 1.0 eq) was added to a solution of *tert*-butyl piperazine-1-carboxylate (1.7 g, 9.3 mmol, 0.95 eq) and DIPEA (2.1 mL, 11.7 mmol, 1.2 eq) in DCM (25 mL) at 0 °C. The mixture was warmed to 20 °C and stirred for 1 h. The mixture was poured into water (5 mL) and DCM (5 mL). The aqueous solution was extracted with DCM (5 mL x 2). The organic phases were combined, dried over anhydrous $Na_2SO_4$, and concentrated to obtain **SI-9** as a light yellow oil. **SI-9** was used in the next step without purification.

**Step 3: SI-8** (0.5 M, 20 mL, 4.0 eq) was added to a solution of **SI-9** (1.0 g, 2.5 mmol, 1.0 eq) and Ni(dppp)$Cl_2$ (134 mg, 0.25 mmol, 0.1 eq) in THF (20 mL) at 0 °C. After addition, the reaction mixture was warmed to 20 °C and stirred for an additional 2 h. The reaction was poured into water (5 mL) and EtOAc (5 mL). The aqueous solution was extracted with EtOAc (5 mL x 2). The combined organic phases were dried over anhydrous $Na_2SO_4$, and concentrated under reduced pressure. The resulting residue was purified by flash chromatography (PE:EtOAc = 25:1 to 5:1) to provide **SI-10** (340 mg, 0.7 mmol, 28%) as a white solid.

**Step 4:** HCl/dioxane (4N, 5.0 mL, 24 eq) was added to a solution of **SI-10** (340 mg, 0.82 mmol, 1.0 eq) in dioxane (5 mL) at 20 °C and stirred for 2 h. The reaction mixture was concentrated to provide **SI-11** (250 mg, crude) as a light yellow oil. **SI-11** was used for the synthesis of **EV-5** and **EV-6** without purification.

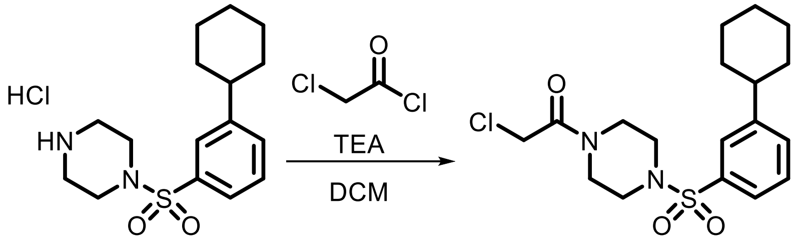

**SI-11** **EV-5**

Following **General Procedure 1**, starting with **SI-11** (50 mg, 0.14 mmol, 1.0 eq), chloroacetyl chloride (1.5 eq), and TEA (3.0 eq), **EV-5** was obtained after prep-HPLC (FA) as a white solid (15 mg, 24%).

**$^1$H NMR (400 MHz, $CDCl_3$) δ** 7.58 – 7.52 (m, 2H), 7.49 – 7.43 (m, 2H), 4.00 (s, 2H), 3.66 (dt, *J* = 39.6, 5.1 Hz, 4H), 3.04 (dt, *J* = 23.0, 5.1 Hz, 4H), 2.64 – 2.54 (m, 1H), 1.93 – 1.81 (m, 4H), 1.81 – 1.74 (m, 1H), 1.49 – 1.33 (m, 4H), 1.32 – 1.19 (m, 1H).

**HRMS ESI-TOF (*m/z*):** $[M+H]^+$ for $C_{18}H_{26}ClN_2O_3S$ 385.1347, found 385.1348.

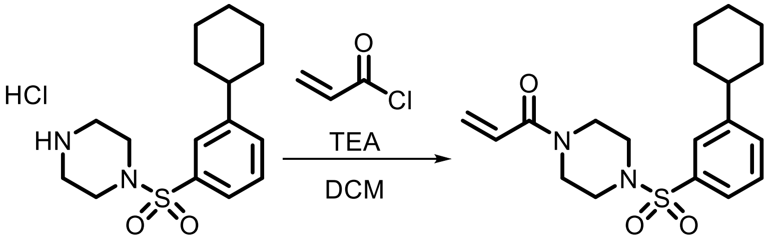

**SI-11** **EV-6**

Following **General Procedure 2**, starting with **SI-11** (150 mg, 0.43 mmol, 1.0 eq), acryloyl chloride (1.5 eq), and TEA (3.0 eq), **EV-6** was obtained after prep-TLC (PE:EtOAc = 1:1) as a white solid (49 mg, 30%).

**$^1$H NMR (400 MHz, $CDCl_3$) δ** 7.59 – 7.51 (m, 2H), 7.49 – 7.40 (m, 2H), 6.46 (dd, *J* = 16.8, 10.6 Hz, 1H), 6.25 (dd, *J* = 16.9, 1.9 Hz, 1H), 5.69 (dd, *J* = 10.5, 1.8 Hz, 1H), 3.71 (d, *J* = 48.6 Hz, 4H), 3.02 (t, *J* = 5.1 Hz, 4H), 2.58 (t, *J* = 10.2 Hz, 1H), 1.97 – 1.72 (m, 5H), 1.49 – 1.20 (m, 5H).

**HRMS ESI-TOF (*m/z*):** $[M+H]^+$ for $C_{19}H_{27}N_2O_3S$ 363.1737, found 363.1738.

**Synthesis of SI-12 as a precursor for EV-7 – EV-10**

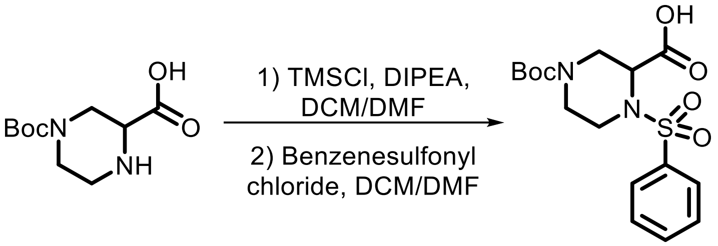

**SI-12**

Under an atmosphere of nitrogen, TMSCl (4.4 mL, 34.7 mmol, 1.6 eq) was added to a stirred solution of 4-Boc-piperazine-2-carboxylic acid (5.0 g, 21.7 mmol, 1.0 eq) and DIPEA (13.2 mL, 76.0 mmol, 3.5 eq) in anhydrous DCM (50 mL) and DMF (20 mL) at 15 °C. The reaction was stirred for 2 h followed by the addition of benzenesulfonyl chloride (3.0 mL, 23.9 mmol, 1.1 eq) and stirred for an additional 2 h. The reaction mixture was diluted with water (50 mL) and extracted with DCM (100 mL x 3). The combined organic phases were washed with 0.5 M HCl (50 mL x 3), dried over anhydrous $Na_2SO_4$, filtered, and concentrated under reduced pressure to give **SI-12** (3.5 g, crude) as a yellow oil which was used without purification.

**Synthesis of SI-14 as a precursor for EV-7 and EV-8**

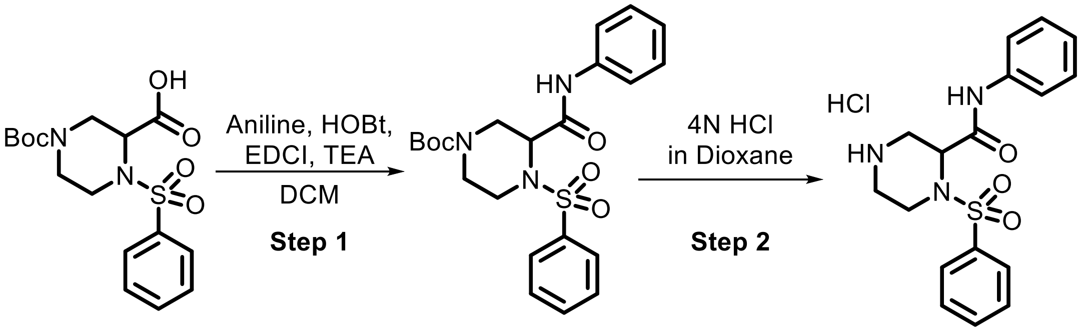

**SI-12** **SI-13** **SI-14**

**Step 1:** HOBt (87 mg, 0.65 mmol, 1.2 eq), EDCI (124 mg, 0.65 mmol, 1.2 eq), and aniline (49 µL, 0.54 mmol, 1.0 eq) were added to a solution of **SI-12** (200 mg, 0.54 mmol, 1.0 eq), and TEA (0.23 mL, 1.62 mmol, 3.0 eq) in anhydrous DCM (1 mL) and stirred for 8 h. Upon completion, 50 mL of water was added to the reaction mixture and then extracted with DCM (75 mL x 3). The combined organic extracts were washed with brine, dried over anhydrous $Na_2SO_4$, filtered, and concentrated under reduced pressure to afford **SI-13** (200 mg, crude) as yellow oil. **SI-13** was used in the next step without purification.

**Step 2:** HCl/dioxane (4 N, 0.84 mL, 10.0 eq) was added to a solution of **SI-13** (150 mg, 0.34 mmol, 1.0 eq) in anhydrous dioxane (1 mL) and stirred for 1h. Upon consumption of the starting material, the reaction was concentrated under reduced pressure to afford **SI-14** (110 mg, crude) as yellow oil which was used without purification.

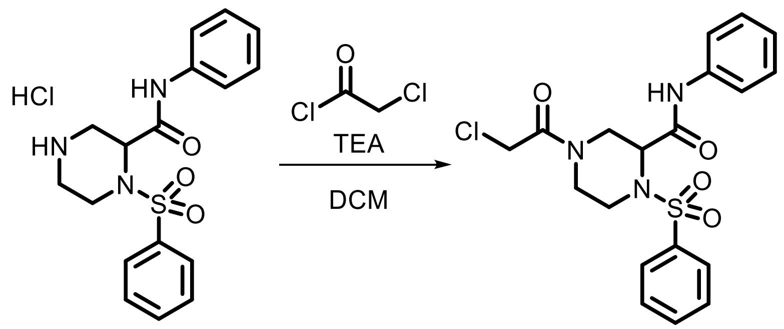

**SI-14** **EV-7**

Following **General Procedure 1**, starting with **SI-14** (50 mg, 0.13 mmol, 1.0 eq), chloroacetyl chloride (2.0 eq), and TEA (4.0 eq), **EV-7** was obtained after prep-HPLC (HCl) as a yellow solid (17 mg, 28%).

**$^1$H NMR (400 MHz, $CDCl_3$) δ** 8.33 (s, 1H), 7.94 (d, *J* = 7.3 Hz, 2H), 7.73 (t, *J* = 7.4 Hz, 1H), 7.65 (t, *J* = 7.7 Hz, 2H), 7.45 (d, *J* = 7.8 Hz, 2H), 7.35 (t, *J* = 7.8 Hz, 2H), 7.17 (t, *J* = 7.4 Hz, 1H), 4.57 (d, *J* = 12.7 Hz, 1H), 4.51 (d, *J* = 13.9 Hz, 1H), 4.35 (d, *J* = 13.6 Hz, 1H), 4.11 (d, *J* = 12.6 Hz, 1H), 4.02 (d, *J* = 14.5 Hz, 1H), 3.24 (ddd, *J* = 15.0, 12.4, 3.4 Hz, 2H), 2.87 (dd, *J* = 14.0, 4.1 Hz, 1H), 2.46 (td, *J* = 12.8, 3.3 Hz, 1H).

**HRMS ESI-TOF (*m/z*):** $[M+H]^+$ for $C_{19}H_{21}ClN_3O_4S$ 422.0936, found 422.0939.

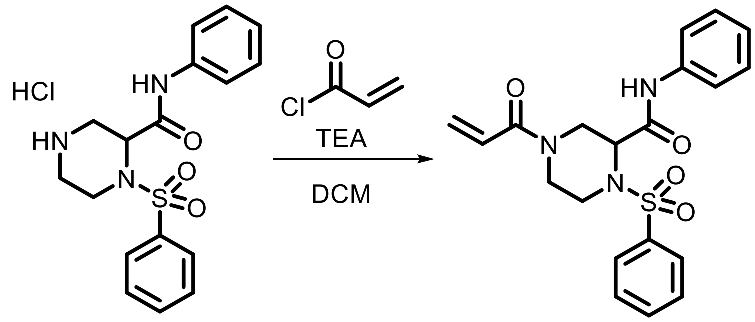

**SI-14** **EV-8**

Following **General Procedure 2**, starting with **SI-14** (50 mg, 0.13 mmol, 1.0 eq), acryloyl chloride (2.0 eq), and TEA (4.0 eq), **EV-8** was obtained after prep-HPLC (FA) as an off-white solid (21 mg, 41%).

**$^1$H NMR (400 MHz, $CDCl_3$) δ** 8.30 (s, 1H), 7.96 – 7.90 (m, 2H), 7.74 – 7.56 (m, 3H), 7.46 (d, *J* = 7.9 Hz, 2H), 7.33 (t, *J* = 7.8 Hz, 2H), 7.15 (t, *J* = 7.5 Hz, 1H), 6.86 – 6.74 (m, 1H), 6.25 (d, *J* = 16.7 Hz, 1H), 5.71 (dd, *J* = 10.5, 1.9 Hz, 1H), 4.73 – 4.55 (m, 2H), 4.37 (d, *J* = 13.6 Hz, 1H), 4.01 (d, *J* = 14.4 Hz, 1H), 3.31 (t, *J* = 13.1 Hz, 1H), 2.80 (d, *J* = 13.8 Hz, 1H), 2.49 (m, 1H).

**HRMS ESI-TOF (*m/z*):** $[M+H]^+$ for $C_{20}H_{22}N_3O_4S$ 400.1326, found 400.1329.

**Synthesis of SI-16 as a precursor for EV-9 and EV-10**

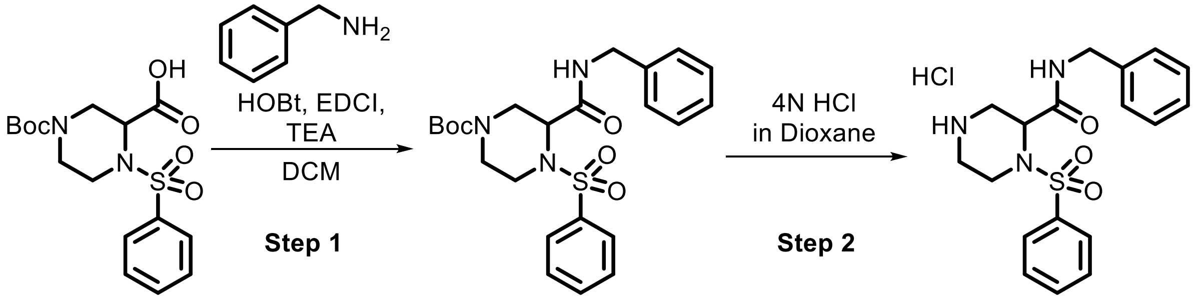

**SI-12** **SI-15** **SI-16**

**Step 1:** HOBt (88 mg, 0.65 mmol, 1.2 eq), EDCI (124 mg, 0.65 mmol, 1.2 eq) and benzylamine (59 µL, 0.54 mmol, 1.0 eq) were added to a solution of **SI-12** (200 mg, 0.54 mmol, 1.0 eq) and TEA (226 µL, 1.6 mmol, 3.0 eq) in anhydrous DCM (1.0 mL) at 15 °C and then stirred for 8 h. 50 mL of water was added to the reaction mixture and then extracted with DCM (75 mL x 3). The combined organic phases were washed with brine (25 mL x 1), dried over anhydrous $Na_2SO_4$, filtered and concentrated under reduced pressure to afford **SI-15** (250 mg, crude) as a yellow oil which was used without purification.

**Step 2:** HCl/dioxane (4 M, 1.4 mL, 10.0 eq) was added to a solution of **SI-15** (250 mg, 0.54 mmol, 1.0 eq) in anhydrous dioxane (1.0 mL) at 15 °C. The reaction was stirred for 1h. The reaction was then concentrated under reduced pressure to afford **SI-16** (200 mg, crude) as a yellow oil which was used without purification.

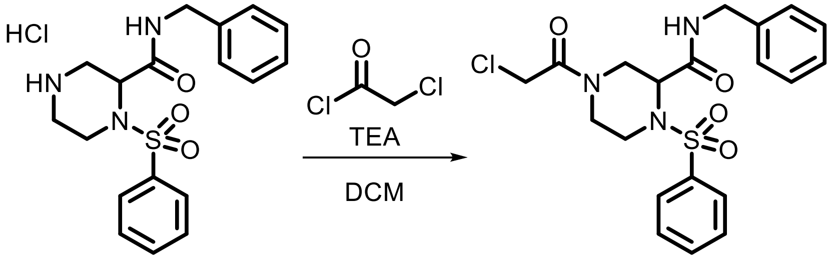

**SI-16** **EV-9**

Following **General Procedure 1**, starting with **SI-16** (100 mg, 0.25 mmol, 1.0 eq), chloroacetyl chloride (2.0 eq), and TEA (4.0 eq), **EV-9** was obtained after prep-HPLC (HCl) as an off-white solid (21 mg, 19%).

**$^1$H NMR (400 MHz, $CDCl_3$) δ** 7.88 – 7.83 (m, 2H), 7.70 – 7.64 (m, 1H), 7.57 (t, *J* = 7.7 Hz, 2H), 7.38 – 7.27 (m, 3H), 7.23 – 7.17 (m, 2H), 6.95 (br s, 1H), 4.53 – 4.42 (m, 5H), 4.28 (d, *J* = 14.4 Hz, 1H), 4.08 (d, *J* = 12.7 Hz, 1H), 3.90 (d, *J* = 14.5 Hz, 1H), 3.14 (ddd, *J* = 14.8, 12.1, 3.5 Hz, 1H), 2.83 (dd, *J* = 13.8, 4.0 Hz, 1H), 2.46 – 2.34 (m, 1H).

**HRMS ESI-TOF (*m/z*):** $[M+H]^+$ for $C_{20}H_{23}ClN_3O_4S$ 436.1093, found 436.1093.

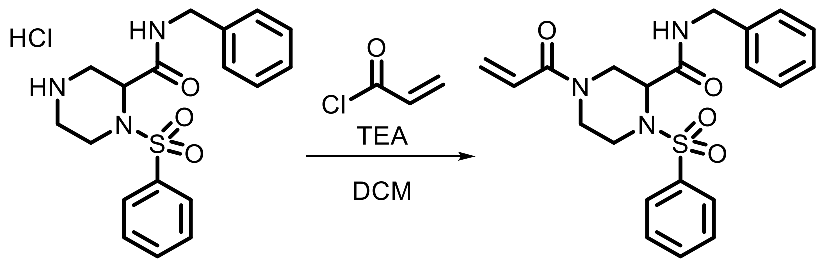

**SI-16** **EV-10**

Following **General Procedure 2**, starting with **SI-16** (100 mg, 0.26 mmol, 1.0 eq), acryloyl chloride (2.0 eq), and TEA (4.0 eq), **EV-10** was obtained after prep-HPLC (FA) as an off white solid (19 mg, 18%).

**$^1$H NMR (400 MHz, $CDCl_3$) δ** 7.89 – 7.82 (m, 2H), 7.64 (t, *J* = 7.4 Hz, 1H), 7.55 (t, *J* = 7.7 Hz, 2H), 7.36 – 7.25 (m, 3H), 7.23 – 7.16 (m, 2H), 6.90 (br s, 1H), 6.76 (dd, *J* = 16.7, 10.8 Hz, 1H), 6.25 (dd, *J* = 16.6, 1.8 Hz, 1H), 5.69 (dd, *J* = 10.5, 1.9 Hz, 1H), 4.64 (d, *J* = 13.8 Hz, 1H), 4.51 (s, 1H), 4.43 (dd, *J* = 5.9, 3.7 Hz, 2H), 4.32 (d, *J* = 13.7 Hz, 1H), 3.89 (d, *J* = 14.5 Hz, 1H), 3.20 (t, *J* = 13.3 Hz, 1H), 2.78 (d, *J* = 13.6 Hz, 1H), 2.44 (t, *J* = 12.6 Hz, 1H).

**HRMS ESI-TOF (*m/z*):** $[M+H]^+$ for $C_{21}H_{24}N_3O_4S$ 414.1482, found 414.1486.

**Synthesis of SI-19 as a precursor for EV-11 – EV-31**

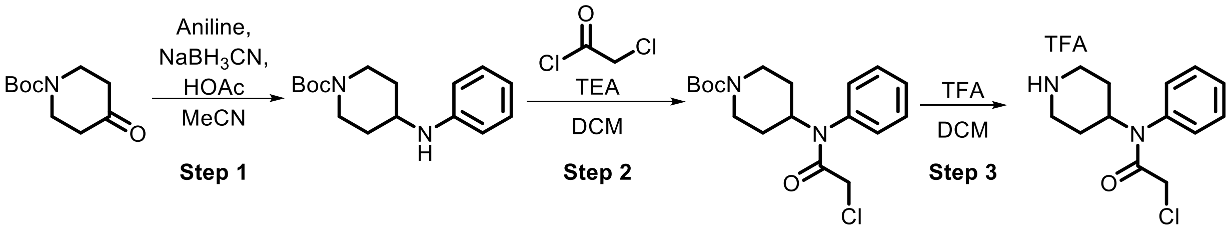

**SI-17** **SI-18** **SI-19**

**Step 1:** Following **General Procedure 5**, starting with 1-Boc-4-piperidone (19 g, 95 mmol, 1.0 eq) and aniline (8.7 mL, 95.4 mmol, 1.0 eq), **SI-17** (13.0 g, 49%) was obtained as a white solid after purification via prep-HPLC (basic).

**Step 2:** Following General Procedure 1, starting with **SI-17** (13.5 g, 48.9 mmol, 1.0 eq), chloroacetyl chloride (2.0 eq), and TEA (2.0 eq), **SI-18** (12.8 g, 74%) was obtained as a yellow solid after purification by prep-HPLC (basic).

**Step 3:** TFA (34.7 mL, 454 mmol, 10.0 eq) was added to a solution of **SI-18** (16.0 g, 45.4 mmol, 1.0 eq) in DCM (20 mL) at 18 °C and stirred for 3 h. Upon completion, the mixture was concentrated under reduced pressure to provide **SI-19** (23.0 g, crude) as a yellow oil which was used without purifiaction.

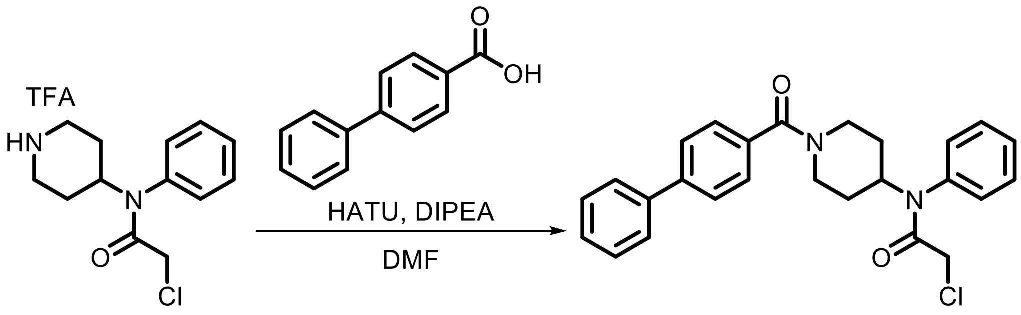

**SI-19** **EV-11**

Following **General Procedure 3**, starting with **SI-19** (100 mg, 0.27 mmol, 1.0 *eq)* and 4-phenylbenzoic acid (1.1 *eq*), **EV-11** was obtained after prep-HPLC (HCl) as a white solid (41 mg, 35%).

**$^1$H NMR (400 MHz, CDCl$_3$) δ** 7.61 – 7.52 (m, 4H), 7.51 – 7.41 (m, 5H), 7.41 – 7.33 (m, 3H), 7.14 (br s, 2H), 4.87 (m, 2H), 3.90 (br s, 1H), 3.71 (s, 2H), 3.18 (br s, 1H), 2.87 (br s, 1H), 2.03 – 1.78 (m, 2H), 1.49 – 1.19 (m, 2H).

**HRMS ESI-TOF (*m/z*):** [M+H]$^+$ for $C_{26}H_{26}ClN_2O_2$ 433.1678, found 433.1677.

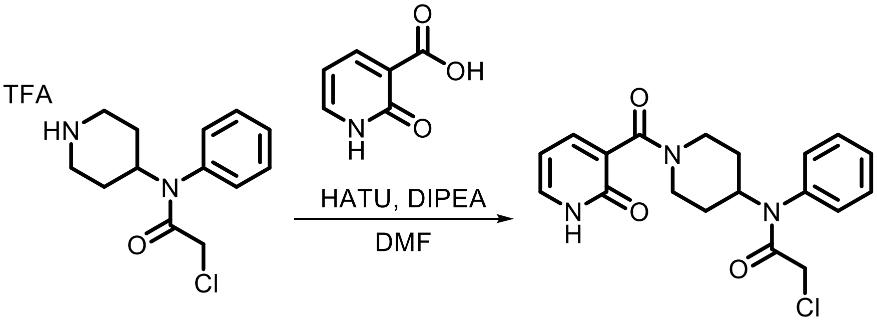

**SI-19** **EV-12**

Following **General Procedure 3**, starting with **SI-19** (150 mg, 0.41 mmol, 1.0 eq) and 2-hydroxynicotinic acid (1.1 eq), **EV-12** was obtained after prep-HPLC (HCl) as a solid (3 mg, 2%).

**$^1$H NMR (400 MHz, CDCl$_3$) δ** 7.70 (br s, 1H), 7.59 (br s, 1H), 7.46 (br s, 3H), 7.15 (br s, 2H), 6.60 (br s, 1H), 4.77 (br s, 1H), 3.71 (s, 2H), 2.95 (br s, 2H), 1.91 (br s, 2H), 1.48 (br s, 2H).

**HRMS ESI-TOF (*m/z*):** [M+H]$^+$ for $C_{19}H_{21}ClN_3O_2$ 374.1266, found 374.1266.

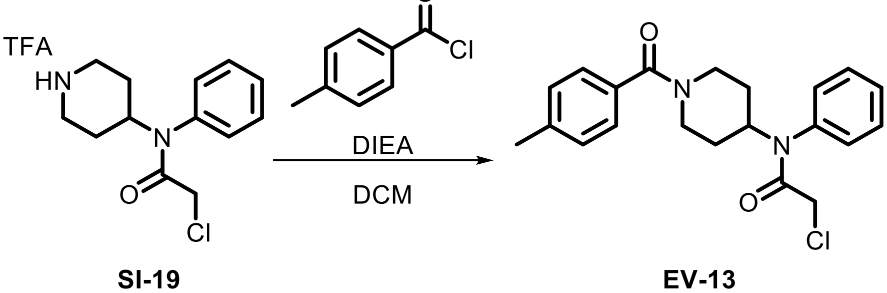

**SI-19** **EV-13**

Following **General Procedure 4**, starting with **SI-19** (150 mg, 0.41 mmol, 1.0 eq) and 4-methylbenzoyl chloride (2.0 eq), **EV-13** was obtained after prep-HPLC (HCl) as an off-white solid (66 mg, 44%).

**$^1$H NMR (400 MHz, CDCl$_3$) δ** 7.50 – 7.45 (m, 3H), 7.22 – 7.10 (m, 6H), 4.90-4.78 (m, 1H), 3.71 (s, 2H), 2.99 (br s, 2H), 2.35 (s, 3H), 1.95 – 1.77 (m, 4H), 1.37 – 1.25 (m, 2H).

**HRMS ESI-TOF (*m/z*):** $[M+H]^+$ for $C_{21}H_{24}ClN_2O_2$ 371.1521, found 371.1522.

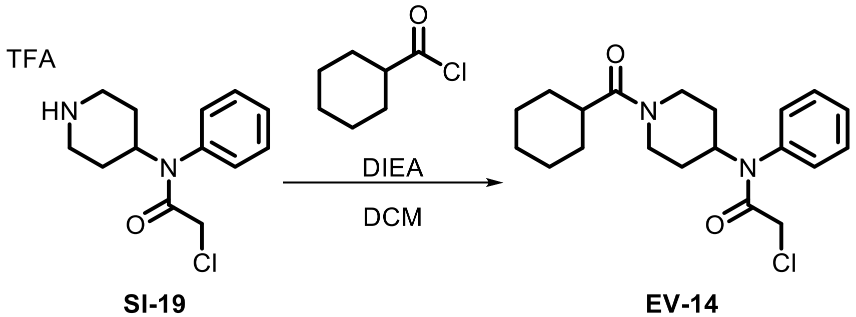

**SI-19** **EV-14**

Following **General Procedure 4**, starting with **SI-19** (150 mg, 0.41 mmol, 1.0 eq) and cyclohexylcarbonyl chloride (2.0 eq), **EV-14** was obtained after prep-HPLC (HCl) as an off-white solid (57 mg, 38%).

**$^1$H NMR (400 MHz, CDCl$_3$) δ** 7.50 – 7.42 (m, 3H), 7.12 (dd, *J* = 6.3, 3.0 Hz, 2H), 4.82 (t, *J* = 12.2 Hz, 1H), 4.32 (br s, 2H), 3.71 (s, 2H), 2.85 (br s, 2H), 2.38 (t, *J* = 11.9 Hz, 1H), 1.89 (d, *J* = 12.0 Hz, 2H), 1.77 (s, 3H), 1.69 – 1.56 (m, 1H), 1.51 – 1.34 (m, 2H), 1.29 – 1.10 (m, 4H).

**HRMS ESI-TOF (*m/z*):** $[M+H]^+$ for $C_{20}H_{28}ClN_2O_2$ 363.1834, found 363.1833.

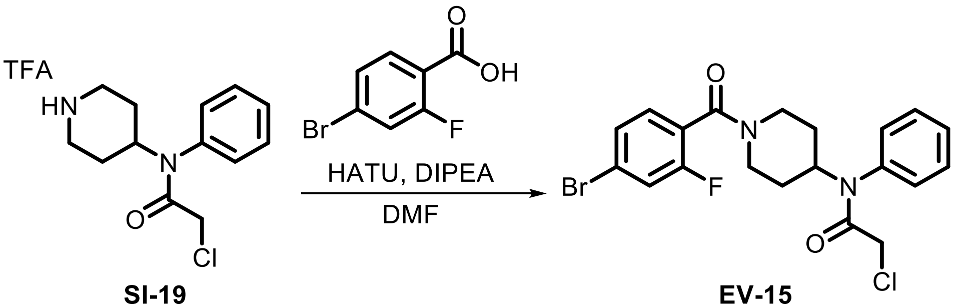

**SI-19** **EV-15**

Following **General Procedure 3**, starting with **SI-19** (150 mg, 0.41 mmol, 1.0 eq) and 4-bromo-2-fluorobenzoic acid (2.0 eq), **EV-15** (9 mg, 5%) was obtained after prep-HPLC (HCl).

**$^1$H NMR (400 MHz, CDCl$_3$) δ** 7.50 – 7.44 (m, 3H), 7.31 (dd, *J* = 8.1, 1.8 Hz, 1H), 7.24 (dd, *J* = 8.9, 1.7 Hz, 1H), 7.20 – 7.13 (m, 2H), 7.09 (br s, 1H), 4.93 – 4.72 (m, 2H), 3.71 (s, 2H), 3.59 – 3.43 (m, 1H), 3.18 (br s, 1H), 2.83 (t, *J* = 12.5 Hz, 1H), 1.89 (dd, *J* = 39.4, 12.6 Hz, 2H), 1.46 – 1.15 (m, 2H).

**HRMS ESI-TOF (*m/z*):** $[M+H]^+$ for $C_{20}H_{20}BrClFN_2O_2$ 453.0375, found 453.0374.

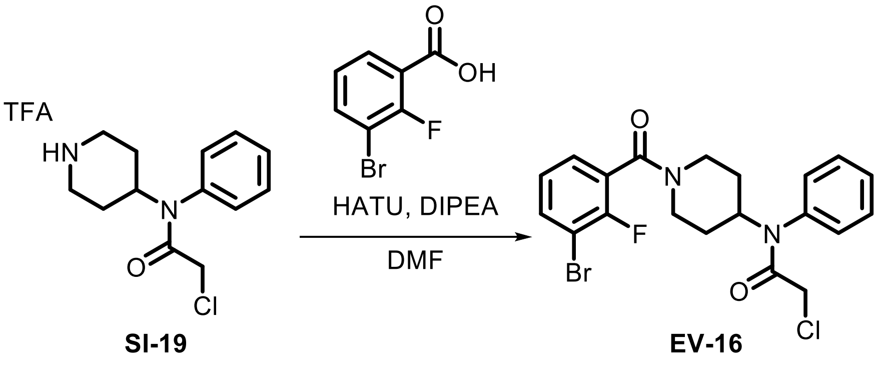

SI-19 EV-16

Following **General Procedure 3**, starting with **SI-19** (150 mg, 0.41 mmol, 1.0 eq) and 3-bromo-2-fluorobenzoic acid (2.0 eq), **EV-16** (27 mg, 15%) was obtained after prep-HPLC.

**$^1$H NMR (400 MHz, CDCl$_3$) δ** 7.57 (t, *J* = 7.4 Hz, 1H), 7.51 – 7.44 (m, 3H), 7.25 – 7.09 (m, 3H), 7.05 (t, *J* = 7.7 Hz, 1H), 4.91 – 4.72 (m, 2H), 3.71 (s, 2H), 3.60 – 3.45 (m, 1H), 3.18 (br s, 1H), 2.85 (s, 1H), 1.98 – 1.77 (m, 3H), 1.37 (qd, *J* = 12.8, 11.9, 3.9 Hz, 1H), 1.22 (br s, 1H).

**HRMS ESI-TOF (*m/z*):** $[M+H]^+$ for $C_{20}H_{20}BrClFN_2O_2$ 453.0375, found 453.0377.

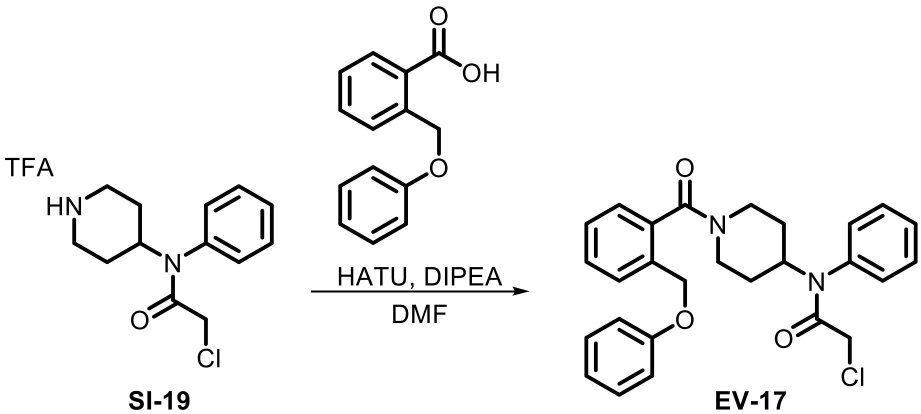

SI-19 EV-17

Following **General Procedure 3**, starting with **SI-19** (150 mg, 0.41 mmol, 1.0 eq) and 2-(phenoxymethyl)benzoic acid (2.0 eq), **EV-17** (37 mg, 19%) was obtained after prep-HPLC.

**$^1$H NMR (400 MHz, CDCl$_3$) δ** 7.50 (d, *J* = 7.5 Hz, 2H), 7.46 – 7.23 (m, 6H), 7.21 – 6.94 (m, 4H), 6.89 (s, 1H), 6.79 (s, 1H), 5.16 – 4.86 (m, 2H), 4.80 (s, 2H), 3.80 – 3.46 (m, 3H), 3.07 (d, *J* = 63.3 Hz, 1H), 2.76 (d, *J* = 50.2 Hz, 1H), 1.80 (s, 2H), 1.55 – 1.05 (m, 2H).

**HRMS ESI-TOF (*m/z*):** $[M+H]^+$ for $C_{27}H_{28}ClN_2O_3$ 463.1783, found 463.1785.

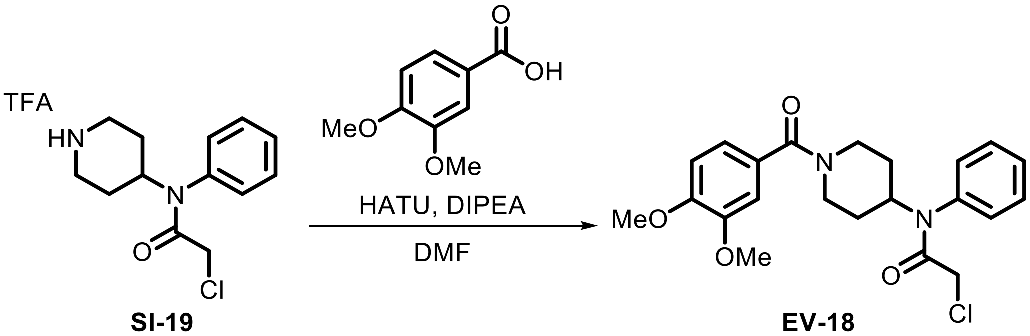

**SI-19** **EV-18**

Following **General Procedure 3**, starting with **SI-19** (150 mg, 0.41 mmol, 1.0 eq) and 3,4-dimethoxybenzoic acid (2.0 eq), **EV-18** (44 mg, 26%) was obtained after prep-HPLC.

**$^1$H NMR (400 MHz, $CDCl_3$) δ** 7.48 (br s, 3H), 7.14 (br s, 2H), 6.86 (dd, *J* = 24.0, 9.5 Hz, 3H), 4.87 (br s, 1H), 3.89 (s, 3H), 3.85 (s, 3H), 3.71 (s, 2H), 2.03 – 1.90 (m, 6H), 1.33 (br s, 2H).

**HRMS ESI-TOF (*m/z*):** $[M+H]^+$ for $C_{22}H_{26}ClN_2O_4$ 417.1576, found 417.1577.

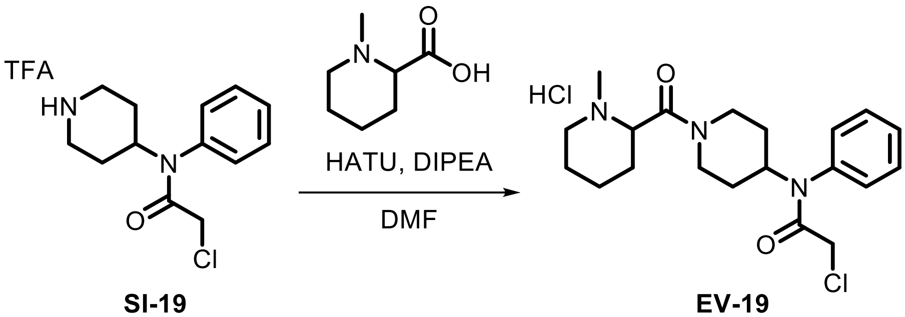

**SI-19** **EV-19**

Following **General Procedure 3**, starting with **SI-19** (150 mg, 0.41 mmol, 1.0 eq) and 1-methylpiperidine-2-carboxylic acid (2.0 eq), **EV-19** (20 mg, 13%) was obtained after prep-HPLC (HCl).

**$^1$H NMR (400 MHz, $CDCl_3$) δ** 12.03 (br s, 0.5H), 9.77 (d, *J* = 38.5 Hz, 0.5H), 7.46 (s, 3H), 7.13 (s, 2H), 4.89 – 4.49 (m, 2H), 4.41 – 3.95 (m, 1H), 3.71 (s, 3H), 3.35 – 3.06 (m, 2H), 2.87 – 2.61 (m, 4H), 2.56 – 2.30 (m, 2H), 2.22 – 1.90 (m, 3H), 1.92 – 1.59 (m, 3H), 1.52 – 1.35 (m, 1H), 1.36 – 1.14 (m, 2H).

**HRMS ESI-TOF (*m/z*):** $[M+H]^+$ for $C_{20}H_{29}ClN_3O_2$ 378.1943, found 378.1950.

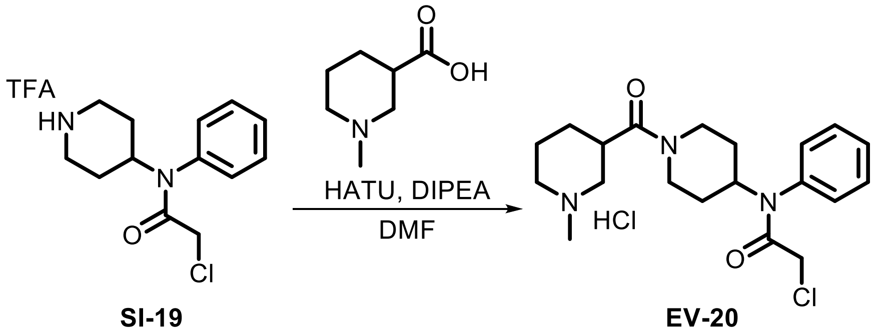

SI-19 EV-20

Following **General Procedure 3**, starting with **SI-19** (150 mg, 0.41 mmol, 1.0 eq) and 1-methylpiperidine-3-carboxylic acid (2.0 eq), **EV-20** (12 mg, 8%) was obtained after prep-HPLC (HCl).

**$^1$H NMR (400 MHz, CDCl$_3$) δ** 12.34 (br s, 0.8H), 9.44 (d, $J$ = 46.2 Hz, 0.2H), 7.46 (br s, 3H), 7.12 (br s, 2H), 4.87 – 4.72 (m, 1H), 4.61 (d, $J$ = 13.4 Hz, 1H), 4.04 (dd, $J$ = 45.2, 13.8 Hz, 1H), 3.70 (d, $J$ = 4.4 Hz, 2H), 3.68 – 3.51 (m, 1H), 3.49 – 3.42 (m, 1H), 3.35 – 3.10 (m, 2H), 3.02 – 2.81 (m, 1H), 2.74 (t, $J$ = 5.0 Hz, 3H), 2.70 – 2.57 (m, 1H), 2.41 – 2.22 (m, 1H), 2.06 – 1.85 (m, 5H), 1.61 – 1.39 (m, 1H), 1.37 – 1.08 (m, 2H).

**HRMS ESI-TOF (*m/z*):** $[M+H]^+$ for $C_{20}H_{29}ClN_3O_2$ 378.1943, found 378.1949.

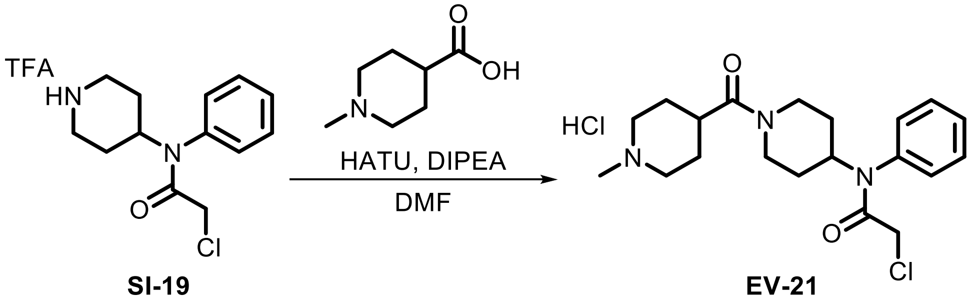

SI-19 EV-21

Following **General Procedure 3**, starting with **SI-19** (150 mg, 0.41 mmol, 1.0 eq) and 1-methylpiperidine-4-carboxylic acid (2.0 eq), **EV-21** (16 mg, 10%) was obtained after prep-HPLC (HCl).

**$^1$H NMR (400 MHz, CDCl$_3$) δ** 11.57 (s, 0.4H), 10.76 (s, 0.6H), 7.46 (br s, 3H), 7.13 (br s, 2H), 4.82 – 4.70 (m, 1H), 4.59 (br s, 2H), 4.09 – 3.79 (m, 1H), 3.70 (s, 2H), 3.48 (br s, 1H), 3.41 – 3.25 (m, 1H), 3.22 – 2.95 (m, 2H), 2.94 – 2.84 (m, 1H), 2.85 – 2.69 (m, 3H), 2.61 (t, $J$ = 12.8 Hz, 1H), 2.51 – 2.26 (m, 1H), 2.17 (s, 1H), 2.06 – 1.70 (m, 4H), 1.36 – 1.13 (m, 2H).

**HRMS ESI-TOF (*m/z*):** $[M+H]^+$ for $C_{20}H_{29}ClN_3O_2$ 378.1943, found 378.1950.

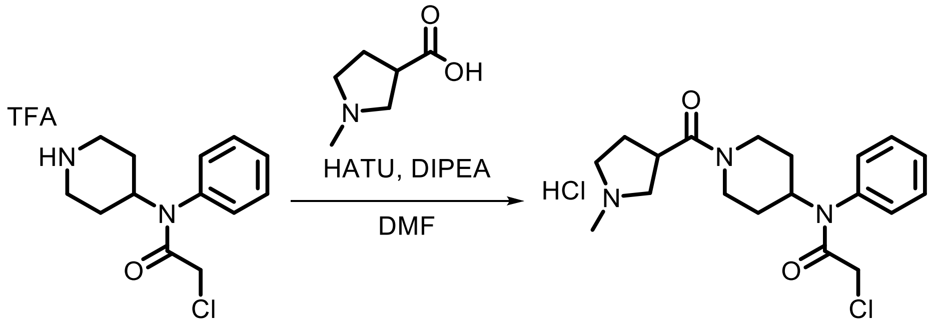

**SI-19** **EV-22**

Following **General Procedure 3**, starting with **SI-19** (150 mg, 0.41 mmol, 1.0 eq) and 1-methylpyrrolidine-3-carboxylic acid (2.0 eq), **EV-22** (17 mg, 11%) was obtained after prep-HPLC (HCl).

**$^1$H NMR (400 MHz, D$_2$O) δ** 7.54 – 7.48 (m, 3H), 7.28 (br s, 2H), 4.72 (t, *J* = 12.1 Hz, 1H), 4.49 – 4.34 (m, 1H), 3.98 (d, *J* = 14.0 Hz, 1H), 3.88 (s, 2H), 3.85 – 3.56 (m, 3H), 3.39 – 3.02 (m, 3H), 2.95 – 2.88 (m, 3H), 2.84 – 2.71 (m, 1H), 2.63 – 2.22 (m, 1H), 2.18 – 1.77 (m, 3H), 1.39 – 1.14 (m, 2H).

**HRMS ESI-TOF (*m/z*):** [M+H]$^+$ for $C_{19}H_{27}ClN_3O_2$ 364.1787, found 364.1794.

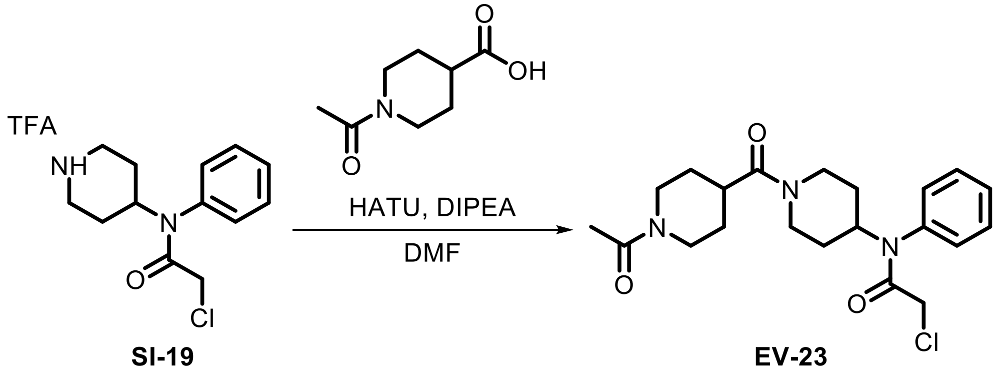

**SI-19** **EV-23**

Following **General Procedure 3**, starting with **SI-19** (150 mg, 0.41 mmol, 1.0 eq) and *N*-acetylpiperidine-4-carboxylic acid (2.0 eq), **EV-23** (37 mg, 22%) was obtained after prep-HPLC.

**$^1$H NMR (400 MHz, CDCl$_3$)** δ 7.48 – 7.44 (m, 3H), 7.14 – 7.10 (m, 2H), 4.82 (t, *J* = 12.2 Hz, 1H), 4.74 – 4.25 (m, 2H), 3.90 (br s, 1H), 3.71 (s, 2H), 3.29 – 2.74 (m, 3H), 2.63 (br s, 4H), 2.13 (s, 3H), 1.96 (br s, 1H), 1.88 (br s, 1H), 1.68 (br s, 3H), 1.23 (d, *J* = 13.1 Hz, 2H).

**HRMS ESI-TOF (*m/z*):** [M+H]$^+$ for $C_{21}H_{29}ClN_3O_3$ 406.1892, found 406.1893.

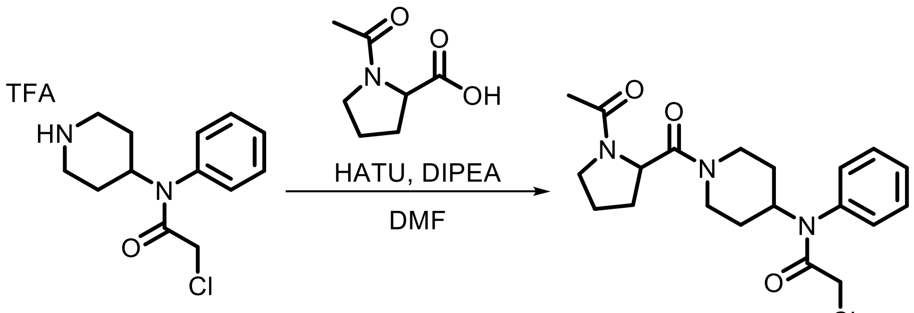

**SI-19** **EV-24**

Following **General Procedure 3**, starting with **SI-19** (150 mg, 0.41 mmol, 1.0 eq) and *N*-acetylpyrrolidine-2-carboxylic acid (2.0 eq), **EV-24** (28 mg, 17%) was obtained after prep-HPLC (HCl).

**$^1$H NMR (400 MHz, $CDCl_3$) δ** 7.46 (br s, 3H), 7.14 (br s, 2H), 6.22 (br s, 1H), 4.89 (br s, 1H), 4.75 (br s, 1H), 4.58 (d, *J* = 12.1 Hz, 1H), 3.92 (br s, 1H), 3.76 – 3.55 (m, 4H), 3.21 (d, *J* = 41.3 Hz, 1H), 2.81 – 2.58 (m, 1H), 2.28 – 2.14 (m, 3H), 2.15 – 1.69 (m, 5H), 1.60 – 1.16 (m, 2H).

**HRMS ESI-TOF (*m/z*):** $[M+H]^+$ for $C_{20}H_{27}ClN_3O_3$ 392.1736, found 392.1737.

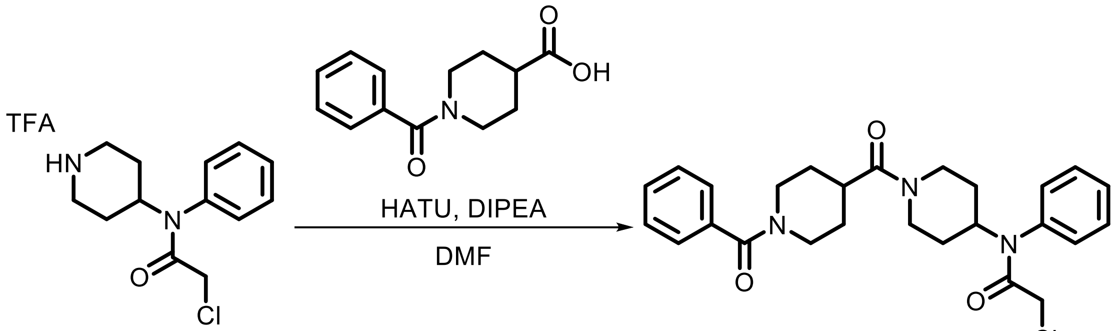

**SI-19** **EV-25**

Following **General Procedure 3**, starting with **SI-19** (150 mg, 0.41 mmol, 1.0 eq) and *N*-benzoylpiperidine-4-carboxylic acid (2.0 eq), **EV-25** (24 mg, 12%) was obtained after prep-HPLC.

**$^1$H NMR (400 MHz, $CDCl_3$) δ** 7.46 (br s, 3H), 7.38 (br s, 5H), 7.12 (br s, 2H), 4.82 (t, *J* = 12.0 Hz, 1H), 4.68 (br s, 1H), 3.92 (br s, 1H), 3.71 (s, 2H), 3.17 (br s, 1H), 2.94 (br s, 2H), 2.69 (br s, 2H), 2.32 – 2.10 (m, 3H), 1.92 (br s, 2H), 1.74 (br s, 3H), 1.25 (m, 2H).

**HRMS ESI-TOF (*m/z*):** $[M+H]^+$ for $C_{26}H_{31}ClN_3O_3$ 468.2049, found 468.2051.

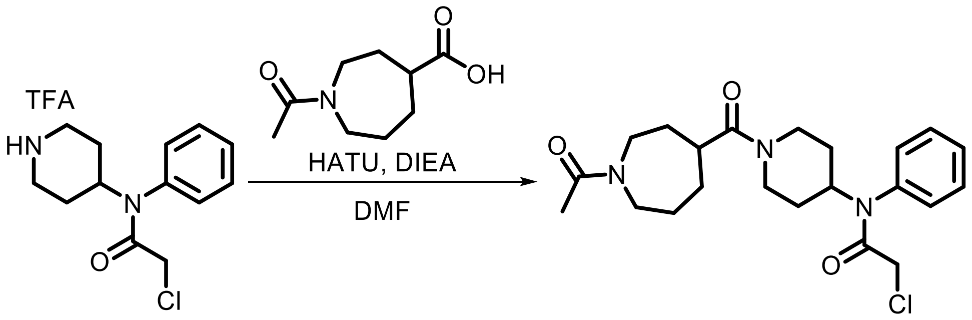

**SI-19** **EV-26**

Following **General Procedure 3**, starting with **SI-19** (100 mg, 0.27 mmol, 1.0 eq) and 1-acetylazepane-4-carboxylic acid (1.1 eq), **EV-26** was obtained after prep-HPLC (HCl) as a white solid (12 mg, 12%).

**$^1$H NMR (400 MHz, CDCl$_3$) δ** 7.50 – 7.42 (m, 3H), 7.19 – 7.07 (m, 2H), 4.78 (t, *J* = 10.8 Hz, 1H), 4.64 (br s, 1H), 3.93 (m, 1H), 3.70 (m, 3H), 3.58 (m, 1H), 3.36 (m, 1H), 3.14 (m, 1H), 2.67 (m, 2H), 2.41 (br s, 3H), 1.97 (m, 5H), 1.67 (m, 3H), 1.25 (m, 2H).

**HRMS ESI-TOF (*m/z*):** $[M+H]^+$ for $C_{22}H_{31}ClN_3O_3$ 420.2049, found 420.2049.

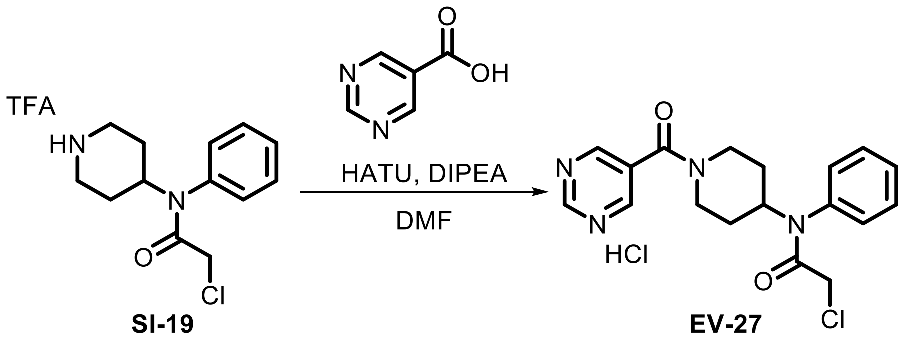

**SI-19** **EV-27**

Following **General Procedure 3**, starting with **SI-19** (150 mg, 0.41 mmol, 1.0 eq) and pyrimidine-5-carboxylic acid (2.0 eq), **EV-27** (13 mg, 9%) was obtained after prep-HPLC (HCl).

**$^1$H NMR (400 MHz, CDCl$_3$) δ** 9.23 (s, 1H), 8.70 (s, 2H), 7.49 (s, 3H), 7.13 (br s, 2H), 4.88 (tt, *J* = 12.2, 3.8 Hz, 1H), 4.76 (br s, 1H), 3.75 – 3.63 (m, 3H), 3.28 (br s, 1H), 2.89 (br s, 1H), 1.99 (br s, 1H), 1.91 (br s, 1H), 1.39 (br s, 1H), 1.31 (dd, *J* = 10.5, 6.8 Hz, 1H).

**HRMS ESI-TOF (*m/z*):** $[M+H]^+$ for $C_{19}H_{20}ClN_4O_2$ 359.1270, found 359.1270.

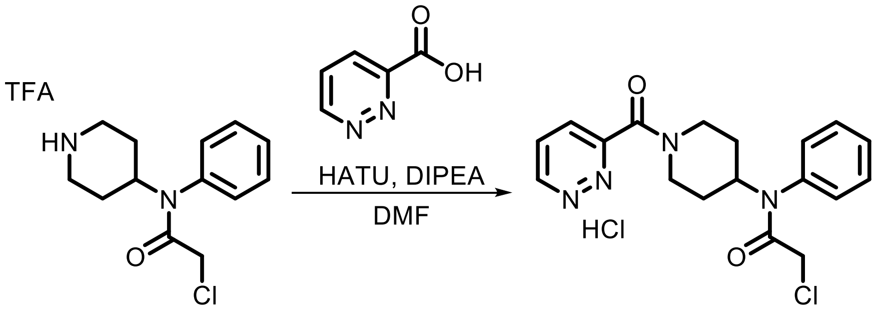

SI-19 EV-28

Following **General Procedure 3**, starting with **SI-19** (150 mg, 0.41 mmol, 1.0 eq) and pyridazine-3-carboxylic acid (2.0 eq), **EV-28** (14 mg, 9%) was obtained after prep-HPLC (HCl).

**$^1$H NMR (400 MHz, $CDCl_3$) δ** 9.69 (br s, 1H), 8.36 (br s, 2H), 7.46 (br s, 3H), 7.16 (br s, 2H), 4.94 – 4.66 (m, 2H), 4.08 (br s, 1H), 3.71 (s, 2H), 3.30 (br s, 1H), 2.97 (t, *J* = 11.4 Hz, 1H), 2.26 – 1.79 (m, 2H), 1.66 – 1.17 (m, 2H).

**HRMS ESI-TOF (*m/z*):** $[M+H]^+$ for $C_{19}H_{20}ClN_4O_2$ 359.1270, found 359.1273.

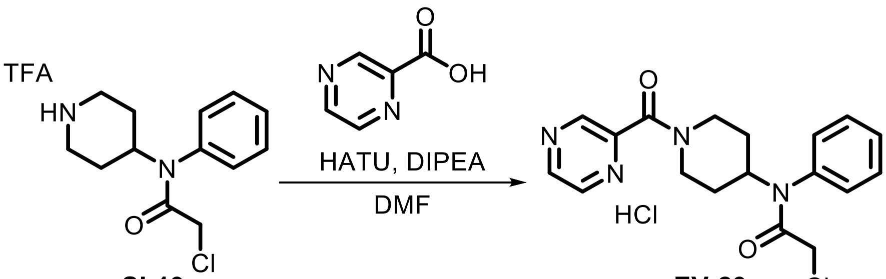

SI-19 EV-29

Following **General Procedure 3**, starting with **SI-19** (150 mg, 0.41 mmol, 1.0 eq) and pyrazine-2-carboxylic acid (2.0 eq), **EV-29** (24 mg, 16%) was obtained after prep-HPLC (HCl).

**$^1$H NMR (400 MHz, $CDCl_3$) δ** 8.85 (s, 1H), 8.58 (d, *J* = 21.5 Hz, 2H), 7.47 (s, 3H), 7.13 (dd, *J* = 17.3, 5.6 Hz, 2H), 4.95 – 4.85 (m, 1H), 4.81 (d, *J* = 12.9 Hz, 1H), 4.06 (d, *J* = 13.4 Hz, 1H), 3.71 (s, 2H), 3.23 (t, *J* = 12.9 Hz, 1H), 2.90 (t, *J* = 13.0 Hz, 1H), 1.99 (d, *J* = 12.6 Hz, 1H), 1.88 (d, *J* = 12.2 Hz, 1H), 1.50 – 1.34 (m, 2H).

**HRMS ESI-TOF (*m/z*):** $[M+H]^+$ for $C_{19}H_{20}ClN_4O_2$ 359.1270, found 359.1270.

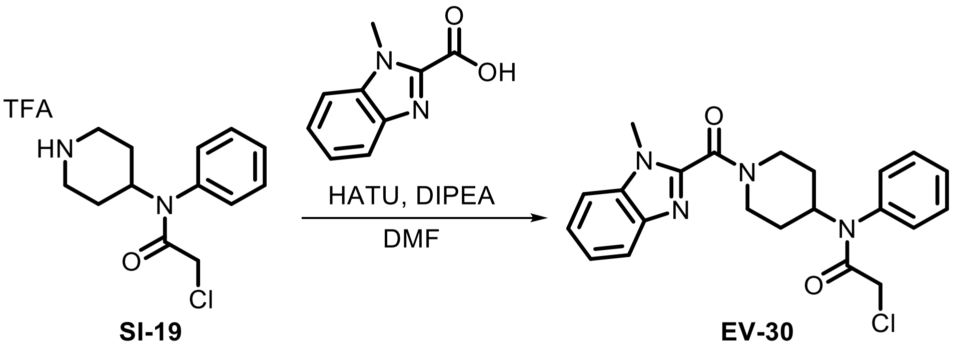

SI-19 EV-30

Following **General Procedure 3**, starting with **SI-19** (150 mg, 0.41 mmol, 1.0 eq) and 1-methyl-1*H*-benzimidazole-2-carboxylic acid (2.0 eq), **EV-30** (12 mg, 7%) was obtained after prep-HPLC.

**$^1$H NMR (400 MHz, CDCl$_3$) δ** 8.11 (br s, 1H), 7.63 (s, 3H), 7.46 (s, 3H), 7.26 (br s, 1H), 7.16 (br s, 1H), 4.72 (d, *J* = 12.1 Hz, 2H), 4.01 (s, 3H), 3.72 (s, 4H), 3.05 (t, *J* = 12.6 Hz, 1H), 1.98 (d, *J* = 12.7 Hz, 3H), 1.57 (d, *J* = 12.4 Hz, 1H).

**HRMS ESI-TOF (*m/z*):** $[M+H]^+$ for $C_{22}H_{24}ClN_4O_2$ 411.1583, found 411.1588.

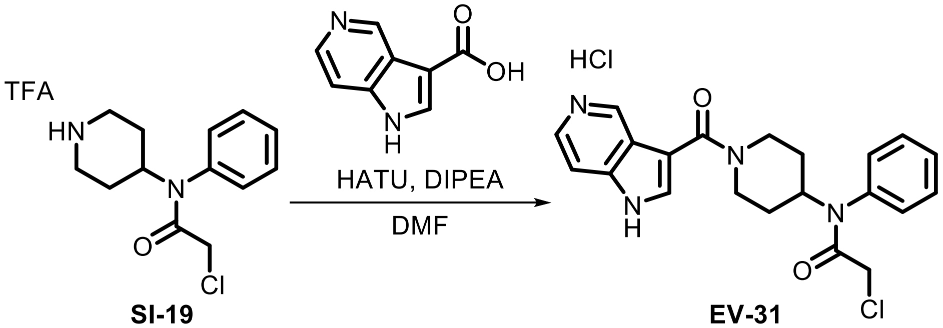

**SI-19** **EV-31**

Following **General Procedure 3**, starting with **SI-19** (150 mg, 0.41 mmol, 1.0 eq) and 1*H*-pyrrolo[3,2-*c*]pyridine-3-caroxylic acid (2.0 eq), **EV-31** (3 mg, 2%) was obtained after prep-HPLC (HCl).

**$^1$H NMR (400 MHz, CD$_3$CN) δ** 12.53 (br s, 1H), 10.91 (br s, 1H), 9.07 (d, *J* = 6.7 Hz, 1H), 8.26 (t, *J* = 6.3 Hz, 1H), 7.94 (d, *J* = 6.6 Hz, 1H), 7.89 (d, *J* = 2.5 Hz, 1H), 7.50 – 7.45 (m, 3H), 7.27 – 7.20 (m, 2H), 4.84 – 4.71 (m, 1H), 4.43 (s, 2H), 3.77 (s, 2H), 3.21 – 3.03 (m, 2H), 1.91 (br s, 2H), 1.33 (dd, *J* = 12.5, 4.3 Hz, 2H).

**HRMS ESI-TOF (*m/z*):** $[M+H]^+$ for $C_{21}H_{22}ClN_4O_2$ 397.1426, found 397.1428.

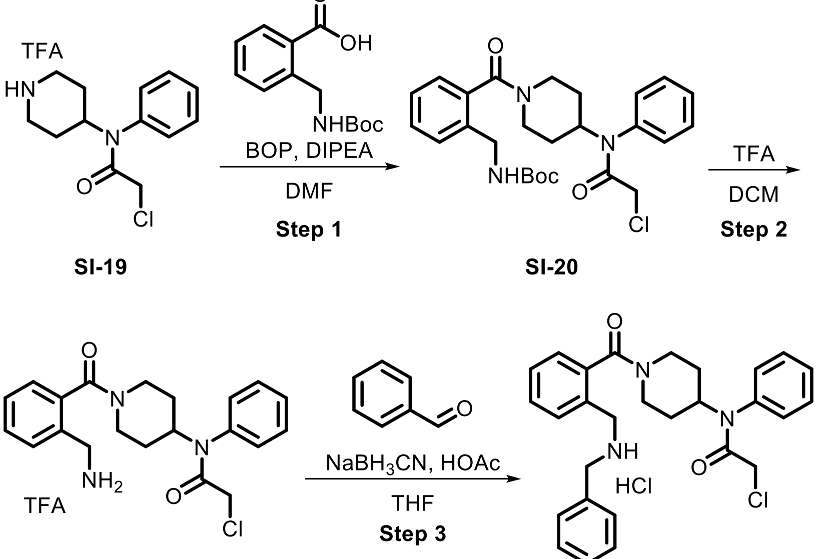

**SI-19** **SI-20** **SI-21** **EV-32**

**Step 1:** BOP (880 mg, 2.0 mmol, 1.0 eq) and DIPEA (1.0 mL, 6.0 mmol, 3.0 eq) were added to a solution of 2-(Boc-aminomethyl)benzoic acid (500 mg, 2.0 mmol, 1.0 eq) and **SI-19** (730 mg, 2.0 mmol, 1.0 eq) in DMF (2 mL). The mixture was stirred at 25 °C for 10 h. The mixture was diluted with DCM (30 mL) and washed with brine (30 mL x 3). The organic layer was dried over anhydrous $Na_2SO_4$, filtered, and concentrated. The resulting residue was purified by flash chromatography to afford **SI-20** (500 mg, 11%) as a white solid.

**Step 2:** TFA (0.32 mL, 4.1 mmol, 5.0 eq) was added to a solution of **SI-20** (400 mg, 0.82 mmol, 1.0 eq) in DCM (3 mL) and the mixture was stirred at 25 °C for 10 h. The reaction was diluted with $H_2O$ (20 mL) and extracted with DCM (30 mL x 3). The combined organic phases were washed with brine (20 mL), dried over anhydrous $Na_2SO_4$, filtered, and concentrated. The residue was purified by prep-HPLC (HCl) to provide **SI-21** (130 mg, 35%) as a white solid.

**Step 3:** HOAc (7.4 µL, 0.13 mmol, 1.0 eq) was added to a solution of **SI-21** (50 mg, 0.13 mmol, 1.0 eq) and benzaldehyde (13 µL, 0.13 mmol, 1.0 eq) in THF (1 mL) and the mixture was stirred at 25 °C for 1 h. $NaBH_3CN$ (8 mg, 0.13 mmol, 1.0 eq) was then added and the mixture was stirred at 0 °C for 2 h. The reaction mixture was quenched by the addition of 1 M HCl (10 mL) and then diluted with $H_2O$ (10 mL). The mixture was extracted with DCM (20 mL x 2) and the combined organic layers were washed with brine (10 mL), dried over anhydrous $Na_2SO_4$, filtered and concentrated. The resulting residue was purified by prep-HPLC (HCl) to afford **EV-32** (11 mg, 18%) as a white solid.

**$^1$H NMR (400 MHz, $CDCl_3$) δ** 10.00 (br s, 1H), 7.73 (br s, 1H), 7.61 – 7.38 (m, 7H), 7.33 (br s, 3H), 7.23 – 7.05 (m, 3H), 4.89 – 4.66 (m, 2H), 4.02 (br s, 3H), 3.79 – 3.66 (m, 3H), 3.19 (br s, 1H), 2.84 (t, *J* = 13.8 Hz, 1H), 2.07 – 1.95 (m, 1H), 1.86 (br s, 1H), 1.69 (br s, 1H), 1.48 – 1.16 (m, 2H).

**HRMS ESI-TOF (*m/z*):** $[M+H]^+$ for $C_{28}H_{31}ClN_3O_2$ 476.2100, 476.2104.

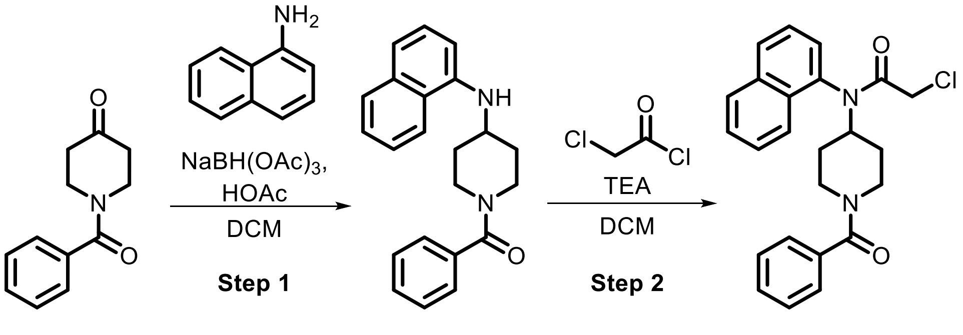

**SI-22** **EV-33**

**Step 1:** Following **General Procedure 5**, starting with 1-benzoylpiperidin-4-one (300 mg, 1.5 mmol, 1.0 eq) and naphthalen-1-amine (1.2 eq) in DCM (2 mL), **SI-22** was obtained as a yellow oil and used without purification.

**Step 2:** Following **General Procedure 1**, starting with **SI-22** (425 mg, 1.2 mmol, 1.0 eq), chloroacetyl chloride (3.0 eq), and TEA (6.0 eq). **EV-33** was obtained after prep-HPLC (FA) as a yellow oil (68 mg, 13%).

**$^1$H NMR (300 MHz, $CDCl_3$) δ** 8.01 – 7.90 (m, 2H), 7.82 (br s, 1H), 7.63 – 7.48 (m, 3H), 7.43 – 7.12 (br s, 6H), 4.95 (tt, *J* = 12.2, 3.9 Hz, 1H), 4.73 (br , 1H), 3.82 (br s, 1H), 3.70 – 3.55 (m, 2H), 3.09 (br s, 1H), 2.88 (br s, 1H), 2.32 – 2.04 (m, 1H), 1.78 – 1.43 (m, 2H), 1.04 (br, 1H).

**HRMS ESI-TOF (*m/z*):** $[M+H]^+$ for $C_{24}H_{24}ClN_2O_2$ 407.1521, found 407.1523.

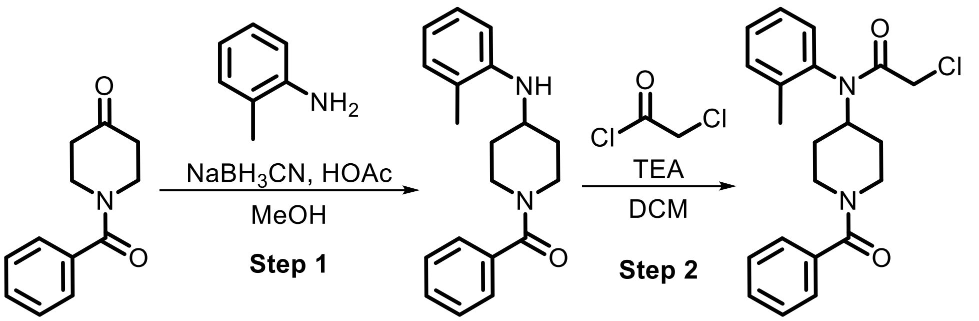

**SI-23** **EV-34**

**Step 1:** Following **General Procedure 5**, starting with 1-benzoylpiperidin-4-one (300 mg, 1.5 mmol, 1.0 eq) and 2-methylaniline (1.2 eq) in MeOH, **SI-23** was obtained as a yellow oil and used without purification.

**Step 2:** Following **General Procedure 1**, starting with **SI-23** (1.0 eq), chloroacetyl chloride (3.0 eq), and TEA (6.0 eq), **EV-34** (43 mg, 10%) was obtained after prep-HPLC.

**$^1$H NMR (400 MHz, $CDCl_3$) δ** 7.37 – 7.32 (m, 5H), 7.32 – 7.23 (m, 3H), 7.05 (s, 1H), 4.71 (tt, *J* = 12.1, 3.9 Hz, 2H), 3.80 (br s, 1H), 3.64 (s, 2H), 3.10 (br s, 1H), 2.82 (br s, 1H), 2.24 (br s, 3H), 2.15 – 1.98 (m, 1H), 1.83 (br s, 1H), 1.64 – 1.42 (m, 1H), 1.35 – 1.11 (m, 1H).

**HRMS ESI-TOF (*m/z*):** $[M+H]^+$ for $C_{21}H_{24}ClN_2O_2$ 371.1521, found 371.1520.

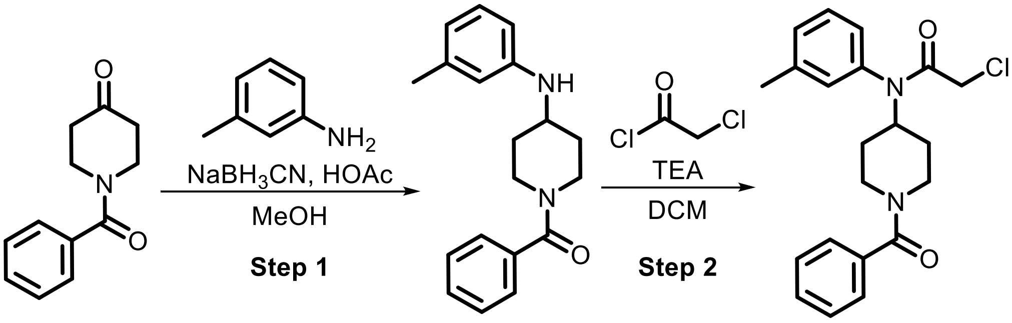

**SI-24** **EV-35**

**Step 1:** Following **General Procedure 5**, starting with 1-benzoylpiperidin-4-one (300 mg, 1.5 mmol, 1.0 eq) and 3-methylaniline (1.2 eq) in MeOH, **SI-24** was obtained as a yellow oil and used without purification.

**Step 2:** Following **General Procedure 1**, starting with **SI-24** (1.0 eq), chloroacetyl chloride (3.0 eq), and TEA (6.0 eq), **EV-35** (71 mg, 16%) was obtained after prep-HPLC.

**$^1$H NMR (400 MHz, $CDCl_3$) δ** 7.32 – 7.15 (m, 7H), 6.83 (s, 2H), 4.79 – 4.66 (m, 2H), 3.70 (br s, 1H), 3.62 (s, 2H), 3.04 (br s, 1H), 2.79 (br s, 1H), 2.31 (s, 3H), 1.87 (br s, 1H), 1.73 (br s, 1H), 1.31 (br s, 1H), 1.18 – 1.09 (m, 1H).

**HRMS ESI-TOF (*m/z*):** $[M+H]^+$ for $C_{21}H_{24}ClN_2O_2$ 371.1521, found 371.1522.

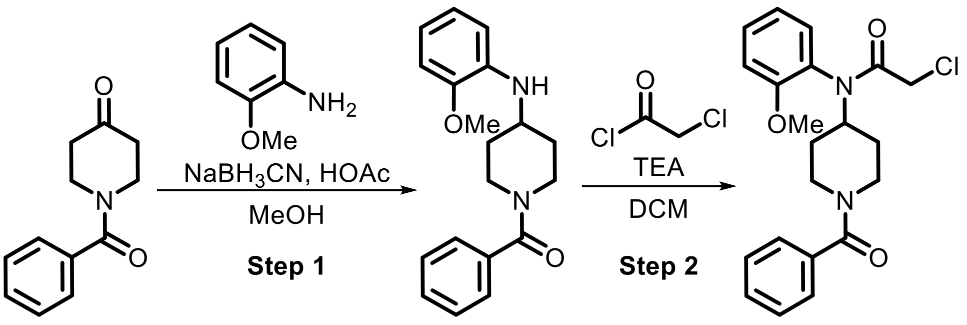

**SI-25** **EV-36**

**Step 1:** Following **General Procedure 5**, starting with 1-benzoylpiperidin-4-one (300 mg, 1.5 mmol, 1.0 eq) and 2-methoxyaniline (1.2 eq) in MeOH, **SI-25** was obtained as a yellow oil and used without purification.

**Step 2:** Following **General Procedure 1**, starting with **SI-25** (1.0 eq), chloroacetyl chloride (3.0 eq), and TEA (6.0 eq), **EV-36** (400 mg, 86%) was obtained after prep-HPLC.

**$^1$H NMR (400 MHz, CDCl$_3$) δ** 7.47 – 7.34 (m, 4H), 7.33 – 7.28 (m, 2H), 7.07 (m, 3H), 4.86 – 4.74 (m, 2H), 3.82 (d, *J* = 17.1 Hz, 4H), 3.73 (d, *J* = 4.2 Hz, 2H), 3.15 (d, *J* = 13.7 Hz, 1H), 2.85 (d, *J* = 15.7 Hz, 1H), 2.11 – 1.79 (m, 2H), 1.60 – 1.00 (m, 2H).

**HRMS ESI-TOF (*m/z*):** $[M+H]^+$ for $C_{21}H_{24}ClN_2O_3$ 387.1470, found 387.1472.

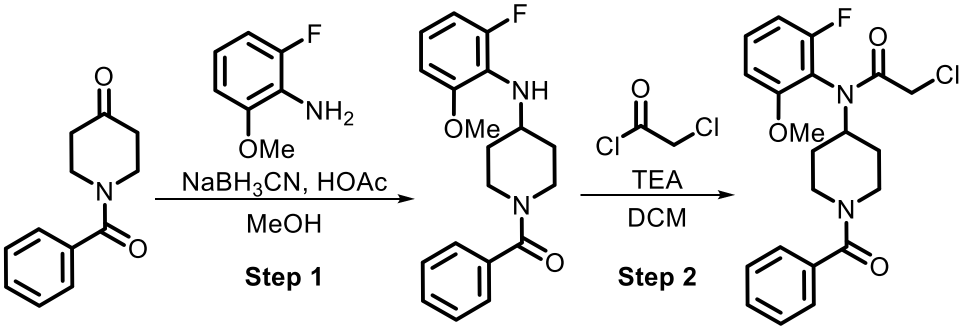

**SI-26** **EV-37**

**Step 1:** Following **General Procedure 5**, starting with 1-benzoylpiperidin-4-one (300 mg, 1.5 mmol, 1.0 eq) and 2-fluoro-6-methoxyaniline (1.2 eq) in MeOH, **SI-26** was obtained as a yellow oil and used without purification.

**Step 2:** Following **General Procedure 1**, starting with **SI-26** (1.0 eq), chloroacetyl chloride (3.0 eq), and TEA (6.0 eq), **EV-37** (71 mg, 15%) was obtained after prep-HPLC.

**$^1$H NMR (400 MHz, CDCl$_3$) δ** 7.44 – 7.34 (m, 4H), 7.35 – 7.30 (m, 2H), 6.89 – 6.77 (m, 2H), 4.75 (br s, 2H), 3.90 – 3.78 (m, 4H), 3.73 (s, 2H), 3.14 (br s, 1H), 2.84 (br s, 1H), 1.98 (d, *J* = 12.7 Hz, 2H), 1.54 – 1.20 (m, 2H).

**HRMS ESI-TOF (*m/z*):** $[M+H]^+$ for $C_{21}H_{23}ClFN_2O_3$ 405.1376, found 405.1381.

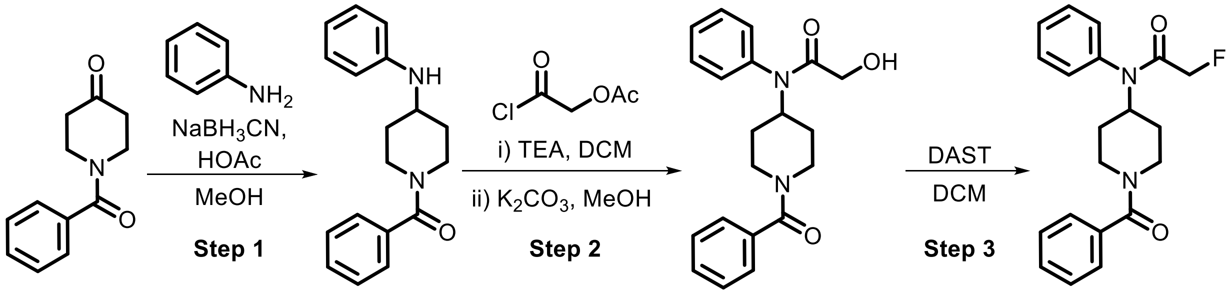

**Step 1:** Following **General Procedure 5**, starting with 1-benzoylpiperidin-4-one (300 mg, 1.5 mmol, 1.0 eq) and aniline (1.2 eq) in MeOH, **SI-27** was obtained as a yellow oil and used without purification.

**Step 2:** Acetoxyacetyl chloride (115 µL, 1.1 mmol, 1.5 eq) was added to a solution of **SI-27** (200 mg, 0.71 mmol, 1.0 eq) and TEA (297 µL, 2.1 mmol, 3.0 eq) in DCM (3 mL). The reaction was stirred for 1 h before being diluted with DCM (15 mL). The mixture was washed with water (10 mL), dried over anhydrous $Na_2SO_4$, and concentrated. The resulting residue was dissolved in MeOH (5 mL), $K_2CO_3$ (196 mg, 1.4 mmol, 2.0 eq) was added, and the reaction was stirred for 1 h. The reaction was concentrated under reduced pressure and the residue was dissolved in water (10 mL) and DCM (15 mL). The aqueous phase was extracted with DCM (2 x 15 mL). The combined organic phases were dried over anhydrous $Na_2SO_4$ and concentrated to provide **SI-28** which was used without purification.

**Step 3:** DAST (47 µL, 0.35 mmol, 1.2 eq) was added to a solution of **SI-28** (100 mg, 0.30 mmol, 1.0 eq) in DCM (2 mL) at 0 °C. The reaction was warmed to 25 °C and stirred for 2 h. Upon completion, the reaction mixture was diluted with DCM (15 mL) and washed with $H_2O$ (10 mL), dried over anhydrous $Na_2SO_4$, and concentrated. The resulting residue was purified by prep-TLC (PE:EtOAc = 1:2) to provide **EV-38** (30 mg, 29%) as a white solid.

**$^1$H NMR (400 MHz, MeOD) δ** 7.55 – 7.50 (m, 3H), 7.46 – 7.39 (m, 3H), 7.32 – 7.25 (m, 4H), 4.80 (tt, *J* = 12.1, 3.9 Hz, 1H), 4.69 (br s, 1H), 4.60 (s, 1H), 4.48 (s, 1H), 3.74 (br s, 1H), 3.24 (br s, 1H), 2.93 (br s, 1H), 2.00 (br s, 1H), 1.86 (br s, 1H), 1.50 – 1.07 (m, 2H).

**HRMS ESI-TOF (*m/z*):** $[M+H]^+$ for $C_{20}H_{22}FN_2O_2$ 341.1660, found 341.1663.

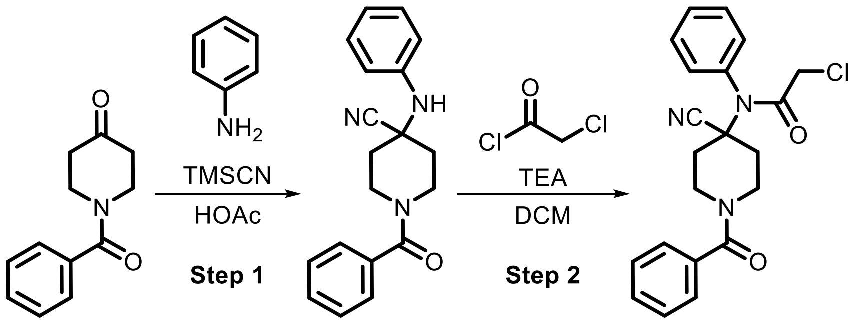

**Step 1:** TMSCN (107 mg, 1.1 mmol, 1.1 eq) was added dropwise to a solution of 1-benzoylpiperidin-4-one (200 mg, 0.98 mmol, 1.0 eq) and aniline (90 µL, 0.98 mmol, 1.0 eq) at 0 °C. The reaction was warmed to 25 °C and stirred for 3 h. The reaction was cooled to 0 °C and aqueous $NH_4OH$ was added to the reaction until pH 10 and then extracted with DCM (3 x 10 mL). The combined organic phases were dried over anhydrous $Na_2SO_4$ and

concentrated. The resulting residue was purified by flash chromatography (PE:EtOAc = 50:1-2:1) to provide **SI-29** (260 mg, 74%) as a yellow oil.

**Step 2:** Following **General Procedure 1**, starting with **SI-29** (260 mg, 0.85 mmol, 1.0 eq), chloroacetyl chloride (1.5 eq), and TEA (3.0 eq), **EV-39** was obtained after prep-TLC (PE:EtOAc = 1:1) as a light yellow solid (70 mg, 23%).

**$^1$H NMR (400 MHz, MeOD) δ** 7.59 – 7.54 (m, 3H), 7.48 – 7.37 (m, 7H), 4.71 (br s, 1H), 3.85 (s, 3H), 3.46 (br s, 1H), 3.15 (br s, 1H), 2.41 (dd, *J* = 13.4, 2.6 Hz, 2H), 1.68 (t, *J* = 12.2 Hz, 2H).

**HRMS ESI-TOF (*m/z*):** $[M+H]^+$ for $C_{21}H_{21}ClN_3O_2$ 382.1317, found 382.1316.

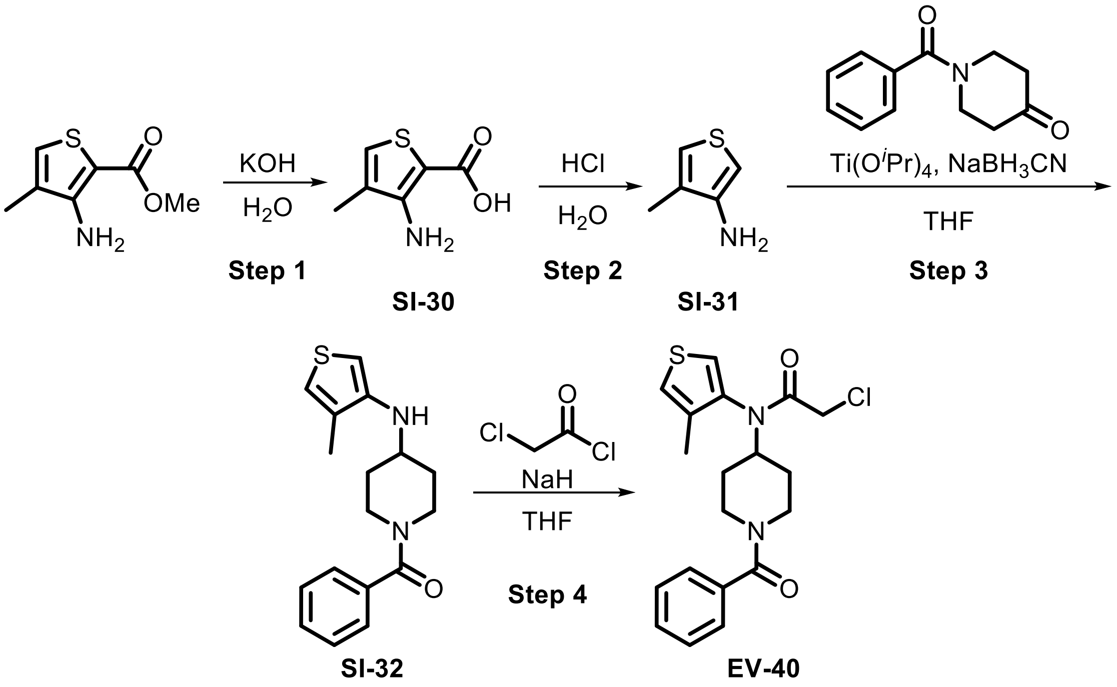

**Step 1:** Methyl 3-amino-4-methylthiophene-2-carboxylate (1.0 g, 5.8 mmol, 1.0 eq) was added to a solution of KOH (459 mg, 8.2 mmol, 1.4 eq) in $H_2O$ (3 mL) at 28 °C. The mixture was heated to 80 °C and stirred for 30 min. The mixture was not worked up and used directly in the next step.

**Step 2:** The solution of **SI-30** (1.0 g, 5.1 mmol, 1.0 eq) in $H_2O$ (3 mL) and HCl (6 M, 2.8 mL, 3.3 eq) was stirred at 50 °C for 16 h. The mixture was slowly poured into aqueous $NaHCO_3$ (20 mL) and extracted with EtOAc (10 mL x 2). The combined organic phases were dried over anhydrous $Na_2SO_4$ and concentrated. **SI-31** (500 mg, crude) was obtained as brown liquid and used without purification.

**Step 3:** $Ti(O^iPr)_4$ (786 µL, 2.7 mmol, 1.5 eq) was added to a solution of 1-benzoylpiperidin-4-one (468 mg, 2.3 mmol, 1.3 eq) and **SI-31** (200 mg, 1.8 mmol, 1.0 eq) in THF (5 mL) and the mixture was stirred at 70 °C for 16 h. The reaction was cooled to 0 °C and $NaBH_3CN$ (167 mg, 2.7 mmol, 1.5 eq) was added. The mixture was stirred for an additional 1 h and then poured into aqueous $NaHCO_3$ (10 mL) and extracted with EtOAc (15 mL x 3). The combined organic phases were dried over anhydrous $Na_2SO_4$ and concentrated. The resulting residue was purified by flash chromatography (PE:EtOAc = 15:1-3:1) to provide **SI-32** (200 mg, 25%) as yellow oil.

**Step 4:** NaH (68 mg, 1.7 mmol, 60% dispersion in mineral oil, 3.0 eq) was added to a solution of **SI-32** (170 mg, 0.57 mmol, 1.0 eq) in THF (3 mL) at 0 °C. The mixture was warmed to 28 °C and stirred for 0.5 h. The reaction was cooled to 0 °C and chloroacetyl chloride (90 µL, 1.1 mmol, 2.0 eq) was added. The mixture was warmed to 28 °C and stirred for an additional 1.5 h. The reaction was cooled to 0 °C and water (5 mL) was added. The mixture was extracted with EtOAc (5 mL x 3) and the combined organic phases were dried over anhydrous $Na_2SO_4$ and concentrated. The resulting residue was purified by prep-TLC (PE:EtOAc = 1:1) and then by prep-HPLC (HCl) to provide **EV-40** (24 mg, 11%) as a light yellow solid.

**$^1$H NMR (400 MHz, $CDCl_3$) δ** 7.46 – 7.28 (m, 5H), 7.11 (br s, 2H), 4.76 (tt, *J* = 12.1, 4.0 Hz, 2H), 3.78 (br s, 1H), 3.76 – 3.62 (m, 2H), 3.13 (br s, 1H), 2.84 (br s, 1H), 2.10 (br s, 3H), 1.97 – 1.68 (m, 2H), 1.60 – 1.05 (m, 2H).

**HRMS ESI-TOF (*m/z*):** $[M+H]^+$ for $C_{19}H_{22}ClN_2O_2S$ 377.1085, found 377.1086.

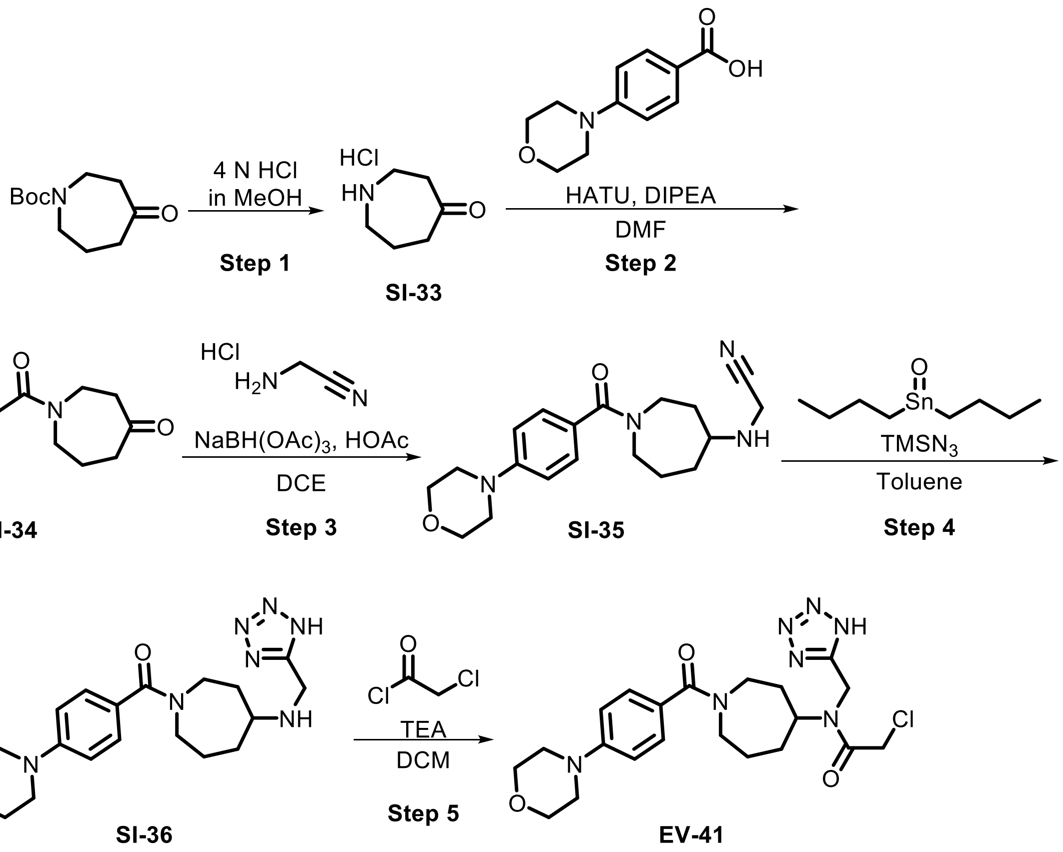

**Step 1:** A mixture of *tert*-butyl 4-oxoazepane-1-carboxylate (1.0 g, 4.7 mmol, 1.0 eq) in 4N HCl/MeOH (5.0 mL, 4.3 eq) was stirred at 15 °C for 2 h. The reaction mixture was concentrated under reduced pressure to afford **SI-33** (700 mg, crude) as a white solid which was used in the next step without purification.

**Step 2:** Following **General Procedure 3**, starting with **SI-33** (51 mg, 0.34 mmol, 1.0 eq) and 4-morpholinobenzoic acid (1.0 eq), **SI-34** was obtained as a yellow oil and used in the next step without purification.

**Step 3:** HOAc (642 µL, 11.2 mmol, 2.0 eq) and 2-aminoacetonitrile hydrochloride (2.9 g, 22.5 mmol, 4.0 eq) were added to a solution of **SI-34** (1.70 g, 5.6 mmol, 1.0 eq) in DCE (4 mL) and the mixture was stirred at 30 °C for 30 min. The reaction was cooled to 0 °C and $NaBH(OAc)_3$ (1.79 g, 8.4 mmol, 1.5 eq) was added portion-wise. The reaction was heated to 20 °C and stirred for 16 h. The mixture was quenched with $H_2O$ (20 mL) and extracted with DCM (30 mL x 3). The combined organic layers were dried over anhydrous $Na_2SO_4$ and concentrated under reduced pressure. The resulting residue was purified by prep HPLC (TFA) to provide **SI-35** (560 mg, 1.6 mmol, 29%) as a colorless oil.

**Step 4:** Dibutyltin oxide (22 mg, 0.09 mmol, 0.2 eq) and azido(trimethyl)silane (233 µL, 1.8 mmol, 4.0 eq) were added to a solution of **SI-35** (150 mg, 0.44 mmol, 1.0 eq) in toluene (3 mL). The reaction mixture was stirred at 110 °C for 16 h. Toluene was removed and 1N NaOH (10 mL) was added to the resulting residue. The mixture was washed with DCM (10 mL). The aqueous phase was concentrated and the remaining residue was purified by prep-HPLC (basic) to provide **SI-36** (50 mg, 30%) as a light yellow solid.

**Step 5:** Following **General Procedure 1**, starting with **SI-36** (20 mg, 0.06 mmol, 1.0 eq), chloroacetyl chloride (1.0 eq), and TEA (3.0 eq), **EV-41** was obtained after prep-HPLC (HCl) as a colorless oil (7 mg, 56%).

**$^1$H NMR (400 MHz, DMSO-*d*$_6$) δ** 7.34 – 7.20 (m, 2H), 7.00 (d, *J* = 8.2 Hz, 2H), 4.93 – 4.62 (m, 2H), 4.59 – 4.33 (m, 2H), 4.13 (s, 1H), 3.73 (t, *J* = 4.8 Hz, 5H), 3.58 – 3.42 (m, 1H), 3.38 – 3.21 (m, 2H), 3.17 (t, *J* = 4.8 Hz, 4H), 2.09 – 1.81 (m, 3H), 1.79 – 1.45 (m, 3H).

**HRMS ESI-TOF (*m/z*):** $[M+H]^+$ for $C_{21}H_{29}ClN_7O_3$ 462.2015, found 462.2017.

**Synthesis of SI-37 as a precursor for EV-42 and EV-43**

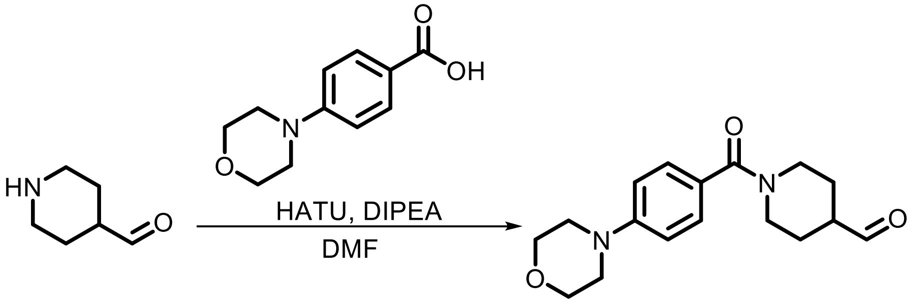

**SI-37**

Following **General Procedure 3**, starting with piperidine-4-carboxaldehyde (2.0 g, 13.4 mmol, 1.0 eq) and 4-morpholinobenzoic acid (1.1 eq), **SI-37** was obtained after prep-HPLC (TFA) as a yellow oil (1.15 g, 20%).

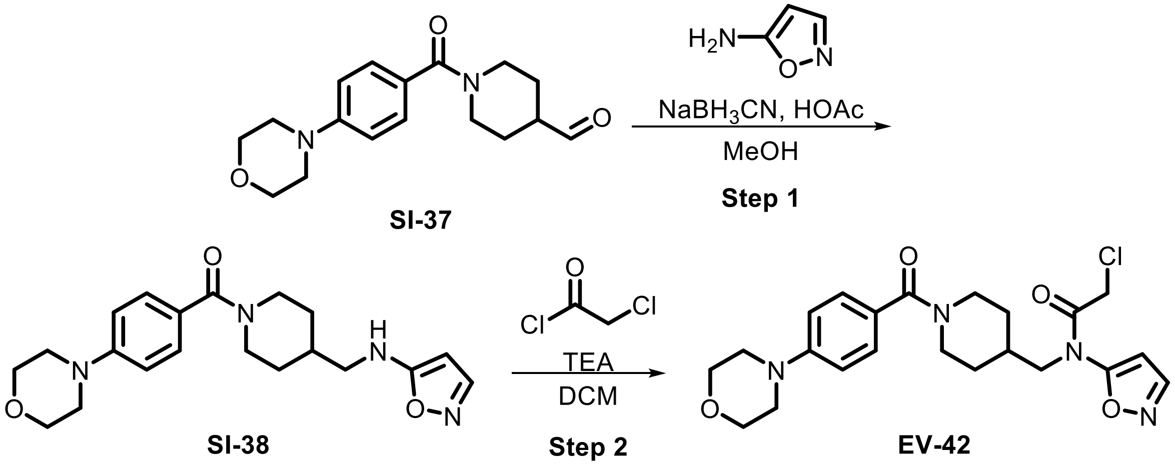

**Step 1:** Following **General Procedure 5**, starting with **SI-37** (40 mg, 0.13 mmol, 1.0 eq) and isoxazol-5-amine (1.1 eq), **SI-38** (30 mg, 45%) was obtained after purification by prep-TLC (EtOAc:MeOH = 10:1).

**Step 2:** Following **General Procedure 1**, starting with **SI-38** (30 mg, 0.08 mmol, 1.0 eq), chloroacetyl chloride (2.0 eq), and TEA (4.0 eq), **EV-42** was obtained after prep-HPLC (HCl) as a colorless oil (13 mg, 29%).

**$^1$H NMR (400 MHz, DMSO-*d*$_6$) δ** 8.64 (d, *J* = 1.9 Hz, 1H), 7.28 – 7.23 (m, 2H), 7.04 – 6.94 (m, 2H), 6.57 (d, *J* = 1.9 Hz, 1H), 4.47 (br s, 2H), 3.76 – 3.71 (m, 5H), 3.68 (d, *J* = 7.3 Hz, 2H), 3.21 – 3.13 (m, 5H), 2.82 (br s, 2H), 1.90 – 1.80 (m, 1H), 1.61 (d, *J* = 12.5 Hz, 2H), 1.26 – 0.96 (m, 3H).

**HRMS ESI-TOF (*m/z*):** $[M+H]^+$ for $C_{22}H_{28}ClN_4O_4$ 447.1794, found 447.1796.

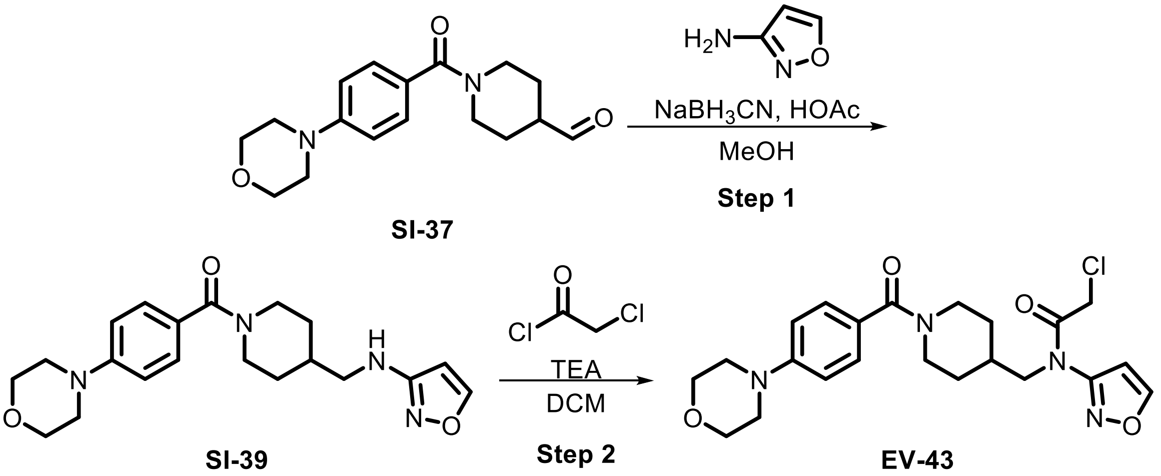

**Step 1:** Following **General Procedure 5**, starting with **SI-37** (30 mg, 0.1 mmol, 1.0 eq) and isoxazol-5-amine (1.1 eq), **SI-39** (13 mg, 29%) was obtained after purification by flash chromatography (DCM:MeOH = 100:1 – 10:1).

**Step 2:** Following **General Procedure 1**, starting with **SI-39** (30 mg, 0.08 mmol, 1.0 eq), chloroacetyl chloride (2.0 eq), and TEA (4.0 eq), **EV-43** was obtained after prep-HPLC (HCl) as a colorless oil (13 mg, 29%).

**$^1$H NMR (400 MHz, DMSO-*d*$_6$) δ** 8.98 (s, 1H), 7.29 (d, *J* = 8.5 Hz, 2H), 7.07 (d, *J* = 8.4 Hz, 2H), 6.93 (d, *J* = 1.7 Hz, 1H), 4.52 (s, 2H), 3.77 (t, *J* = 4.8 Hz, 6H), 3.71 (d, *J* = 7.3 Hz, 2H), 3.21 (t, *J* = 4.8 Hz, 4H), 2.80 (br s, 2H), 1.87 (br s, 1H), 1.59 (d, *J* = 12.6 Hz, 2H), 1.28-1.02 (m, 2H).

**HRMS ESI-TOF (*m/z*):** $[M+H]^+$ for $C_{22}H_{28}ClN_4O_4$ 447.1794, found 447.1796.

#### Synthesis of SI-42 as a precursor for EV-44 and EV-45

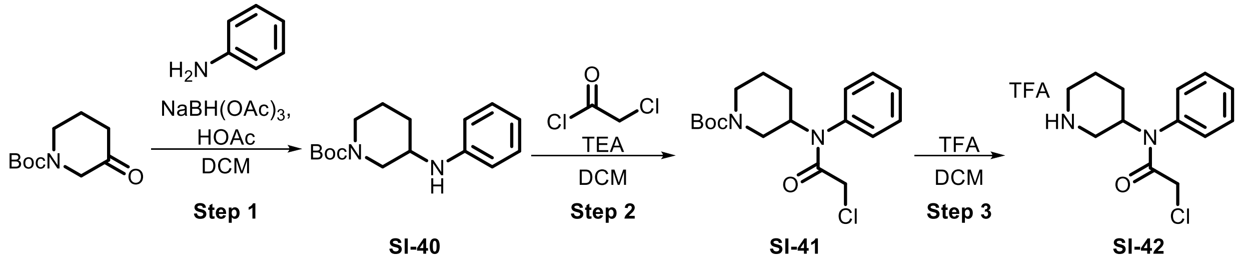

**Step 1:** A solution of HOAc (2.9 mL, 50.2 mmol, 1.0 eq) , aniline (4.6 mL, 50.2 mmol, 1.0 eq) and 1-Boc-3-piperidone (10.0 g, 50.2 mmol, 1.0 eq) in anhydrous DCM (150 mL) was stirred at 25 °C for 16 h. $NaBH(OAc)_3$ (21.3 g, 100 mmol, 2.0 eq) was then added and the reaction was stirred for an additional 3 h. The reaction mixture was quenched with aq. $NaHCO_3$ (50 mL), washed with saturated brine (50 mL), dried over anhydrous $Na_2SO_4$, filtered and concentrated under reduced pressure. **SI-40** was obtained as yellow oil which was used without purification.

**Step 2:** Following **General Procedure 1**, starting with **SI-40** (1.0 eq), chloroacetyl chloride (1.5 eq), and TEA (3.0 eq), **SI-41** was obtained and used in the next step without additional purification.

**Step 3:** TFA (1.5 mL, 20.4 mmol, 3.0 eq) was added dropwise to a solution of **SI-41** (2.4 g, 6.80 mmol, 1.0 eq) in DCM (2 mL) at 0 °C. The mixture was then warmed to 25 °C and stirred for 2 h. The solution was quenched with $H_2O$ (2 mL) and extracted with DCM (2 mL x 3), the combined organic layers were washed with brine (2 mL x 3), dried over anhydrous $Na_2SO_4$ and concentrated under reduced pressure to provide **SI-42** (1.3 g, crude) as a yellow oil which was used without purification.

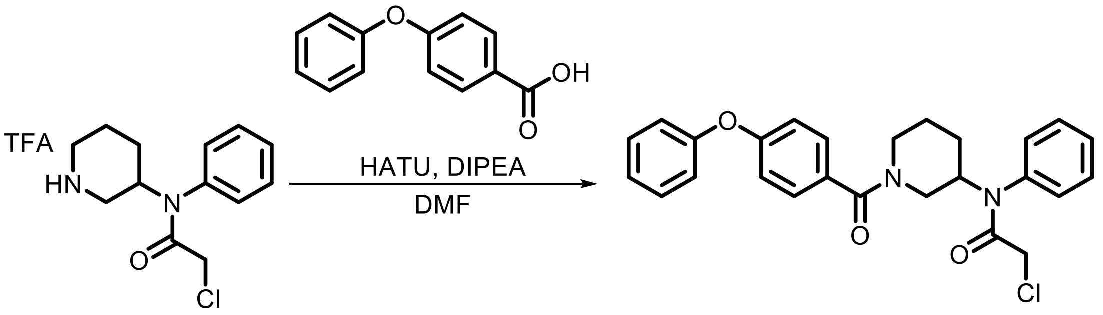

**SI-42** **EV-44**

Following **General Procedure 3**, starting with **SI-42** (248 mg, 0.98 mmol, 1.2 eq) and 4-phenoxybenzoic acid (1.0 eq), **EV-44** (29 mg, 29%) was obtained after prep-HPLC (HCl).

**$^1$H NMR (400 MHz, DMSO-$d_6$) δ** 7.48 (s, 3H), 7.43 (t, $J$ = 7.8 Hz, 4H), 7.40 – 7.24 (m, 2H), 7.20 (t, $J$ = 7.4 Hz, 1H), 7.09 (d, $J$ = 8.0 Hz, 2H), 7.01 (d, $J$ = 8.3 Hz, 2H), 4.43 (s, 2H), 3.83 (s, 2H), 3.55 (br s, 1H), 2.89 – 2.53 (m, 2H), 1.88 (br s, 1H), 1.74 – 1.42 (m, 2H), 1.31 – 1.00 (m, 1H).

**HRMS ESI-TOF ($m/z$):** $[M+H]^+$ for $C_{26}H_{26}ClN_2O_3$ 449.1627, found 449.1626.

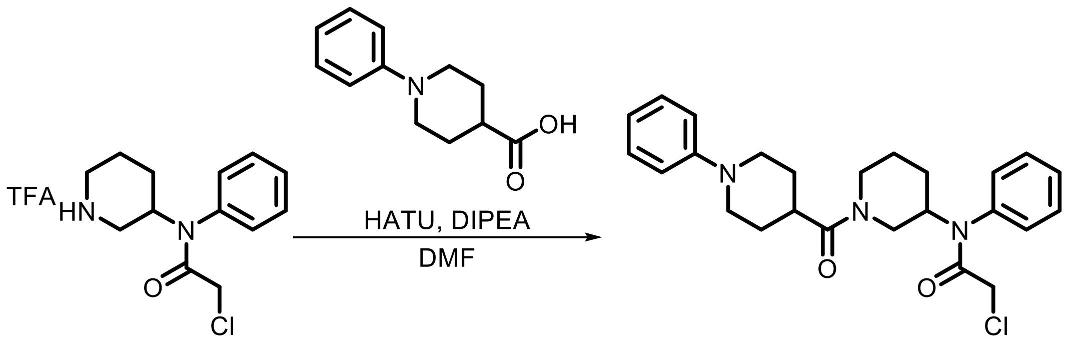

**SI-42** **EV-45**

Following **General Procedure 3**, starting with **SI-42** (248 mg, 0.98 mmol, 1.2 eq) and 1-phenylpiperidine-4-carboxylic acid (1.0 eq), **EV-45** (71 mg, 17%) was obtained after prep-HPLC (HCl).

**$^1$H NMR (400 MHz, DMSO-*d*$_6$) δ** 7.55 – 7.46 (m, 3H), 7.44 – 7.31 (m, 2H), 7.19 (q, *J* = 7.2 Hz, 2H), 6.93 (dd, *J* = 14.5, 8.1 Hz, 2H), 6.74 (q, *J* = 6.8 Hz, 1H), 4.38 – 4.18 (m, 2H), 3.96 – 3.60 (m, 5H), 2.88 – 2.63 (m, 4H), 2.21 (t, *J* = 11.9 Hz, 1H), 2.00 – 1.77 (m, 2H), 1.74 – 1.56 (m, 4H), 1.53 – 1.25 (m, 1H), 1.23 – 0.95 (m, 1H).

**HRMS ESI-TOF (*m/z*):** $[M+H]^+$ for $C_{25}C_{31}ClN_3O_2$ 440.2100, found 440.2104.

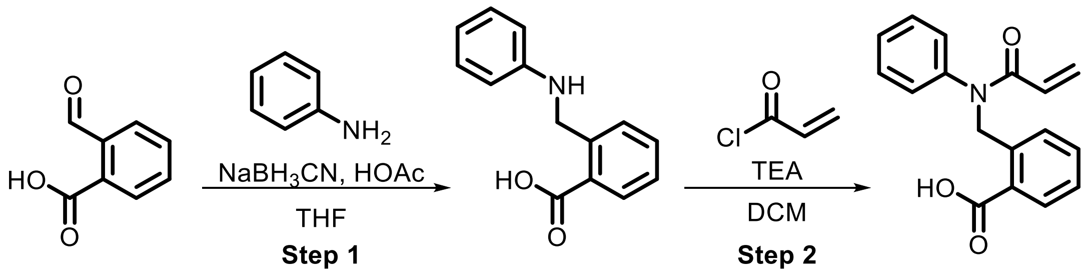

**SI-43** **EV-46**

**Step 1:** Following **General Procedure 5**, starting with 2-formylbenzoic acid (100 mg, 0.67 mmol, 1.0 eq) and aniline (1.0 eq), **SI-43** was obtained as a crude product and used in the next step without purification.

**Step 2:** Following **General Procedure 2**, starting with **SI-43** (190 mg, 0.66 mmol, 1.0 eq), acryloyl chloride (2.0 eq), and TEA (3.0 eq), **EV-46** was obtained after prep-HPLC (FA) as a yellow solid (49 mg, 26%).

**$^1$H NMR (400 MHz, CDCl$_3$) δ** 7.95 (d, *J* = 7.8 Hz, 1H), 7.56 – 7.45 (m, 2H), 7.39 – 7.24 (m, 4H), 7.19 – 7.13 (m, 2H), 6.44 (dd, *J* = 16.8, 2.0 Hz, 1H), 6.14 (dd, *J* = 16.8, 10.3 Hz, 1H), 5.59 (dd, *J* = 10.3, 1.9 Hz, 1H), 5.42 (s, 2H).

**HRMS ESI-TOF (*m/z*):** $[M+H]^+$ for $C_{17}H_{16}NO_3$ 282.1125, found 282.1127.

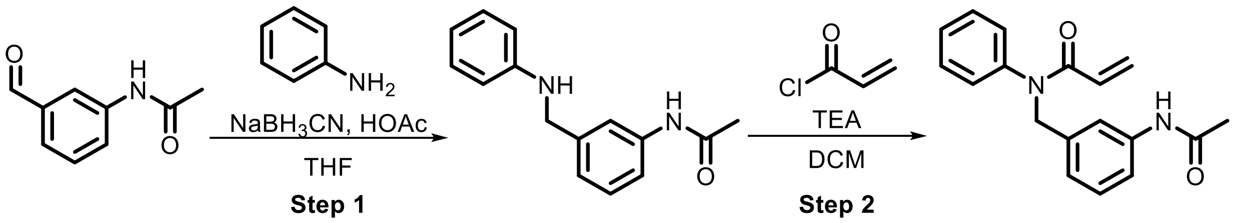

**SI-44** **EV-47**

**Step 1:** Following **General Procedure 5**, starting with *N*-(3-formylphenyl)acetamide (100 mg, 0.61 mmol, 1.0 eq), and aniline (1.0 eq), **SI-44** was obtained as a crude product and used in the next step without purification.

**Step 2:** Following **General Procedure 2**, starting with **SI-44** (1.0 eq), acryloyl chloride (2.0 eq), and TEA (3.0 eq), **EV-47** (86 mg, 48%) was obtained after prep-HPLC (FA).

**$^1$H NMR (400 MHz, CDCl$_3$) δ** 7.59 (d, *J* = 7.7 Hz, 1H), 7.40 (br s, 1H), 7.36 – 7.28 (m, 4H), 7.20 (t, *J* = 7.9 Hz, 1H), 7.03 (d, *J* = 6.4 Hz, 2H), 6.89 (d, *J* = 7.6 Hz, 1H), 6.41 (dd, *J* = 16.8, 2.0 Hz, 1H), 6.05 (dd, *J* = 16.8, 10.3 Hz, 1H), 5.55 (dd, *J* = 10.3, 2.0 Hz, 1H), 4.94 (s, 2H), 2.13 (s, 3H).

**HRMS ESI-TOF (*m/z*):** $[M+H]^+$ for $C_{18}H_{19}N_2O_2$ 295.1441, found 295.1442.

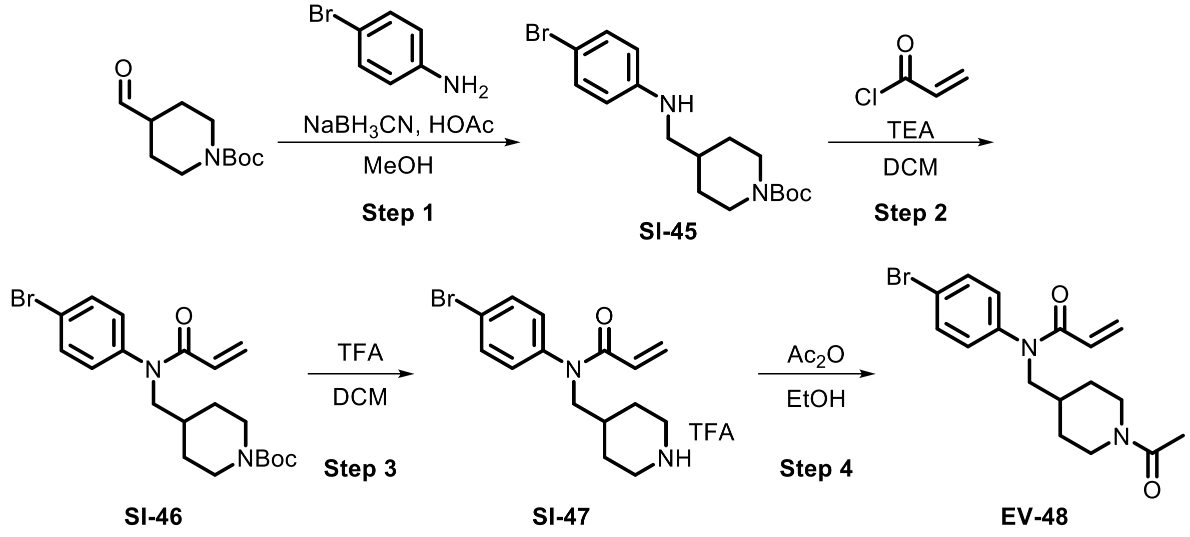

**Step 1:** Following **General Procedure 5**, starting with *tert*-butyl 4-formylpiperidine-1-carboxylate (130 mg, 0.61 mmol, 1.0 eq) and 4-bromoaniline (1.0 eq), **SI-45** (220 mg, crude) was obtained as a crude product and used in the next step without purification

**Step 2:** Following **General Procedure 2**, starting with **SI-45** (220 mg, 0.60 mmol, 1.0 eq), acryloyl chloride (2.0 eq), and TEA (3.0 eq), **SI-46** (250 mg, crude) was obtained as a crude product and used in the next step without purification.

**Step 3:** TFA (452 µL, 5.9 mmol, 10.0 eq) was added to a solution of **SI-46** (250 mg, 0.59 mmol, 1.0 eq) in DCM (2 mL). The mixture was stirred at 15 °C for 16 h. The reaction mixture was concentrated under reduced pressure to afford **SI-47** (250 mg, crude) as a yellow oil which was used without purification.

**Step 4:** $Ac_2O$ (108 µL, 1.1 mmol, 2.0 eq) was added to a solution of **SI-47** (250 mg, 0.57 mmol, 1.00 eq) in EtOH (4 mL) at room temperature. The mixture was heated to 60 °C and stirred for 3 h. The reaction mixture was purified by prep. HPLC (basic). The eluent was evaporated to remove organic solvents followed by lyophilization to afford **EV-48** (25 mg, 12%) as a yellow solid.

**$^1$H NMR (400 MHz, $CDCl_3$) δ** 7.58 – 7.52 (m, 2H), 7.09 – 7.00 (m, 2H), 6.37 (dd, *J* = 16.7, 1.9 Hz, 1H), 6.00 (dd, *J* = 16.7, 10.3 Hz, 1H), 5.56 (dd, *J* = 10.3, 1.9 Hz, 1H), 4.60 – 4.49 (m, 1H), 3.88 – 3.74 (m, 2H), 3.55 (dd, *J* = 13.5, 6.6 Hz, 1H), 3.03 – 2.91 (m, 1H), 2.51 (td, *J* = 12.8, 3.0 Hz, 1H), 2.05 (s, 3H), 1.92 – 1.78 (m, 1H), 1.77 – 1.70 (m, 1H), 1.65 – 1.60 (m, 1H), 1.33 – 1.08 (m, 2H).

**HRMS ESI-TOF (*m/z*):** $[M+H]^+$ for $C_{17}H_{22}BrN_2O_2$ 365.0859, found 365.0861.

**Synthesis of SI-48 as a precursor for EV-49 – EV-52**

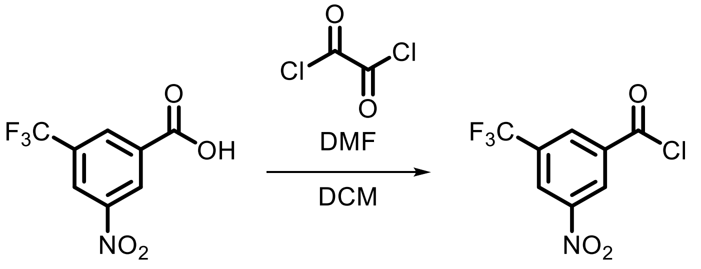

**SI-48**

Oxalyl chloride (93 µL, 1.1 mmol, 1.3 eq) and DMF (50 µL) were added to a solution of 3-nitro-5-(trifluoromethyl)benzoic acid (200 mg, 0.85 mmol, 1.0 eq) in DCM (2 mL). The mixture was stirred at 40 °C for 3 h. The reaction mixture was concentrated under reduced pressure to afford **SI-48** (250 mg, crude) as a light yellow oil, which was used without purification.

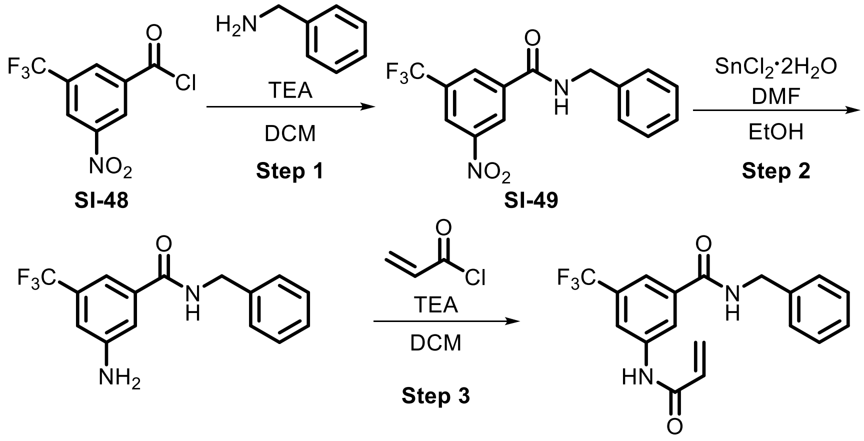

**SI-50** **EV-49**

**Step 1:** Following **General Procedure 4**, starting with **SI-48** (60 mg, 0.24 mmol, 1.0 eq), benzylamine (1.0 eq), and TEA (3.0 eq), **SI-49** (80 mg, crude) was obtained as crude product and used without purification.

**Step 2:** $SnCl_2 \cdot 2H_2O$ (223 mg, 1.0 mmol, 4.0 eq) and DMF (1 mol %) were added to a solution of **SI-49** (80 mg, 0.25 mmol, 1.0 eq) in EtOH (1 mL). The mixture was heated to 80 °C and stirred for 2 h. The reaction was then cooled, sat. aqueous $NaHCO_3$ (2 mL) was added, and the reaction was stirred for 5 min. The mixture was extracted with DCM (2 mL x 3). The combined organic layers were dried over anhydrous $Na_2SO_4$ and concentrated to afford **SI-50** (90 mg, crude) as light yellow oil.

**Step 3:** Following **General Procedure 2**, starting with **SI-50** (90 mg, 0.31 mmol, 1.0 eq), acryloyl chloride (0.8 eq), and DMF (1 mol %), **EV-49** was obtained as a white solid (22 mg, 21%) after prep-HPLC (FA).

**$^1$H NMR (400 MHz, DMSO-*d*$_6$) δ** 10.68 (s, 1H), 9.31 (t, *J* = 5.9 Hz, 1H), 8.35 (d, *J* = 10.8 Hz, 2H), 7.96 (s, 1H), 7.37 – 7.31 (m, 4H), 7.31 – 7.21 (m, 1H), 6.50 – 6.26 (m, 2H), 5.84 (dd, *J* = 9.9, 2.1 Hz, 1H), 4.50 (d, *J* = 5.8 Hz, 2H).

**HRMS ESI-TOF (*m/z*):** $[M+H]^+$ for $C_{18}H_{16}F_3N_2O_2$ 349.1159, found 349.1161.

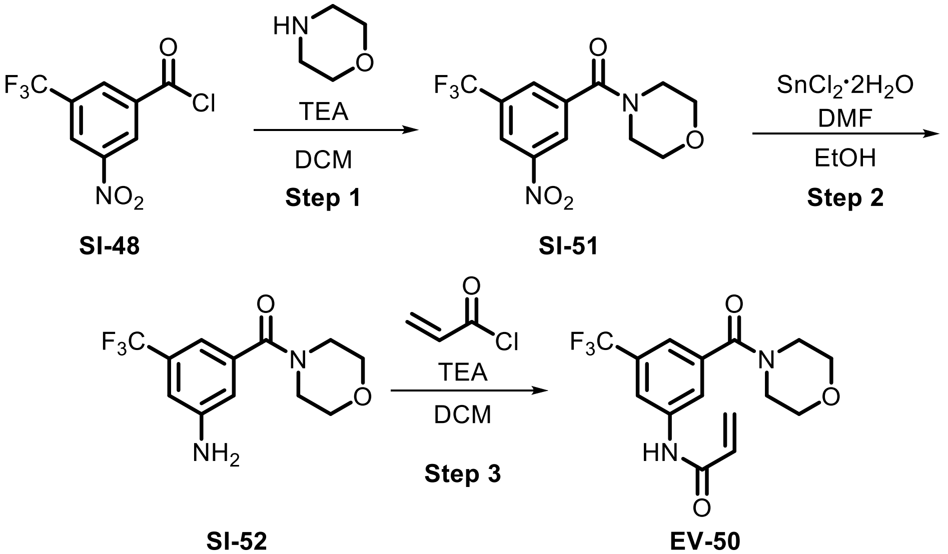

**Step 1:** Following **General Procedure 4**, starting with **SI-48** (60 mg, 0.24 mmol, 1.0 eq), benzylamine (1.0 eq), and TEA (3.0 eq), **SI-51** (80 mg, crude) was obtained as crude product and used without purification.

**Step 2:** $SnCl_2 \cdot 2H_2O$ (237 mg, 1.1 mmol, 4.0 eq) and DMF (1 mol %) were added to a solution of **SI-51** (80 mg, 0.26 mmol, 1.0 eq) in EtOH (1 mL). The mixture was stirred at 80 °C for 2 h. The reaction was cooled, sat. aqueous $NaHCO_3$ (2 mL) was added, and the mixture was stirred for 5 min. The reaction mixture was extracted with DCM (2 mL x 3) and the combined organic layers were dried over anhydrous $Na_2SO_4$ and concentrated to afford **SI-52** (90 mg, crude) as a light yellow oil.

**Step 3:** Following **General Procedure 2**, starting with **SI-52** (90 mg, 0.31 mmol, 1.0 eq), acryloyl chloride (0.8 eq), and DMF (1 mol %), **EV-50** was obtained as a white solid (22 mg, 25%) after prep-HPLC (FA)

**$^1$H NMR (400 MHz, DMSO-*d*$_6$) δ** 8.13 (s, 1H), 7.85 (s, 1H), 7.43 (s, 1H), 6.44 – 6.26 (m, 2H), 5.83 (dd, *J* = 9.7, 2.0 Hz, 1H), 3.65 – 3.59 (m, 4H), 3.52 (br s, 2H), 3.30 (br s, 2H)

**HRMS ESI-TOF (*m/z*):** $[M+H]^+$ for $C_{15}H_{16}F_3N_2O_3$ 329.1108, found 329.1107.

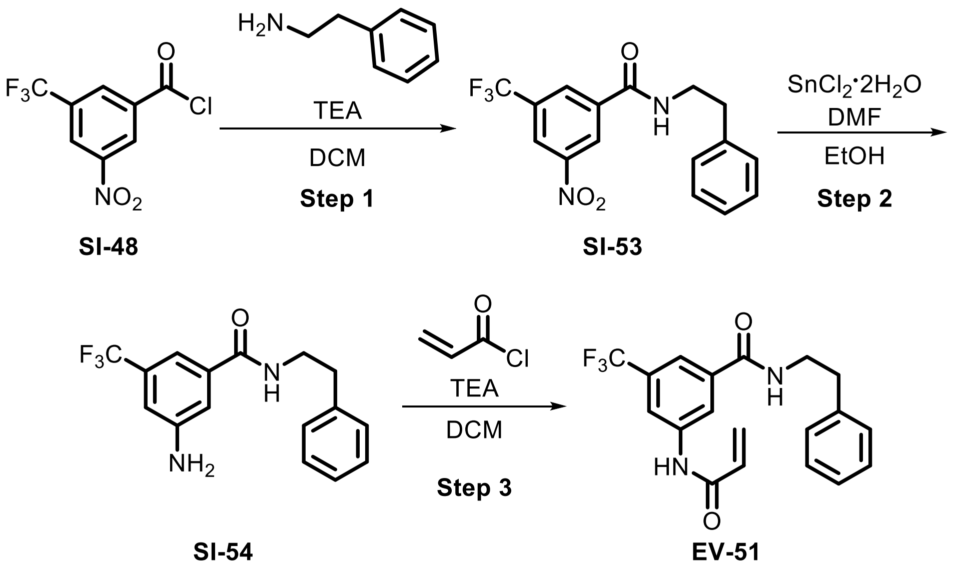

**Step 1:** Following **General Procedure 4**, starting with **SI-48** (60 mg, 0.24 mmol, 1.0 eq), phenethylamine (1.0 eq), and TEA (3.0 eq), **SI-53** (80 mg, crude) was obtained as crude product and used without purification.

**Step 2:** $SnCl_2 \cdot 2H_2O$ (213 mg, 0.95 mmol, 4.0 eq) and DMF (1 mol %) were added to a solution of **SI-53** (80 mg, 0.24 mmol, 1.0 eq) in EtOH (1 mL). The mixture was stirred at 80 °C for 2 h. The reaction was cooled to rt, sat. aqueous $NaHCO_3$ (2 mL) was added, and the mixture was stirred for 5 min. The mixture was extracted with DCM (2 mL x 3), the combined organic layers were dried over anhydrous $Na_2SO_4$ and concentrated to afford **SI-54** (90 mg, crude) as light yellow oil.

**Step 3:** Following **General Procedure 2**, starting with **SI-54** (90 mg, 0.31 mmol, 1.0 eq), acryloyl chloride (0.8 eq), and DMF (1 mol %), **EV-51** was obtained after prep-HPLC (FA) as a white solid (22 mg, 17%).

**$^1$H NMR (400 MHz, DMSO-*d*$_6$) δ** 10.66 (s, 1H), 8.84 (t, *J* = 5.6 Hz, 1H), 8.31 (d, *J* = 20.2 Hz, 2H), 7.86 (s, 1H), 7.34 – 7.15 (m, 5H), 6.50 – 6.28 (m, 2H), 5.84 (dd, *J* = 9.9, 2.1 Hz, 1H), 3.50 (q, *J* = 6.8 Hz, 2H), 2.86 (t, *J* = 7.5 Hz, 2H).

**HRMS ESI-TOF (*m/z*):** $[M+H]^+$ for $C_{19}H_{18}F_3N_2O_2$ 363.1315, found 363.1317.

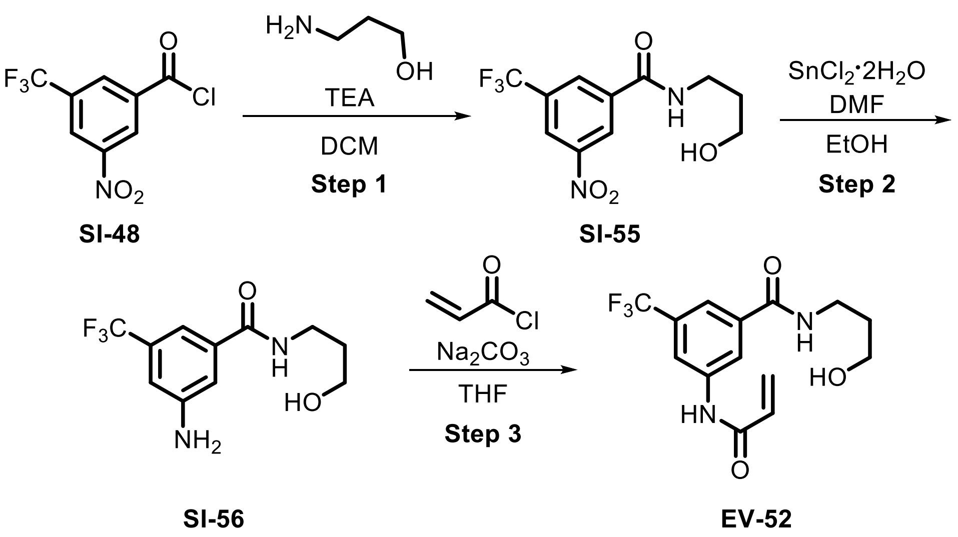

**Step 1:** Following **General Procedure 4**, starting with **SI-48** (215 mg, 0.85 mmol, 1.0 eq), phenethylamine (1.0 eq), and TEA (3.0 eq), **SI-55** (200 mg, crude) was obtained as crude product and used without purification.

**Step 2:** $SnCl_2{\cdot}2H_2O$ (247 mg, 1.1 mmol, 4.0 eq) and DMF (1 mol %) were added to a solution of **SI-56** (80 mg, 0.27 mmol, 1.0 eq) in EtOH (1.00 mL) at 25 °C. Following the addition, the mixture was stirred at 80 °C for 2 h. The reaction mixture was quenched with $H_2O$ (1 mL) and saturated $NaHCO_3$ (2 mL). The mixture was then extracted with DCM (2 mL x 3). The combined organic layers were dried over anhydrous $Na_2SO_4$ and concentrated under reduced pressure to provide **SI-56** (70 mg, 85%) as a yellow oil.

**Step 3:** A solution of $Na_2CO_3$ (28 mg, 0.27 mmol, 2.0 eq) in $H_2O$ (0.2 mL) was added to a solution of **SI-56** (35 mg, 0.13 mmol, 1.0 eq) in THF (0.4 mL) in one portion. The mixture was cooled to 10 °C and acryloyl chloride (12 µL, 0.15 mmol, 1.1 eq) was added. Following the addition, the mixture was warmed to 25 °C and stirred for 16 h. The reaction mixture was quenched with $H_2O$ (2 mL) and extracted with DCM (2 ml x 4). The combined organic layers were dried over anhydrous $Na_2SO_4$ and concentrated under reduced pressure. The resulting residue was diluted with MeCN (2 mL) and purified by prep-HPLC (FA) to afford **EV-52** (15 mg, 35%) as colorless oil.

**$^1$H NMR (400 MHz, DMSO-*d*$_6$) δ** 10.70 (s, 1H), 8.72 (t, *J* = 5.5 Hz, 1H), 8.32 (d, *J* = 20.8 Hz, 2H), 7.89 (s, 1H), 6.50 – 6.28 (m, 2H), 5.83 (dd, *J* = 9.8, 2.0 Hz, 1H), 4.50 (br s, 1H), 3.47 (t, *J* = 6.3 Hz, 2H), 3.33 (d, *J* = 6.7 Hz, 2H), 1.69 (p, *J* = 6.7 Hz, 2H).

**HRMS ESI-TOF (*m/z*):** $[M+H]^+$ for $C_{14}H_{16}F_3N_2O_3$ 317.1108, found 317.1109.

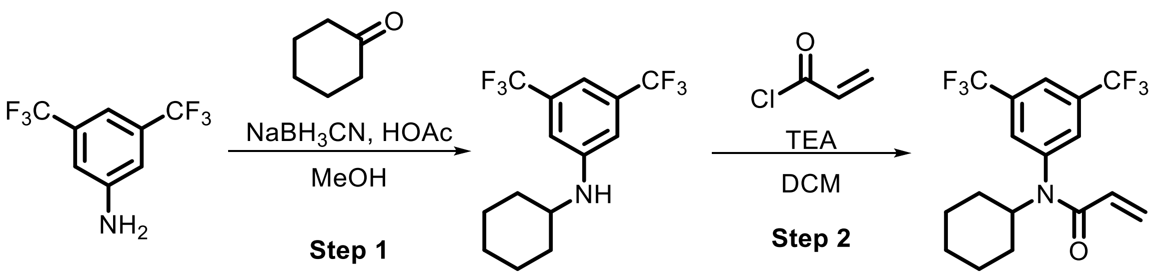

**SI-57** **EV-53**

**Step 1:** Following **General Procedure 5**, starting with 3,5-bis(trifluoromethyl)aniline (200 mg, 0.87 mmol, 1.0 eq) and cyclohexanone (1.0 eq), **SI-57** (77 mg, 28%) was obtained as a white solid.

**Step 2:** Following **General Procedure 2**, starting with **SI-57** (77 mg, 0.25 mmol, 1.0 eq), acryloyl chloride (3.0 eq), and TEA (3.0 eq), **EV-53** was obtained after prep-HPLC (FA) as a white solid (45 mg, 47%).

**$^1$H NMR (400 MHz, $CDCl_3$) δ** 7.93 (s, 1H), 7.55 (s, 2H), 6.40 (dd, *J* = 16.5, 2.0 Hz, 1H), 5.69 (s, 1H), 5.56 (dd, *J* = 10.4, 2.1 Hz, 1H), 4.67 (s, 1H), 1.89 (d, *J* = 11.9 Hz, 2H), 1.79 (d, *J* = 13.6 Hz, 2H), 1.62 (d, *J* = 15.5 Hz, 1H), 1.43 (q, *J* = 14.0, 12.0 Hz, 2H), 1.08 – 0.86 (m, 3H).

**HRMS ESI-TOF (*m/z*):** $[M+H]^+$ for $C_{17}H_{18}F_6NO$ 366.1287, found 366.1288.

**Synthesis of SI-58 as a precursor for EV-54 – EV-60**

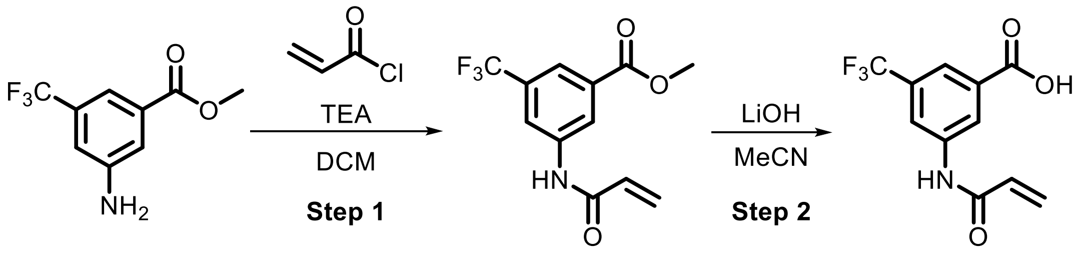

**SI-58** **SI-59**

**Step 1:** Following **General Procedure 2**, starting with methyl 3-amino-5-(trifluoromethyl)benzoate (1.0 g, 4.6 mmol, 1.0 *eq*), acryloyl chloride (1.2 *eq*), and TEA (3.0 *eq*), **SI-58** was obtained after flash chromatography as a yellow oil (2.8 g, 56%).

**Step 2:** A solution of LiOH (210 mg, 8.8 mmol, 1.2 *eq*) in $H_2O$ (5 mL) was added to a solution of **SI-58** (2.0 g, 7.3 mmol, 1.0 *eq*) in MeCN (15 mL) and stirred at 25 °C for 3 h. The reaction mixture was concentrated and the pH was adjusted to pH 3 with citric acid. The precipitate was collected by filtration to provide **SI-59** (1.0 g, 53%) as a light yellow solid. **SI-59** was used without purification.

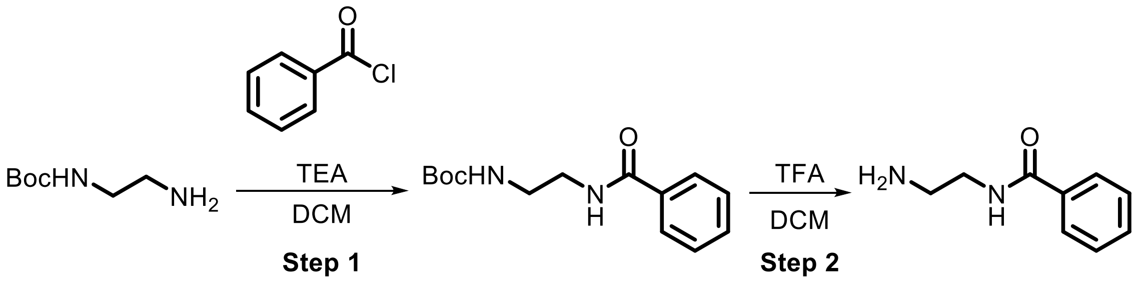

**SI-60** **SI-61**

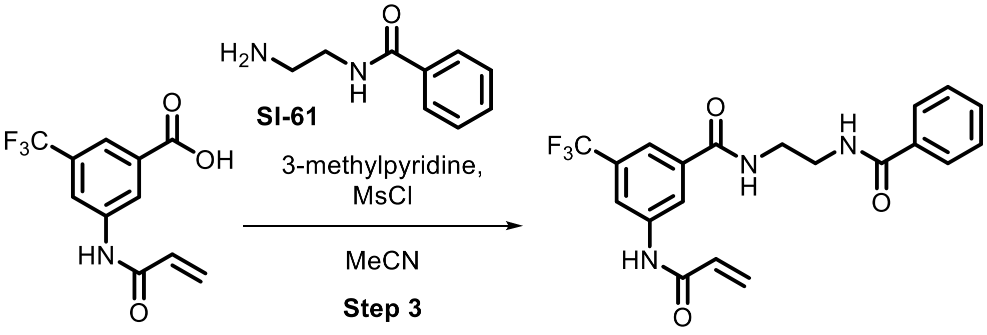

**SI-59** **EV-54**

**Step 1:** Following **General Procedure 4**, starting with *tert*-butyl *N*-(2-aminoethyl)carbamate (2.0 g, 12.5 mmol, 1.0 *eq*) and benzoyl chloride (1.5 *eq*), **SI-60** was obtained as a crude product and used in the next step without purification.

**Step 2:** TFA (3.6 mL, 49.2 mmol, 5.0 *eq*) was added to a solution of **SI-60** (2.6 g, 9.8 mmol, 1.0 *eq*) in DCM (10 mL) and the reaction mixture was stirred at 25 °C for 2 h. The reaction mixture was quenched with aqueous $NaHCO_3$ (20 mL) and extracted with DCM (5 mL x 3). The combined organic layers were dried over anhydrous $Na_2SO_4$ and concentrated under reduced pressure to afford **SI-61** (1.20 g, crude) as a yellow oil which was used for the synthesis of **EV-54** without purification.

**Step 3:** MsCl (36 µL, 0.46 mmol, 1.2 *eq*) was added to a solution of **SI-59** (100 mg, 0.62 mmol, 1.0 *eq*), **SI-61** (70 mg, 0.42 mmol, 1.1 *eq*), and 3-methylpyridine (112 µL, 1.2 mmol, 3.0 *eq*) in MeCN (2 mL) and the resulting mixture was stirred at 25 °C for 2 h. The reaction mixture was quenched with $H_2O$ (5 mL) and extracted with DCM (5 mL x 3). The combined organic layers were dried over anhydrous $Na_2SO_4$ and concentrated under reduced pressure. The resulting residue was purified by prep-HPLC (basic) to provide **EV-54** (90 mg, 56%) as a white solid.

**$^1$H NMR (400 MHz, MeOD) δ** 8.25 (d, *J* = 1.6 Hz, 2H), 7.87 – 7.79 (m, 3H), 7.53 (t, *J* = 7.3 Hz, 1H), 7.48 – 7.42 (m, 2H), 6.47 – 6.40 (m, 2H), 5.84 (dd, *J* = 6.7, 5.1 Hz, 1H), 4.59 (br s, 1H), 3.64 (s, 4H), 3.48 (t, *J* = 1.7 Hz, 1H), 3.13 (t, *J* = 1.7 Hz, 1H).

**HRMS ESI-TOF (*m/z*):** $[M+H]^+$ for $C_{20}H_{19}F_3N_3O_3$ 406.1373, found 406.1375.

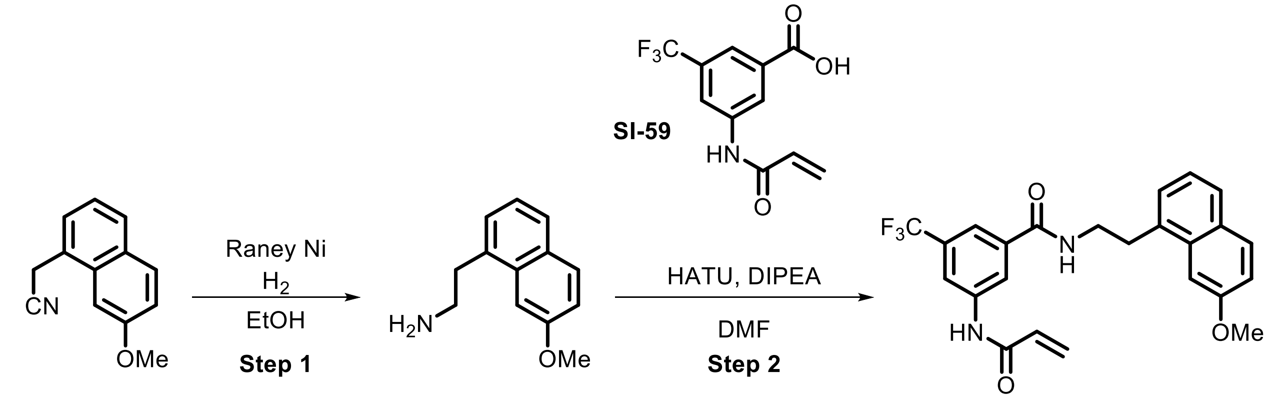

**SI-62** **EV-55**

**Step 1:** Raney-Ni (20 mg, 0.23 mmol, 0.2 *eq*) was added to a solution of 7-methoxy-1-naphthylacetonitrile (200 mg, 1.0 mmol, 1.0 eq) and $NH_3 \cdot H_2O$ (500 µL) in EtOH (2 mL). The reaction vessel was evacuated and backfilled with $H_2$ (3x). The reaction was stirred for 16 h at 45 psi $H_2$ at 60 °C. The mixture was filtered and the filtrate was concentrated to provide **SI-62** (200 mg, crude) as a colorless oil. **SI-62** was used for the synthesis of **EV-55** without purification.

**Step 2:** Following **General Procedure 3**, starting with **SI-59** (100 mg, 0.39 mmol, 1.0 *eq*) and **SI-62** (1.1 *eq*), **EV-55** was obtained as a white solid (42 mg, 24%).

**$^1$H NMR (400 MHz, MeOD) δ** 8.24 (d, *J* = 7.2 Hz, 2H), 7.80 – 7.71 (m, 2H), 7.67 (d, *J* = 8.3 Hz, 1H), 7.60 (s, 1H), 7.36 (d, *J* = 6.8 Hz, 1H), 7.29 – 7.22 (m, 1H), 7.12 (dd, *J* = 8.9, 2.6 Hz, 1H), 6.43 (d, *J* = 6.6 Hz, 2H), 5.88 – 5.80 (m, 1H), 3.95 (s, 3H), 3.72 (t, *J* = 7.9 Hz, 2H), 3.37 (t, *J* = 7.3 Hz, 2H).

**HRMS ESI-TOF (*m/z*):** $[M+H]^+$ for $C_{24}H_{22}F_3N_2O_3$ 443.1577, found 443.1579.

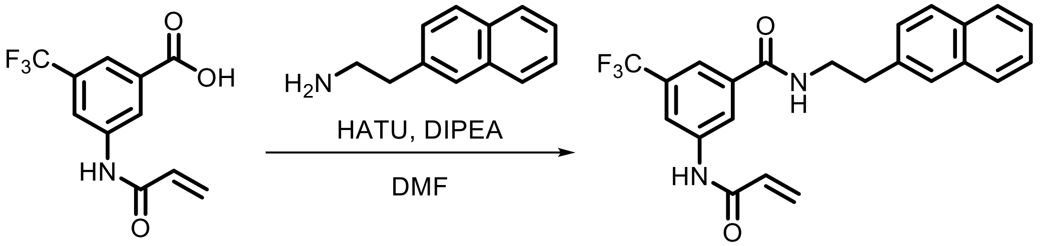

**SI-59** **EV-56**

Following **General Procedure 3**, starting with **SI-59** (50 mg, 0.19 mmol, 1.0 *eq*) and 2-(naphthalen-2-yl)ethan-1-amine (1.0 *eq*), **EV-56** was obtained after prep-HPLC (basic) as a white solid (5 mg, 6%).

**$^1$H NMR (400 MHz, $CDCl_3$) δ** 8.18 (s, 1H), 8.06 (s, 1H), 7.93 (s, 1H), 7.86 – 7.77 (m, 3H), 7.66 (d, *J* = 20.1 Hz, 2H), 7.52 – 7.42 (m, 2H), 7.38 (d, *J* = 8.1 Hz, 1H), 6.48 (d, *J* = 16.8 Hz, 1H), 6.41 – 6.33 (m, 1H), 6.30 (dd, *J* = 16.8, 10.2 Hz, 1H), 5.83 (d, *J* = 10.2 Hz, 1H), 3.80 (q, *J* = 6.7 Hz, 2H), 3.11 (t, *J* = 7.0 Hz, 2H).

**HRMS ESI-TOF (*m/z*):** $[M+H]^+$ for $C_{23}H_{19}F_3N_2O_2$ 413.1472, found 413.1470.

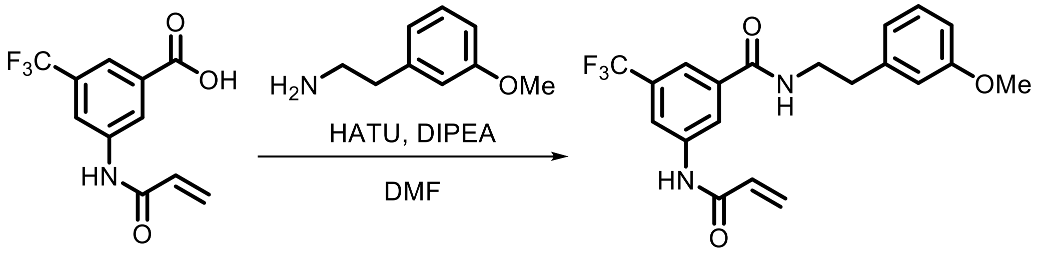

**SI-59** **EV-57**

Following **General Procedure 3**, starting with **SI-59** (100 mg, 0.39 mmol, 1.0 *eq*) and 3-methoxyphenethylamine (1.2 *eq*), **EV-57** was obtained after prep-HPLC (basic) as a white solid (28 mg, 17%).

**$^1$H NMR (400 MHz, MeOD) δ** 8.24 (d, *J* = 8.1 Hz, 2H), 7.76 (s,10H), 7.20 (t, *J* = 8.2 Hz, 1H), 6.84 (d, *J* = 8.1 Hz, 2H), 6.77 (dd, *J* = 7.6, 2.1 Hz, 1H), 6.46 – 6.40 (m, 2H), 5.84 (dd, *J* = 7.0, 4.9 Hz, 1H), 3.76 (s, 3H), 3.61 (t, *J* = 7.4 Hz, 2H), 2.90 (t, *J* = 7.4 Hz, 2H).

**HRMS ESI-TOF (*m/z*):** $[M+H]^+$ for $C_{20}H_{20}F_3N_2O_3$ 393.1421, found 393.1423.

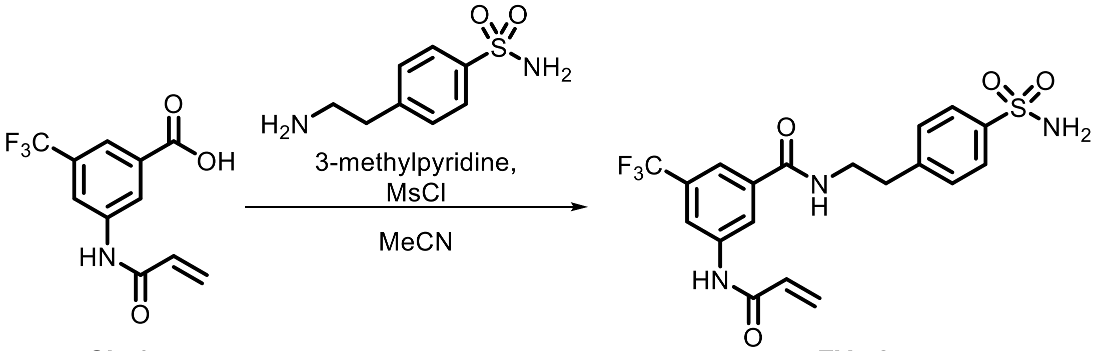

**SI-59** **EV-58**

3-methylpyridine (108 mg, 1.2 mmol, 3.0 *eq*) and MsCl (30 µL, 0.39 mmol, 1.0 *eq*) were added to a solution of **SI-59** (100 mg, 0.39 mmol, 1.0 *eq*) and 4-(2-Aminoethyl)benzenesulfonamide (77 mg, 0.39 mmol, 1.0 *eq*) in MeCN (1 mL) at 0 °C. The reaction was warmed to 25 °C and stirred for 3 h. The reaction mixture was quenched with water (1 mL) and concentrated. The resulting residue was purified via prep-HPLC (basic) to provide **EV-58** (64 mg, 37%) as an off-white solid.

**$^1$H NMR (400 MHz, MeOD) δ** 8.23 (d, *J* = 22.5 Hz, 2H), 7.84 (d, *J* = 8.4 Hz, 2H), 7.78 (s, 1H), 7.45 (d, *J* = 8.3 Hz, 2H), 6.47 – 6.40 (m, 2H), 5.84 (t, *J* = 5.7 Hz, 1H), 4.60 (s, 2H), 3.66 (t, *J* = 7.2 Hz, 2H), 3.03 (t, *J* = 7.2 Hz, 2H).

**HRMS ESI-TOF (*m/z*):** $[M+H]^+$ for $C_{19}H_{19}F_3N_2O_4S$ 442.1043, found 442.1043.

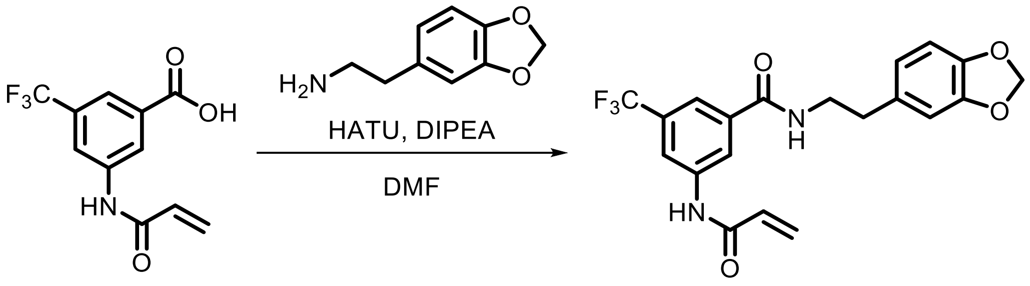

**SI-59** **EV-59**

Following **General Procedure 3**, starting with **SI-59** (100 mg, 0.39 mmol, 1.0 *eq*) and homopiperonylamine (1.0 *eq*), **EV-59** was obtained after prep-HPLC (basic) as a white solid (8 mg, 5%).

**$^1$H NMR (400 MHz, DMSO-*d*$_6$) δ** 10.63 (s, 1H), 8.79 (t, *J* = 5.6 Hz, 1H), 8.30 (d, *J* = 25.7 Hz, 2H), 7.85 (s, 1H), 6.86 – 6.80 (m, 2H), 6.69 (dd, *J* = 7.9, 1.7 Hz, 1H), 6.43 (dd, *J* = 17.0, 9.9 Hz, 1H), 6.32 (dd, *J* = 17.0, 2.1 Hz, 1H), 5.96 (s, 2H), 5.84 (dd, *J* = 9.9, 2.1 Hz, 1H), 3.46 (q, *J* = 7.0 Hz, 2H), 2.77 (t, *J* = 7.3 Hz, 2H).

**HRMS ESI-TOF (*m/z*):** $[M+H]^+$ for $C_{20}H_{18}F_3N_2O_4$ 407.1213, found 407.1213.

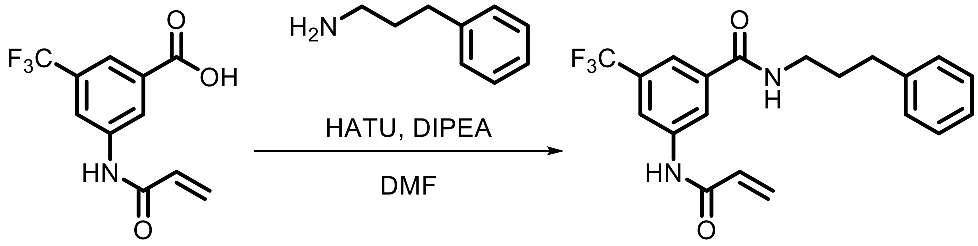

**SI-59** **EV-60**

Following **General Procedure 3**, starting with **SI-59** (100 mg, 0.39 mmol, 1.0 *eq*) and 3-phenylpropylamine (1.2 *eq*), **EV-60** was obtained after prep-HPLC (basic) as a white solid (5 mg, 3%).

**$^1$H NMR (400 MHz, MeOD) δ** 8.25 (s, 2H), 7.80 (s, 1H), 7.29 – 7.19 (m, 4H), 7.19 – 7.10 (m, 1H), 6.48 – 6.38 (m, 2H), 5.84 (dd, *J* = 7.1, 4.7 Hz, 1H), 4.60 (br s, 1H), 3.42 (t, *J* = 7.1 Hz, 2H), 2.71 (t, *J* = 7.6 Hz, 2H), 1.96 (p, *J* = 7.5 Hz, 2H).

**HRMS ESI-TOF (*m/z*):** $[M+H]^+$ for $C_{20}H_{20}F_3N_2O_2$ 377.1472, found 377.1474.

**Synthesis of SI-63 as a precursor for EV-61 and EV-62**

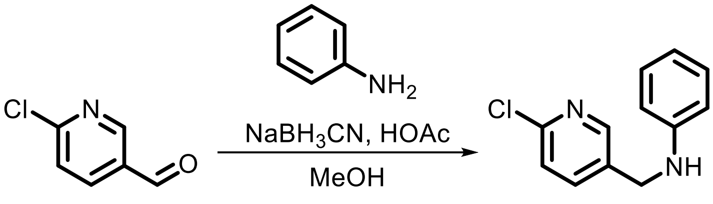

**SI-63**

Following **General Procedure 5**, starting with 6-chloropyridine-3-carbaldehyde (100 mg, 0.71 mmol, 1.0 eq) and aniline (1.0 eq), **SI-63** was obtained as crude product and used without purification.

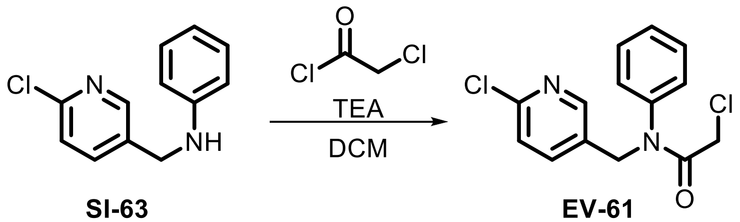

**SI-63** **EV-61**

Following **General Procedure 1**, starting with **SI-63** (70 mg, 0.32 mmol, 1.0 eq), chloroacetyl chloride (2.0 eq), and TEA (3.0 eq), **EV-61** was obtained after prep-HPLC (HCl) as a colorless oil (56 mg, 59%).

**$^1$H NMR (400 MHz, $CDCl_3$) δ** 8.13 (s, 1H), 7.72 (d, *J* = 7.6 Hz, 1H), 7.40 (s, 3H), 7.32 (d, *J* = 7.5 Hz, 1H), 7.03 (s, 2H), 4.87 (s, 2H), 3.83 (s, 2H).

**HRMS ESI-TOF (*m/z*):** $[M+H]^+$ for $C_{14}H_{13}Cl_2N_2O$ 295.0400, found 295.0404.

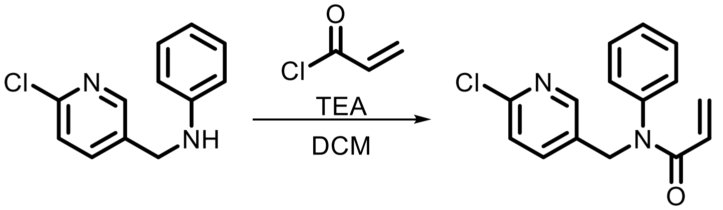

**SI-63** **EV-62**

Following **General Procedure 2**, starting with **SI-63** (70 mg, 0.32 mmol, 1.0 eq), acryloyl chloride (2.0 eq), and TEA (3.0 eq), **EV-62** (19 mg, 22%) was obtained after prep-HPLC.

**$^1$H NMR (400 MHz, $CDCl_3$) δ** 8.13 (d, *J* = 2.5 Hz, 1H), 7.67 (dd, *J* = 8.2, 2.5 Hz, 1H), 7.39 – 7.33 (m, 3H), 7.26 (d, *J* = 8.2 Hz, 1H), 6.99 (dd, *J* = 7.9, 1.8 Hz, 2H), 6.42 (dd, *J* = 16.8, 1.9 Hz, 1H), 6.01 (dd, *J* = 16.8, 10.3 Hz, 1H), 5.57 (dd, *J* = 10.3, 2.0 Hz, 1H), 4.94 (s, 2H).

**HRMS ESI-TOF (*m/z*):** $[M+H]^+$ for $C_{15}H_{14}ClN_2O$ 273.0789, found 273.0791.

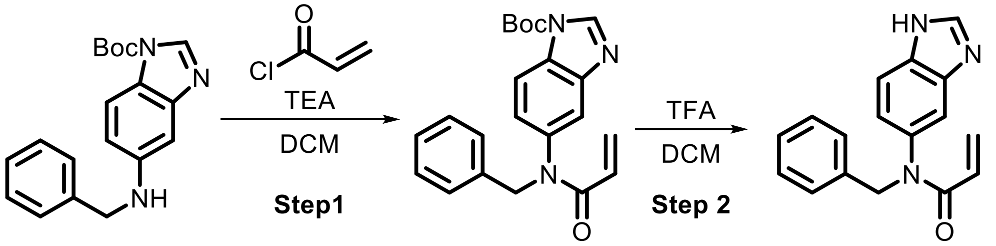

**SI-64** **EV-63**

**Step 1:** Following **General Procedure 2**, starting with *tert*-butyl 5-(benzylamino)-1*H*-benzo[*d*]imidazole-1-carboxylate (100 mg, 0.3 mmol, 1.0 eq.),(Bar-Peled et al., 2017) acryloyl chloride (2.0 eq), and TEA (3.0 eq), **SI-64** was obtained as crude product and used in the next step without purification.

**Step 2:** Following **General Procedure 6**, starting with **SI-64** (1.0 eq) and TFA (24 eq), **EV-63** (20 mg, 24% over 2 steps) after prep-HPLC (basic).

**$^1$H NMR (400 MHz, DMSO-$d_6$) δ** 8.24 (s, 1H), 7.55 (d, *J* = 8.4 Hz, 1H), 7.30 – 7.25 (m, 3H), 7.24 – 7.18 (m, 3H), 6.93 (dd, *J* = 8.4, 2.0 Hz, 1H), 6.22 (dd, *J* = 16.8, 2.5 Hz, 1H), 6.00 (dd, *J* = 16.8, 10.2 Hz, 1H), 5.56 (dd, *J* = 10.2, 2.5 Hz, 1H), 4.98 (s, 2H).

**HRMS ESI-TOF (*m/z*):** $[M+H]^+$ for $C_{17}H_{16}N_3O$ 278.1288, found 278.1290.

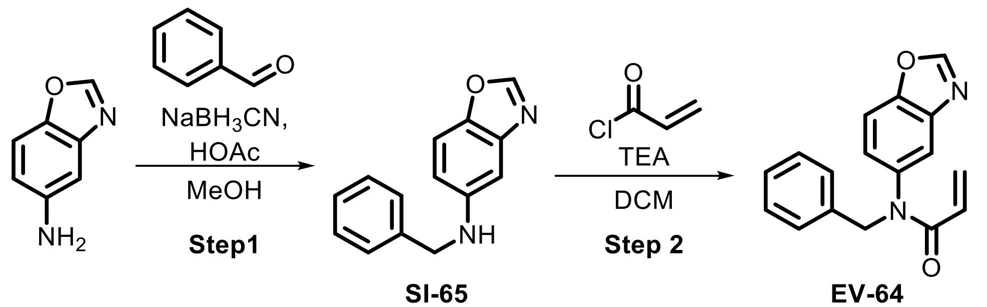

**SI-65** **EV-64**

**Step 1:** Following **General Procedure 5**, starting with benzo[*d*]oxazol-5-amine (1.0 eq) and benzaldehyde (1.1 eq), **SI-65** (55 mg, crude) was obtained and used in the next step without purification.

**Step 2:** Following **General Procedure 2**, starting with **SI-65** (55 mg, 0.24 mmol, 1.0 eq.), acryloyl chloride (2.0 eq), and TEA (3.0 eq), **EV-64** (30 mg, 44%) was obtained after prep-HPLC (basic).

**$^1$H NMR (400 MHz, $CDCl_3$) δ** 8.15 (s, 1H), 7.54 (d, *J* = 8.6 Hz, 1H), 7.46 (s, 1H), 7.31 – 7.20 (m, 5H), 7.04 (dd, *J* = 8.6, 2.1 Hz, 1H), 6.46 (dd, *J* = 16.8, 2.0 Hz, 1H), 6.00 (dd, *J* = 16.8, 10.3 Hz, 1H), 5.56 (dd, *J* = 10.3, 2.0 Hz, 1H), 5.04 (s, 2H).

**HRMS ESI-TOF (*m/z*):** $[M+H]^+$ for $C_{17}H_{15}N_2O_2$ 279.1128, found 279.1129.

### Synthesis of SI-67 as a precursor for EV-65 and EV-66

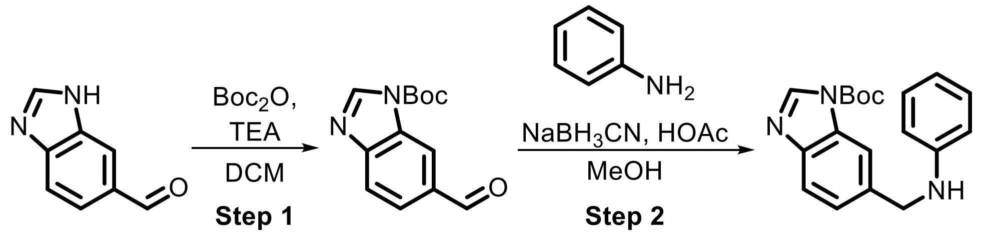

**SI-66** **SI-67**

**Step 1:** $Boc_2O$ (1.57 mL, 6.8 mmol, 2.0 eq) was added in one portion to a mixture of 3H-benzimidazole-5-carbaldehyde (500 mg, 3.4 mmol, 1.0 eq) and TEA (948 µL, 6.8 mmol, 2.0 eq) in DCM (5 mL) at 25 °C and was stirred for 16 h. The reaction mixture was filtered and the filtrate was concentrated. The resulting residue was purified by flash chromatography to afford **SI-66** (800 mg, crude) as a yellow oil.

**Step 2:** Following **General Procedure 5**, starting with **SI-66** (180 mg, 0.73 mmol, 1.0 eq) and aniline (1.1 eq), **SI-67** was obtained as crude product and used without purification.

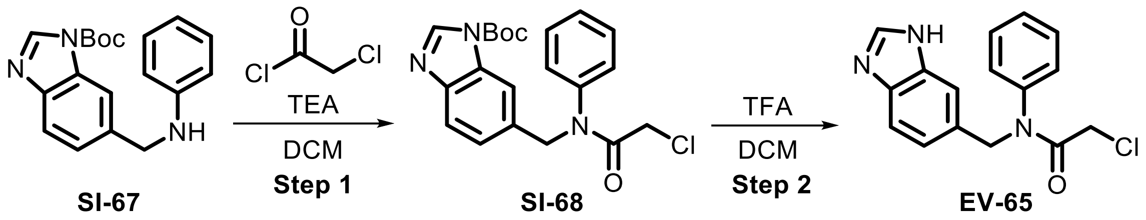

**Step 1:** Following **General Procedure 1**, starting with **SI-67** (200 mg, 0.62 mmol, 1.0 eq), chloroacetyl chloride (3.0 eq), and TEA (5.0 eq), **SI-68** was obtained as crude product and used in the next step without purification.

**Step 2:** Following **General Procedure 6**, starting with **SI-68** (1.0 eq) and TFA (24 eq), **EV-65** (46 mg, 13% over 2 steps) was obtained after prep-HPLC (basic).

**NMR (400 MHz, DMSO-$d_6$) δ** 9.58 (s, 1H), 7.79 (d, *J* = 8.4 Hz, 1H), 7.65 (s, 1H), 7.45 – 7.31 (m, 4H), 7.23 (d, *J* = 7.3 Hz, 2H), 5.06 (s, 2H), 4.07 (s, 2H).

**HRMS ESI-TOF (*m/z*):** $[M+H]^+$ for $C_{16}H_{15}ClN_3O$ 300.0898, found 300.0899.

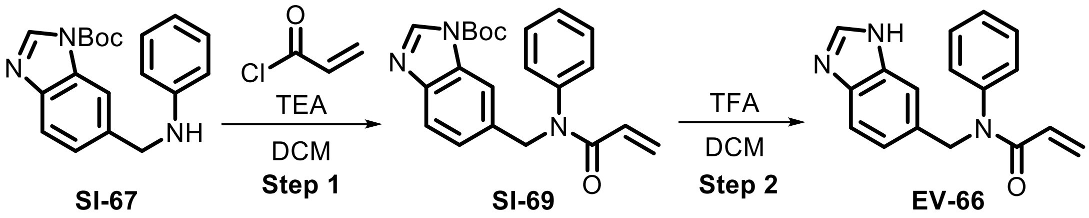

**Step 1:** Following **General Procedure 2**, starting with **SI-67** (200 mg, 0.62 mmol, 1.0 eq), acryloyl chloride (3.0 eq), and TEA (5.0 eq), **SI-69** was obtained as crude product and used in the next step without purification.

**Step 2:** Following **General Procedure 6**, starting with **SI-69** (1.0 eq) and TFA (24 eq), **EV-66** was obtained after prep-HPLC (basic) as a white solid (40 mg, 27% over 2 steps).

**$^1$H NMR (400 MHz, DMSO-$d_6$) δ** 8.16 (s, 1H), 7.49 (d, *J* = 8.2 Hz, 1H), 7.40 – 7.33 (m, 3H), 7.33 – 7.26 (m, 1H), 7.09 (d, *J* = 7.6 Hz, 2H), 7.03 (d, *J* = 8.3 Hz, 1H), 6.26 (dd, *J* = 16.8, 2.4 Hz, 1H), 6.02 (br s, 1H), 5.62 (dd, *J* = 10.2, 2.4 Hz, 1H), 5.06 (s, 2H).

**HRMS ESI-TOF (*m/z*):** $[M+H]^+$ for $C_{17}H_{16}N_3O$ 278.1288, found 278.1292.

**Synthesis of SI-70 as a precursor for EV-67 and EV-68**

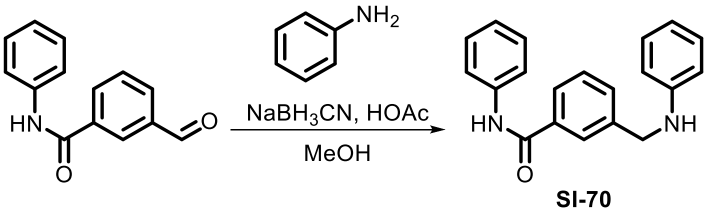

Following **General Procedure 5**, starting with 3-formyl-*N*-phenylbenzamide (750 mg, 3.3 mmol, 1.0 eq) and aniline (1.1 eq), **SI-70** was obtained and used in the next step without purification.

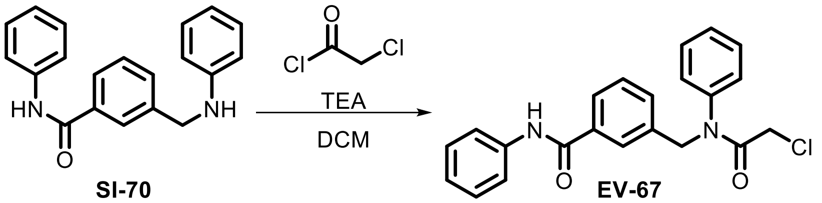

SI-70 EV-67

Following **General Procedure 1**, starting with **SI-70** (600 mg, 2.0 mmol, 1.0 eq), chloroacetyl chloride (2.0 eq), and TEA (5.0 eq), **EV-67** was obtained after prep-HPLC (HCl) as a yellow oil (40 mg, 5%).

**$^1$H NMR (400 MHz, DMSO-*d*$_6$) δ** 10.24 (s, 1H), 7.79 (d, *J* = 7.7 Hz, 1H), 7.74 – 7.60 (m, 3H), 7.43 (t, *J* = 7.6 Hz, 1H), 7.40 – 7.30 (m, 6H), 7.23 (d, *J* = 6.8 Hz, 2H), 7.09 (t, *J* = 7.4 Hz, 1H), 4.95 (s, 2H), 4.02 (s, 2H).

**HRMS ESI-TOF (*m/z*):** $[M+H]^+$ for $C_{22}H_{19}ClN_2O_2$ 379.1208, found 379.1210.

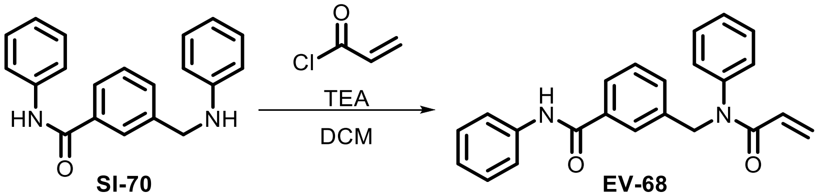

SI-70 EV-68

Following **General Procedure 2**, starting with **SI-70** (600 mg, 2.0 mmol, 1.0 eq), acryloyl chloride (2.0 eq), and TEA (3.0 eq), **EV-68** (121 mg, 17%) was obtained after prep-HPLC (basic).

**$^1$H NMR (400 MHz, DMSO-*d*$_6$) δ** 7.78 (d, *J* = 7.7, 1.5 Hz, 1H), 7.74 – 7.65 (m, 3H), 7.42 (t, *J* = 7.6 Hz, 1H), 7.39 – 7.27 (m, 6H), 7.16 – 7.12 (m, 2H), 7.09 (t, *J* = 7.3 Hz, 1H), 6.24 (dd, *J* = 16.8, 2.1 Hz, 1H), 6.09 – 5.95 (m, 1H), 5.62 (dd, *J* = 10.2, 2.1 Hz, 1H), 5.03 (s, 2H).

**HRMS ESI-TOF (*m/z*):** $[M+H]^+$ for $C_{23}H_{21}N_2O_2$ 357.1598, found 357.1598.

**Synthesis of SI-71 as a precursor for EV-69 and EV-70**

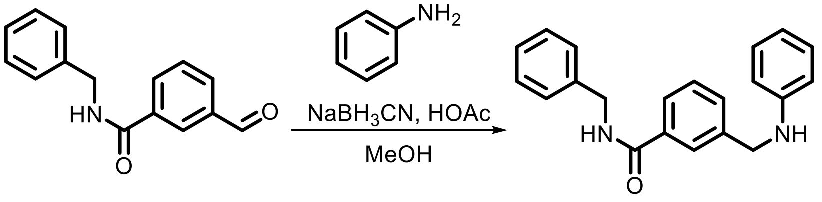

SI-71

Following **General Procedure 5**, starting with 3-formyl-*N*-benzylbenzamide (750 mg, 3.1 mmol, 1.0 eq) and aniline (1.1 eq), **SI-71** was obtained and used in the next step without purification.

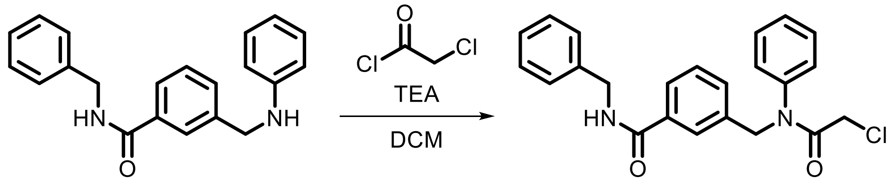

SI-71 EV-69

Following **General Procedure 1**, starting with **SI-71** (600 mg, 1.9 mmol, 1.0 eq), chloroacetyl chloride (2.0 eq), and TEA (5.0 eq), **EV-69** (94 mg, 12%) was obtained after prep-HPLC (basic).

**$^1$H NMR (400 MHz, DMSO-*d*$_6$) δ** 7.73 (d, *J* = 7.5 Hz, 1H), 7.68 (s, 1H), 7.42 – 7.31 (m, 4H), 7.32 – 7.24 (m, 3H), 7.22 (d, *J* = 6.9 Hz, 3H), 4.92 (s, 2H), 4.43 (s, 2H), 4.03 (s, 2H).

**HRMS ESI-TOF (*m/z*):** $[M+H]^+$ for $C_{23}H_{22}ClN_2O_2$ 393.1365, found 393.1365.

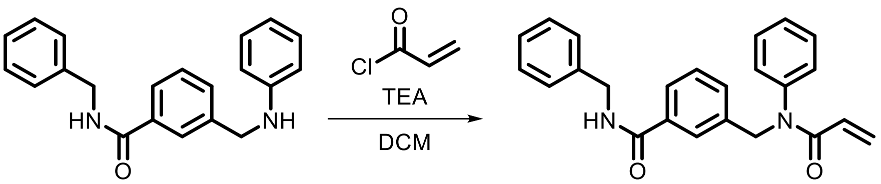

SI-71 EV-70

Following **General Procedure 2**, starting with **SI-71** (600 mg, 1.9 mmol, 1.0 eq), acryloyl chloride (2.0 eq), and TEA (5.0 eq), **EV-70** (150 mg, 20%) was obtained after prep-HPLC (basic).

**$^1$H NMR (400 MHz, DMSO-*d*$_6$) δ** 9.07 (t, *J* = 6.0 Hz, 0.3H), 8.38 (br s, 0.2H), 7.75 – 7.57 (m, 2H), 7.41 – 7.30 (m, 4H), 7.26 (br s, 5H), 7.20 (br s, 1H), 7.11 (d, *J* = 7.6 Hz, 2H), 6.33 – 6.18 (m, 1H), 6.08 – 5.92 (m, 1H), 5.60 (d, *J* = 10.3 Hz, 1H), 4.99 (s, 2H), 4.43 (s, 2H).

**HRMS ESI-TOF (*m/z*):** $[M+H]^+$ for $C_{24}H_{23}N_2O_2$ 371.1754, found 371.1757.

#### Synthesis of SI-72 as a Precursor for EV-71 and EV-72

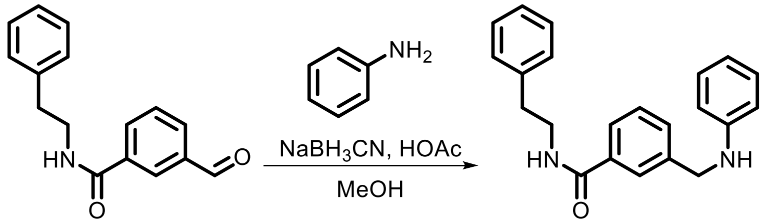

SI-72

Following **General Procedure 5**, starting with 3-formyl-*N*-phenethylbenzamide (750 mg, 3.0 mmol, 1.0 eq) and aniline (1.1 eq), **SI-72** was obtained as crude product and used in the next step without purification.

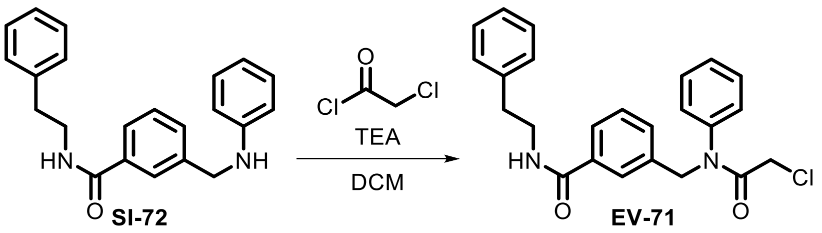

Following **General Procedure 1**, starting with **SI-72** (600 mg, 1.8 mmol, 1.0 eq), chloroacetyl chloride (2.0 eq), and TEA (5.0 eq), **EV-71** (203 mg, 25%) was obtained after prep-HPLC (basic).

**$^1$H NMR (400 MHz, DMSO-$d_6$) δ** 7.69 – 7.61 (m, 2H), 7.40 – 7.29 (m, 5H), 7.29 – 7.20 (m, 6H), 7.20 – 7.14 (m, 1H), 4.92 (s, 2H), 4.04 (s, 2H), 3.45 (t, *J* = 7.2 Hz, 2H), 2.81 (t, *J* = 7.6 Hz, 2H).

**HRMS ESI-TOF (*m/z*):** $[M+H]^+$ for $C_{24}H_{24}ClN_2O_2$ 407.1521, found 407.1521.

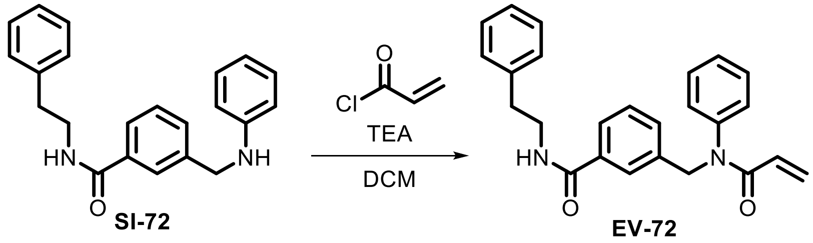

Following **General Procedure 2**, starting with **SI-72** (600 mg, 1.8 mmol, 1.0 eq), acryloyl chloride (2.0 eq), and TEA (5.0 eq), **EV-72** (135 mg, 18%) was obtained after prep-HPLC (basic).

**$^1$H NMR (400 MHz, DMSO-$d_6$) δ** 8.58 (t, *J* = 5.6 Hz, 1H), 7.68 – 7.59 (m, 2H), 7.37 – 7.27 (m, 5H), 7.27 – 7.18 (m, 4H), 7.15 (t, *J* = 7.0 Hz, 1H), 7.10 (d, *J* = 7.1 Hz, 2H), 6.24 (dd, *J* = 16.8, 2.2 Hz, 1H), 6.04 – 5.93 (m, 1H), 5.60 (dd, *J* = 10.1, 2.2 Hz, 1H), 4.98 (s, 2H), 3.50 – 3.37 (m, 2H), 2.80 (t, *J* = 7.4 Hz, 2H).

**HRMS ESI-TOF (*m/z*):** $[M+H]^+$ for $C_{25}H_{25}N_2O_2$ 385.1911, found 385.1910.

**Synthesis of SI-73 as a precursor for EV-73 and EV-74**

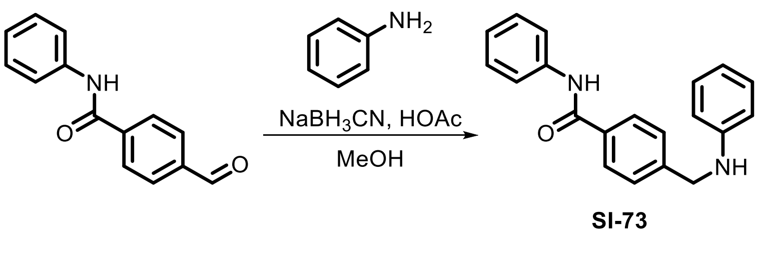

Following **General Procedure 5**, starting with 4-formyl-*N*-phenylbenzamide (750 mg, 3.3 mmol, 1.0 eq) and aniline (1.1 eq), **SI-73** was obtained as crude product and used without purification.

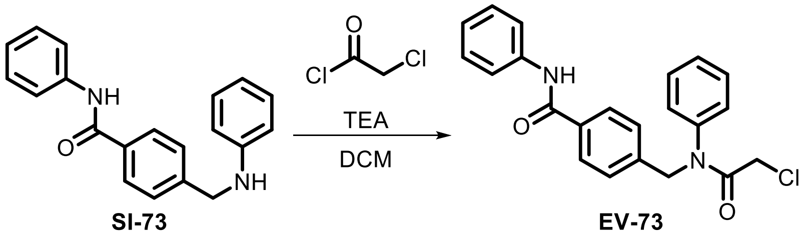

Following **General Procedure 1**, starting with **SI-73** (600 mg, 2.0 mmol, 1.0 eq), chloroacetyl chloride (2.0 eq), and TEA (5.0 eq), **EV-73** (136 mg, 18%) was obtained after prep-HPLC (basic).

**$^1$H NMR (400 MHz, DMSO-$d_6$) δ** 10.21 (s, 1H), 7.87 (d, *J* = 8.1 Hz, 2H), 7.75 (d, *J* = 8.0 Hz, 2H), 7.44 – 7.28 (m, 9H), 7.09 (t, *J* = 7.4 Hz, 1H), 4.98 (s, 2H), 4.11 (s, 2H).

**HRMS ESI-TOF (*m/z*):** $[M+H]^+$ for $C_{22}H_{20}ClN_2O_2$ 379.1208, found 379.1204.

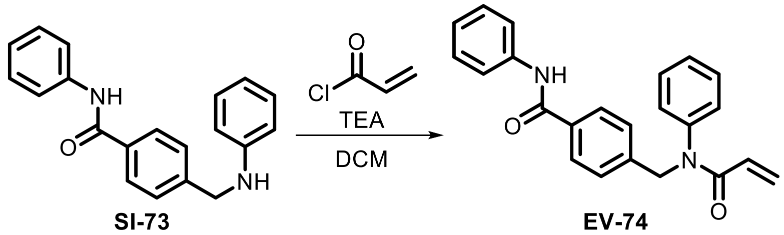

Following **General Procedure 2**, starting with **SI-73** (600 mg, 2.0 mmol, 1.0 eq), acryloyl chloride (2.0 eq), and TEA (5.0 eq), **EV-74** (72 mg, 10%) was obtained after prep-HPLC (basic).

**$^1$H NMR (400 MHz, DMSO-$d_6$) δ** 10.21 (s, 1H), 7.86 (d, *J* = 8.0 Hz, 2H), 7.75 (d, *J* = 8.0 Hz, 2H), 7.43 – 7.29 (m, 7H), 7.19 (d, *J* = 7.6 Hz, 2H), 7.09 (t, *J* = 7.4 Hz, 1H), 6.27 (dd, *J* = 16.8, 2.4 Hz, 1H), 6.15 – 6.00 (m, 1H), 5.65 (dd, *J* = 10.2, 2.4 Hz, 1H), 5.05 (s, 2H).

**HRMS ESI-TOF (*m/z*):** $[M+H]^+$ for $C_{23}H_{21}N_2O_2$ 357.1598, found 357.1599.

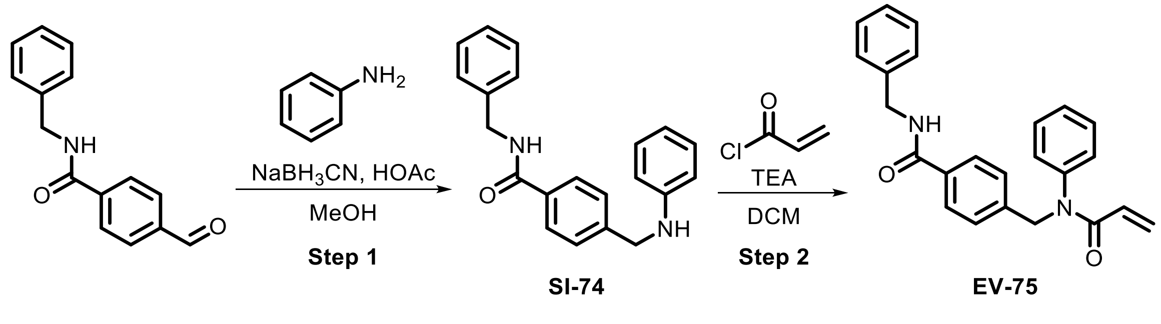

**Step 1:** Following **General Procedure 5**, starting with 4-formyl-*N*-benzylbenzamide (750 mg, 3.1 mmol, 1.0 eq) and aniline (1.1 eq), **SI-74** (600 mg, crude) was obtained and used in the next step without purification.

**Step 2:** Following **General Procedure 2**, starting with **SI-74** (600 mg, 1.9 mmol, 1.0 eq), acryloyl chloride (2.0 eq), and TEA (5.0 eq), **EV-75** (233 mg, 31%) was obtained after prep-HPLC (basic).

**¹H NMR (400 MHz, DMSO-*d*$_6$) δ** 9.04 (t, *J* = 6.1 Hz, 1H), 7.77 (d, *J* = 8.0 Hz, 2H), 7.36 (t, *J* = 7.5 Hz, 2H), 7.32 – 7.24 (m, 7H), 7.23 – 7.18 (m, 1H), 7.13 (d, *J* = 7.2 Hz, 2H), 6.23 (dd, *J* = 16.8, 2.2 Hz, 1H), 6.13 – 5.91 (m, 1H), 5.63 (dd, *J* = 10.2, 1.6 Hz, 1H), 4.99 (s, 2H), 4.43 (d, *J* = 6.0 Hz, 2H).

**HRMS ESI-TOF (*m/z*):** $[M+H]^+$ for $C_{24}H_{23}N_2O_2$ 371.1754, found 371.1754.

**Synthesis of SI-75 as a precursor for EV-76 and EV-77**

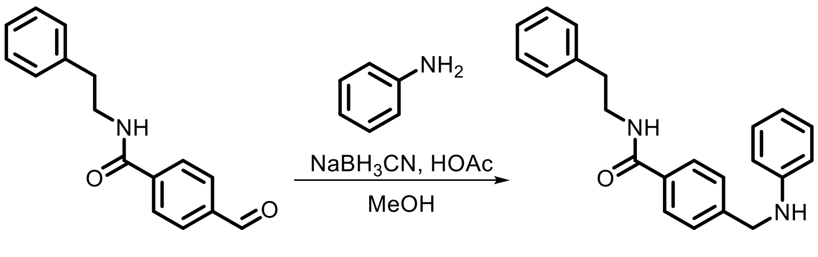

**SI-75**

Following **General Procedure 5**, starting with 4-formyl-*N*-phenethylbenzamide (760 mg, 3.0 mmol, 1.0 eq) and aniline (1.1 eq), **SI-75** was obtained as crude product and used in the next step without purification.

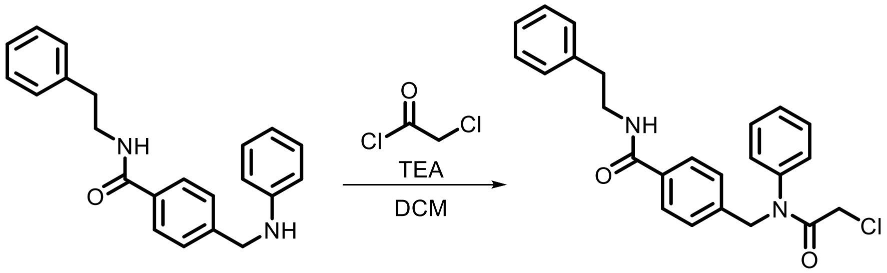

**SI-75** **EV-76**

Following **General Procedure 1**, starting with **SI-75** (600 mg, 1.8 mmol, 1.0 eq), chloroacetyl chloride (2.0 eq), and TEA (5.0 eq), **EV-76** (55 mg, 7%) was obtained after prep-HPLC (basic).

**¹H NMR (400 MHz, DMSO-*d*$_6$) δ** 8.55 (t, *J* = 5.6 Hz, 0.1H), 7.69 (d, *J* = 8.4 Hz, 2H), 7.41 – 7.32 (m, 3H), 7.29 – 7.21 (m, 7H), 7.21 – 7.14 (m, 2H), 4.91 (s, 2H), 4.04 (s, 2H), 3.44 (t, *J* = 7.3 Hz, 2H), 2.80 (t, *J* = 7.5 Hz, 2H).

**HRMS ESI-TOF (*m/z*):** $[M+H]^+$ for $C_{24}H_{24}ClN_2O_2$ 407.1521, found 407.1521.

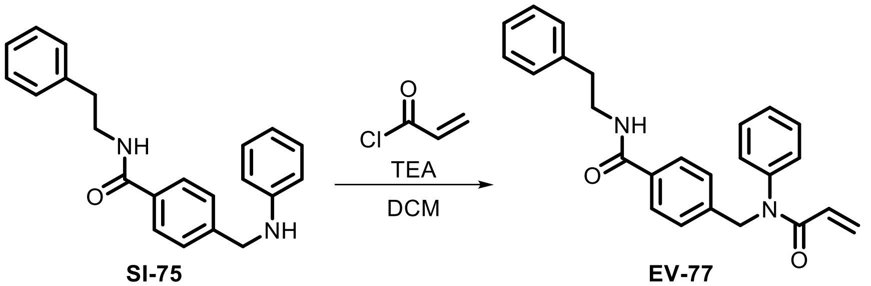

**SI-75** **EV-77**

Following **General Procedure 1**, starting with **SI-75** (600 mg, 1.8 mmol, 1.0 eq), chloroacetyl chloride (2.0 eq), and TEA (5.0 eq), **EV-77** (203 mg, 26%) was obtained after prep-HPLC (basic).

**$^{1}$H NMR (400 MHz, DMSO-$d_6$) δ** 8.55 (t, *J* = 5.6 Hz, 1H), 7.69 (d, *J* = 8.1 Hz, 2H), 7.36 (d, *J* = 7.4 Hz, 2H), 7.33 – 7.26 (m, 2H), 7.27 – 7.16 (m, 6H), 7.16 – 7.09 (m, 2H), 6.23 (dd, *J* = 16.8, 2.2 Hz, 1H), 6.07 – 5.95 (m, 1H), 5.62 (dd, *J* = 10.2, 2.2 Hz, 1H), 4.98 (s, 2H), 3.45 (q, *J* = 6.5 Hz, 2H), 2.80 (t, *J* = 7.5 Hz, 2H).

**HRMS ESI-TOF (*m/z*):** $[M+H]^+$ for $C_{25}H_{25}N_2O_2$ 385.1911, found 385.1913.

**Synthesis of SI-77 as a precursor for EV-78 – EV-81**

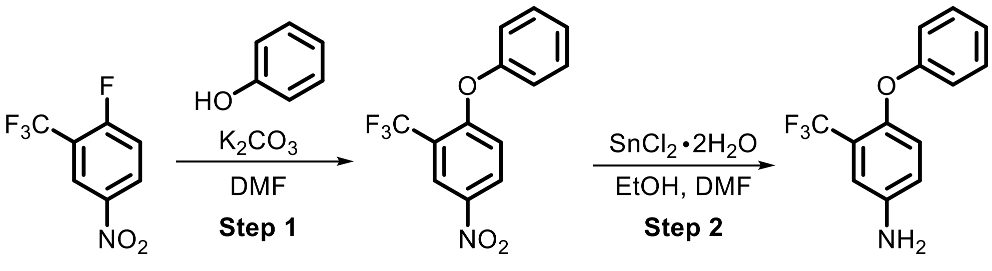

**SI-76** **SI-77**

**Step 1:** $K_2CO_3$ (6.6 g, 48 mmol, 1.0 eq) was added in one portion to a solution of phenol (4.5 g, 48 mmol, 1.0 eq) in DMF (50 mL) at 25 °C and the mixture was stirred for 1 h. Fluoro-4-nitro-2-(trifluoromethyl)benzene (6.6 mL, 48 mmol, 1.0 eq) was then added and the reaction was heated to 80 °C and stirred for 16 h. Upon completion, the mixture was poured into $H_2O$ (300 mL) and extracted with EtOAc (100 mL x 3). The combined organic layers were washed with brine (100 mL x 3), dried over anhydrous $Na_2SO_4$ and concentrated under reduced pressure to give **SI-76** (13 g, 96%) as a yellow solid.

**Step 2:** DMF (35 µL, 0.46 mmol, 0.01 eq) and $SnCl_2 \cdot 2H_2O$ (41.4 g, 184 mmol, 4.0 eq) were added sequentially to a solution of **SI-76** (13.0 g, 46 mmol, 1.0 eq) in EtOH (130 mL) at 25 °C. The reaction mixture was heated to 80 °C and stirred for 2 h. The reaction was cooled and concentrated under reduced pressure and the resulting residue was diluted with $H_2O$ (300 mL). The suspension was filtered and the filtrate was extracted with DCM (100 mL x 5). The combined layers were dried over anhydrous $Na_2SO_4$, and concentrated under reduced pressure to provide **SI-77** (11.0 g, 67%) as a yellow oil.

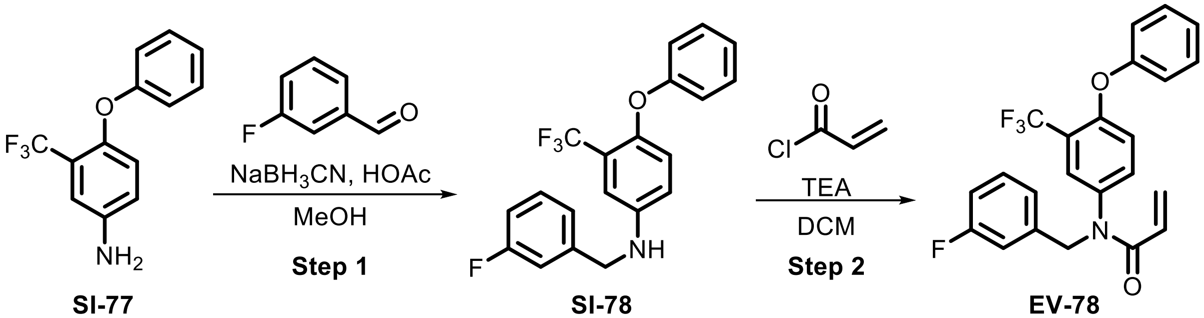

**SI-77** **SI-78** **EV-78**

**Step 1:** Following **General Procedure 5**, starting with **SI-77** (200 mg, 0.79 mmol, 1.0 eq) and 3-fluorobenzaldehyde (1.1 eq), **SI-78** was obtained as crude product and used in the next step without purification.

**Step 2:** Following **General Procedure 2**, starting with **SI-78** (140 mg, 0.39 mmol, 1.0 eq), acryloyl chloride (2.0 eq), and TEA (5.0 eq), **EV-78** was obtained after prep-HPLC (basic) as a colorless oil (30 mg, 18%).

**$^{1}$H NMR (400 MHz, $CDCl_3$) δ** 7.42(dd, *J* = 8.6, 7.4 Hz, 2H), 7.36 (d, *J* = 2.6 Hz, 1H), 7.31 – 7.19 (m, 2H), 7.12 – 7.06 (m, 2H), 7.06 – 6.94 (m, 4H), 6.84 (d, *J* = 8.8 Hz, 1H), 6.49 (dd, *J* = 16.7, 1.9 Hz, 1H), 6.05 (dd, *J* = 16.7, 10.3 Hz, 1H), 5.66 (dd, *J* = 10.3, 1.9 Hz, 1H), 4.97 (s, 2H).

**HRMS ESI-TOF (*m/z*):** $[M+H]^+$ for $C_{23}H_{18}F_3NO_2$ 416.1268, found 416.1268.

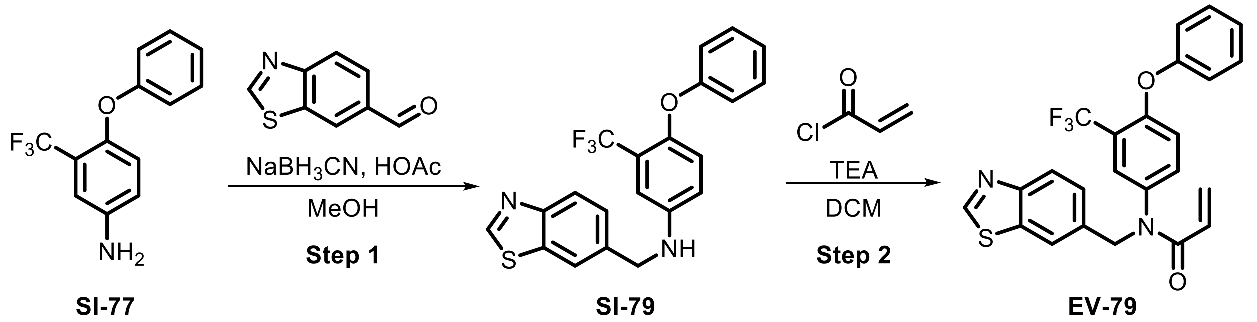

**Step 1:** Following **General Procedure 5**, starting with **SI-77** (300 mg, 1.2 mmol, 1.0 eq) and 1,3-benzothiazole-6-carbaldehyde (1.0 eq), **SI-79** was obtained as crude product and used without purification.

**Step 2:** Following **General Procedure 2**, starting with **SI-79** (125 mg, 0.31 mmol, 1.0 eq), acryoyl chloride (2.0 eq), and TEA (5.0 eq), **EV-79** was obtained after prep-HPLC (basic) as a light yellow oil (86 mg, 57%).

**$^1$H NMR (400 MHz, $CDCl_3$) δ** 8.98 (s, 1H), 8.05 (d, *J* = 8.4 Hz, 1H), 7.88 (d, *J* = 1.4 Hz, 1H), 7.44 – 7.32 (m, 4H), 7.20 (t, *J* = 7.4 Hz, 1H), 7.05 (dd, *J* = 8.6, 1.1 Hz, 2H), 6.96 (d, *J* = 8.6 Hz, 1H), 6.78 (d, *J* = 8.8 Hz, 1H), 6.49 (dd, *J* = 16.7, 1.9 Hz, 1H), 6.03 (dd, *J* = 16.5, 10.6 Hz, 1H), 5.65 (dd, *J* = 10.3, 1.9 Hz, 1H), 5.10 (s, 2H).

**HRMS ESI-TOF (*m/z*):** $[M+H]^+$ for $C_{24}H_{18}F_3N_2O_2S$ 455.1036, found 455.1037.

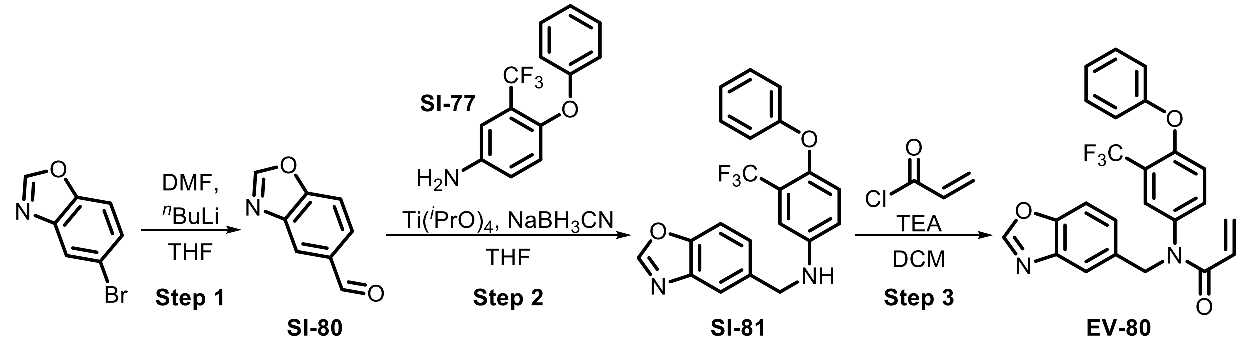

**Step 1:** $^n$BuLi (2.5 M, 4.0 mL, 2.0 *eq*) was added to a solution of 5-bromobenzoxazole (1.0 g, 5.1 mmol, 1.0 *eq*) in THF (8 mL) at -78 °C under a nitrogen atmosphere. After stirring for 1 h at -78 °C, DMF (777 µL, 10.1 mmol, 2.0 *eq*) was added and the reaction mixture was stirred at -78 °C for an additional 2 h. The reaction mixture was warmed to 25 °C, quenched by the addition of aq. $NH_4Cl$ (10 mL) and brine (20 mL) and then extracted with DCM (30 ml x 3). The combined organic layers were dried over anhydrous $Na_2SO_4$, filtered and concentrated. The resulting residue was purified by flash chromatography to provide **SI-80** (100 mg, 11%) as a brown solid.

**Step 2:** $Ti(O^iPr)_4$ (201 µL, 0.68 mmol, 1.0 *eq*) was added to a solution of **SI-80** (100 mg, 0.68 mmol, 1.0 *eq*) and **SI-77** (207 mg, 0.82 mmol, 1.2 *eq*) in THF (2 mL) and stirred at 25 °C for 1 h. The reaction was cooled to 0 °C then $NaBH_3CN$ (85 mg, 1.4 mmol, 2.0 *eq*) and the reaction was stirred at 0 °C for 2 h. The reaction mixture was diluted with $H_2O$ (10 mL) and extracted with DCM (20 mL x 3). The combined organic layers were washed with brine (10 mL), dried over anhydrous $Na_2SO_4$ and concentrated to provide **SI-81** (200 mg, crude) as a yellow oil which was used in the next step without purification.

**Step 3:** Following **General Procedure 2**, starting with **SI-81** (90 mg, 0.23 mmol, 1.0 *eq*), acryloyl chloride (1.5 *eq*), and TEA (5.0 *eq*), **EV-80** was obtained after prep-HPLC (basic) as a yellow solid (24 mg, 22%).

**$^1$H NMR (400 MHz, $CDCl_3$) δ** 8.08 (s, 1H), 7.59 (d, *J* = 1.7 Hz, 1H), 7.52 (d, *J* = 8.4 Hz, 1H), 7.43 – 7.31 (m, 4H), 7.23 – 7.17 (m, 1H), 7.07 – 7.02 (m, 2H), 6.96 (dd, *J* = 9.0, 2.6 Hz, 1H), 6.77 (d, *J* = 8.7 Hz, 1H), 6.48 (dd, *J* = 16.7, 1.9 Hz, 1H), 6.02 (dd, *J* = 16.7, 10.2 Hz, 1H), 5.64 (dd, *J* = 10.3, 1.9 Hz, 1H), 5.08 (s, 2H).

**HRMS ESI-TOF (*m/z*):** $[M+H]^+$ for $C_{24}H_{18}F_3N_2O_3$ 439.1264, found 439.1269.

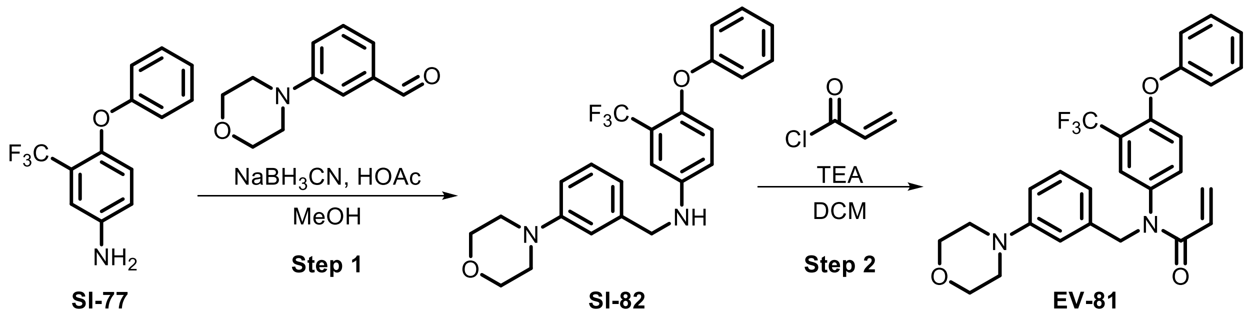

**Step 1:** Following **General Procedure 5**, **SI-77** (300 mg, 1.2 mmol, 1.0 eq) and 3-morpholinobenzaldehyde (1.0 eq), **SI-82** was obtained as a crude product and used without purification.

**Step 2:** Following **General Procedure 2**, starting with **SI-82** (125 mg, 0.29 mmol, 1.0 eq), acryloyl chloride (2.0 eq), and TEA (5.0 eq), **EV-81** was obtained after prep-HPLC (HCl) as a light yellow oil (118 mg, 81%).

**$^1$H NMR (400 MHz, $CDCl_3$) δ** 7.47 – 7.33 (m, 2H), 7.31 (br s, 1H), 7.24 – 7.14 (m, 2H), 7.09 – 6.98 (m, 3H), 6.83 – 6.76 (m, 3H), 6.69 (d, *J* = 7.5 Hz, 1H), 6.45 (dd, *J* = 16.7, 1.9 Hz, 1H), 6.10 – 5.96 (m, 1H), 5.62 (d, *J* = 10.3 Hz, 1H), 4.90 (s, 2H), 3.88 – 3.80 (m, 4H), 3.17 – 3.06 (m, 4H).

**HRMS ESI-TOF (*m/z*):** $[M+H]^+$ for $C_{27}H_{26}F_3N_2O_3$ 483.1890, found 483.1901.

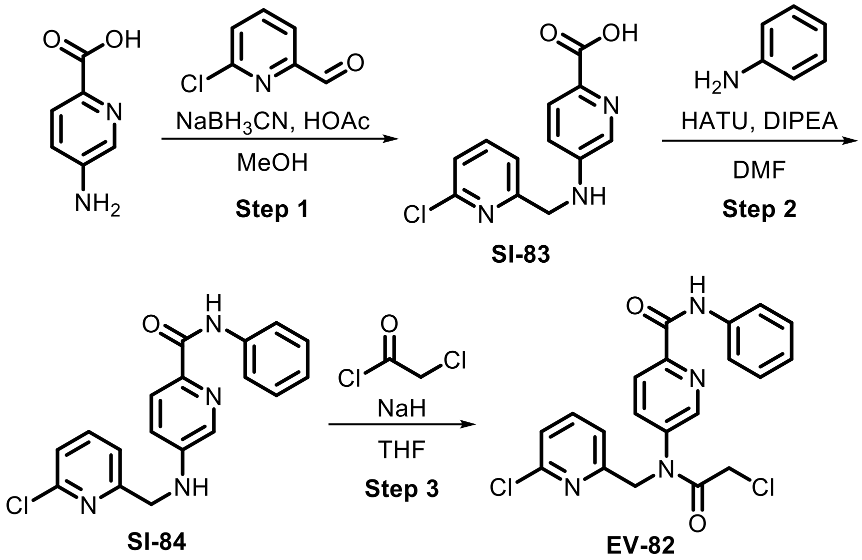

**Step 1:** Following **General Procedure 5**, starting with 6-chloropyridine-2-carbaldehyde (150 mg, 1.1 mmol, 1.0 eq) and 5-aminopicolinic acid (1.0 eq), **SI-83** was obtained as a crude product and used in the next step without purification.

**Step 2:** Following **General Procedure 3**, starting with **SI-83** (1.0 eq) and aniline (1.0 eq), **SI-84** (150 mg, crude) was obtained as a crude product and used in the next step without purification.

**Step 3:** NaH (21 mg, 0.53 mmol, 60% dispersion in mineral oil, 2.0 eq) was added to a solution of **SI-84** (150 mg, 0.27 mmol, 1.0 eq) in anhydrous THF (2 mL) at 0 °C and then stirred for 1 h. Chloroacetyl chloride (127 µL, 1.6 mmol, 6.0 eq) was then added at 0 °C. The mixture was warmed to 25 °C and stirred for 15 h. The mixture was diluted with MeCN (3 mL) and the product was purified by prep-HPLC (HCl). The eluent was evaporated to remove organic solvents followed by lyophilization to provide **EV-82** (26 mg, 21%) as a yellow oil.

**$^1$H NMR (400 MHz, DMSO-$d_6$) δ** 10.64 (s, 1H), 8.81 (br zs, 1H), 8.23 – 8.10 (m, 2H), 7.91 – 7.81 (m, 3H), 7.43 (dd, $J$ = 15.4, 7.7 Hz, 2H), 7.35 (t, $J$ = 7.9 Hz, 2H), 7.15 – 7.07 (m, 1H), 5.06 (s, 2H), 4.46 – 4.24 (m, 2H).

**HRMS ESI-TOF (*m/z*):** $[M+H]^+$ for $C_{20}H_{17}Cl_2N_4O_2$ 415.0723, found 415.0725.

**Synthesis of SI-85 as a precursor for EV-83 and EV-84**

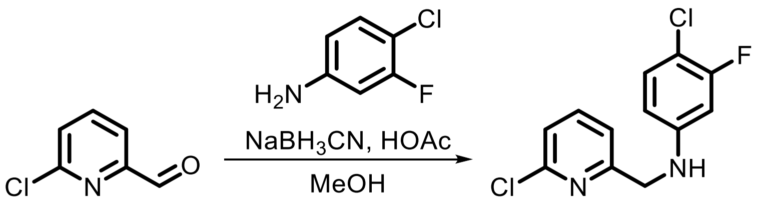

**SI-85**

Following **General Procedure 5**, starting with 6-chloropyridine-2-carbaldehyde (150 mg, 1.1 mmol, 1.0 eq) and 4-chloro-3-fluoro-aniline (1.0 eq), **SI-85** was obtained as a crude product and used in the next step without purification.

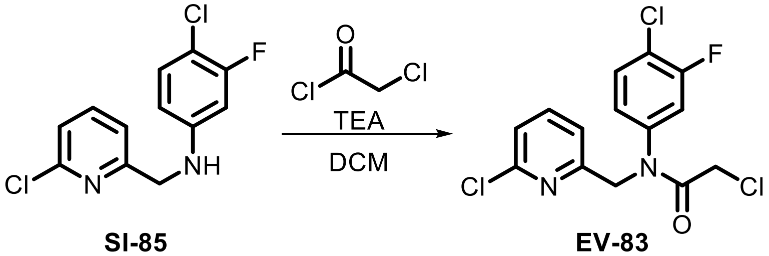

**SI-85** **EV-83**

Following **General Procedure 1**, starting with **SI-85** (140 mg, 0.52 mmol, 1.0 eq), chloroacetyl chloride (2.0 eq), and TEA (5.0 eq), **EV-83** was obtained after prep. HPLC (basic) as a colorless oil (25 mg, 12%).

**$^1$H NMR (400 MHz, DMSO-$d_6$) δ** 7.83 (t, $J$ = 7.8 Hz, 1H), 7.71 – 7.57 (m, 2H), 7.40 (t, $J$ = 7.1 Hz, 2H), 7.32 (br s, 1H), 4.94 (br s, 2H), 4.25 (br s, 2H).

**HRMS ESI-TOF (*m/z*):** $[M+H]^+$ for $C_{14}H_{11}Cl_3FN_2O$ 346.9916, found 346.9916.

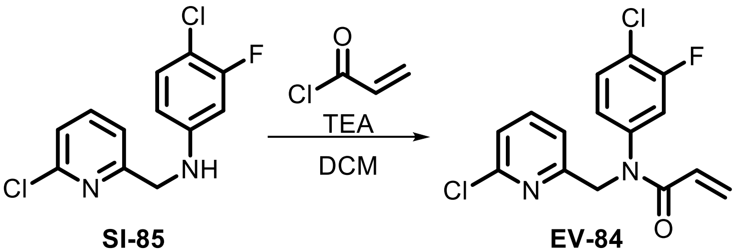

Following **General Procedure 2**, starting with **SI-85** (140 mg, 0.52 mmol, 1.0 eq), acryloyl chloride (2.0 eq), and TEA (5.0 eq), **EV-84** was obtained after prep. HPLC (basic) as a colorless oil (50 mg, 29%).

**$^1$H NMR (400 MHz, $CDCl_3$) δ** 7.62 (t, *J* = 7.7 Hz, 1H), 7.40 (t, *J* = 8.3 Hz, 1H), 7.31 (d, *J* = 6.8 Hz, 1H), 7.21 (d, *J* = 7.1 Hz, 1H), 7.14 (dd, *J* = 9.5, 2.4 Hz, 1H), 7.02 (ddd, *J* = 8.5, 2.4, 1.2 Hz, 1H), 6.42 (dd, *J* = 16.7, 1.8 Hz, 1H), 6.11 (dd, *J* = 16.8, 10.3 Hz, 1H), 5.63 (dd, *J* = 10.3, 1.9 Hz, 1H), 4.99 (s, 2H).

**HRMS ESI-TOF (*m/z*):** $[M+H]^+$ for $C_{15}H_{12}Cl_2FN_2O$ 325.0305, found 325.0304.

**Synthesis of SI-86 as a precursor for EV-85 and EV-86**

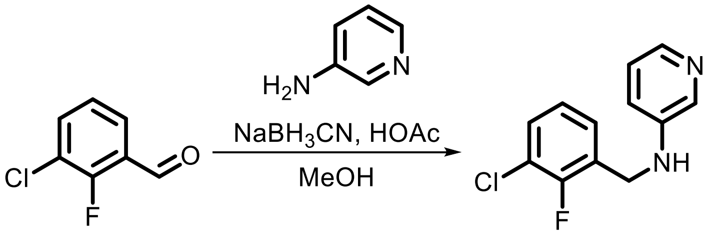

Following **General Procedure 5**, starting with 3-chloro-2-fluorobenzaldehyde (150 mg, 1.1 mmol, 1.0 eq) and 3-aminopyridine (1.0 eq), **SI-86** was obtained as a crude product and used without purification.

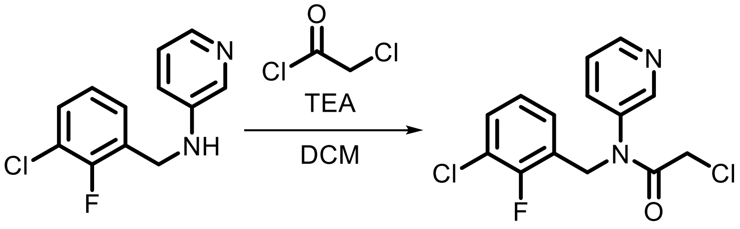

Following **General Procedure 1**, starting with **SI-86** (350 mg, 1.5 mmol, 1.0 eq), chloroacetyl chloride (1.5 eq), and TEA (4.0 eq), **EV-85** was obtained after prep. HPLC (FA) as a white solid (135 mg, 29%).

**$^1$H NMR (400 MHz, $CDCl_3$) δ** 8.86 (d, *J* = 53.0 Hz, 2H), 8.20 (s, 1H), 7.91 (s, 1H), 7.38 (t, *J* = 7.4 Hz, 1H), 7.29 (s, 1H), 7.11 (t, *J* = 7.3 Hz, 1H), 5.12 (s, 2H), 4.08 (s, 2H).

**HRMS ESI-TOF (*m/z*):** $[M+H]^+$ for $C_{14}H_{12}Cl_2FN_2O$ 313.0305, found 313.0308.

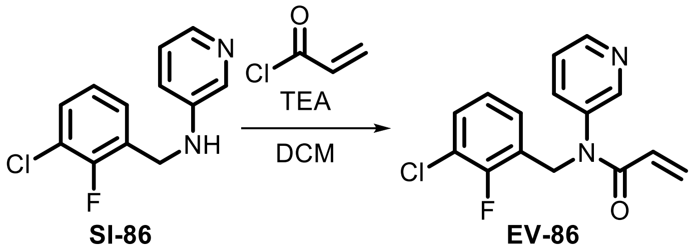

Following **General Procedure 1**, starting with **SI-86** (200 mg, 1.5 mmol, 1.0 eq), chloroacetyl chloride (1.5 eq), and TEA (3.0 eq), **EV-86** was obtained after prep. HPLC (FA) as a white solid (17 mg, 7%).

**$^1$H NMR (400 MHz, DMSO-$d_6$) δ** 8.79 (s, 1H), 8.70 (dd, *J* = 5.3, 1.4 Hz, 1H), 8.16 (d, *J* = 8.2 Hz, 1H), 7.80 (dd, *J* = 8.3, 5.2 Hz, 1H), 7.48 (td, *J* = 7.6, 1.7 Hz, 1H), 7.34 (td, *J* = 7.3, 6.8, 1.6 Hz, 1H), 7.17 (td, *J* = 7.9, 1.1 Hz, 1H), 6.28 (dd, *J* = 16.5, 2.5 Hz, 2H), 5.80 – 5.61 (m, 1H), 5.13 (s, 2H).

**HRMS ESI-TOF (*m/z*):** $[M+H]^+$ for $C_{15}H_{13}ClFN_2O$ 291.0695, found 391.0697.

**Synthesis of SI-89 as a precursor for EV-87 and EV-88**

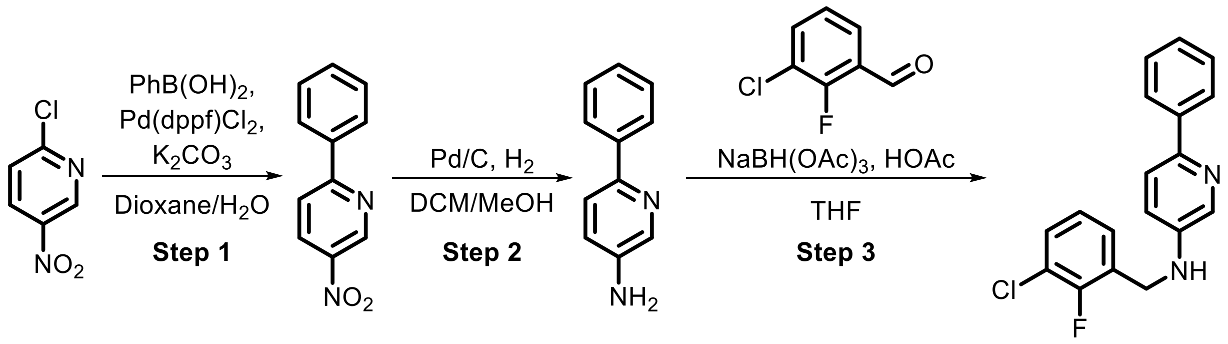

**Step 1:** Under an atmosphere of nitrogen, $Pd(dppf)Cl_2$ (462 mg, 0.63 mmol, 0.05 *eq*) and $K_2CO_3$ (3.5 g, 25 mmol, 2.0 *eq*) were added to a solution of 2-chloro-5-nitropyridine (2.0 g, 12.6 mmol, 1.0 *eq*) and phenylboronic acid (1.7 g, 14 mmol, 1.1 *eq*) in dioxane (24 mL) and $H_2O$ (8 mL). The mixture was stirred at 100 °C for 6 h. The reaction mixture was diluted with $H_2O$ (20 mL) and extracted with DCM (30 mL x 3). The combined organic layers were dried over anhydrous $Na_2SO_4$, filtered and concentrated. The resulting residue was purified by flash chromatography to give **SI-87** (2.4 g, 93%) as a yellow solid.

**Step 2:** A mixture of **SI-87** (1.0 g, 5.0 mmol, 1.0 *eq*), Pd/C (20 mg, 5.0 mmol, 1.0 *eq*) in DCM (3 mL) and MeOH (3 mL) was evacuated and backfilled with $H_2$ (3x). The reaction mixture was stirred at 25 °C for 6 h under $H_2$ atmosphere (15 psi). The reaction mixture was filtered and the filtrate was concentrated to provide **SI-88** (750 mg, 85%) as a light yellow solid.

**Step 3:** Following **General Procedure 5**, starting with **SI-88** (650 mg, 3.8 mmol, 1.0 *eq*) and 3-chloro-2-fluoro-benzaldehyde (606 mg, 3.8 mmol, 1.0 *eq*), **SI-89** (1.2 g, 71%) was obtained and used in the next step without purification.

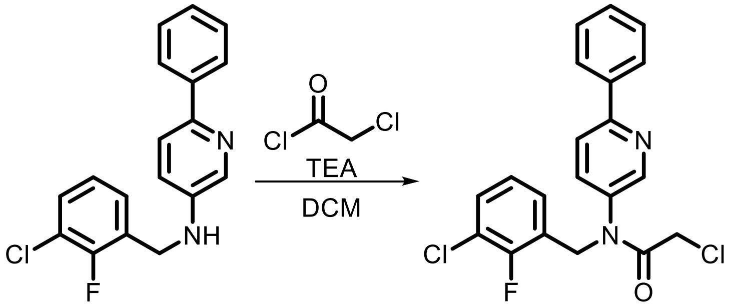

**SI-89** **EV-87**

Following **General Procedure 1**, starting with **SI-89** (150 mg, 0.48 mmol, 1.0 *eq*), chloroacetyl chloride (1.5 *eq*), and TEA (5.0 *eq*), **EV-87** was obtained after prep-HPLC (HCl) as a yellow solid (15 mg, 8%).

**$^1$H NMR (400 MHz, $CDCl_3$) δ** 8.46 (s, 1H), 8.00 (d, *J* = 6.4 Hz, 2H), 7.79 (d, *J* = 8.2 Hz, 1H), 7.60 – 7.42 (m, 4H), 7.41 – 7.30 (m, 2H), 7.07 (t, *J* = 7.8 Hz, 1H), 5.03 (s, 2H), 3.91 (s, 2H).

**HRMS ESI-TOF (*m/z*):** $[M+H]^+$ for $C_{20}H_{16}Cl_2FN_2O$ 389.0618, found 389.0623.

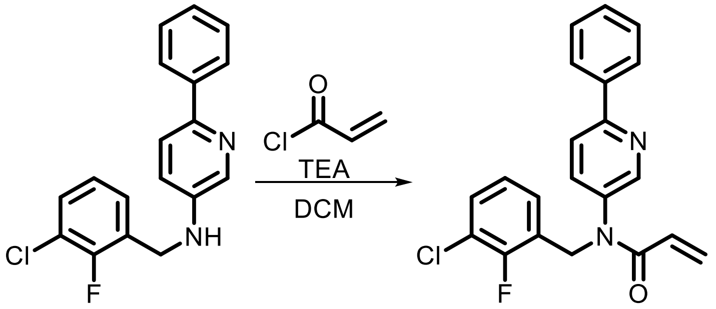

**SI-89** **EV-88**

Following **General Procedure 2**, starting with **SI-89** (600 mg, 1.9 mmol, 1.0 *eq*), acryloyl chloride (1.5 *eq*), and TEA (5.0 *eq*), **EV-88** was obtained after prep-HPLC (HCl) as a yellow solid (212 mg, 29%).

**$^1$H NMR (400 MHz, $CDCl_3$) δ** 8.65 (s, 1H), 8.12 (dd, *J* = 6.6, 2.9 Hz, 2H), 8.04 – 7.87 (m, 2H), 7.64 – 7.52 (m, 3H), 7.43 – 7.31 (m, 2H), 7.09 (t, *J* = 7.8 Hz, 1H), 6.52 (d, *J* = 16.6 Hz, 1H), 6.12 (dd, *J* = 16.5, 10.2 Hz, 1H), 5.77 (d, *J* = 10.1 Hz, 1H), 5.13 (s, 2H).

**HRMS ESI-TOF (*m/z*):** $[M+H]^+$ for $C_{21}H_{17}ClFN_2O$ 367.1008, found 367.1013.

**Synthesis of SI-90 as a precursor for EV-89 and EV-90**

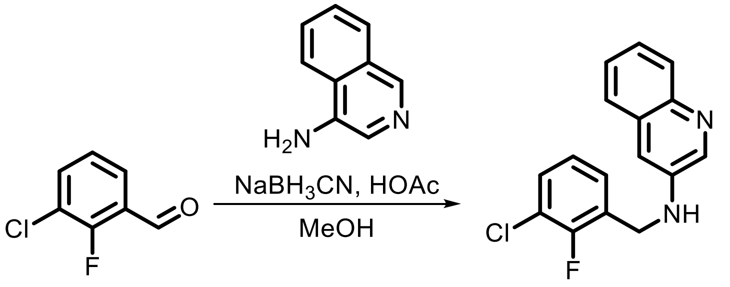

**SI-90**

Following **General Procedure 5**, starting with 3-chloro-2-fluorobenzaldehyde (150 mg, 1.1 mmol, 1.0 eq) and 3-aminoquinoline (1.0 eq), **SI-90** was obtained as a crude product and used without purification.

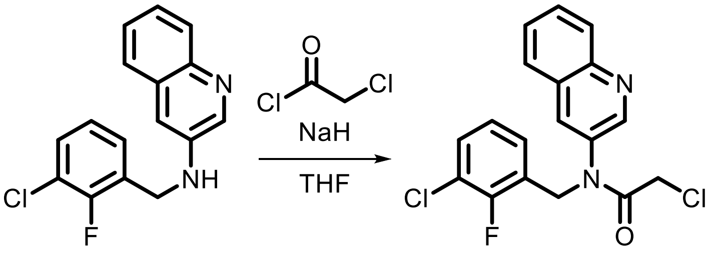

**SI-90** **EV-89**

NaH (17 mg, 0.70 mmol, 2.0 eq) was added to a solution of **SI-90** (100 mg, 0.35 mmol, 1.0 eq) in THF (2 mL) at 0 °C. The reaction was warmed to 25 °C and stirred for 0.5 h. The reaction was cooled to 0 °C and chloroacetyl cloride (28 µL, 0.35 mmol, 1.0 eq) was added. The reaction was warmed to 25 °C and stirred for an additional 1.5 h. The reaction was quenched with $H_2O$ (3 mL) and extracted with DCM (1 mL x 3). The combined organic layers were washed with brine (2 mL x 3), dried over anhydrous $Na_2SO_4$ and concentrated under reduced pressure. The resulting residue was purified by prep-HPLC (FA) to provide **EV-89** (17 mg, 13%) as a yellow oil.

**$^1$H NMR (400 MHz, $CDCl_3$) δ** 9.02 (s, 1H), 8.46 (d, *J* = 31.6 Hz, 2H), 7.96 (d, *J* = 8.0 Hz, 2H), 7.77 (br s, 1H), 7.43 – 7.27 (m, 2H), 7.05 (t, *J* = 7.5 Hz, 1H), 5.13 (s, 2H), 3.97 (s, 2H).

**HRMS ESI-TOF (*m/z*):** $[M+H]^+$ for $C_{18}H_{14}Cl_2FN_2O$ 363.0462, found 363.0462.

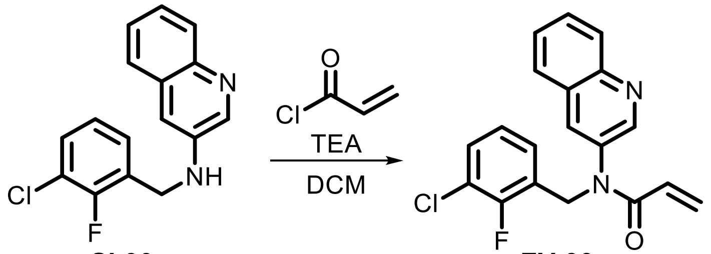

**SI-90** **EV-90**

Following **General Procedure 2**, starting with **SI-90** (150 mg, 0.52 mmol, 1.0 eq), acryloyl chloride (1.0 eq), and TEA (3.0 eq), **EV-90** was obtained after prep-HPLC (basic) as a yellow oil (21 mg, 12%).

**$^1$H NMR (400 MHz, $CDCl_3$) δ** 8.62 (d, *J* = 2.5 Hz, 1H), 8.12 (d, *J* = 8.4 Hz, 1H), 7.88 (d, *J* = 2.5 Hz, 1H), 7.81 – 7.71 (m, 2H), 7.60 (t, *J* = 7.4 Hz, 1H), 7.37 (t, *J* = 6.6 Hz, 1H), 7.33 – 7.24 (m, 1H), 7.05 (t, *J* = 7.9 Hz, 1H), 6.49 (dd, *J* = 16.7, 1.8 Hz, 1H), 6.02 (dd, *J* = 16.8, 10.4 Hz, 1H), 5.62 (dd, *J* = 10.3, 1.8 Hz, 1H), 5.17 (s, 2H).

**HRMS ESI-TOF (*m/z*):** $[M+H]^+$ for $C_{19}H_{15}ClFN_2O$ 341.0852, found 341.0854.

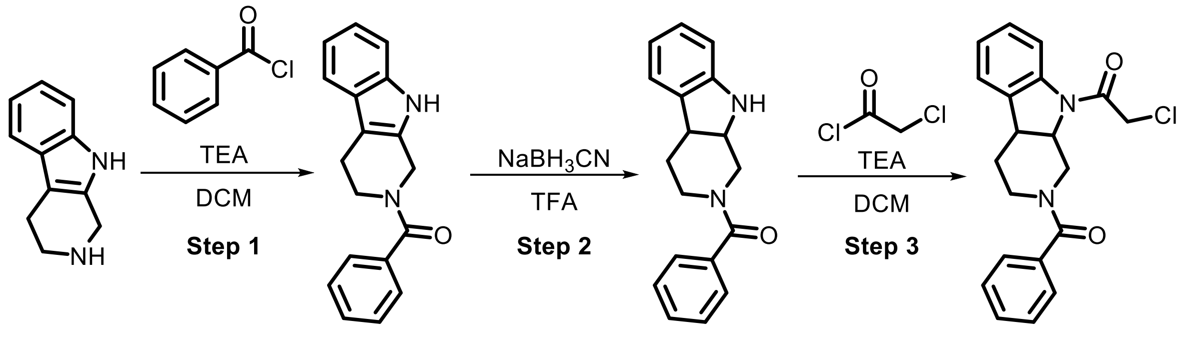

**SI-91** **SI-92** **EV-91**

**Step 1:** Following **General Procedure 4**, starting with 2,3,4,9-tetrahydro-1H-pyrido[3,4-*b*]indole (100 mg, 0.58 mmol, 1.0 eq) and benzoyl chloride (1.0 eq), **SI-91** was obtained as crude product and used in the next step without purification.

**Step 2:** $NaBH_3CN$ (73 mg, 1.2 mmol, 2.0 eq) was added to a solution of **SI-91** (160 mg, 0.58 mmol, 1.0 eq) in TFA (2 mL) at 0 °C. The reaction mixture was warmed to 20 °C and stirred for 2 h. The reaction mixture was concentrated under reduced pressure and the pH of the resulting residue was adjusted to pH 10 with aq. $NaHCO_3$ (30 mL). The aqueous phase was extracted with EtOAc (10 mL x 3) and the combined organic extracts were washed with brine (10 mL), dried over anhydrous $Na_2SO_4$, filtered and concentrated under reduced pressure to afford **SI-92** (160 mg, crude) as a yellow oil which was used without purification.

**Step 3:** Following **General Procedure 1**, starting with **SI-92** (160 mg, 0.57 mmol, 1.0 eq), chloroacetyl chloride (2.0 eq), and TEA (3.0 eq), **EV-91** was obtained after prep-HPLC (FA) as a yellow solid (63 mg, 31%).

**$^1$H NMR (400 MHz, DMSO-*d*$_6$) δ** 8.00 (br s, 1H), 7.39 (br s, 3H), 7.26 (t, *J* = 7.8 Hz, 2H), 7.22 (br s, 1H), 7.12 (br s, 1H), 7.03 (br s, 1H), 4.93 – 4.45 (m, 1H), 4.38 – 3.97 (m, 1H), 3.80 (br s, 1H), 3.67 (br s, 1H), 3.55 – 3.36 (m, 1H), 3.09 (br s, 1H), 2.91 (br s, 1H), 2.39 – 2.03 (m, 2H).

**HRMS ESI-TOF (*m/z*):** $[M+H]^+$ for $C_{20}H_{20}ClN_2O_2$ 355.1208, found 355.1207.

#### Synthesis of SI-97 as a precursor for EV-94 and EV-95

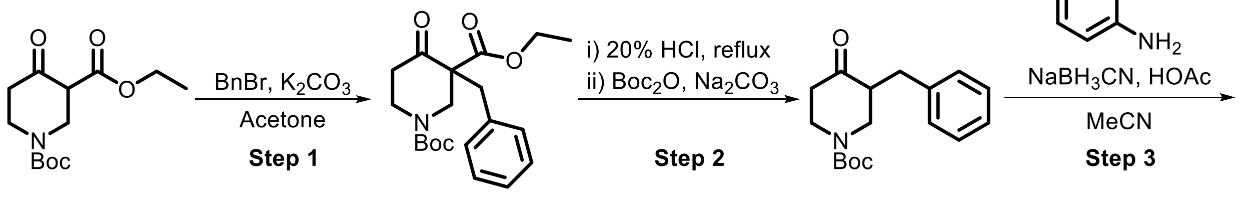

**SI-93** **SI-94**

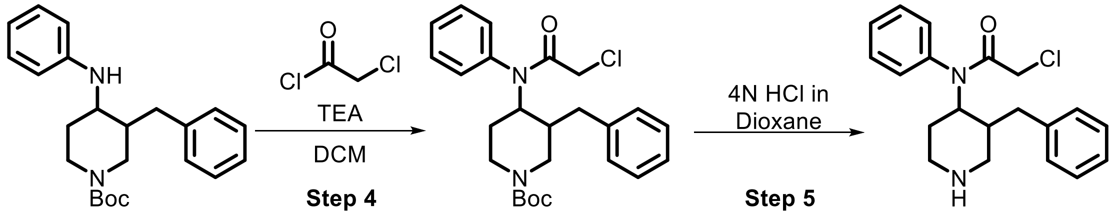

**SI-95** **SI-96** **SI-97**

**Step 1:** Benzyl bromide (1.8 mL, 14.8 mmol, 2.0 eq) was added dropwise to a solution of *N*-Boc-3-carboethoxy-4-piperidone (2.0 g, 7.4 mmol, 1.0 eq) and $K_2CO_3$ (3.1 g, 22 mmol, 3.0 eq) in acetone (80 mL). The mixture was stirred at reflux for 12 h. The reaction was cooled and then filtered. The filtrate was concentrated and $H_2O$ (50 mL) was added. The solution was extracted with EtOAc (40 mL x 2). The combined organic layers were dried over anhydrous $Na_2SO_4$ and concentrated to provide **SI-93** (1.7 g, crude) which was used in the next step without purification.

**Step 2:** **SI-93** (1.7 g, 4.7 mmol, 1.0 eq) was dissolved in 20% HCl (25 mL) and heated to reflux for 48 h. The solvent was removed and the resulting residue was dissolved in THF (50 mL). $Boc_2O$ (2.0 g, 9.2 mmol, 2.0 eq) and $Na_2CO_3$ (1.5 g, 14.1 mmol, 3.0 eq) were added and the mixture was stirred for 12 h. $H_2O$ (50 mL) was added the mixture was extracted with EtOAc (40 mL x 3). The combined organic phases were dried over anhydrous $Na_2SO_4$ and concentrated to provide **SI-94** as a crude product which was used without purification.(Hartman and Flores, 2013)

**Step 3:** Following **General Procedure 5**, starting with **SI-94** (1.0 eq) and aniline (1.0 eq), **SI-95** was obtained as a crude product and was used in the next step without purification.

**Step 4:** Following **General Procedure 1**, starting with **SI-95** (1.0 eq), chloroacetyl chloride (2.0 eq), and TEA (5.0 eq), **SI-96** was obtained as crude product and used without purification.

**Step 5:** HCl/dioxane (4N, 1.1 mL, 5.0 eq) was added to a solution of **SI-95** (400 mg, 0.87 mmol, 1.0 eq) in dioxane (2 mL) at 0 °C. The reaction was warmed to 25 °C and stirred for 3 h. The reaction was concentrated under reduced pressure and partitioned between DCM (20 mL) and saturated $NaHCO_3$ (20 mL). The aqueous phase was extracted with DCM (20 mL x 2) and the combined organic phases were dried over anhydrous $Na_2SO_4$ and concentrated under reduced pressure to provide **SI-96** (300 mg, crude) which was used without purification.

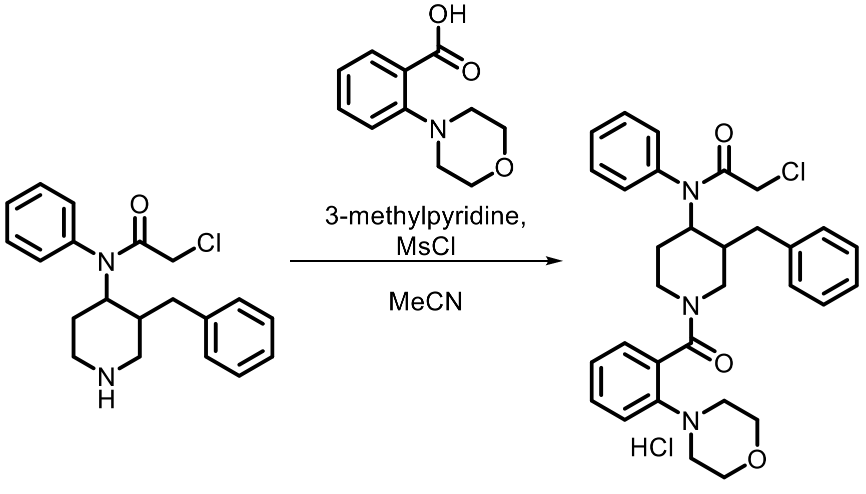

**SI-97** **EV-92**

3-methylpyridine (85 µL, 0.87 mmol, 3.0 eq) and MsCl (34 µL, 0.44 mmol, 1.5 eq) were added to a solution of **SI-97** (100 mg, 0.29 mmol, 1.0 eq) and 2-morpholinobenzoic acid (60 mg, 0.29 mmol, 1.0 eq) in MeCN (1 mL) at 0 °C. The reaction was warmed to 25 °C and stirred for 3 h. The mixture was quenched with $H_2O$ (1 mL) and extracted with EtOAc (0.5 mL x 3). The combined organic layers were dried over anhydrous $Na_2SO_4$ and concentrated under reduced pressure. The resulting residue was purified by prep. HPLC (HCl) to provide **EV-92** (9 mg, 5%) as a white solid.

$^1$**H NMR (400 MHz, MeOD) δ** 7.63 – 7.47 (m, 4H), 7.46 – 7.39 (m, 1H), 7.38 – 7.28 (m, 4H), 7.27 – 7.21 (m, 2H), 7.21 – 7.11 (m, 1H), 7.05 – 6.74 (m, 2H), 4.66 – 4.51 (m, 1H), 3.95 – 3.81 (m, 2H), 3.77 – 3.61 (m, 4H), 3.41 – 3.32 (m, 2H), 3.18 – 2.99 (m, 4H), 2.98 – 2.85 (m, 3H), 2.82 – 2.61 (m, 2H), 1.76 – 1.42 (m, 2H).

**HRMS ESI-TOF (*m/z*):** $[M+H]^+$ for $C_{31}H_{34}ClN_3O_3$ 523.2362, found 532.2363.

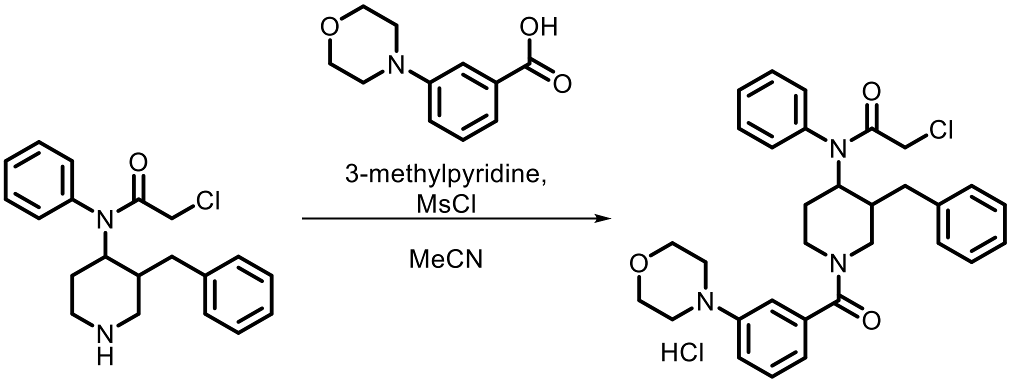

**SI-97** **EV-93**

3-methylpyridine (85 μL, 0.87 mmol, 3.0 eq) and MsCl (34 μL, 0.44 mmol, 1.5 eq) were added to a solution of **SI-97** (100 mg, 0.29 mmol, 1.0 eq) and 3-morpholinobenzoic acid (60 mg, 0.29 mmol, 1.0 eq) in MeCN (1 mL) at 0 °C. The reaction was warmed to 25 °C and stirred for 3 h. The mixture was quenched with $H_2O$ (1 mL) and extracted with EtOAc (0.5 mL x 3). The combined organic layers were dried over anhydrous $Na_2SO_4$ and concentrated under reduced pressure. The resulting residue was purified by prep. HPLC (HCl) to provide **EV-93** (17 mg, 11%) as a white solid.

$^1$**H NMR (400 MHz, MeOD) δ** 7.50 (br s, 5H), 7.39 – 7.26 (m, 3H), 7.23 – 7.09 (m, 3H), 7.06 – 6.97 (m, 2H), 6.92 (d, *J* = 7.5 Hz, 1H), 4.67 – 4.57 (m, 2H), 3.90 – 3.77 (m, 5H), 3.74 – 3.66 (m, 1H), 3.56 – 3.45 (m, 1H), 3.26 – 3.18 (m, 3H), 3.15 – 3.04 (m, 1H), 3.04 – 2.96 (m, 1H), 2.95 – 2.85 (m, 1H), 2.84 – 2.77 (m, 1H), 2.50 (br s, 1H), 1.73 – 1.51 (m, 2H).

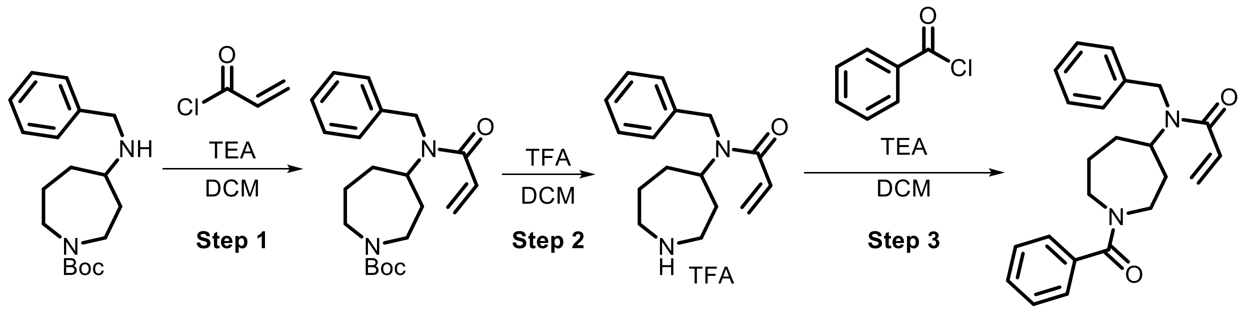

**SI-98** **SI-99** **EV-94**

**Step 1:** Following **General Procedure 2**, starting with *tert*-butyl 4-(benzylamino)azepane-1-carboxylate (1.0 eq)(Bar-Peled et al., 2017), acryloyl chloride (1.0 eq), and TEA (3.0 eq), **SI-98** was obtained as crude product and used without purification.

**Step 2:** TFA (1.5 mL, 19.5 mmol, 5.0 eq) was added to a solution of **SI-98** (1.4 g, 3.9 mmol, 1.0 eq) in DCM (10 mL) and the resulting mixture was stirred for 0.5 h. The reaction was quenched by the addition of water (20 mL) and extracted with DCM (10 mL x 3). The combined organic phases were dried over anhydrous $Na_2SO_4$ and concentrated under reduced pressure to provide **SI-99** (800 mg, crude) as a yellow oil which was used without purification.

**Step 3:** Following **General Procedure 4**, starting with **SI-99** (1.0 eq) and benzoyl chloride (1.2 eq), **EV-94** (50 mg, 3%) was obtained after prep-HPLC (basic) as a white solid.

**$^1$H NMR (400 MHz, $CDCl_3$) δ** 7.40 – 7.25 (m, 8H), 7.21 (t, *J* = 8.1 Hz, 2H), 6.47 – 6.29 (m, 2H), 5.70 – 5.59 (m, 1H), 4.63 – 4.47 (m, 3H), 3.96 (dd, *J* = 89.1, 14.1 Hz, 1H), 3.60 – 3.37 (m, 2H), 3.36 – 3.17 (m, 2H), 2.14 – 1.89 (m, 2H), 1.85 – 1.70 (m, 3H), 1.70 – 1.59 (m, 1H).

**HRMS ESI-TOF (*m/z*):** $[M+H]^+$ for $C_{23}H_{27}N_2O_2$ 363.2067, found 363.2073.

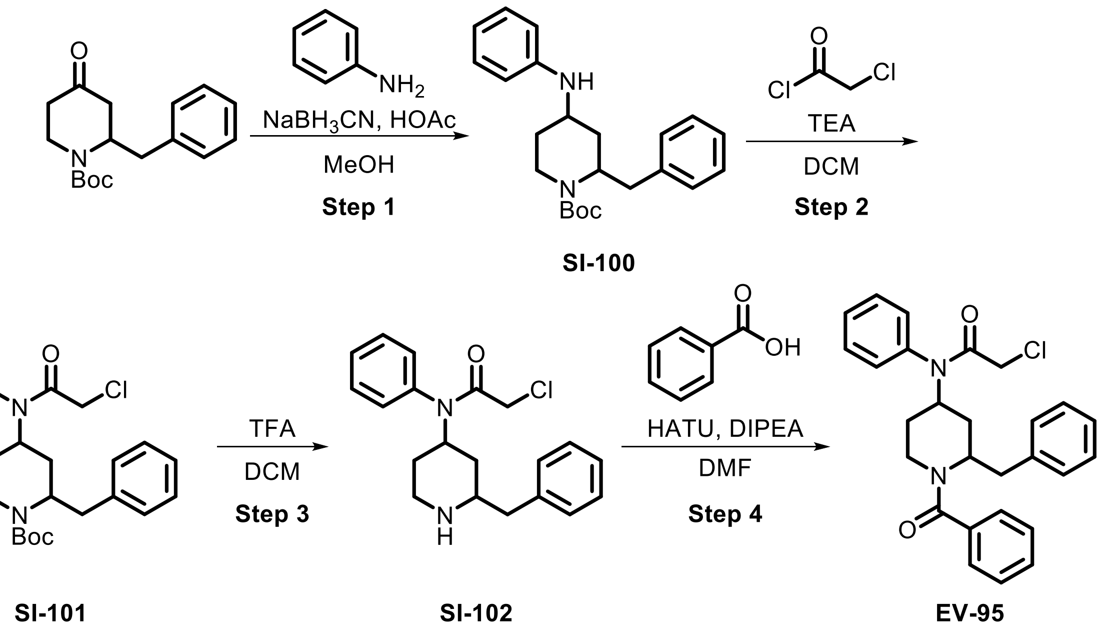

**Step 1:** Following **General Procedure 5**, starting with 1-Boc-2-benzyl-4-piperidinone (1.0 eq) and aniline (1.0 eq), **SI-100** (300 mg, crude) was obtained as a crude product and used without purification.

**Step 2:** Following **General Procedure 1**, starting with **SI-100** (1.0 eq), chloroacetyl chloride (2.0 eq), and TEA (5.0 eq), **SI-101** (150 mg, crude) was obtained and used without purification.

**Step 3:** TFA (25 µL, 0.34 mmol, 1.0 eq) was added to a solution of **SI-101** (150 mg, 0.34 mmol, 1.0 eq) in DCM (1 mL) at 0 °C. The mixture was then warmed to 25 °C and stirred for 2 h. The reaction was quenched with water (3 mL) and extracted with DCM (1 mL x 3). The combined organic layers were washed with brine (2 mL x 3), dried over anhydrous $Na_2SO_4$, and concentrated to provide **SI-102** (100 mg, crude) which was used in the next step without purification.

**Step 4:** Following **General Procedure 3**, starting with **SI-102** (1.0 eq) and benzoic acid (1.2 eq), **EV-95** (27 mg, 13%) was obtained after prep-HPLC (TFA) as a yellow oil.

**$^1$H NMR (400 MHz, DMSO-$d_6$) δ** 7.51 – 7.44 (m, 3H), 7.43 – 7.36 (m, 3H), 7.28 – 7.15 (m, 7H), 7.06 (d, *J* = 7.2 Hz, 2H), 4.50 (d, *J* = 10.4 Hz, 1H), 4.21 (br s, 1H), 3.75 (s, 2H), 3.42 (d, *J* = 7.1 Hz, 1H), 3.07 – 2.97 (m, 1H), 2.91 (dd, *J* = 13.2, 5.4 Hz, 1H), 2.79 (dd, *J* = 13.1, 7.8 Hz, 1H), 2.06 – 1.90 (m, 1H), 1.81 (br s, 1H), 1.50 (q, *J* = 12.2 Hz, 1H), 1.34 (br s, 1H).

**HRMS ESI-TOF (*m/z*):** $[M+H]^+$ for $C_{27}H_{28}ClN_2O_2$ 447.1834, found 447.1836.

### (E) References

Ashburner, M., Ball, C.A., Blake, J.A., Botstein, D., Butler, H., Cherry, J.M., Davis, A.P., Dolinski, K., Dwight, S.S., Eppig, J.T., *et al.* (2000). Gene ontology: tool for the unification of biology. The Gene Ontology Consortium. Nat Genet *25*, 25-29.
Backus, K.M., Correia, B.E., Lum, K.M., Forli, S., Horning, B.D., Gonzalez-Paez, G.E., Chatterjee, S., Lanning, B.R., Teijaro, J.R., Olson, A.J., *et al.* (2016). Proteome-wide covalent ligand discovery in native biological systems. Nature *534*, 570-574.
Bar-Peled, L., Kemper, E.K., Suciu, R.M., Vinogradova, E.V., Backus, K.M., Horning, B.D., Paul, T.A., Ichu, T.A., Svensson, R.U., Olucha, J., *et al.* (2017). Chemical Proteomics Identifies Druggable Vulnerabilities in a Genetically Defined Cancer. Cell *171*, 696-709 e623.
Bianco, G., Forli, S., Goodsell, D.S., and Olson, A.J. (2016). Covalent docking using autodock: Two-point attractor and flexible side chain methods. Protein Sci *25*, 295-301.
Blanc, M., David, F., Abrami, L., Migliozzi, D., Armand, F., Burgi, J., and van der Goot, F.G. (2015). SwissPalm: Protein Palmitoylation database. F1000Res *4*, 261.
Carbon, S., Ireland, A., Mungall, C.J., Shu, S., Marshall, B., Lewis, S., Ami, G.O.H., and Web Presence Working, G. (2009). AmiGO: online access to ontology and annotation data. Bioinformatics *25*, 288-289.
Cock, P.J., Antao, T., Chang, J.T., Chapman, B.A., Cox, C.J., Dalke, A., Friedberg, I., Hamelryck, T., Kauff, F., Wilczynski, B., *et al.* (2009). Biopython: freely available Python tools for computational molecular biology and bioinformatics. Bioinformatics *25*, 1422-1423.
Dobin, A., Davis, C.A., Schlesinger, F., Drenkow, J., Zaleski, C., Jha, S., Batut, P., Chaisson, M., and Gingeras, T.R. (2013). STAR: ultrafast universal RNA-seq aligner. Bioinformatics *29*, 15-21.
Forli, S., Huey, R., Pique, M.E., Sanner, M.F., Goodsell, D.S., and Olson, A.J. (2016). Computational protein-ligand docking and virtual drug screening with the AutoDock suite. Nat Protoc *11*, 905-919.
Gao, D.W., Vinogradova, E.V., Nimmagadda, S.K., Medina, J.M., Xiao, Y., Suciu, R.M., Cravatt, B.F., and Engle, K.M. (2018). Direct Access to Versatile Electrophiles via Catalytic Oxidative Cyanation of Alkenes. J Am Chem Soc *140*, 8069-8073.
Han, X., Wang, J., Wang, J., Liu, S., Hu, J., Zhu, H., and Qian, J. (2017). ScaPD: a database for human scaffold proteins. BMC Bioinformatics *18*, 386.
Hartman, G.D., and Flores, O.A., Hepatitis B antiviral agents, WO2013096744A1, 2013 Jun 27
Heinz, S., Benner, C., Spann, N., Bertolino, E., Lin, Y.C., Laslo, P., Cheng, J.X., Murre, C., Singh, H., and Glass, C.K. (2010). Simple combinations of lineage-determining transcription factors prime cis-regulatory elements required for macrophage and B cell identities. Mol Cell *38*, 576-589.
Lee, H.Y., Suciu, R.M., Horning, B.D., Vinogradova, E.V., Ulanovskaya, O.A., and Cravatt, B.F. (2018). Covalent inhibitors of nicotinamide N-methyltransferase (NNMT) provide evidence for target engagement challenges in situ. Bioorg Med Chem Lett *28*, 2682-2687.

Messina, D.N., Glasscock, J., Gish, W., and Lovett, M. (2004). An ORFeome-based analysis of human transcription factor genes and the construction of a microarray to interrogate their expression. Genome Res *14*, 2041-2047.
Morris, G.M., Huey, R., Lindstrom, W., Sanner, M.F., Belew, R.K., Goodsell, D.S., and Olson, A.J. (2009). AutoDock4 and AutoDockTools4: Automated docking with selective receptor flexibility. J Comput Chem *30*, 2785-2791.
Mortenson, D.E., Brighty, G.J., Plate, L., Bare, G., Chen, W., Li, S., Wang, H., Cravatt, B.F., Forli, S., Powers, E.T., *et al.* (2018). "Inverse Drug Discovery" Strategy To Identify Proteins That Are Targeted by Latent Electrophiles As Exemplified by Aryl Fluorosulfates. J Am Chem Soc *140*, 200-210.
Qinheng, Z., Jordan L., W., Seiya, K., Diogo, S.-M., Christopher J., S., Gencheng, L., Stefano, F., John E., M., Dennis W., W., and K. Barry, S. (2019). “Sleeping Beauty” Phenomenon: SuFEx-Enabled Discovery of Selective Covalent Inhibitors of Human Neutrophil Elastase.
Rieckmann, J.C., Geiger, R., Hornburg, D., Wolf, T., Kveler, K., Jarrossay, D., Sallusto, F., Shen-Orr, S.S., Lanzavecchia, A., Mann, M., *et al.* (2017). Social network architecture of human immune cells unveiled by quantitative proteomics. Nat Immunol *18*, 583-593.
Shifrut, E., Carnevale, J., Tobin, V., Roth, T.L., Woo, J.M., Bui, C.T., Li, P.J., Diolaiti, M.E., Ashworth, A., and Marson, A. (2018). Genome-wide CRISPR Screens in Primary Human T Cells Reveal Key Regulators of Immune Function. Cell *175*, 1958-1971 e1915.
The Gene Ontology, C. (2019). The Gene Ontology Resource: 20 years and still GOing strong. Nucleic Acids Res *47*, D330-D338.
UniProt, C. (2019). UniProt: a worldwide hub of protein knowledge. Nucleic Acids Res *47*, D506-D515.
Wang, Y., Dix, M., Remsberg, J., Lee, H.-y., Kalocsay, M., Gygi, S., Vite, G., Lawrence, M., Parker, C., and Cravatt, B. (2019). Expedited Mapping of the Ligandable Proteome Using Fully Functionalized Enantiomeric Probe Pairs. 7764638.v7764631.
Weerapana, E., Speers, A.E., and Cravatt, B.F. (2007). Tandem orthogonal proteolysis-activity-based protein profiling (TOP-ABPP)--a general method for mapping sites of probe modification in proteomes. Nat Protoc *2*, 1414-1425.
Word, J.M., Lovell, S.C., Richardson, J.S., and Richardson, D.C. (1999). Asparagine and glutamine: using hydrogen atom contacts in the choice of side-chain amide orientation. J Mol Biol *285*, 1735-1747.

### (F) NMR spectra

#### EV-1 $^1$H NMR (MeOD)

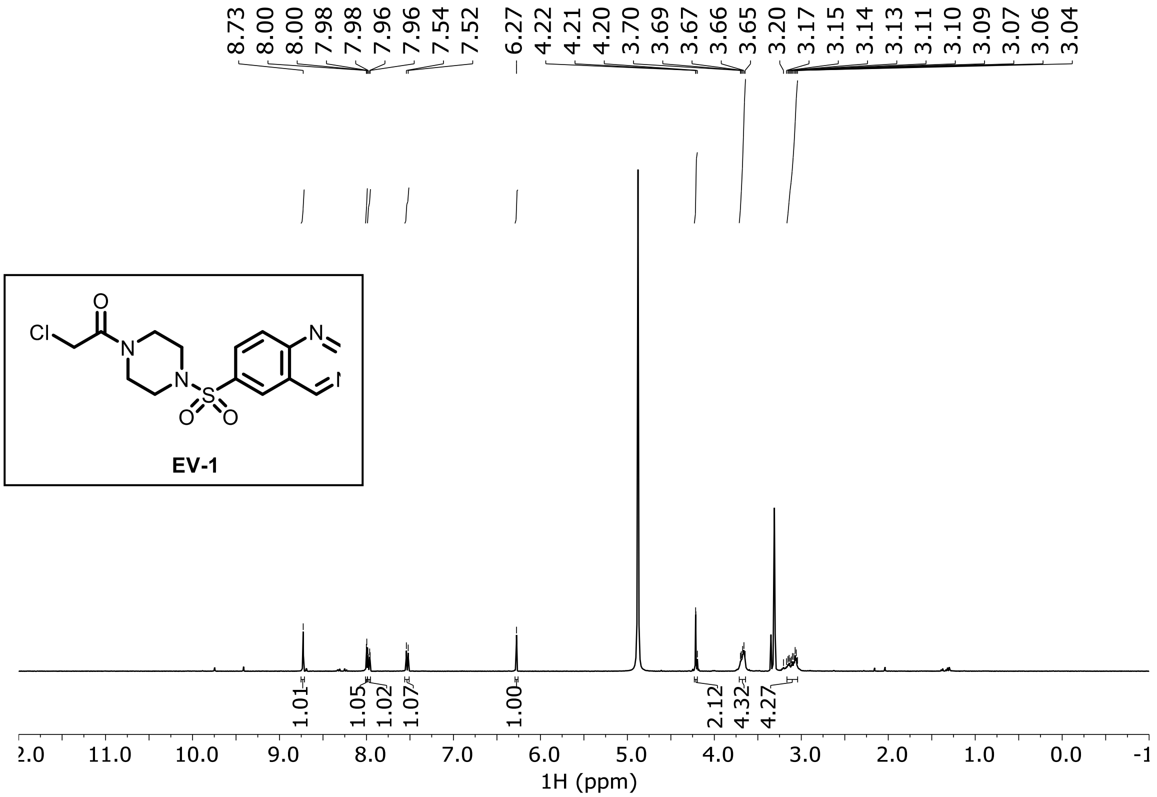

**EV-2 $^{1}$H NMR ($CDCl_3$)**

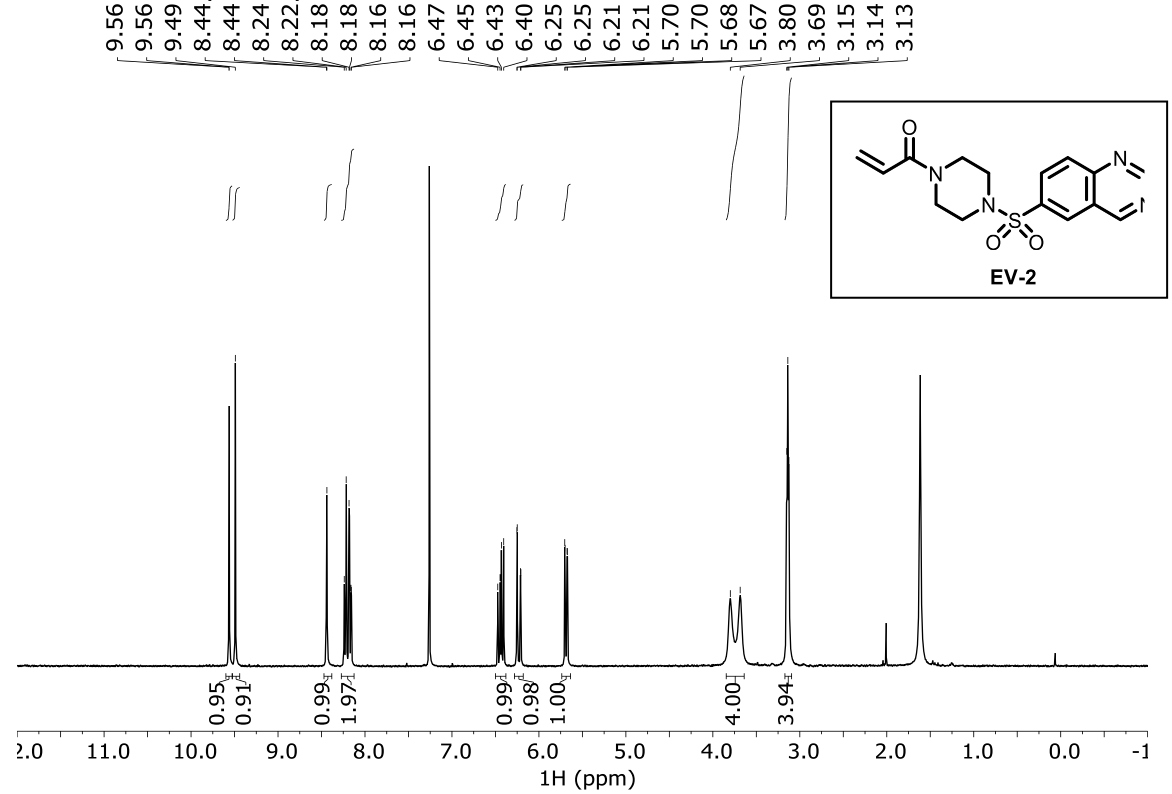

**EV-3 $^{1}$H NMR ($D_2O$)**

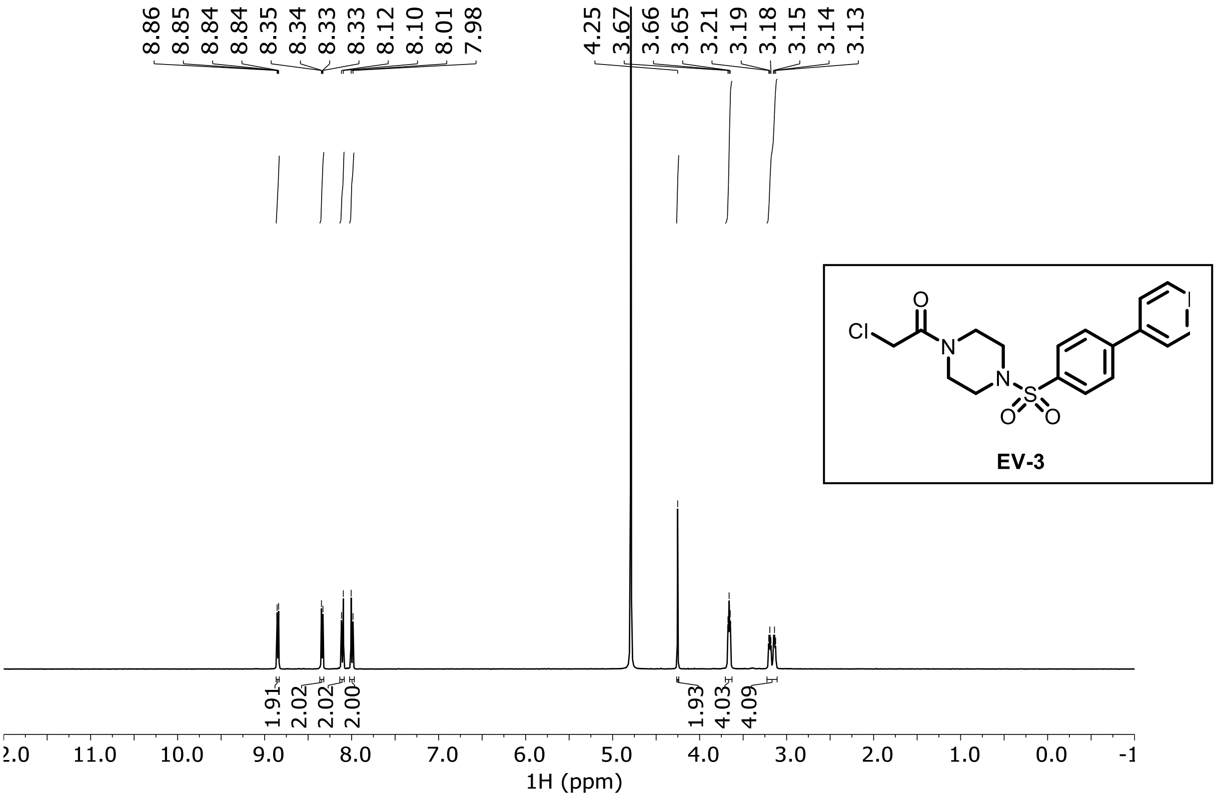

**EV-4 [1]H NMR ($CDCl_3$)**

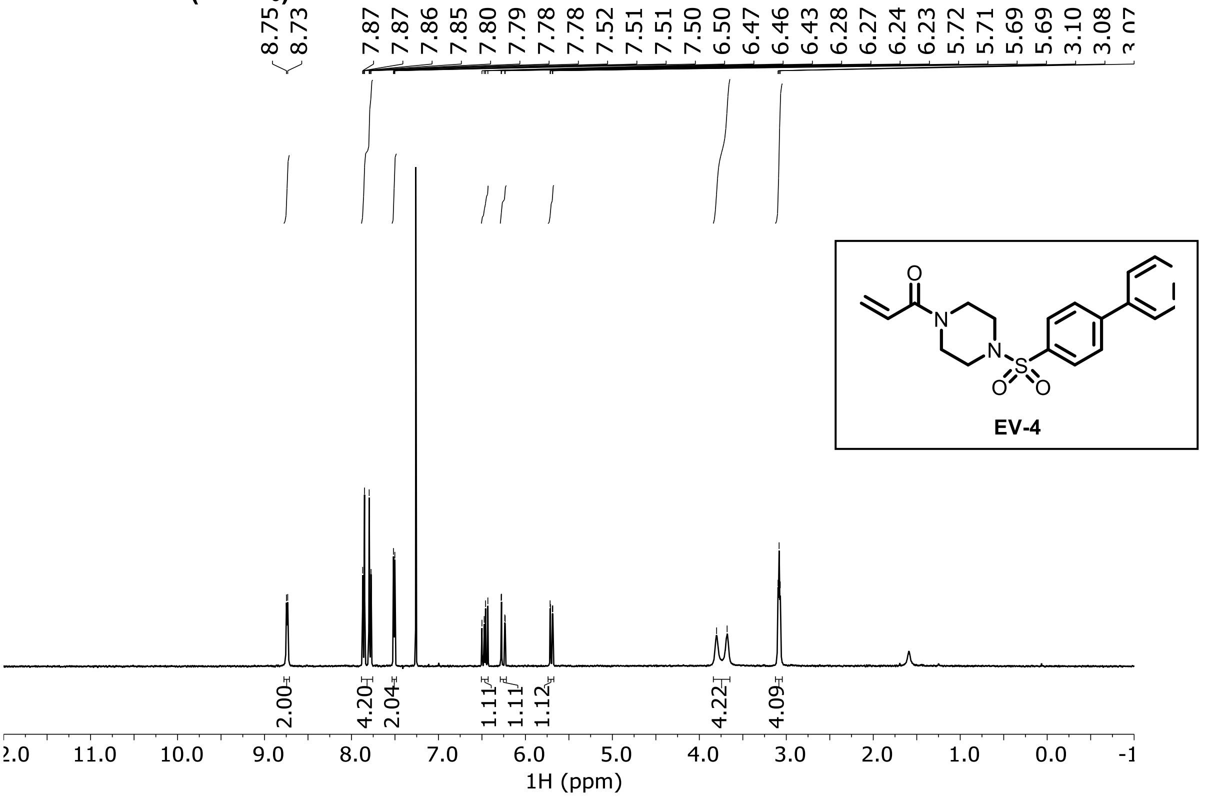

**EV-5 [1]H NMR ($CDCl_3$)**

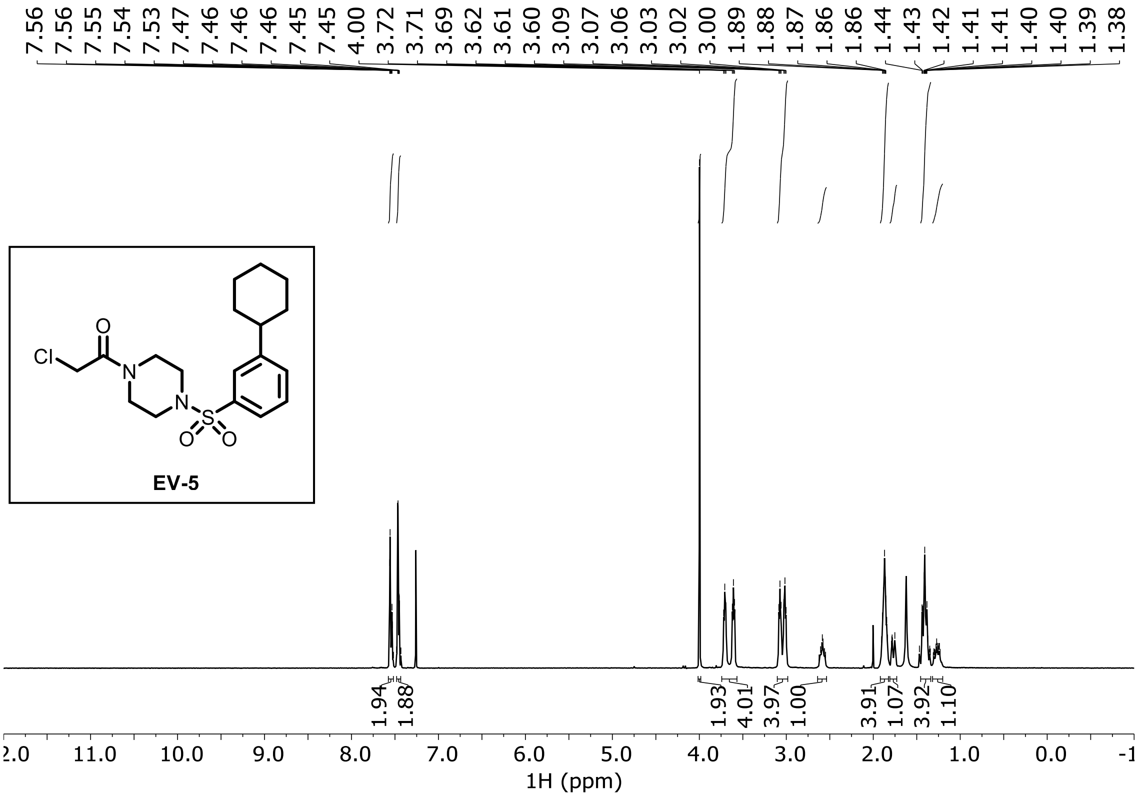

**EV-6 ¹H NMR ($CDCl_3$)**

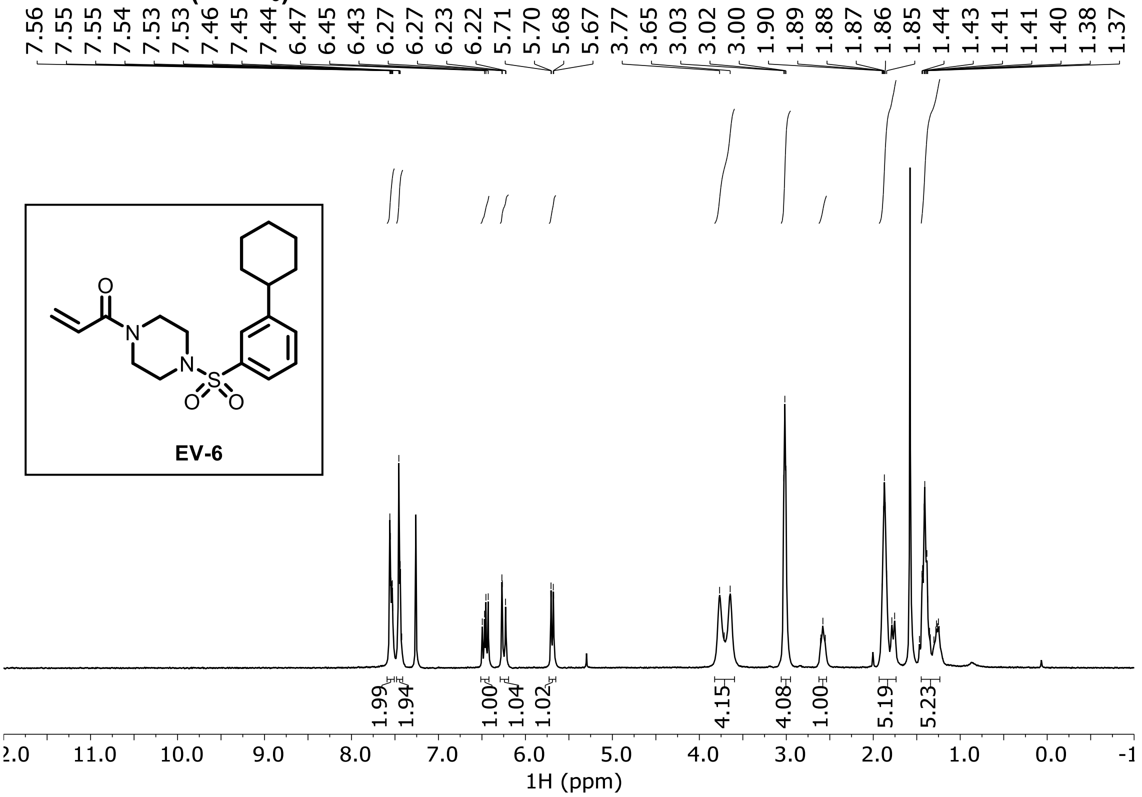

**EV-7 ¹H NMR ($CDCl_3$)**

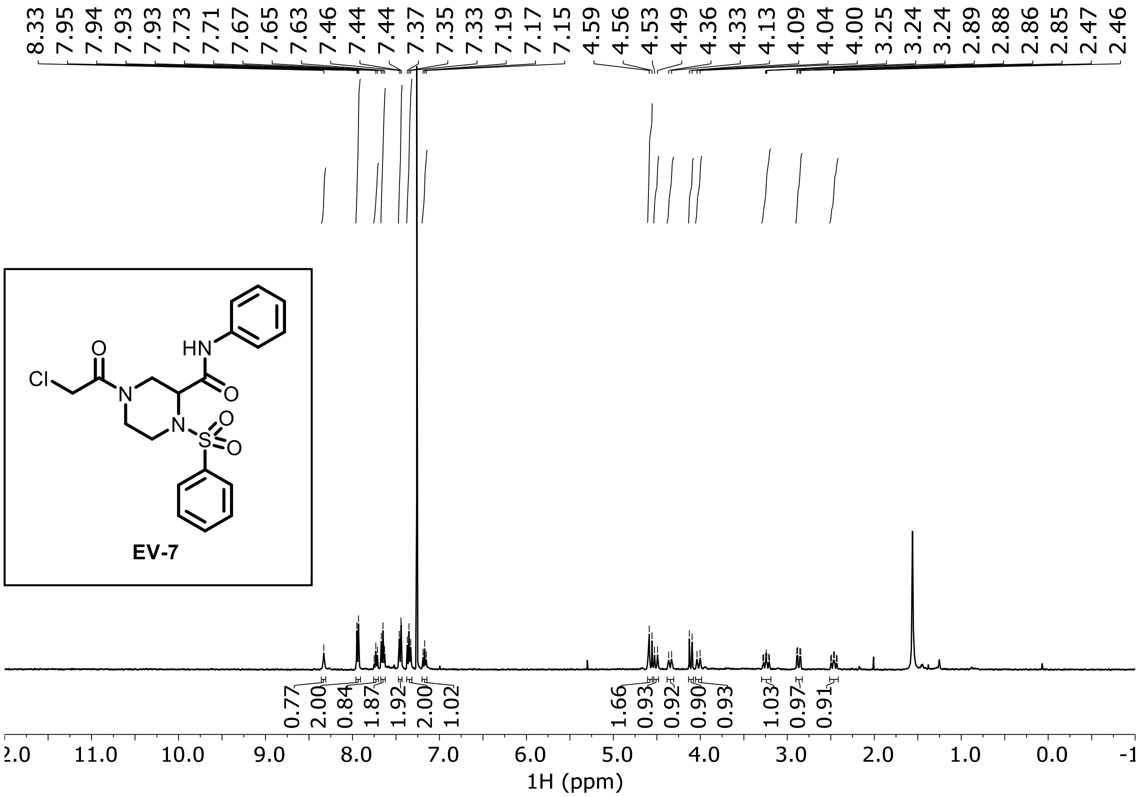

**EV-8 $^1$H NMR ($CDCl_3$)**

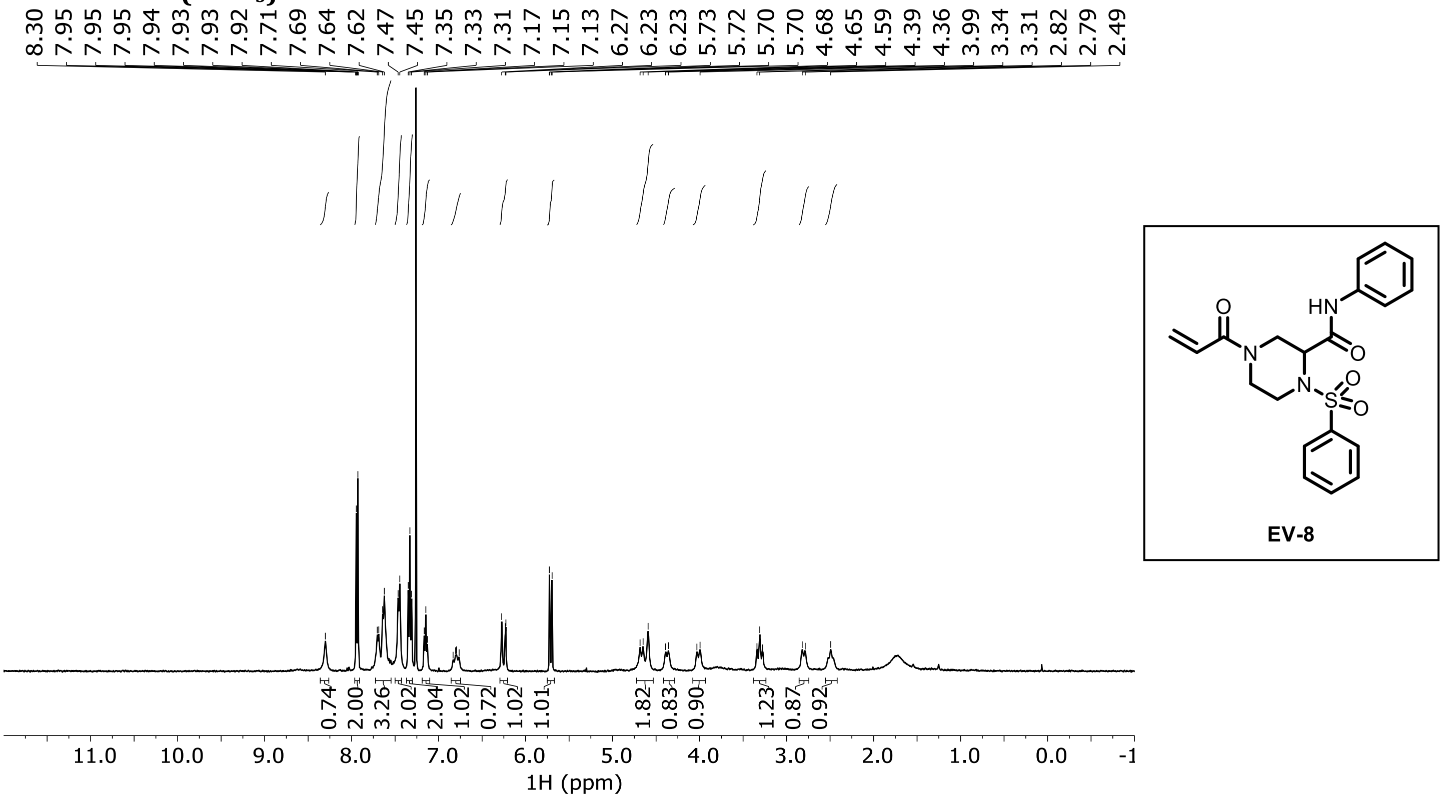

**EV-9 $^1$H NMR ($CDCl_3$)**

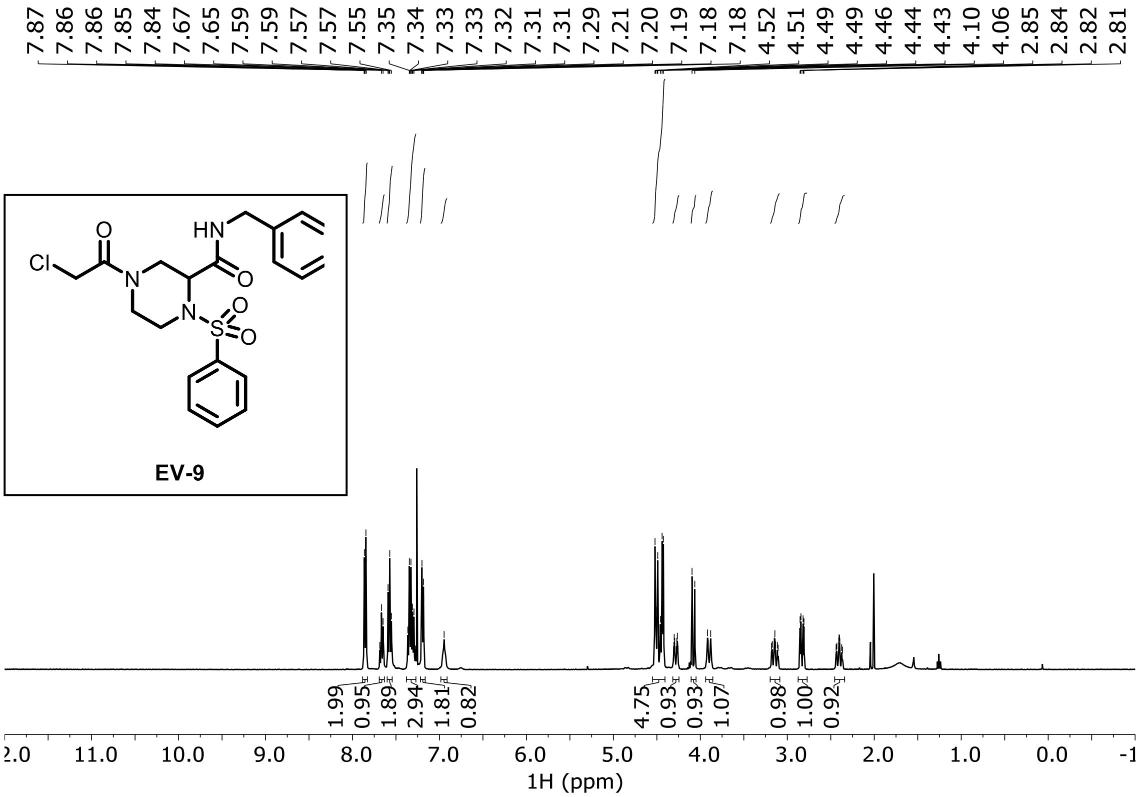

**EV-10 $^{1}H$ NMR ($CDCl_3$)**

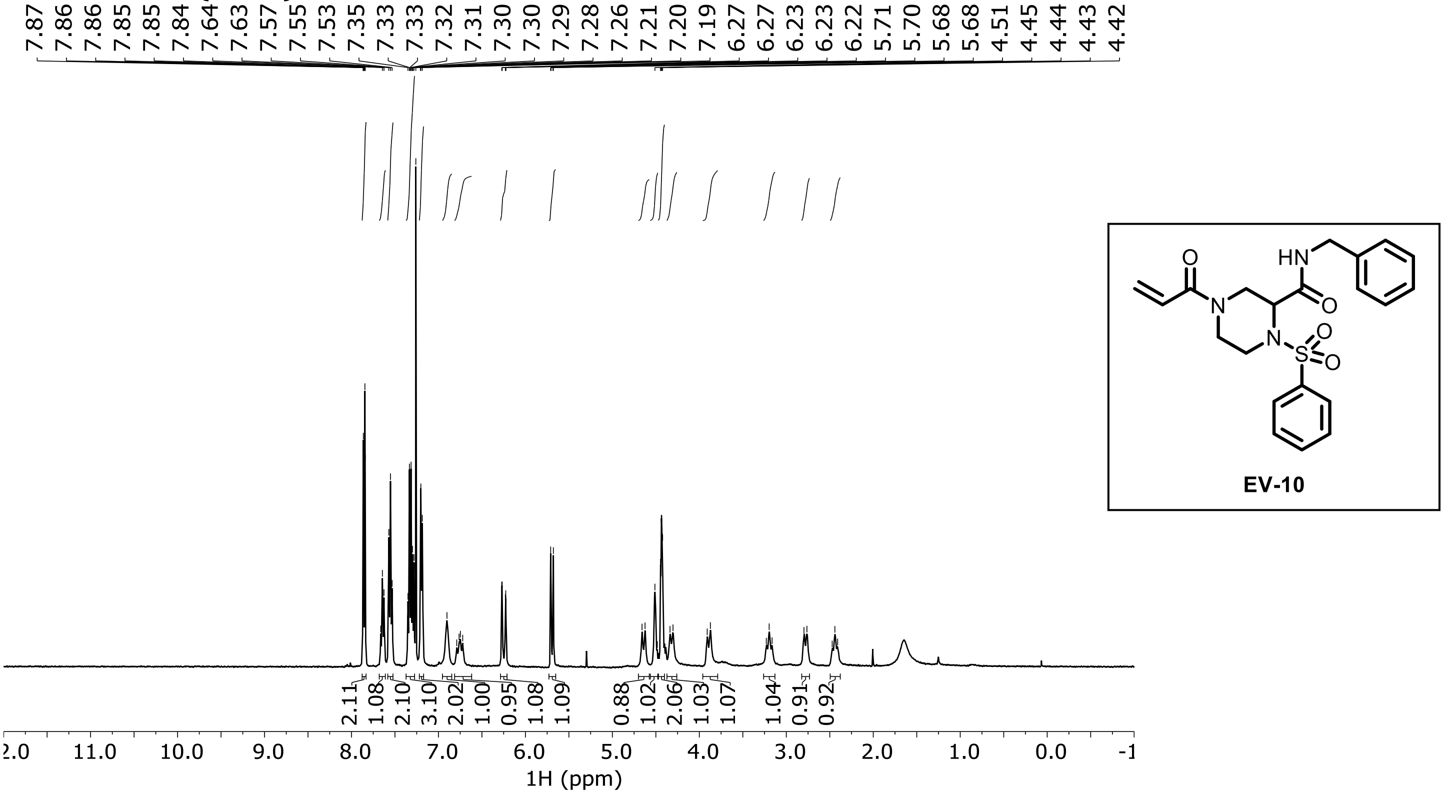

**EV-11 $^{1}H$ NMR ($CDCl_3$)**

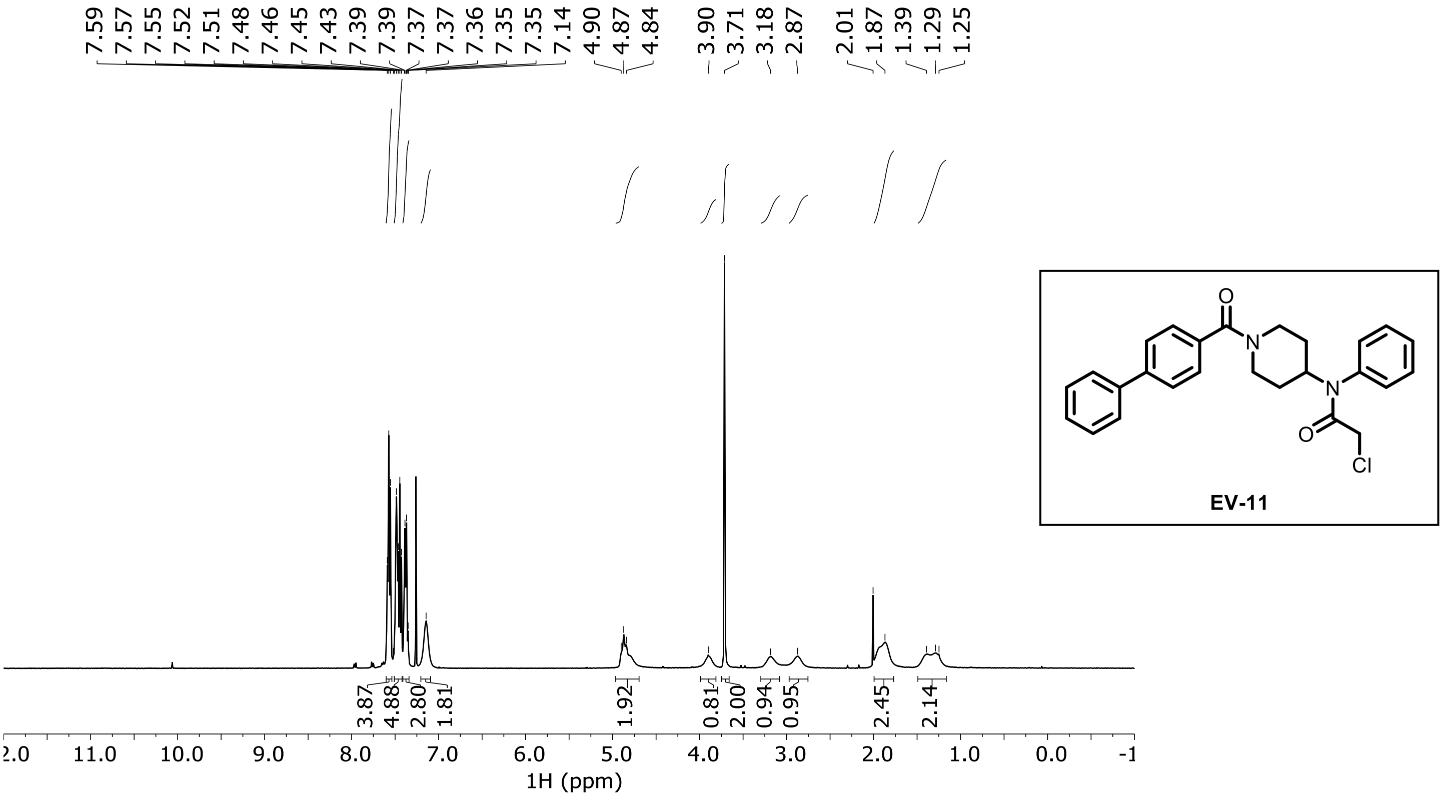

**EV-12 ¹H NMR ($CDCl_3$)**

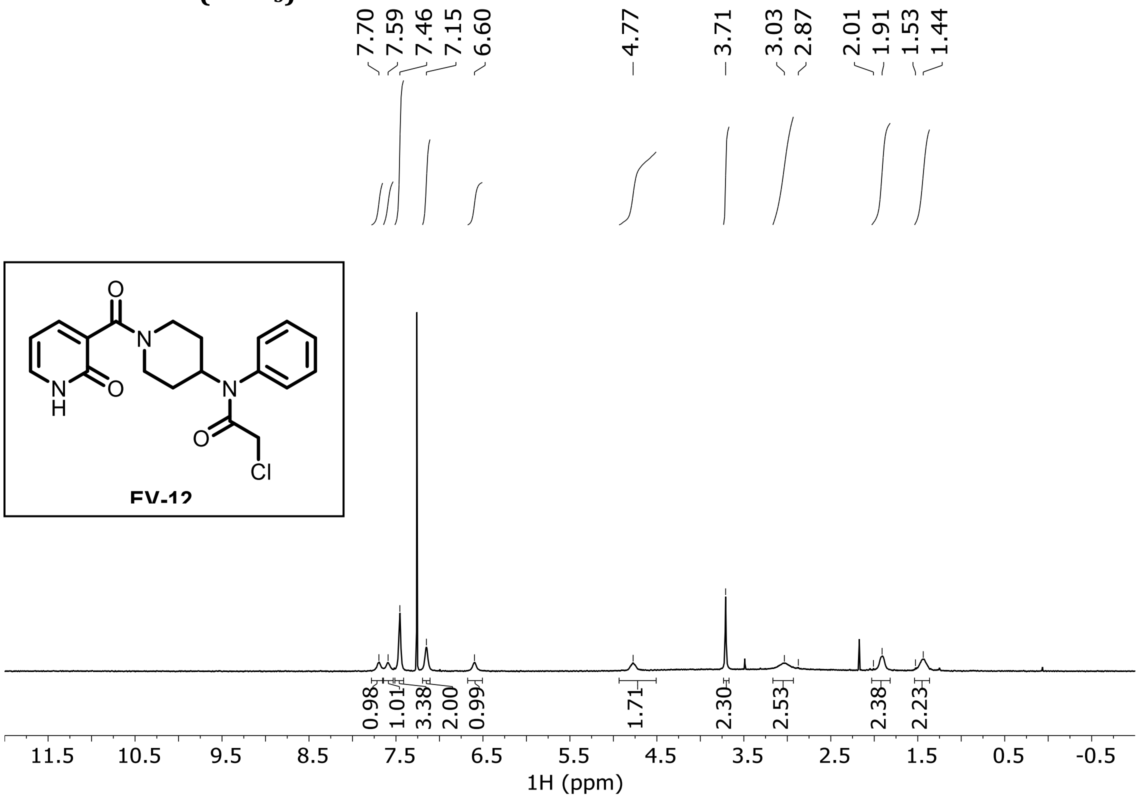

**EV-13 ¹H NMR ($CDCl_3$)**

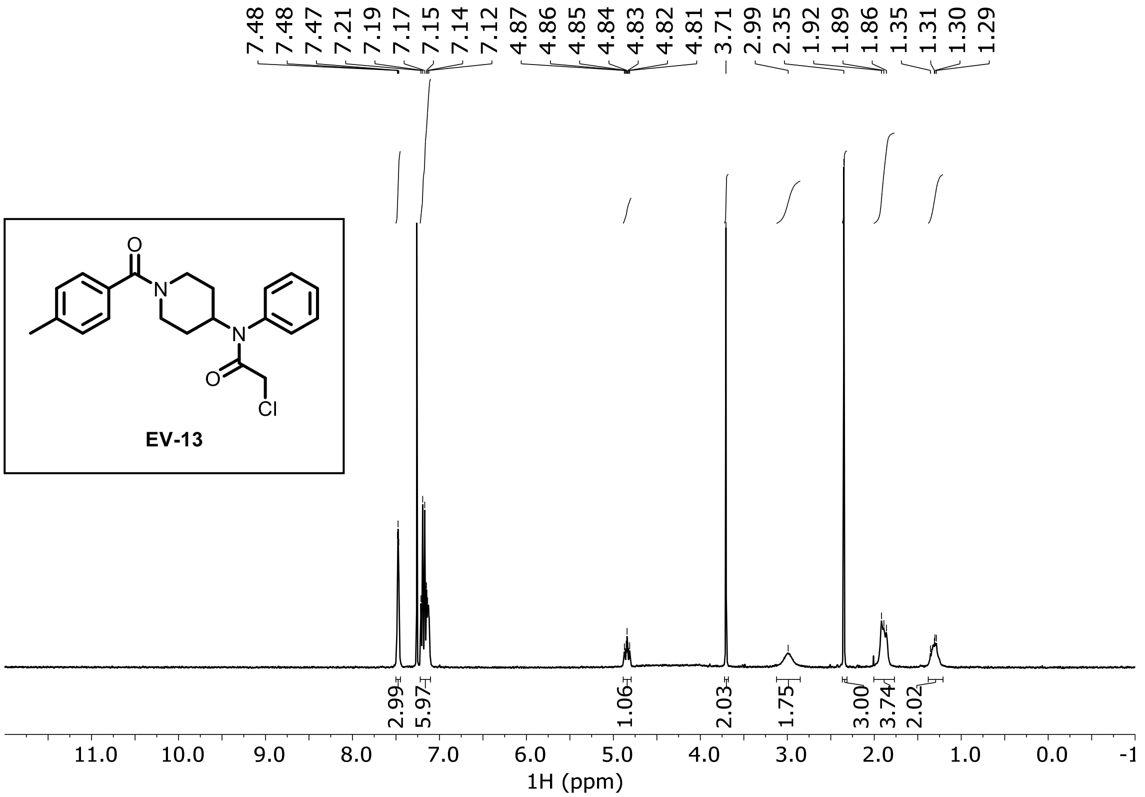

**EV-14 $^{1}$H NMR ($CDCl_3$)**

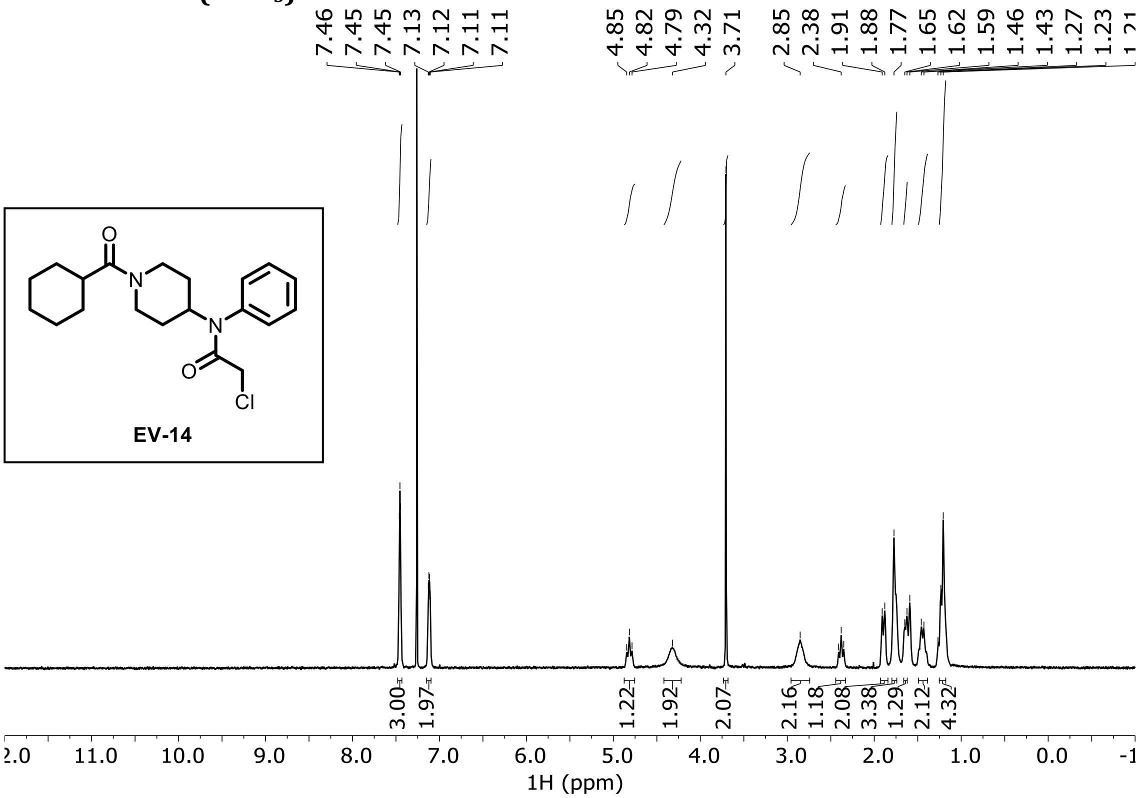

**EV-15 $^{1}$H NMR ($CDCl_3$)**

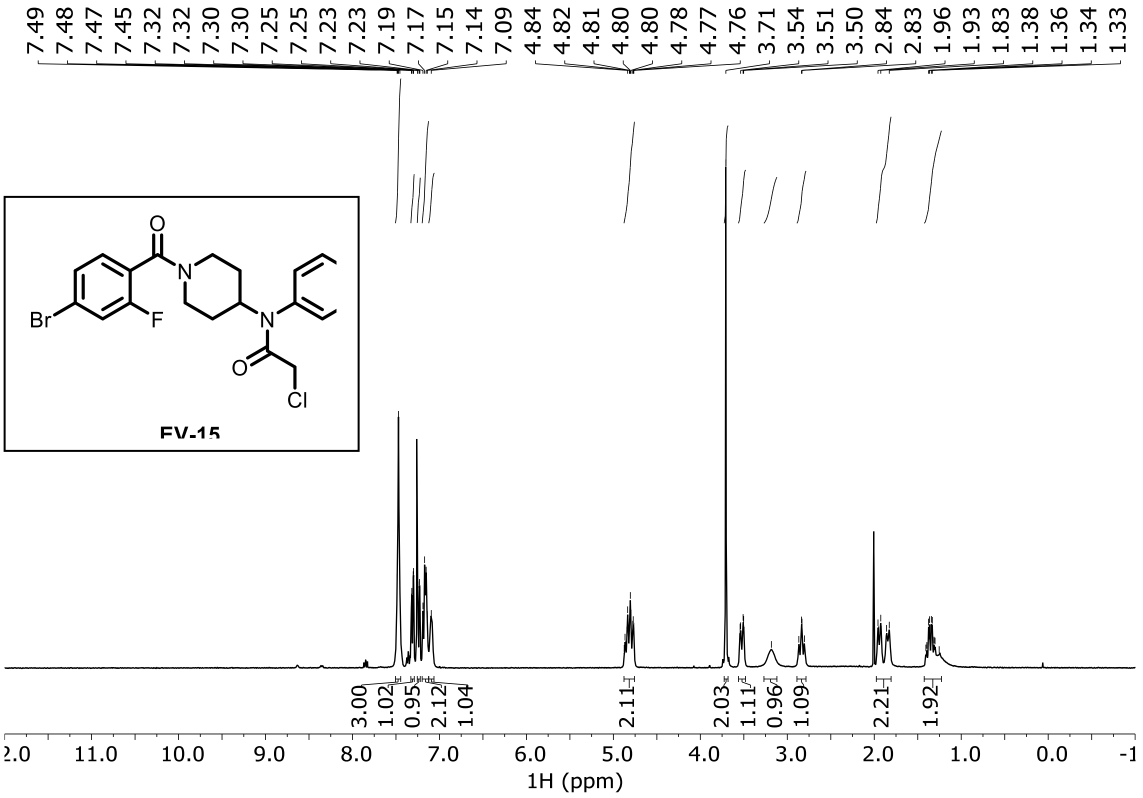

**EV-16 ¹H NMR ($CDCl_3$)**

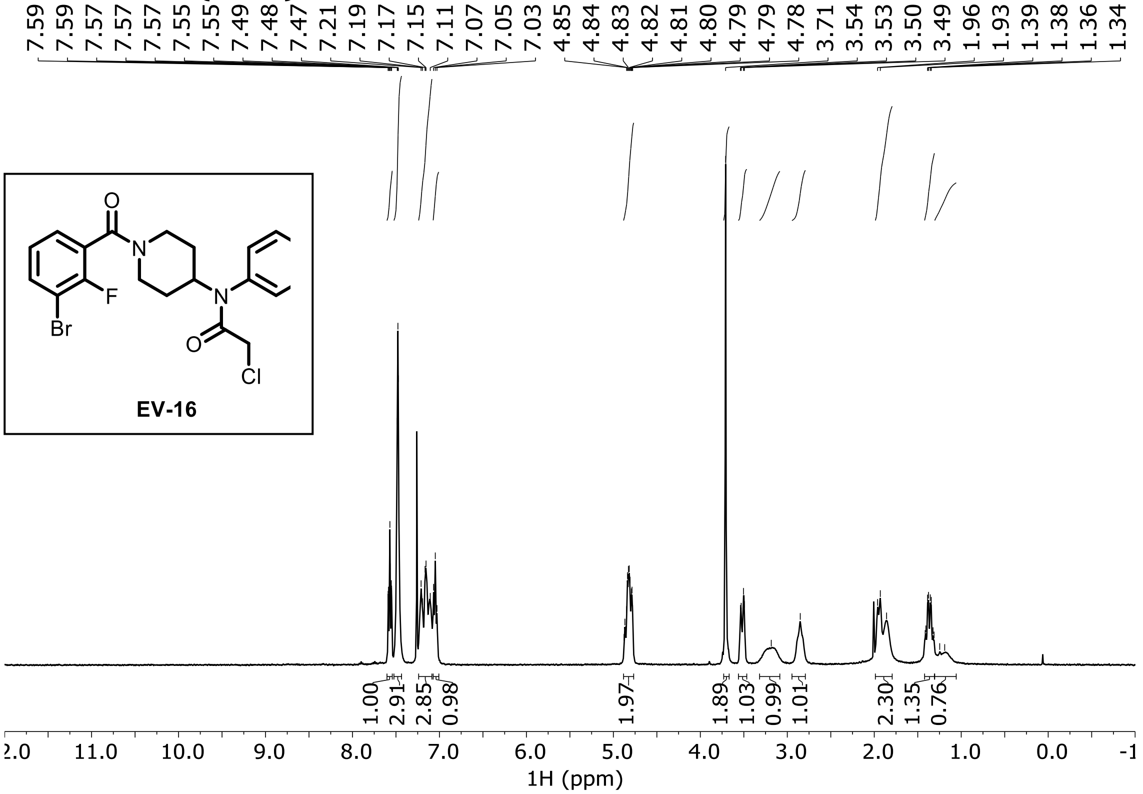

**EV-17 ¹H NMR ($CDCl_3$)**

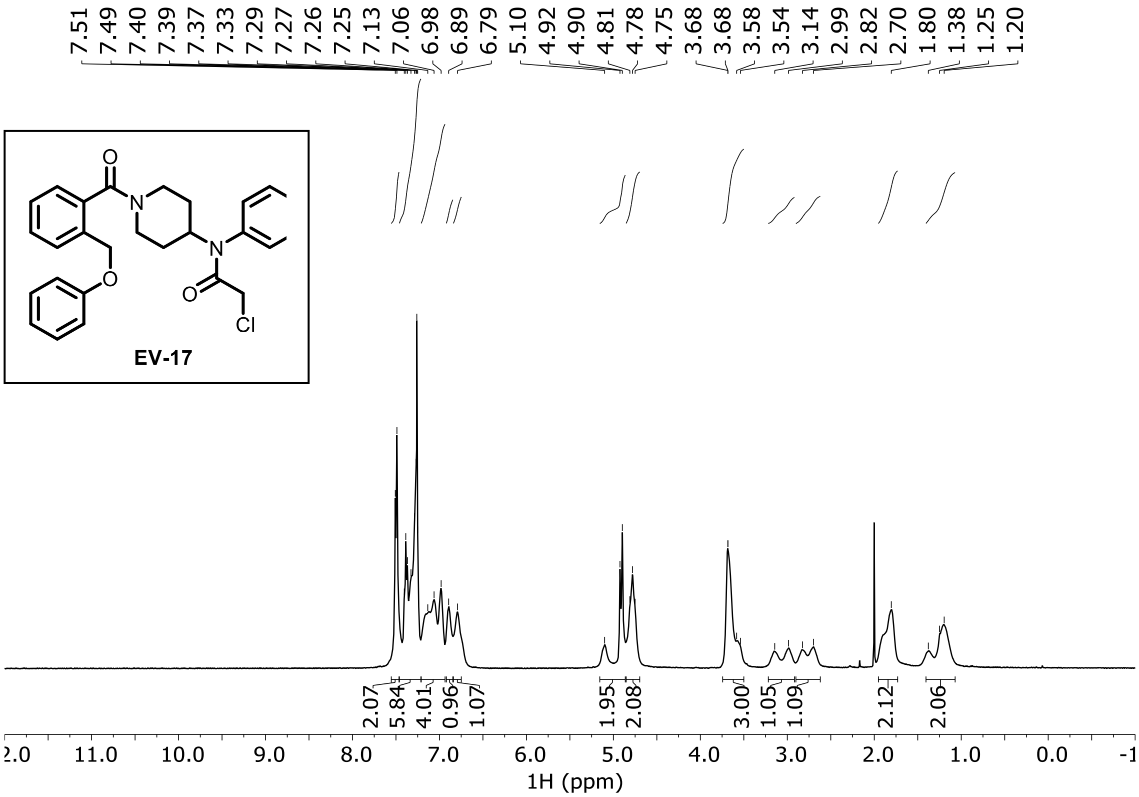

**EV-18 $^{1}$H NMR ($CDCl_3$)**

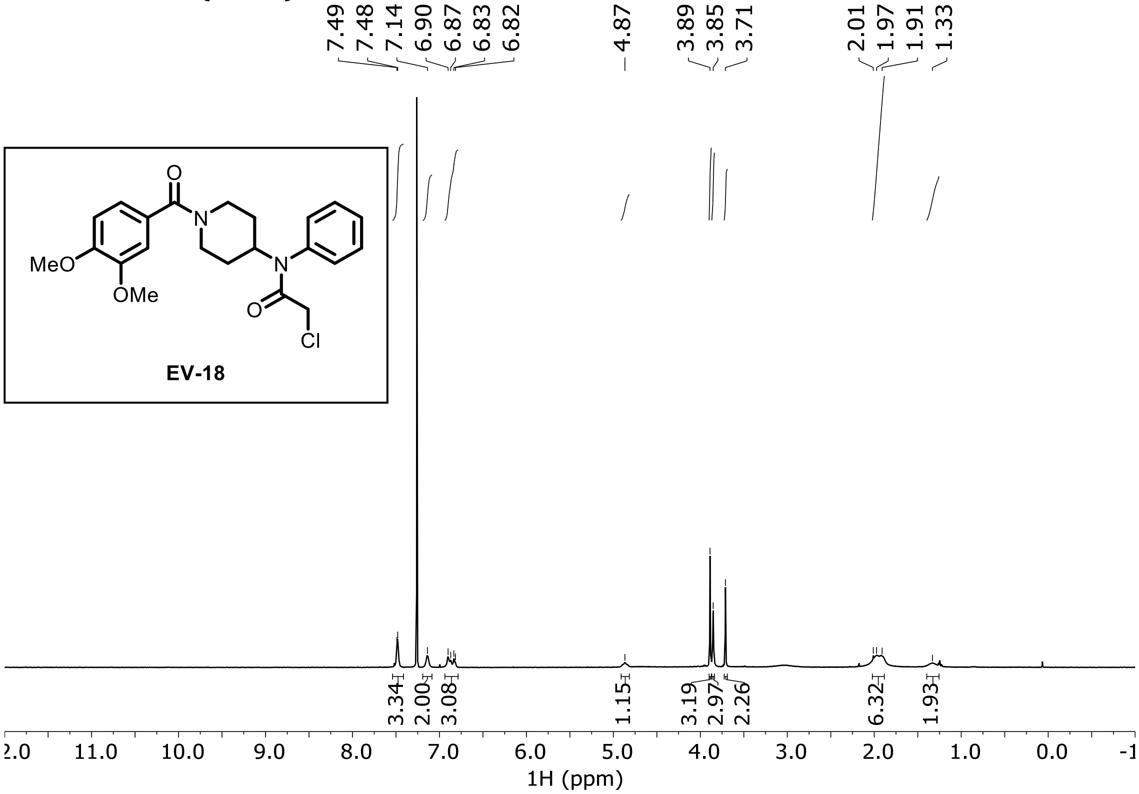

**EV-19 $^{1}$H NMR ($CDCl_3$)**

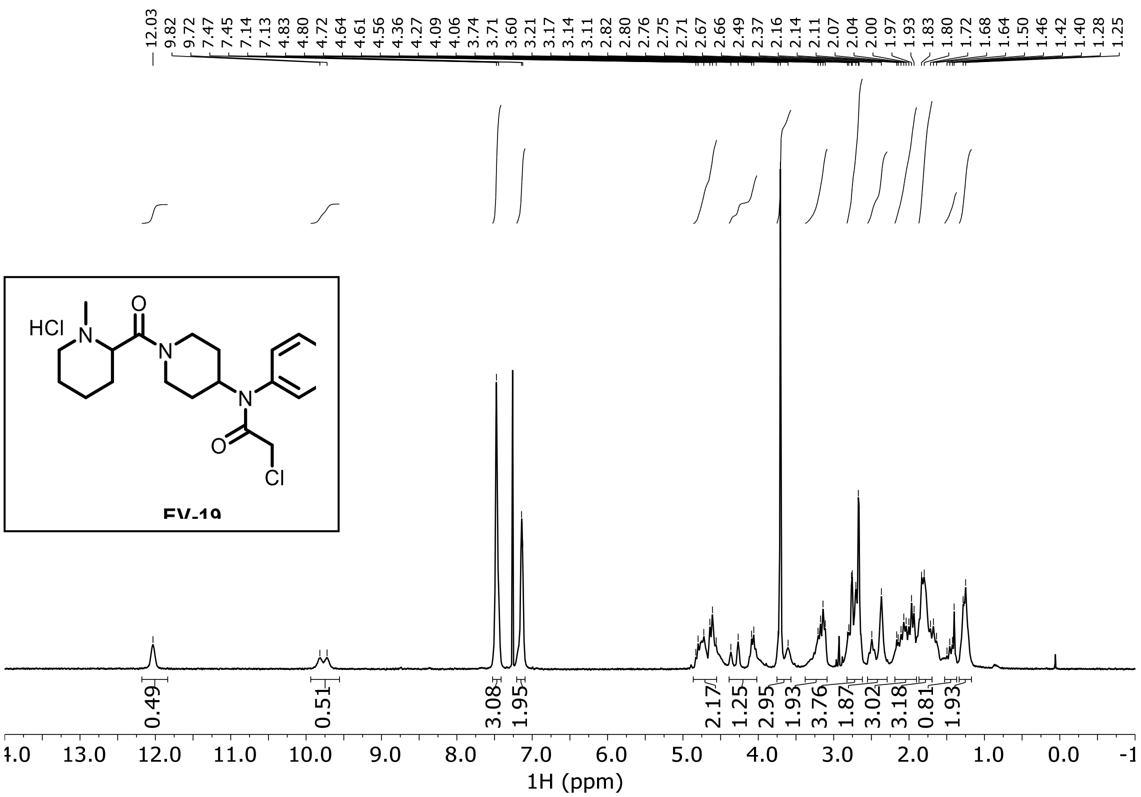

**EV-20 ¹H NMR ($CDCl_3$)**

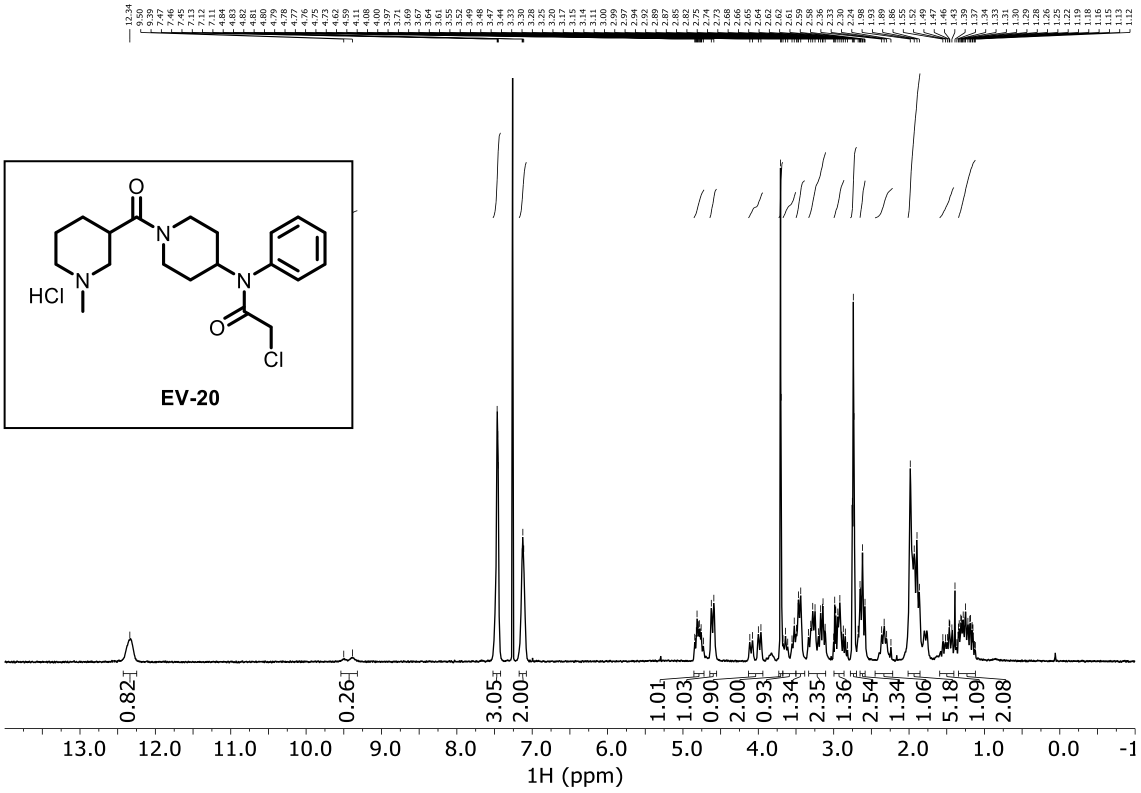

**EV-21 ¹H NMR ($CDCl_3$)**

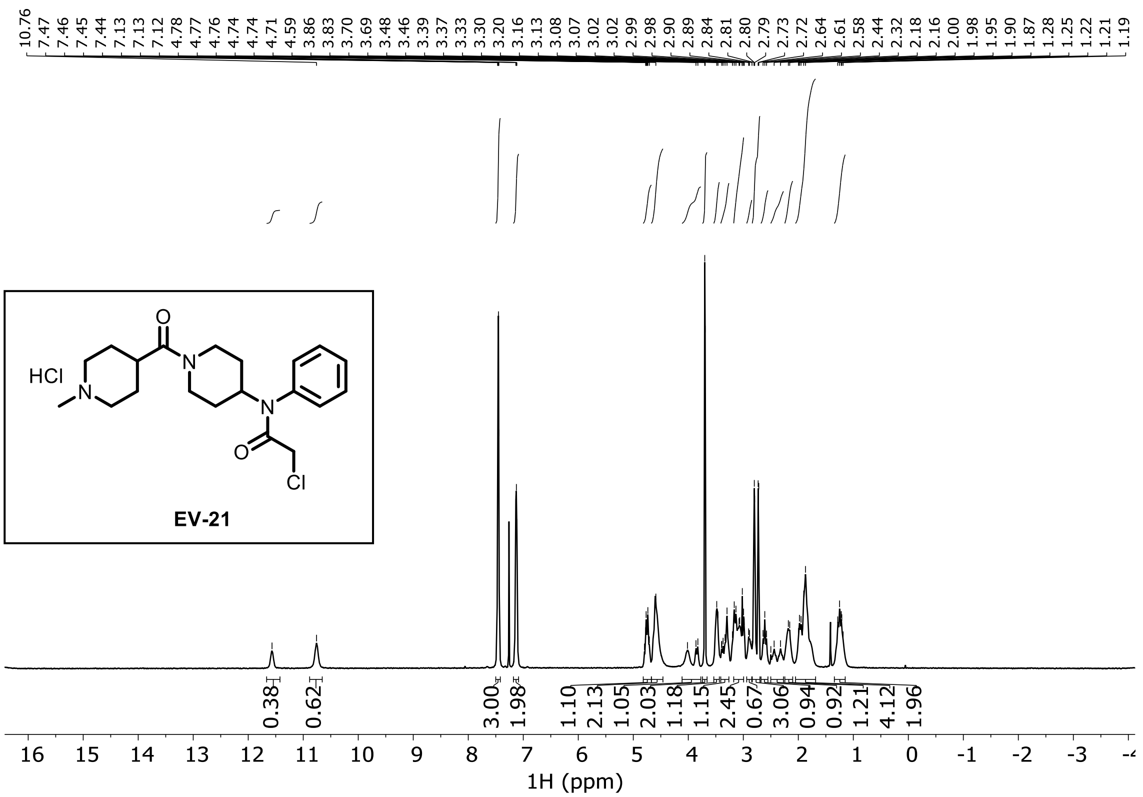

**EV-22 ¹H NMR ($D_2O$)**

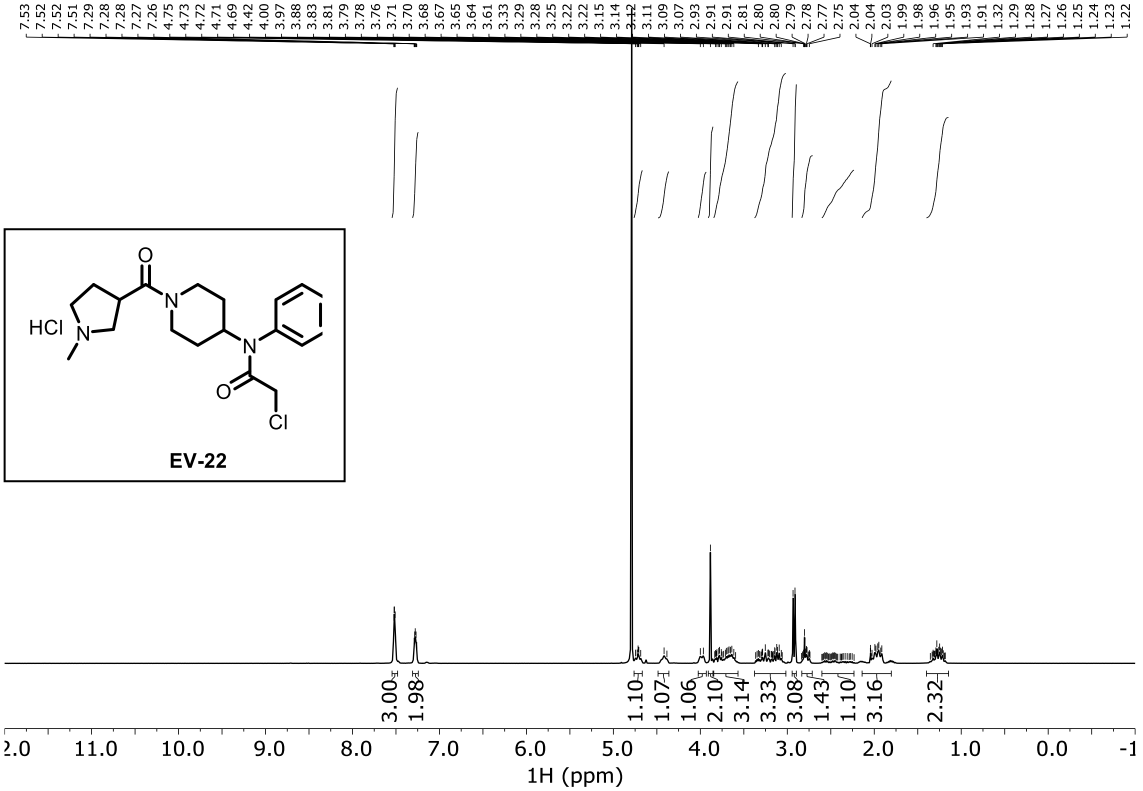

**EV-23 ¹H NMR ($CDCl_3$)**

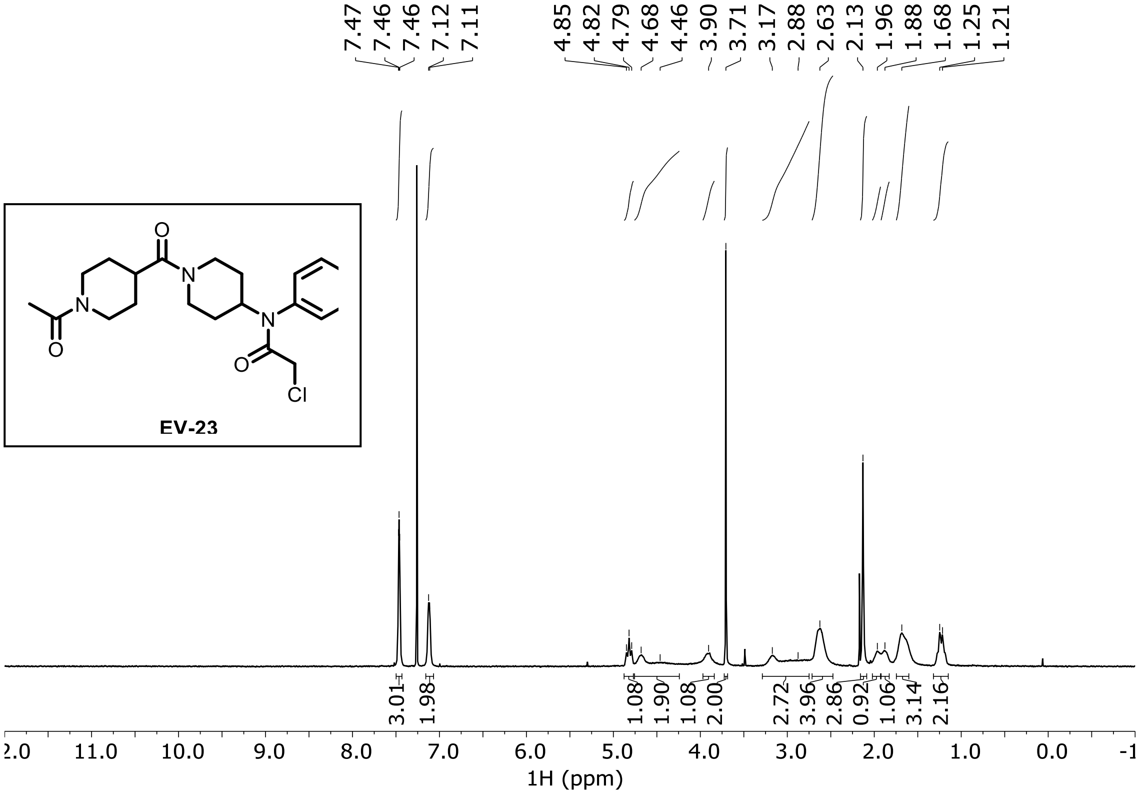

**EV-24 ¹H NMR ($CDCl_3$)**

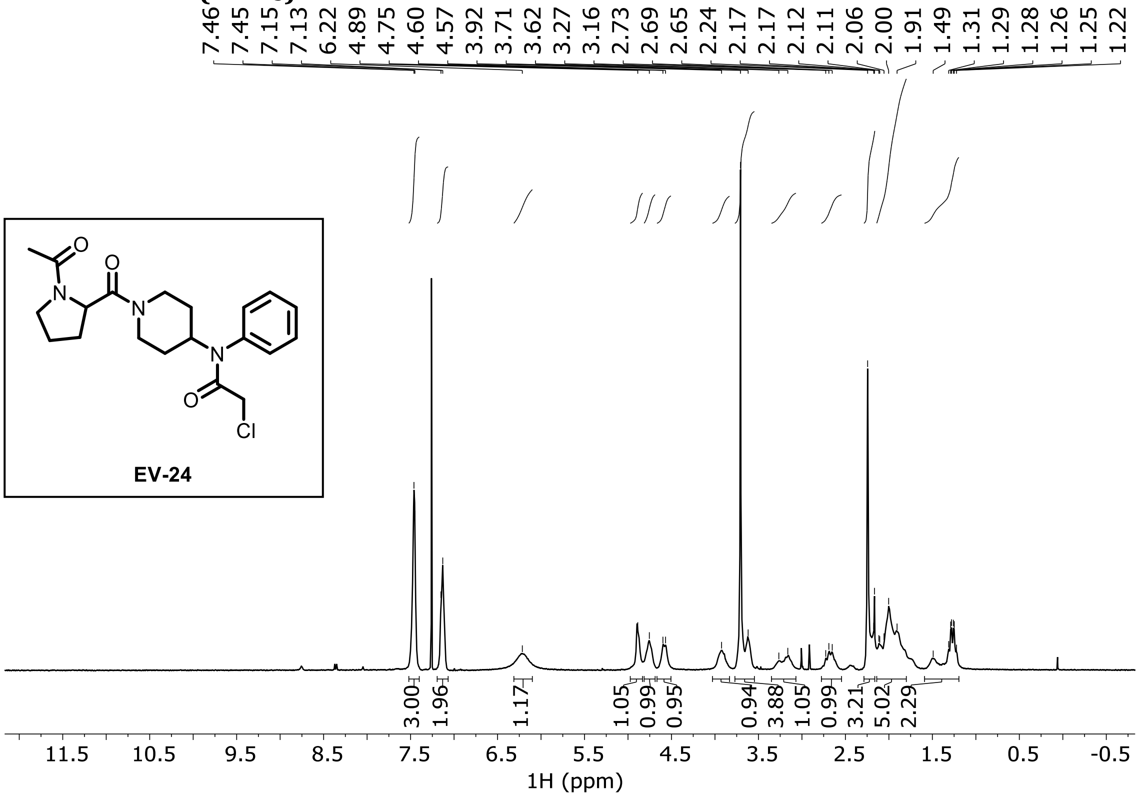

**EV-25 ¹H NMR ($CDCl_3$)**

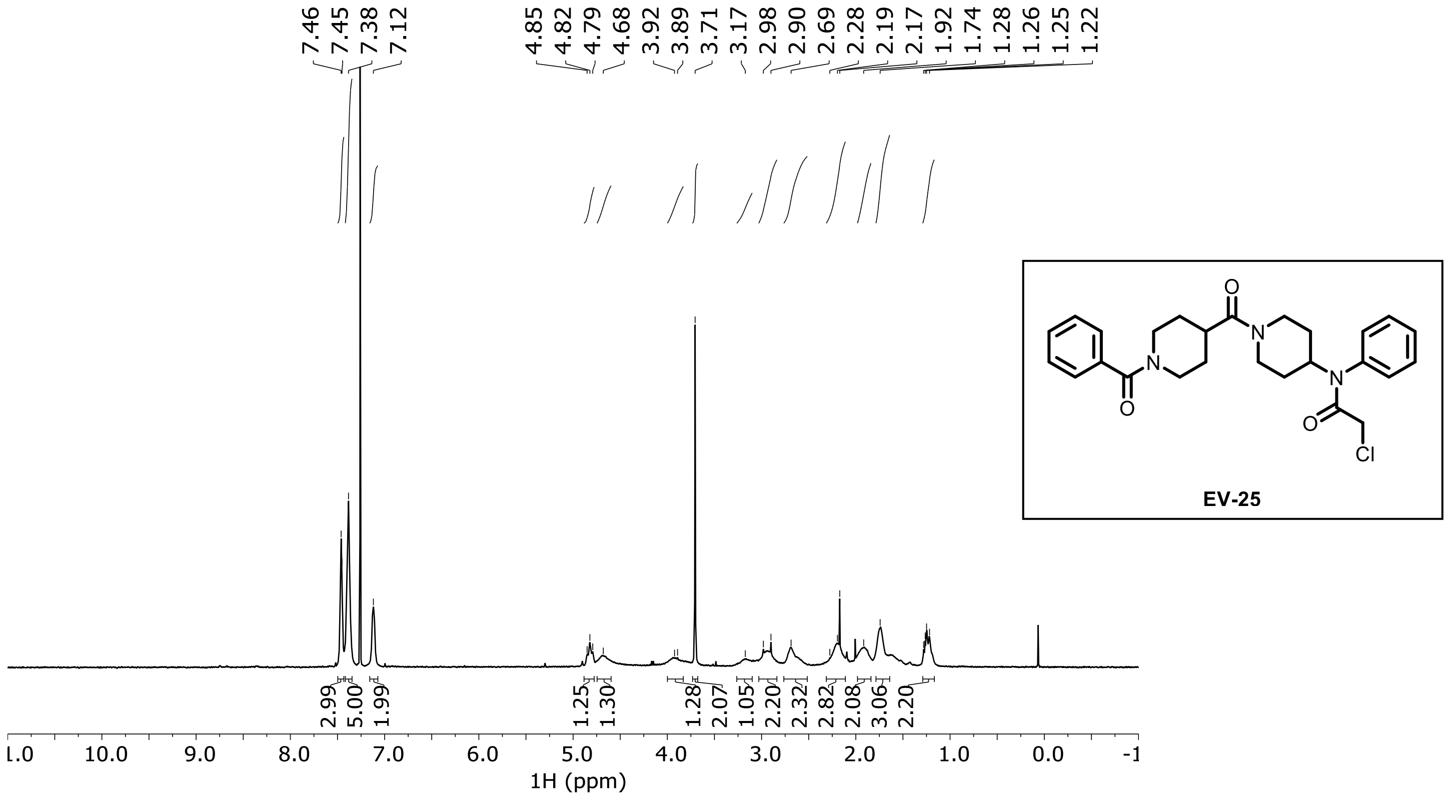

**EV-26 ${}^{1}$H NMR ($CDCl_3$)**

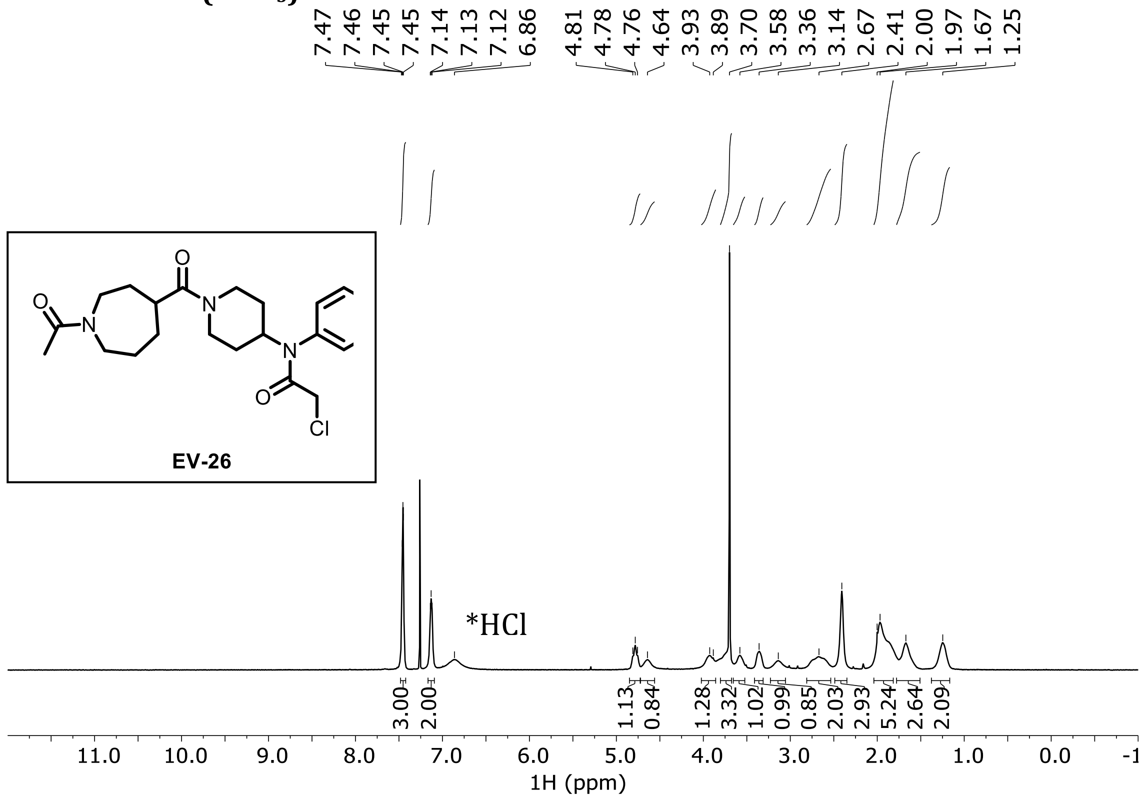

**EV-27 ${}^{1}$H NMR ($CDCl_3$)**

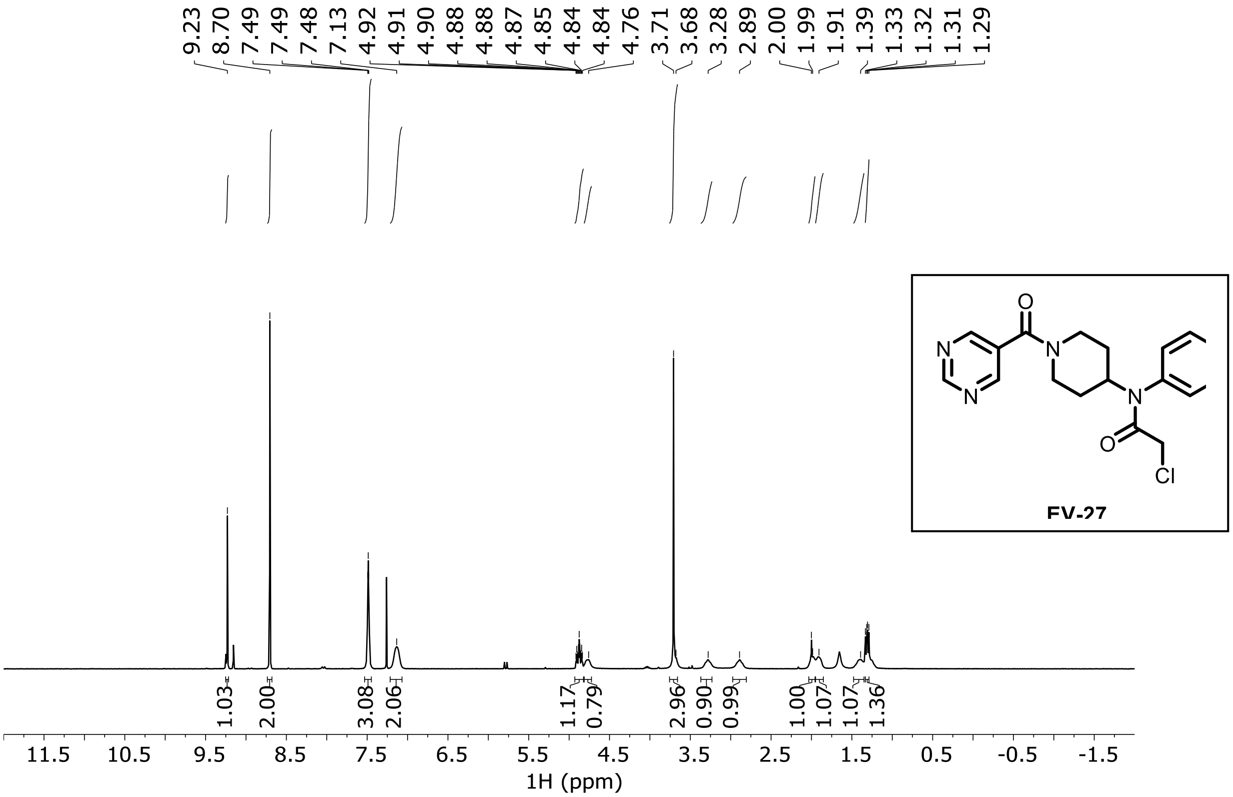

**EV-28 ¹H NMR ($CDCl_3$)**

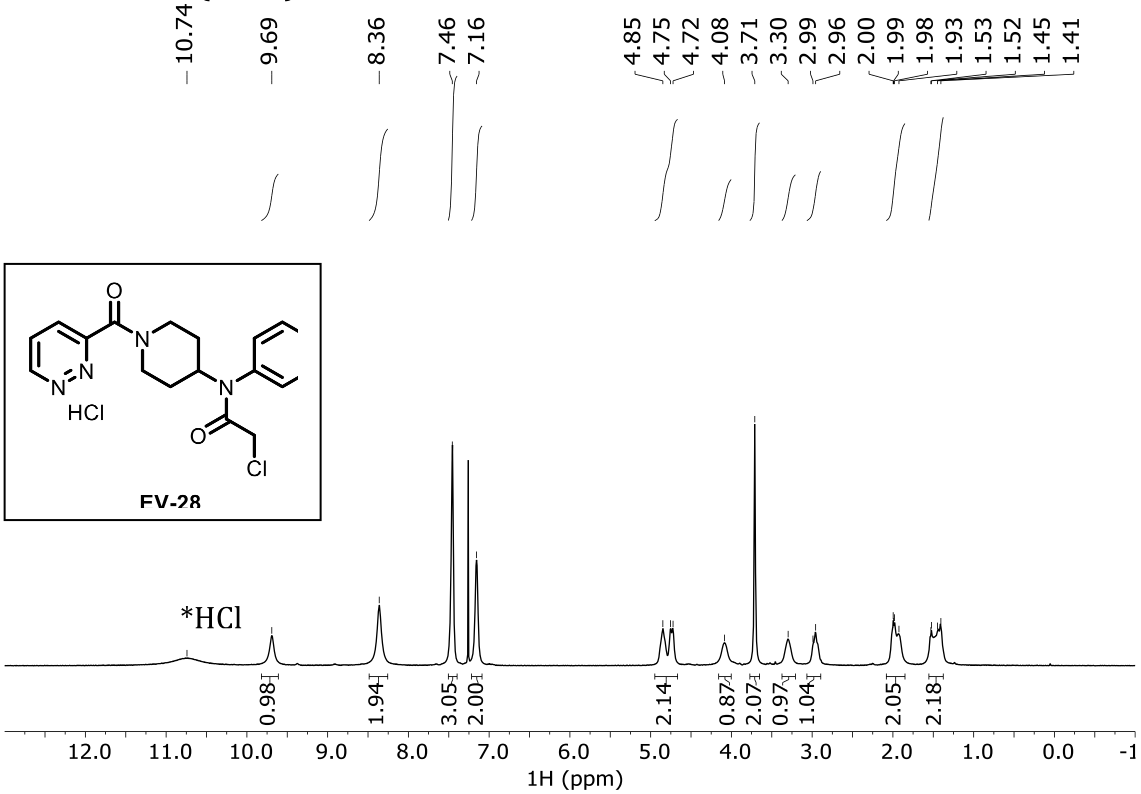

**EV-29 ¹H NMR ($CDCl_3$)**

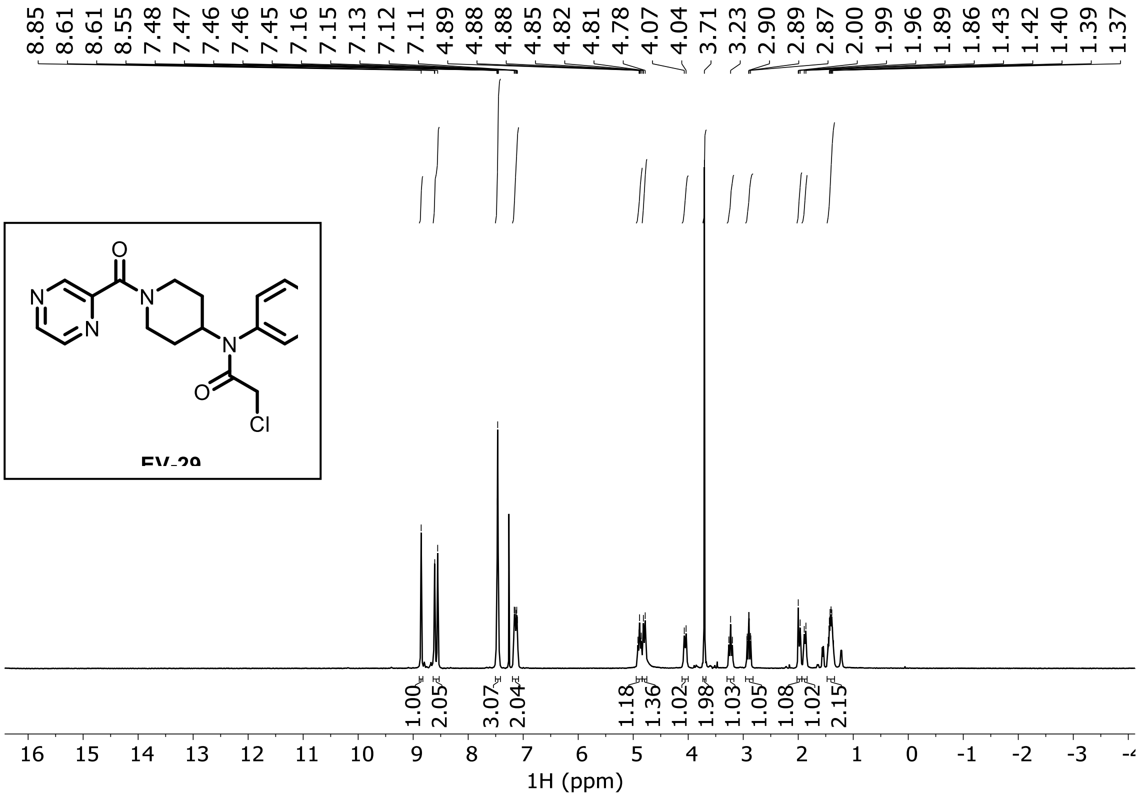

**EV-30 $^{1}$H NMR ($CDCl_3$)**

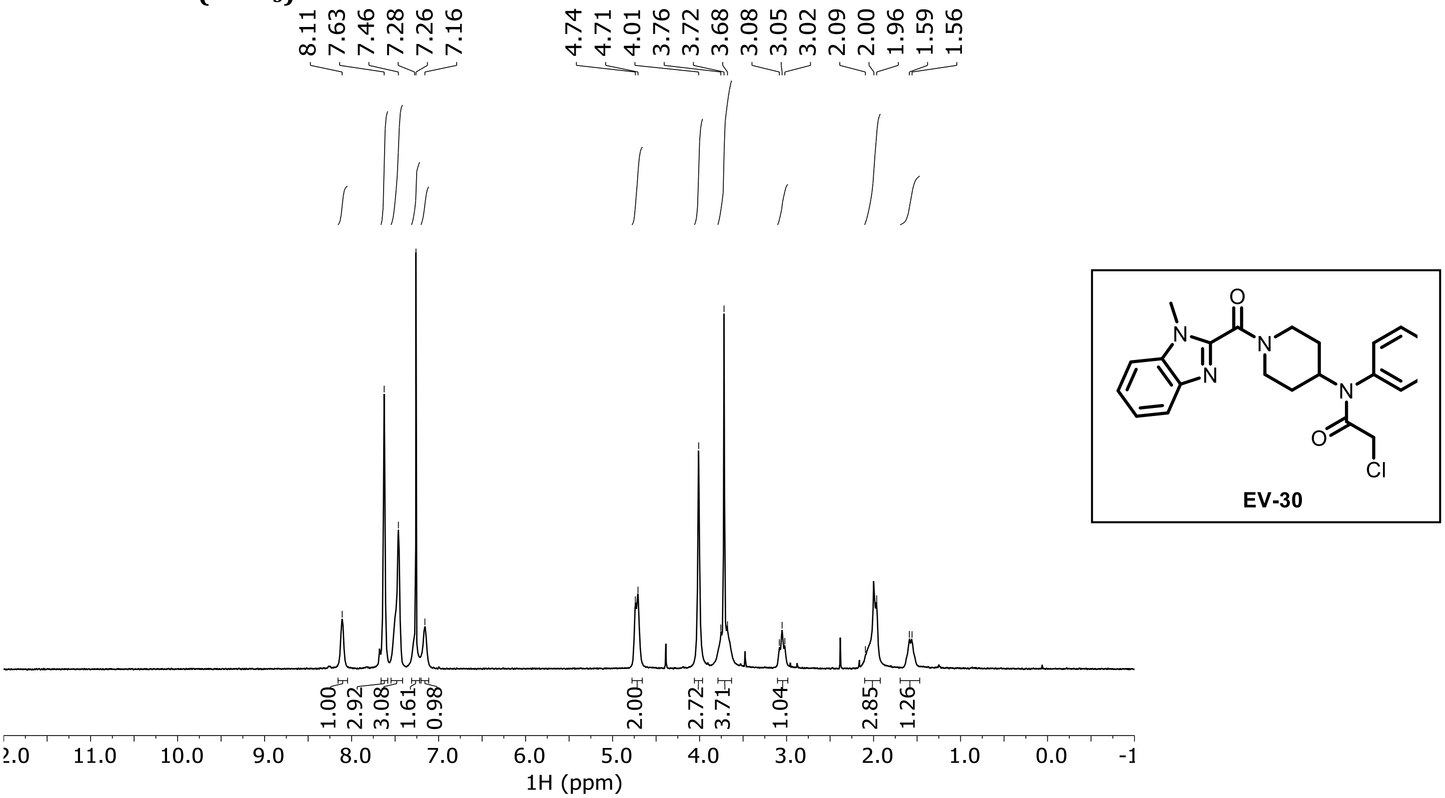

**EV-31 $^{1}$H NMR ($CD_3CN$)**

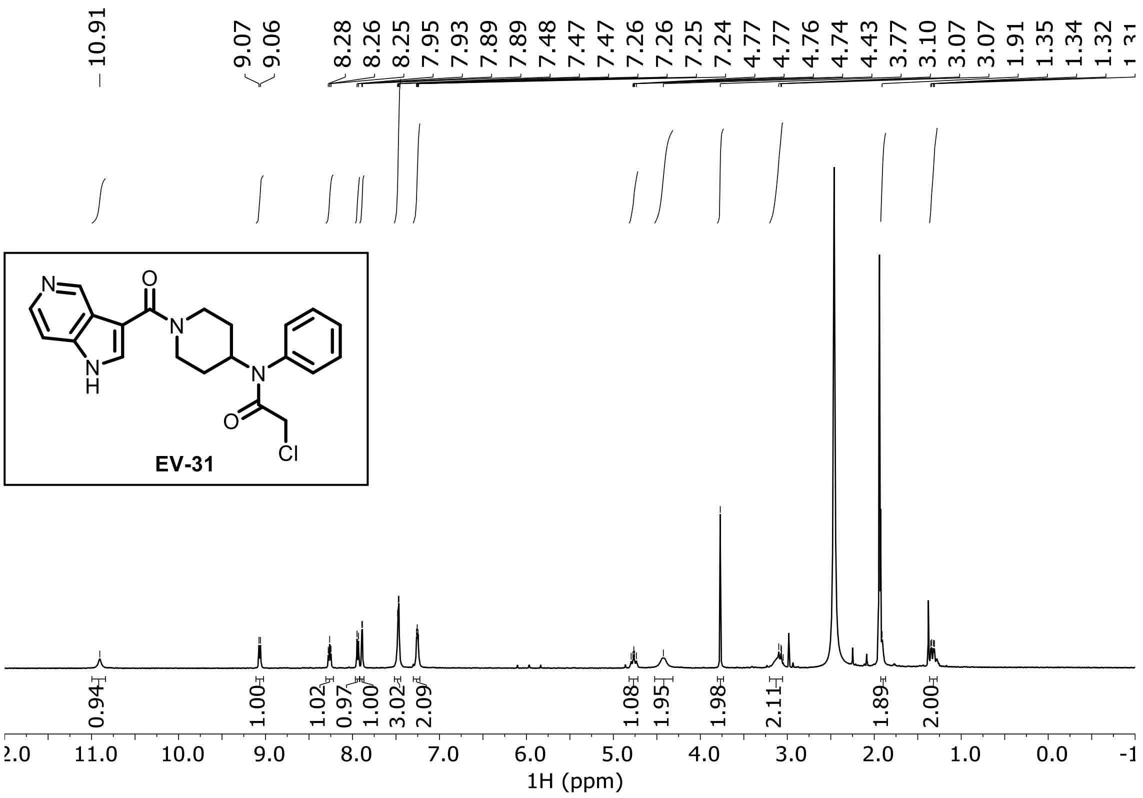

**EV-32 $^1$H NMR ($CDCl_3$)**

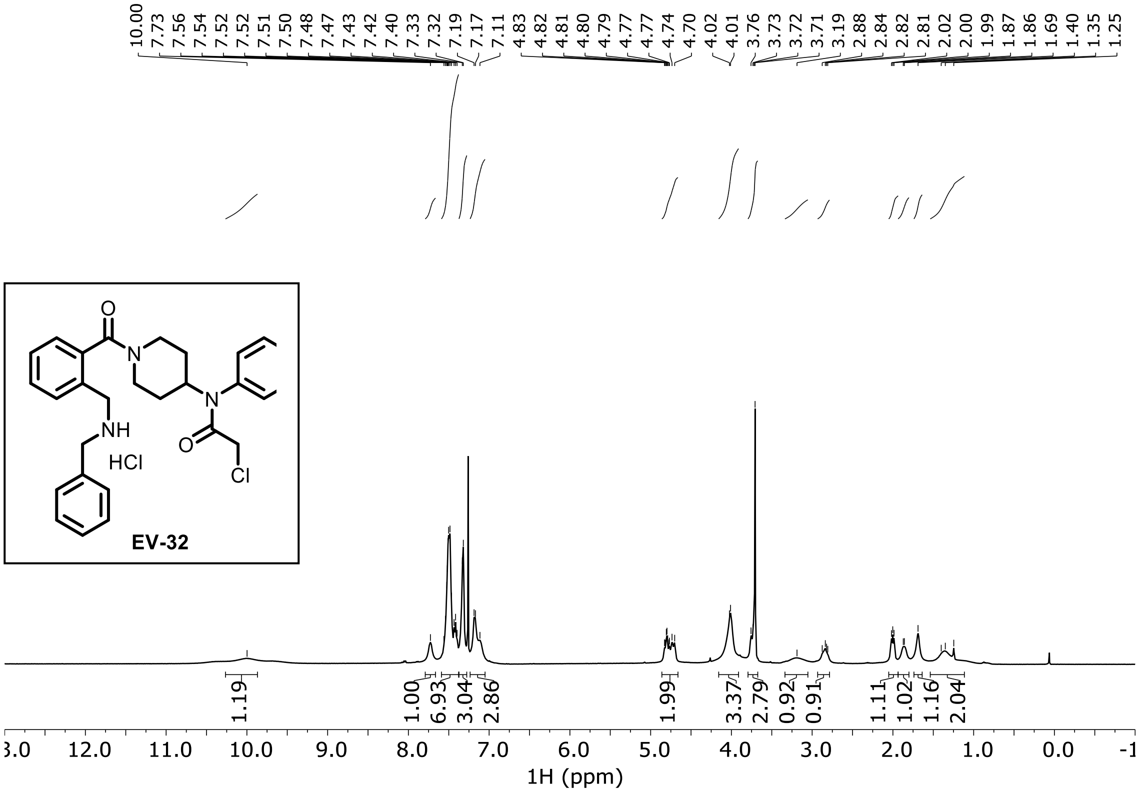

**EV-33 $^1$H NMR ($CDCl_3$)**

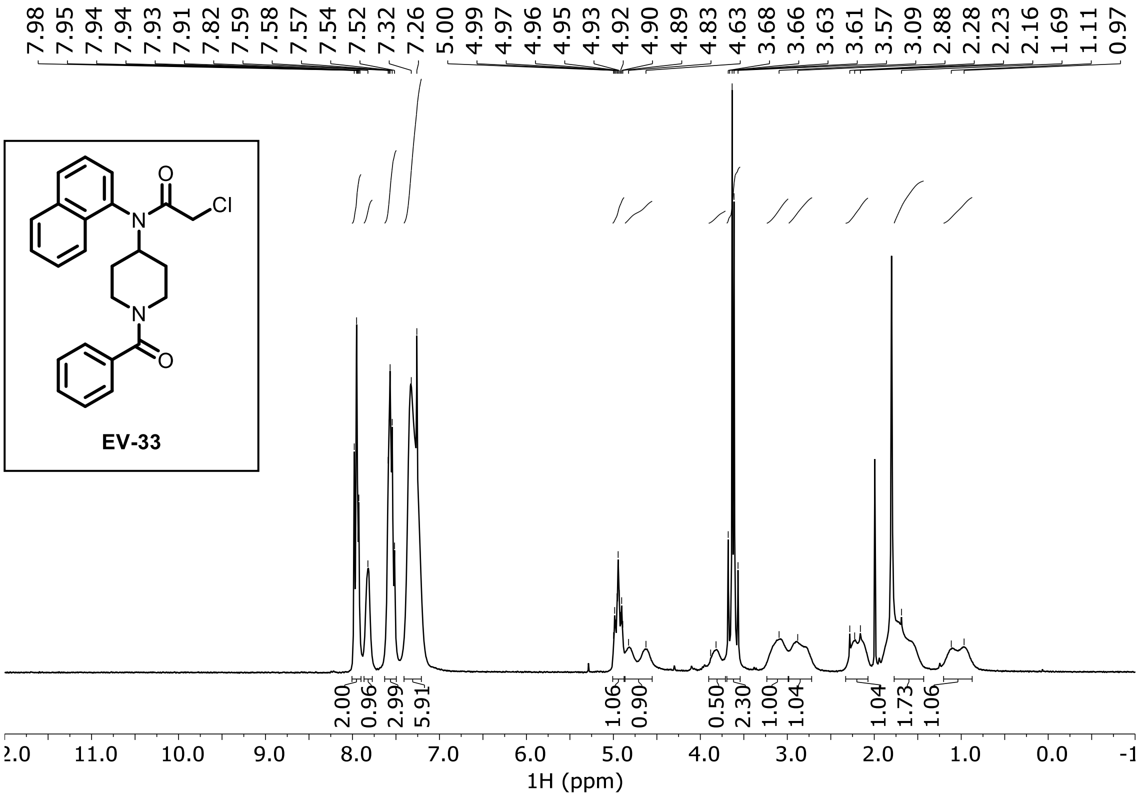

**EV-34 $^{1}$H NMR ($CDCl_3$)**

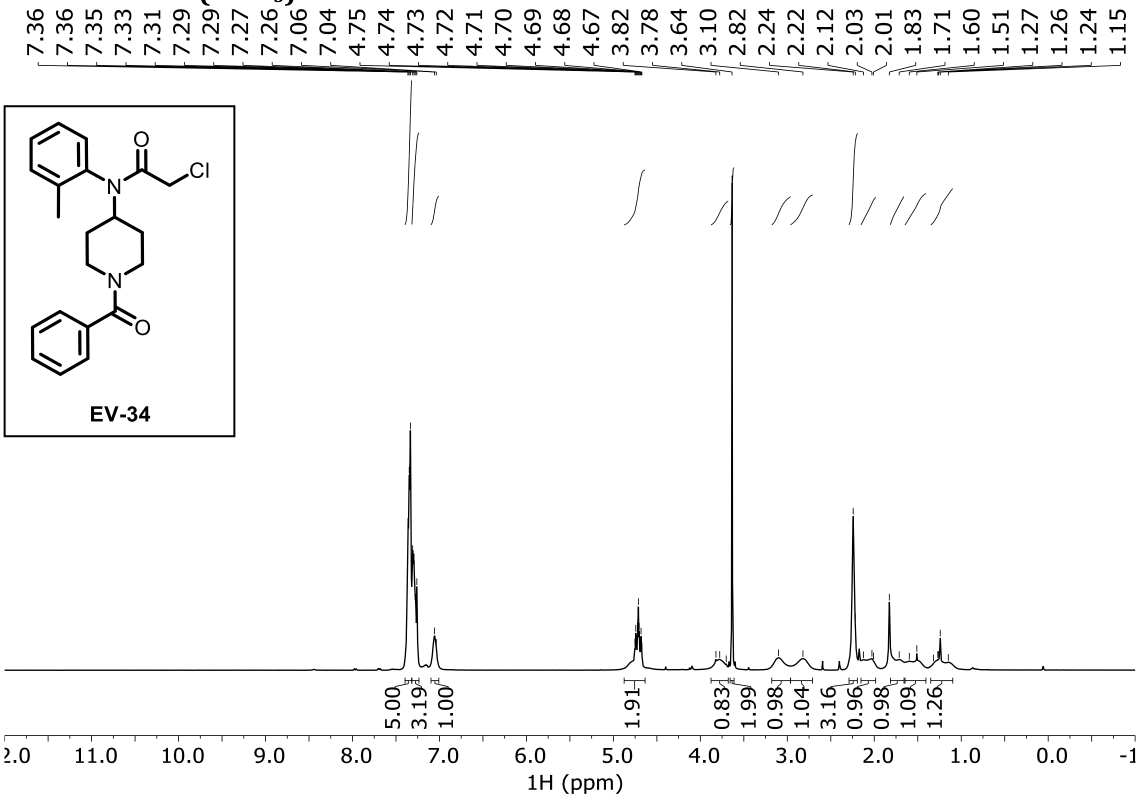

**EV-35 $^{1}$H NMR ($CDCl_3$)**

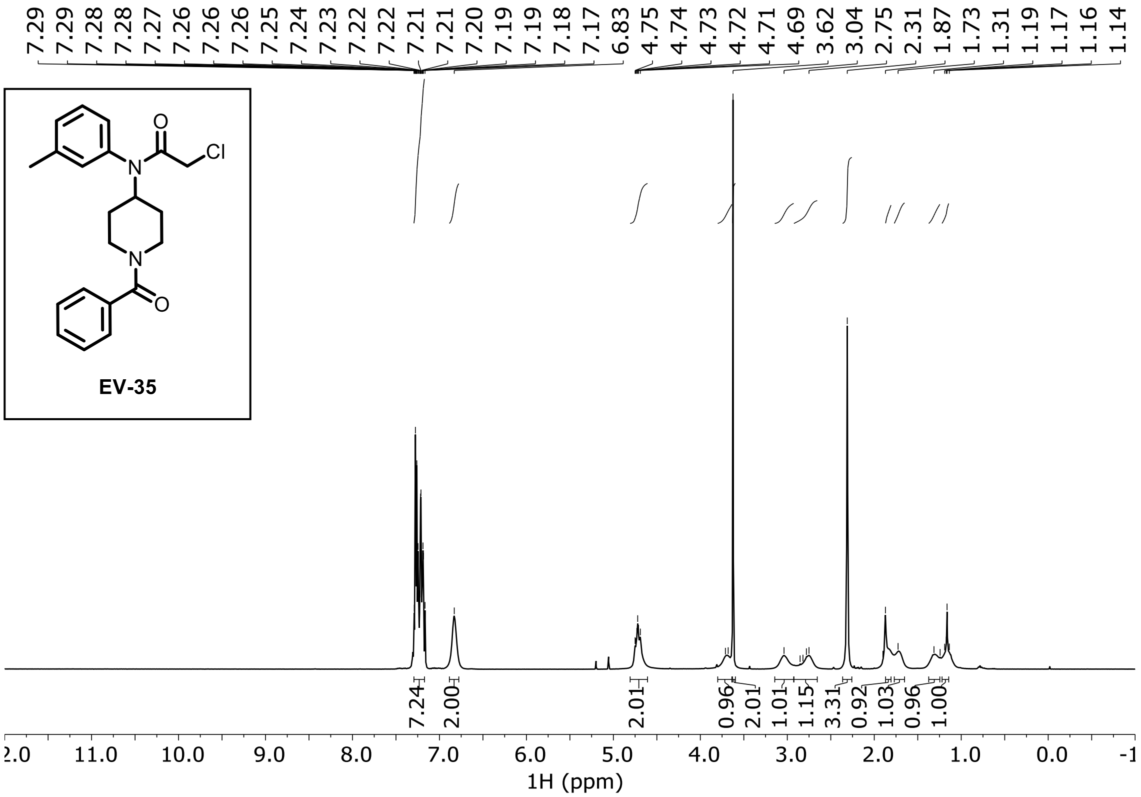

**EV-36 $^{1}$H NMR ($CDCl_3$)**

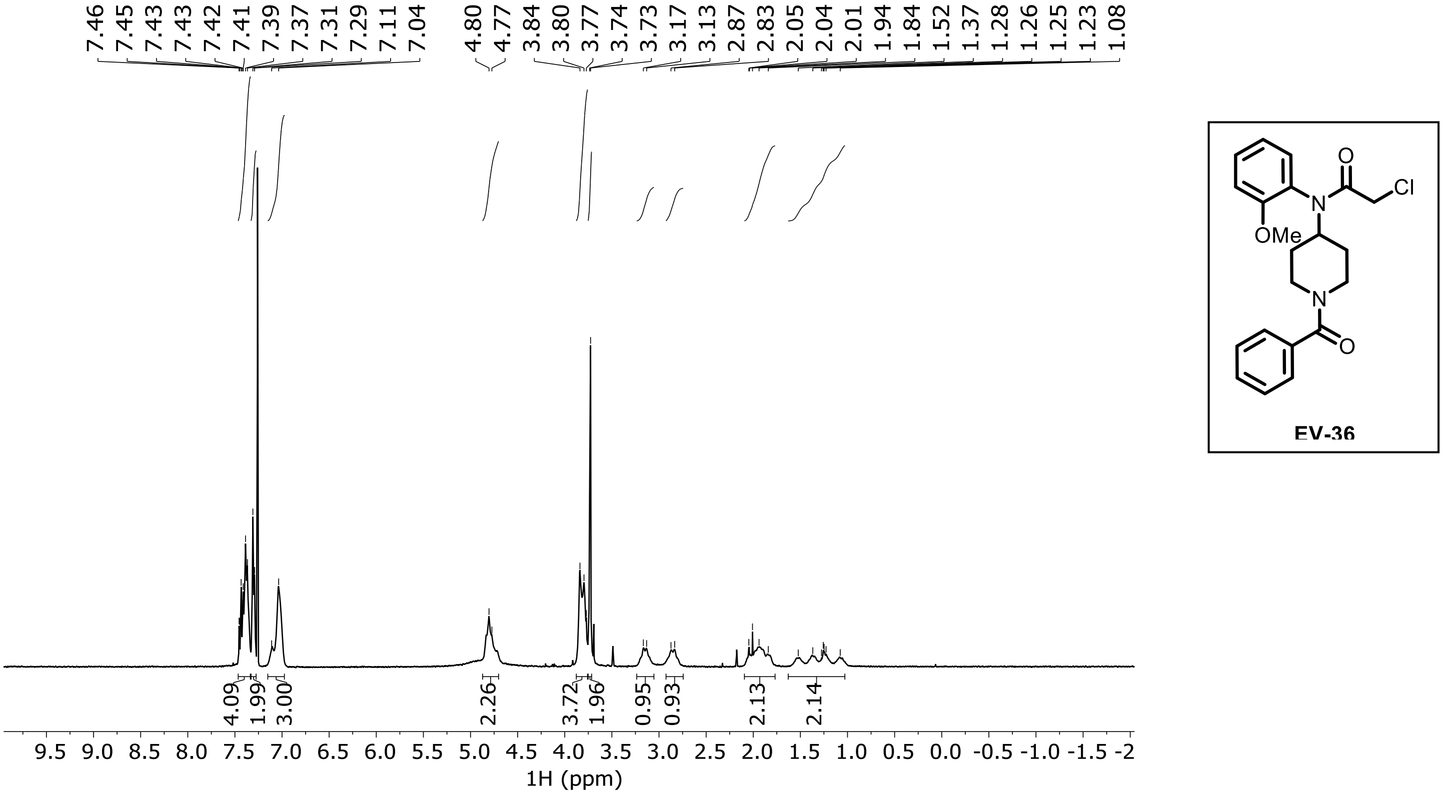

**EV-37 $^{1}$H NMR ($CDCl_3$)**

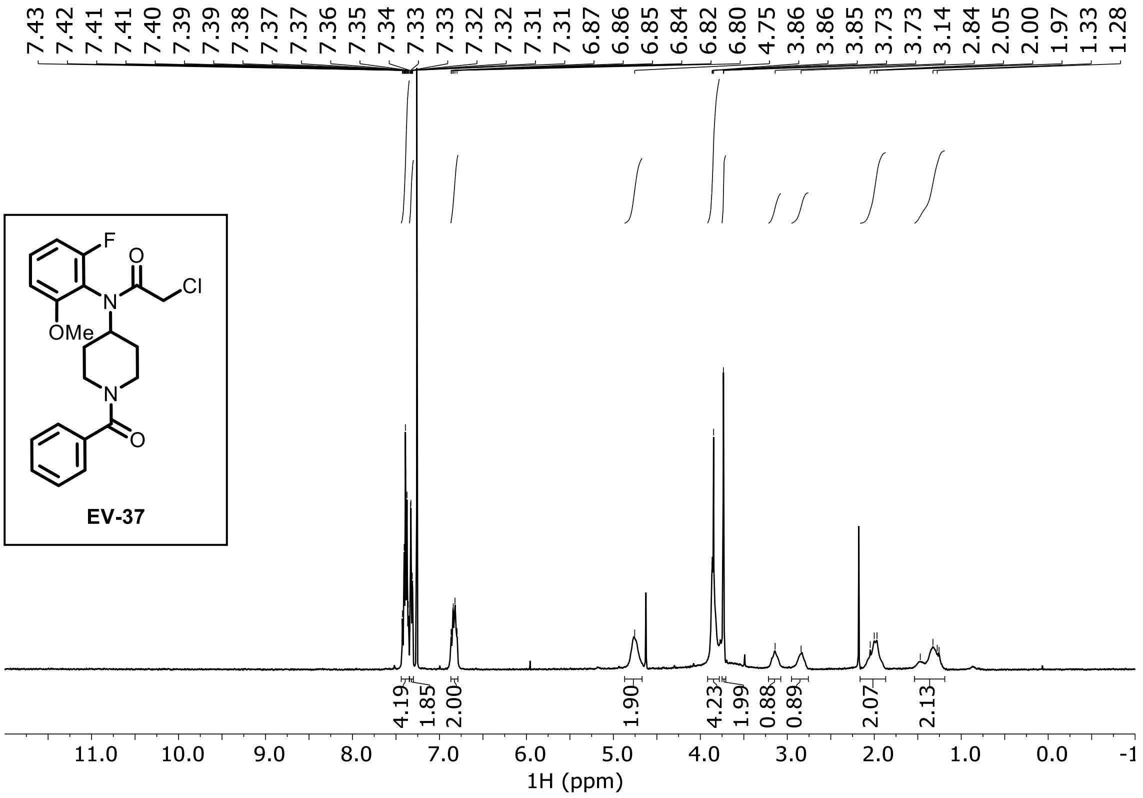

**EV-38 $^1$H NMR (MeOD)**

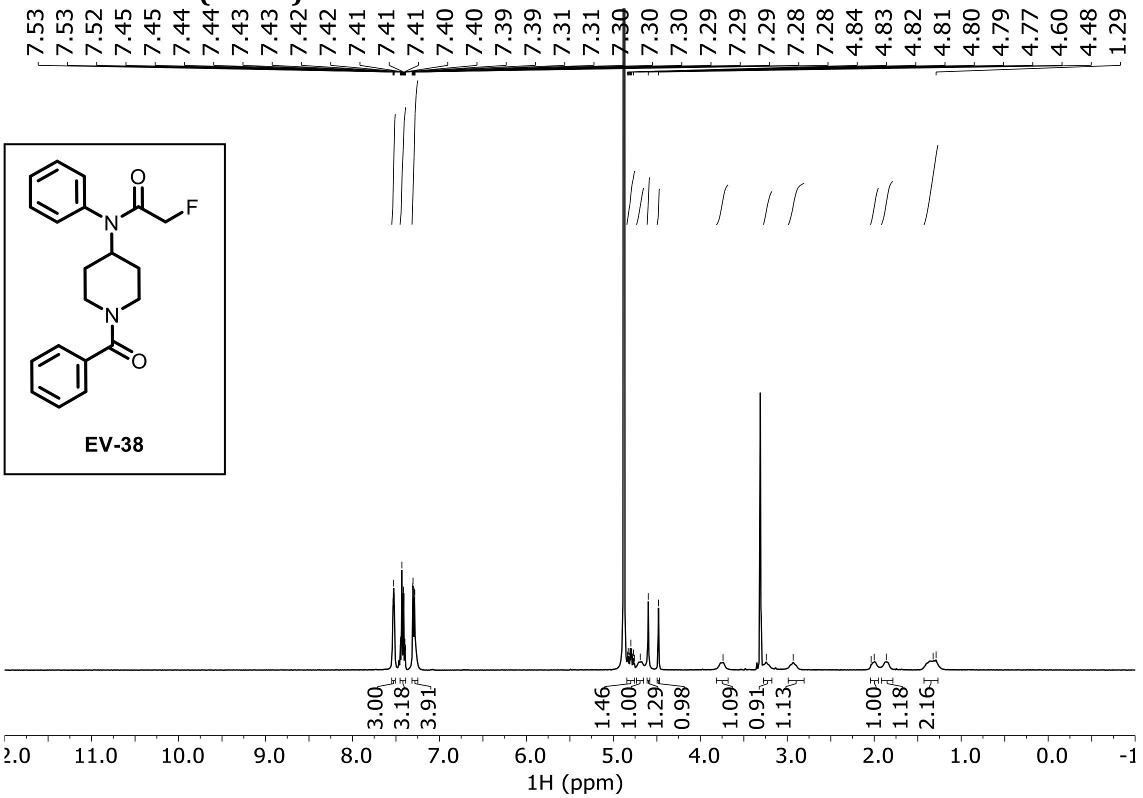

**EV-39 $^1$H NMR (MeOD)**

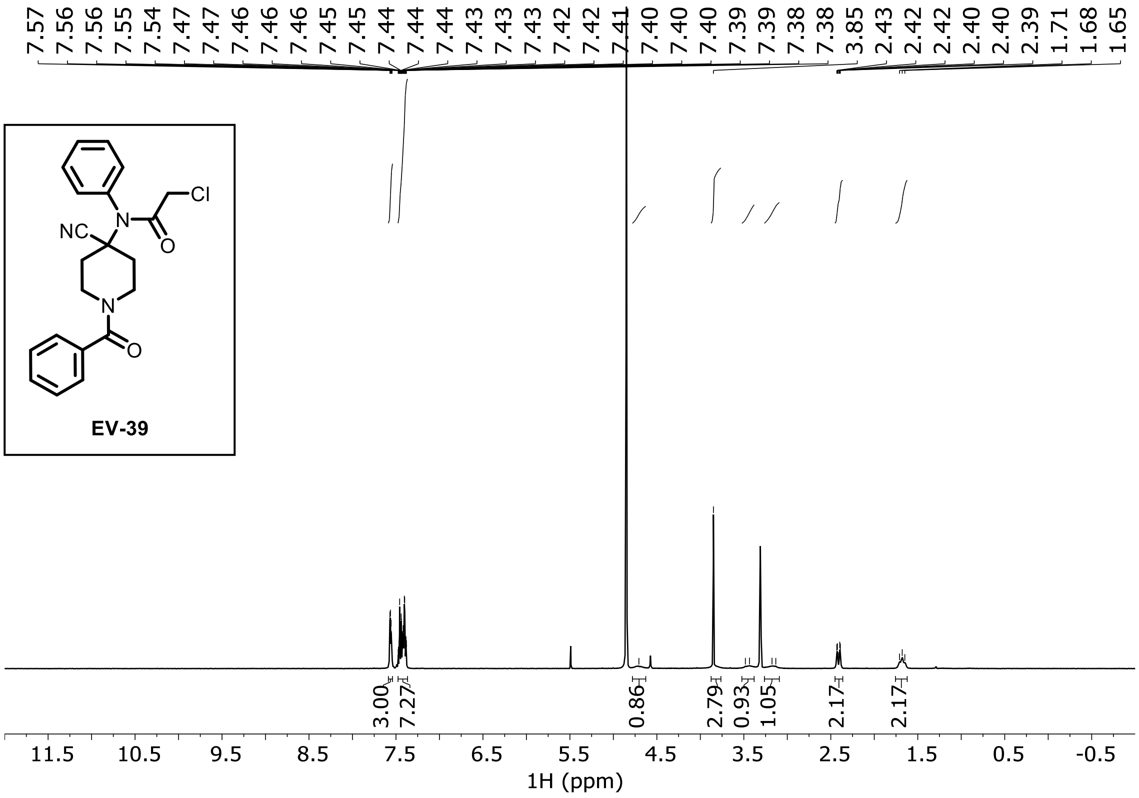

**EV-40 ¹H NMR ($CDCl_3$)**

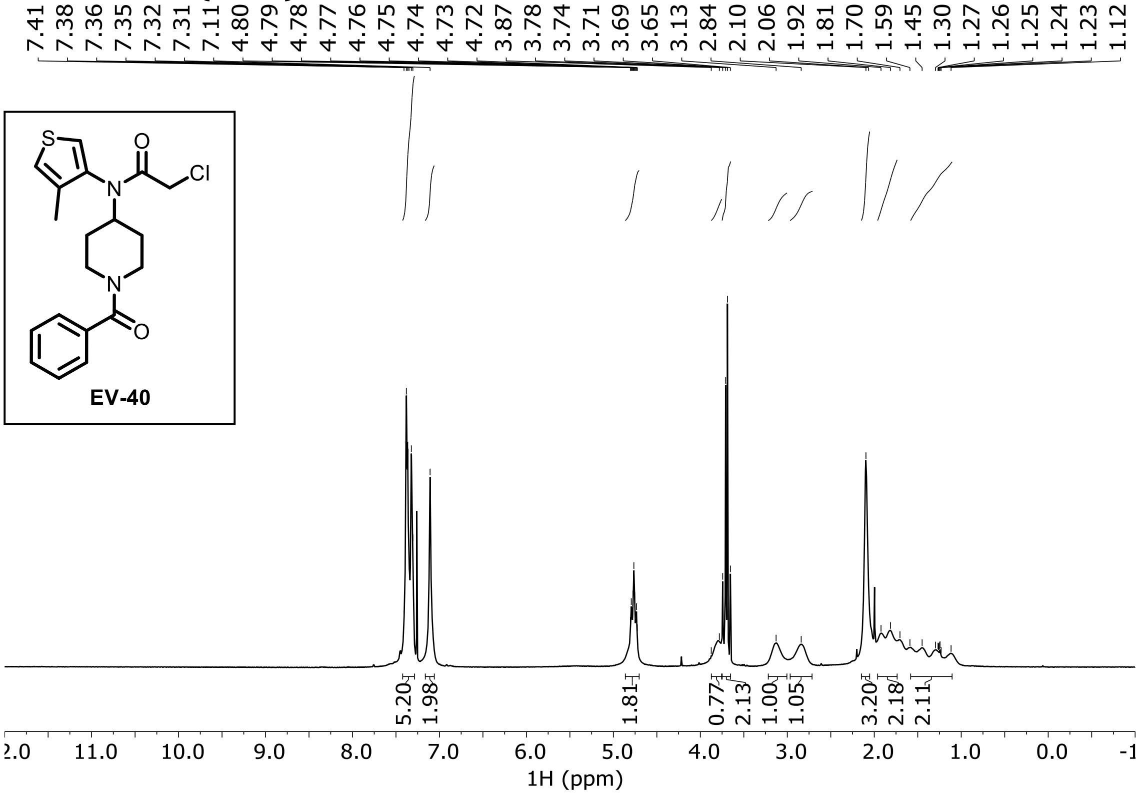

**EV-41 ¹H NMR (DMSO-$d_6$)**

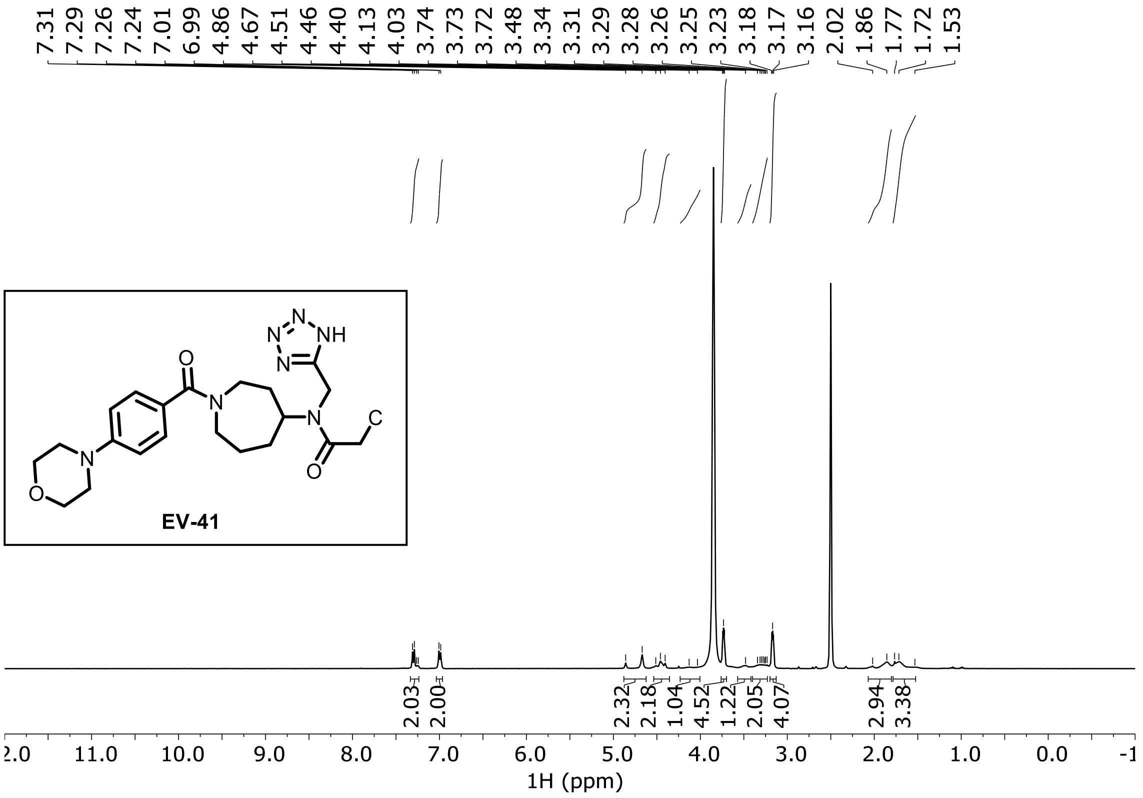

**EV-42 ¹H NMR (DMSO-$d_6$)**

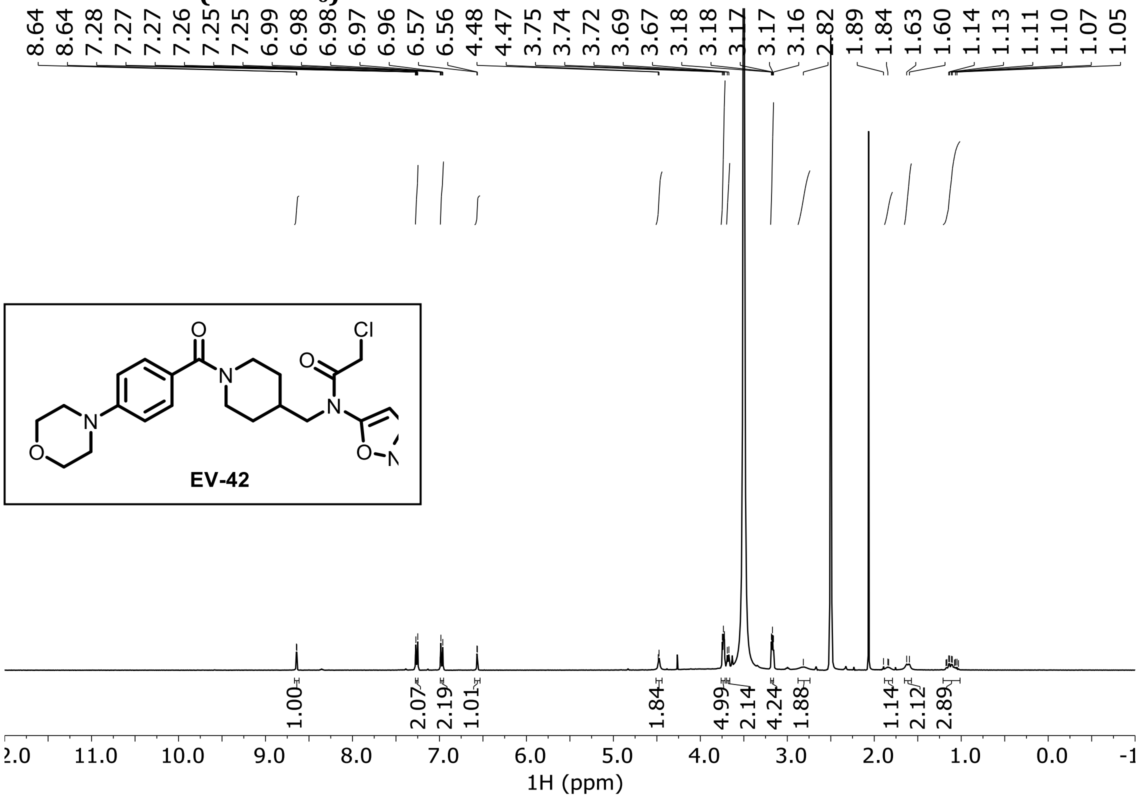

**EV-43 ¹H NMR (DMSO-$d_6$)**

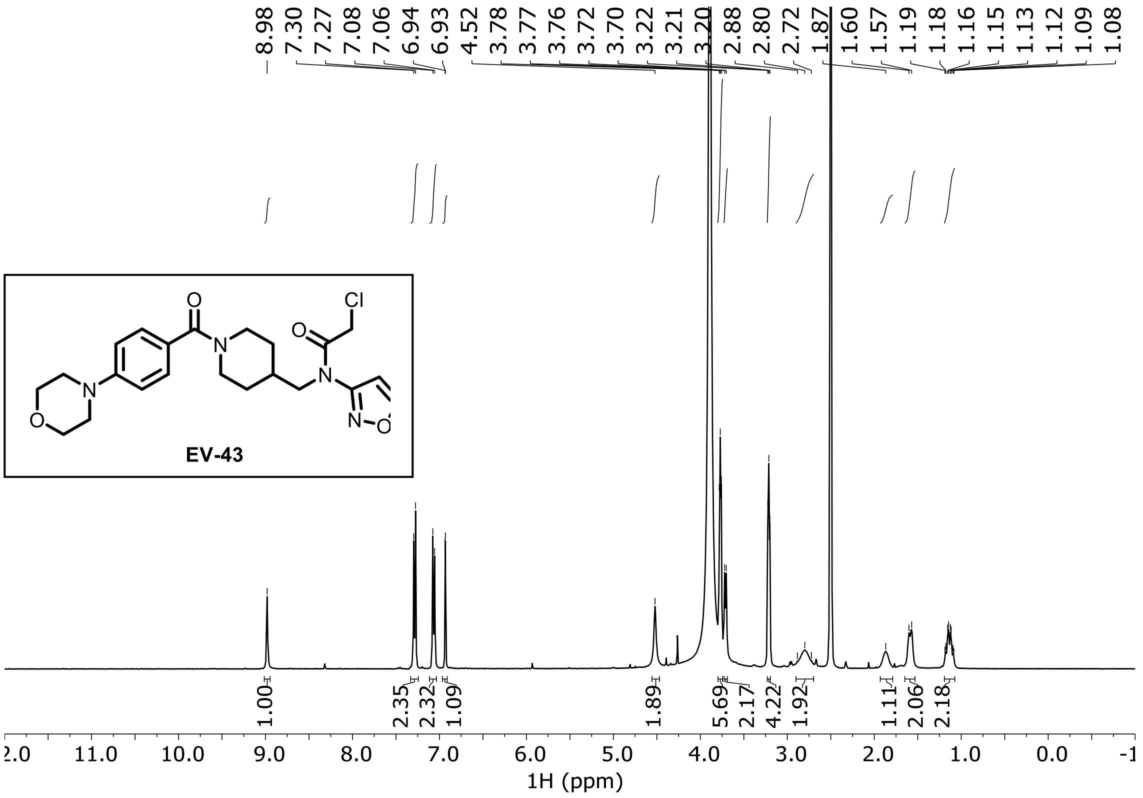

**EV-44 ¹H NMR (DMSO-$d_6$)**

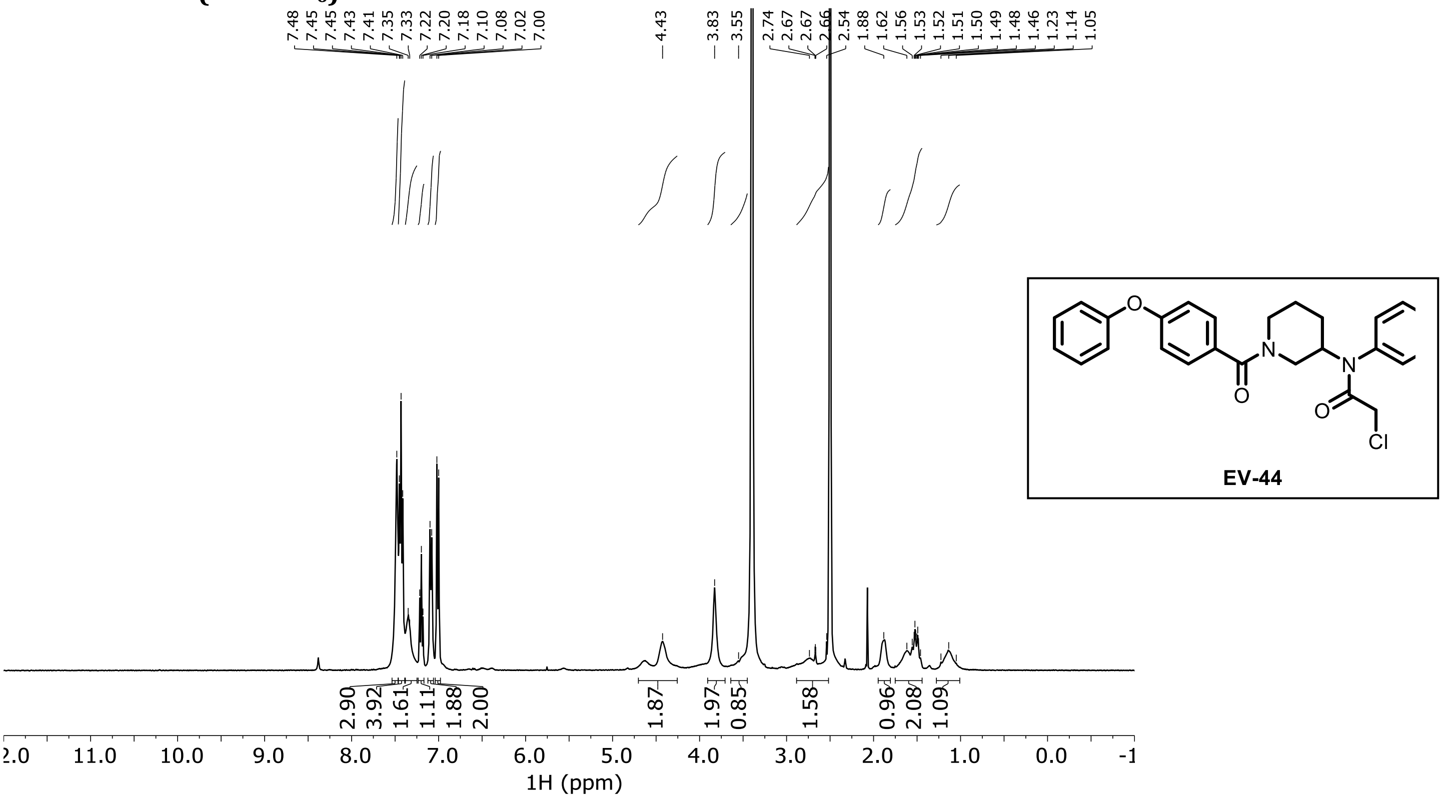

**EV-45 ¹H NMR (DMSO-$d_6$)**

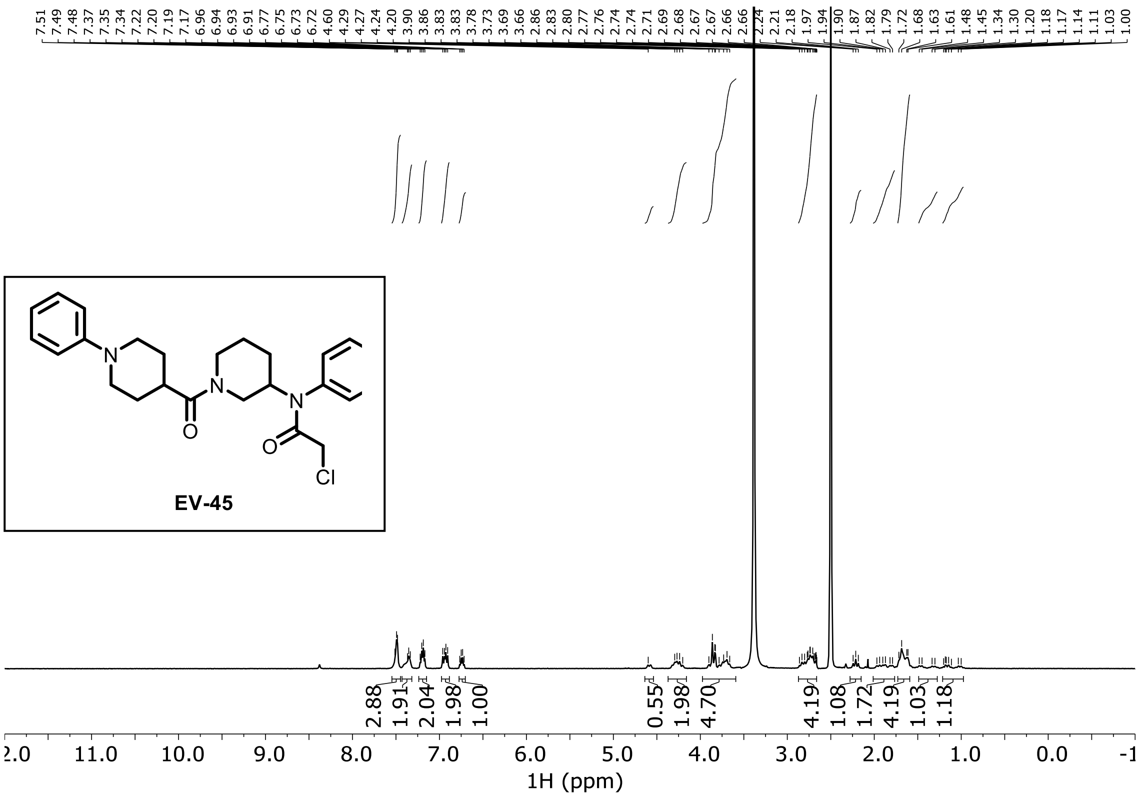

**EV-46 ¹H NMR ($CDCl_3$)**

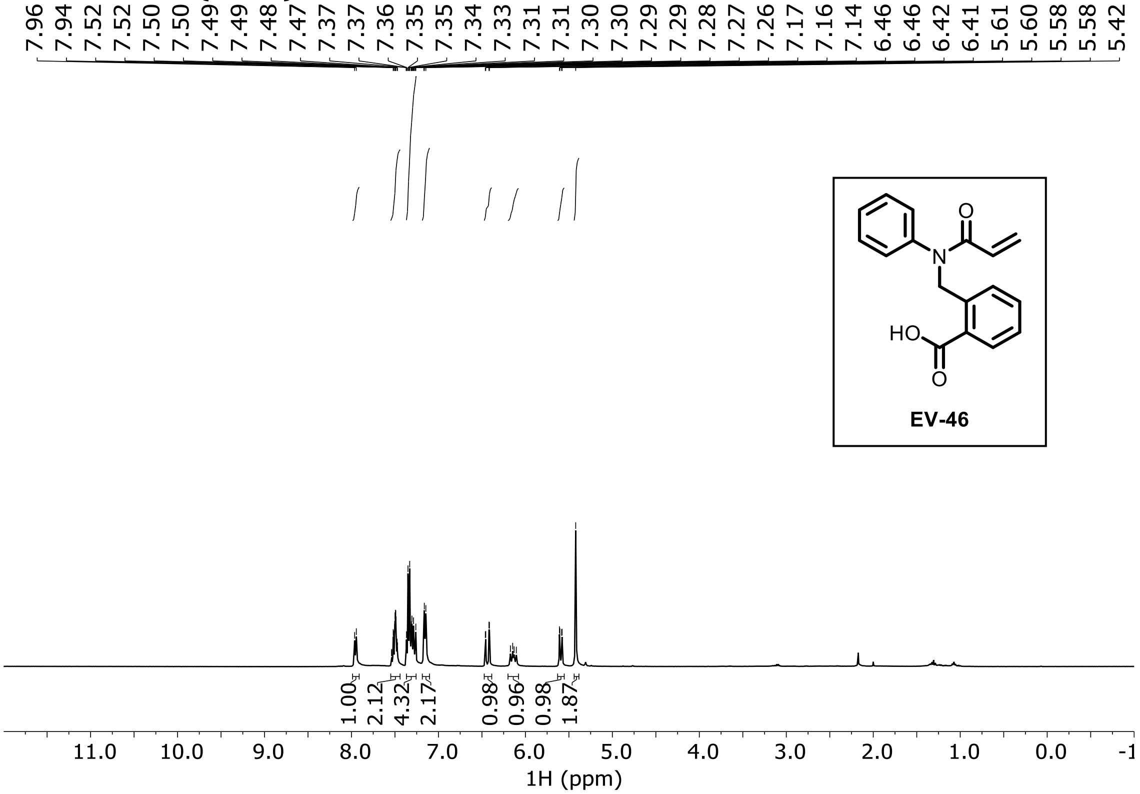

**EV-47 ¹H NMR ($CDCl_3$)**

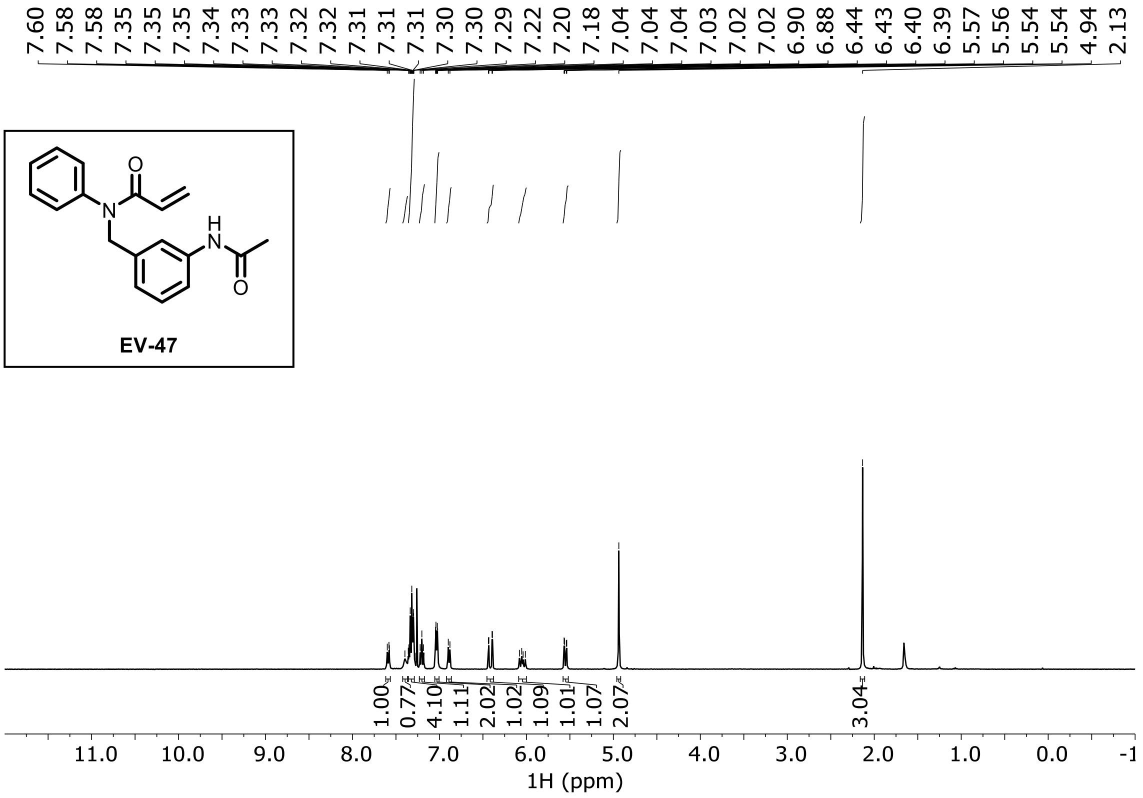

**EV-48 $^{1}$H NMR ($CDCl_3$)**

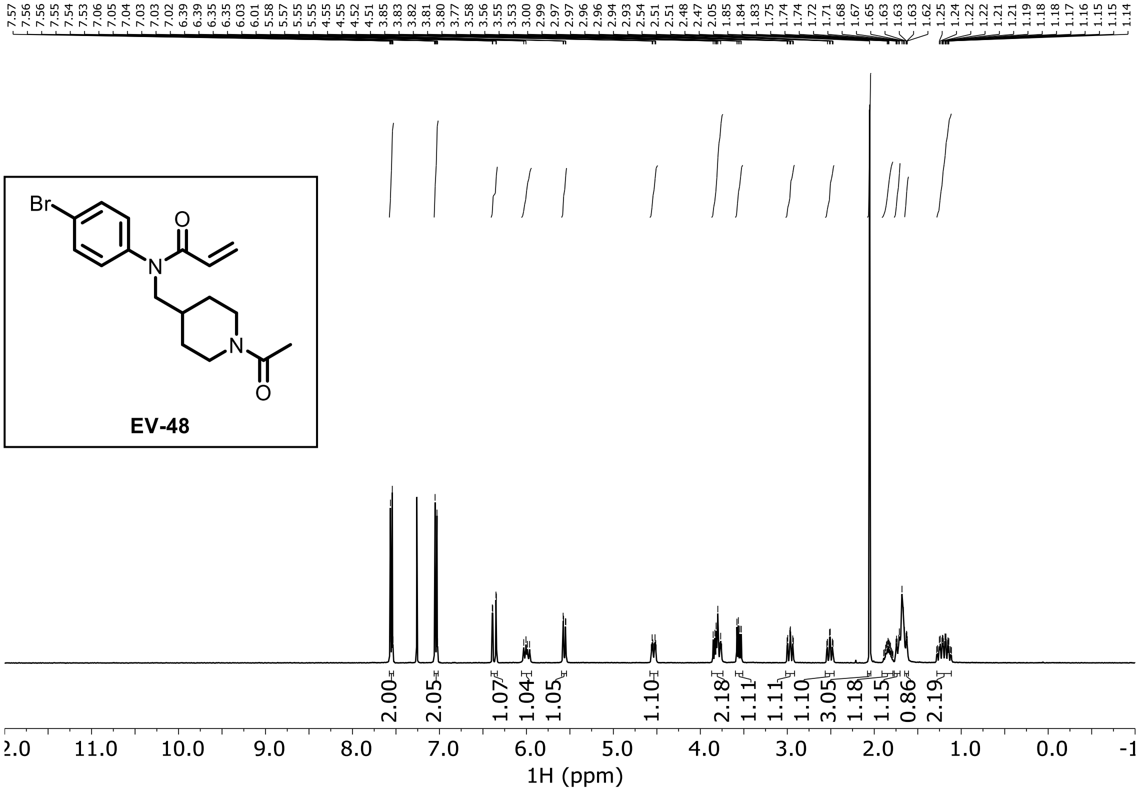

**EV-49 $^{1}$H NMR (DMSO-$d_6$)**

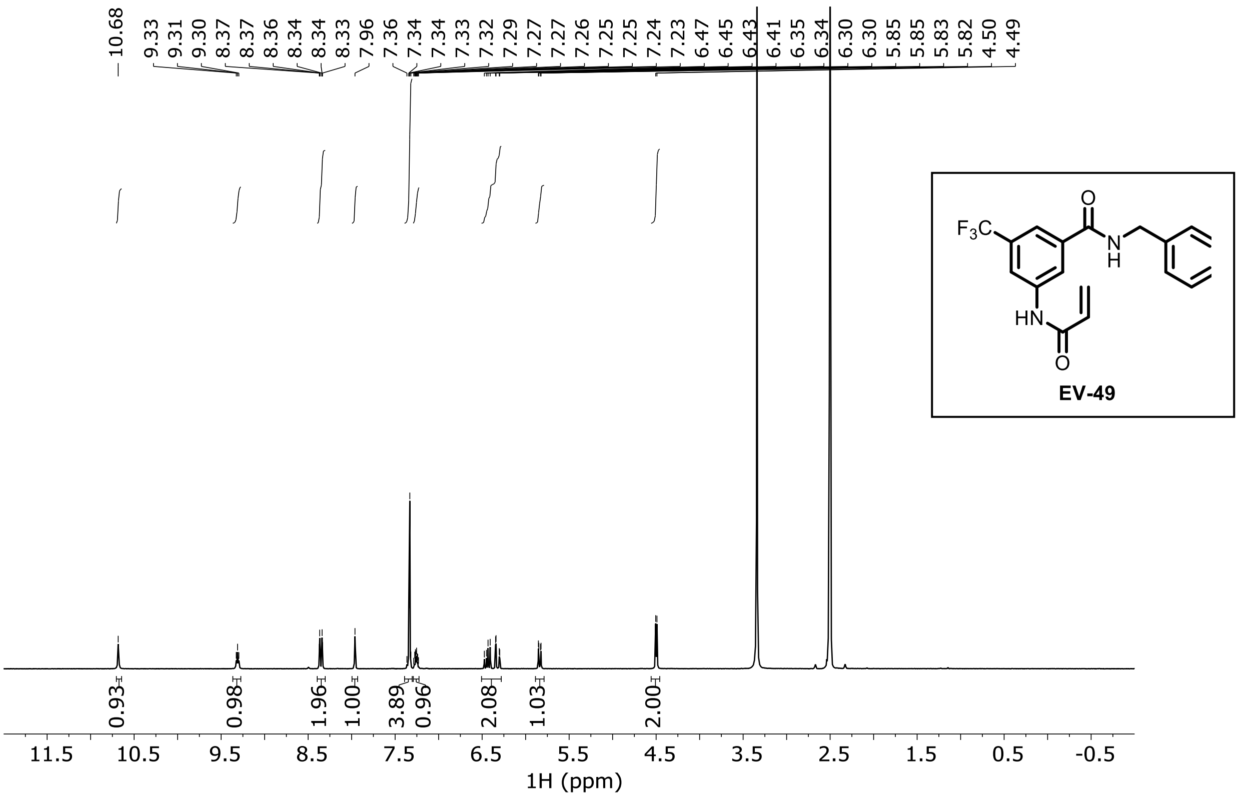

**EV-50 1H NMR (DMSO-$d_6$)**

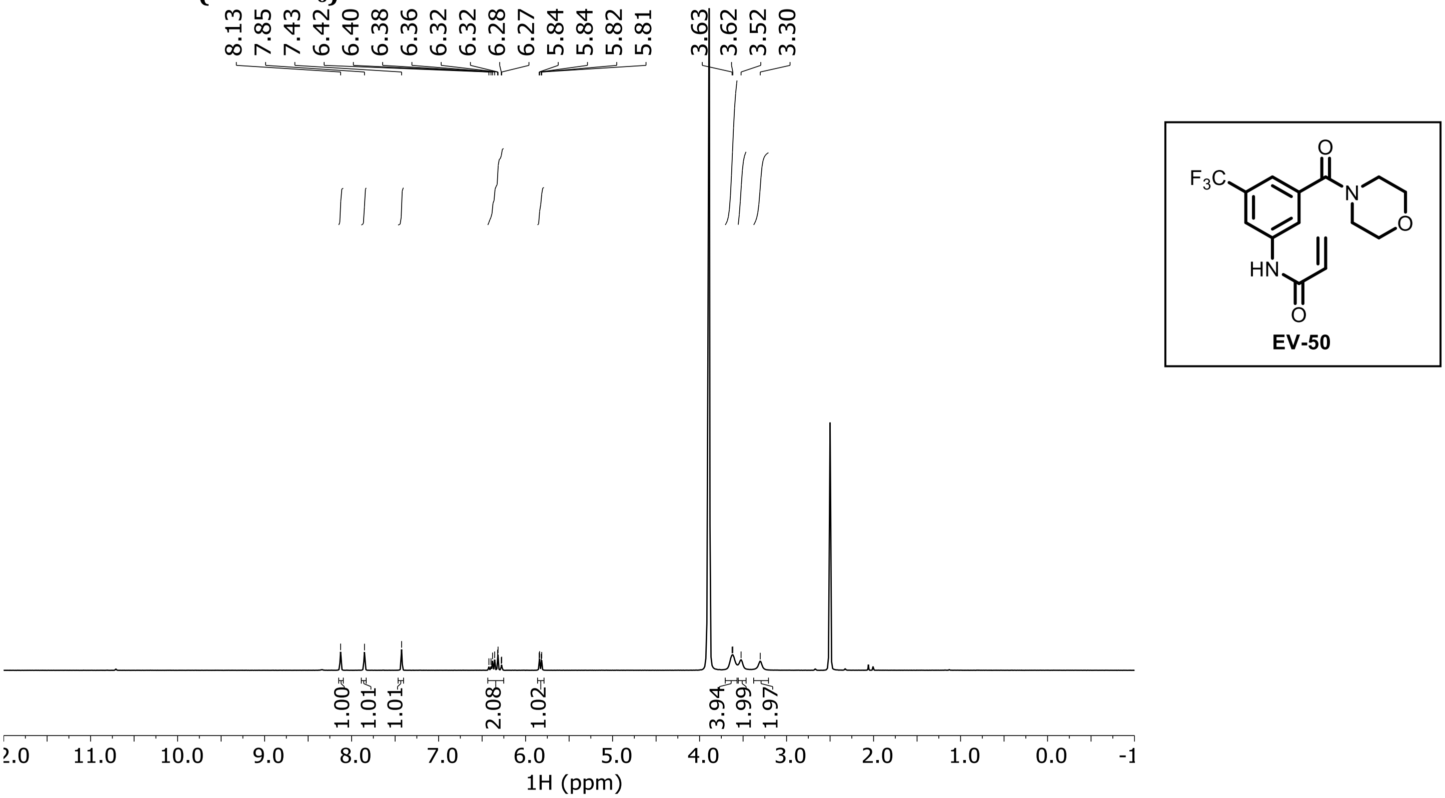

**EV-51 1H NMR (DMSO-$d_6$)**

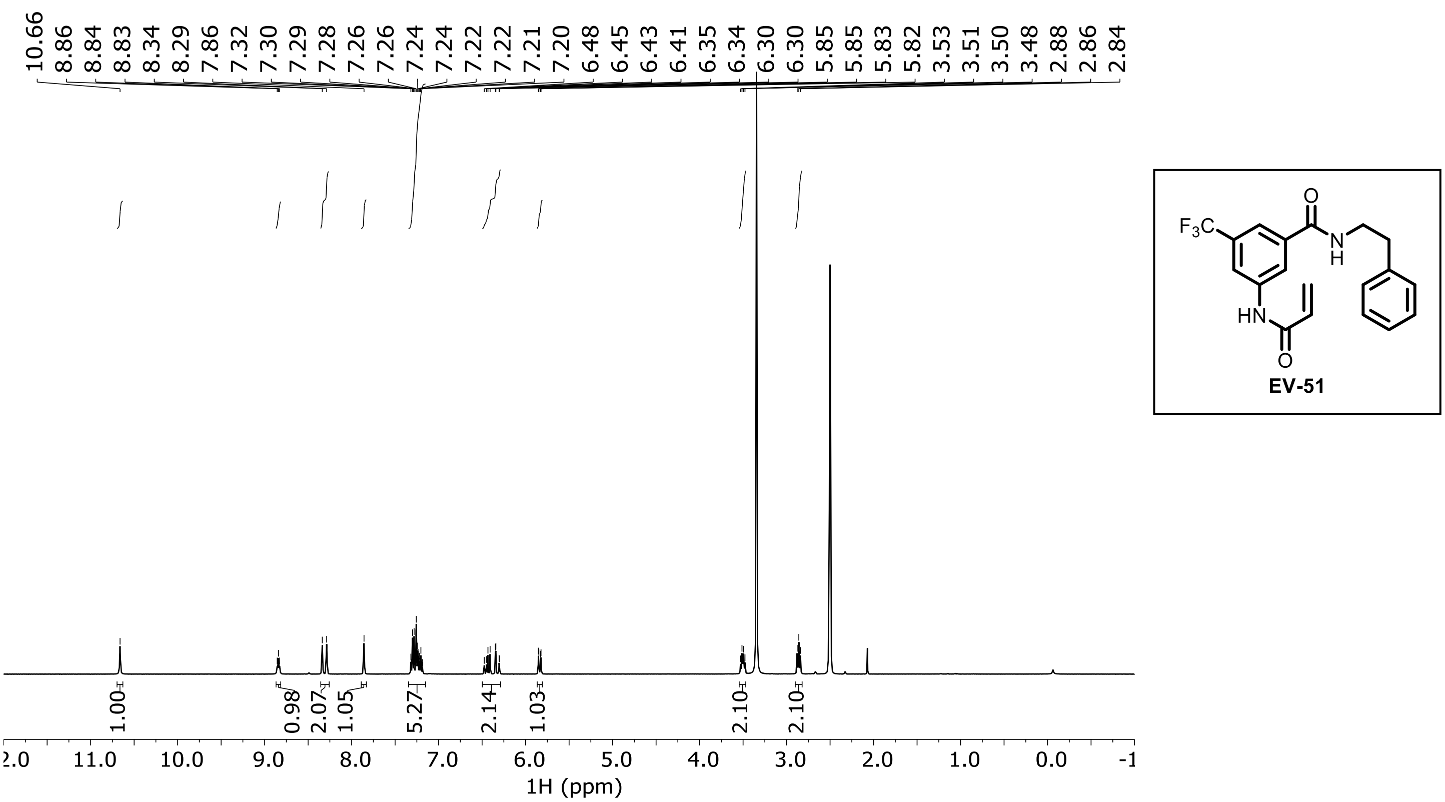

**EV-52 $^1$H NMR (DMSO-$d_6$)**

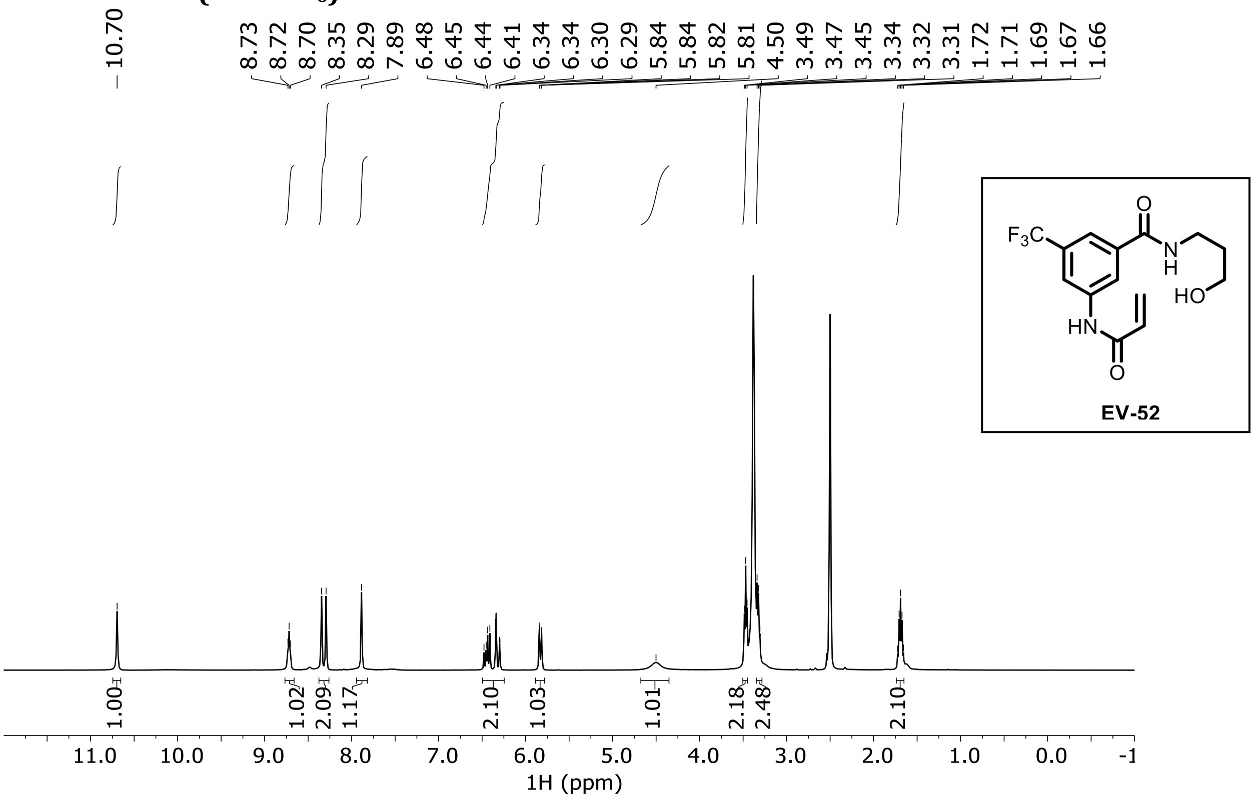

**EV-53 $^1$H NMR (DMSO-$d_6$)**

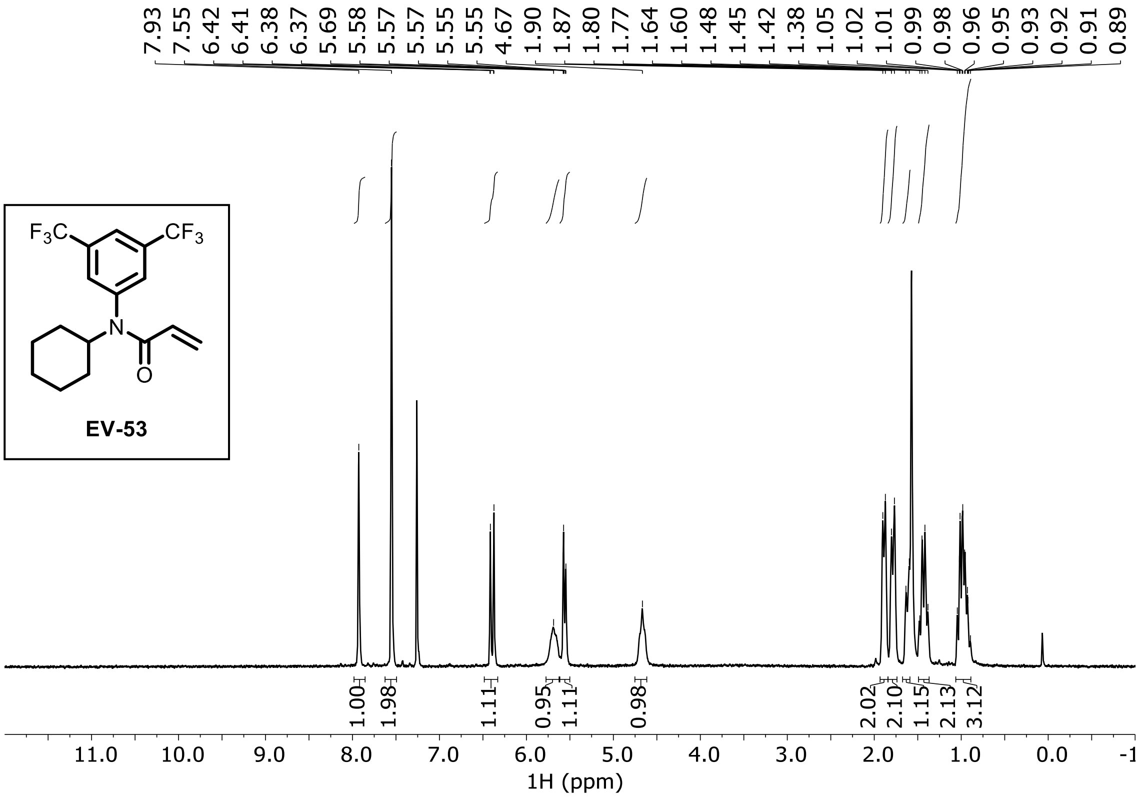

EV-54 ¹H NMR (MeOD)

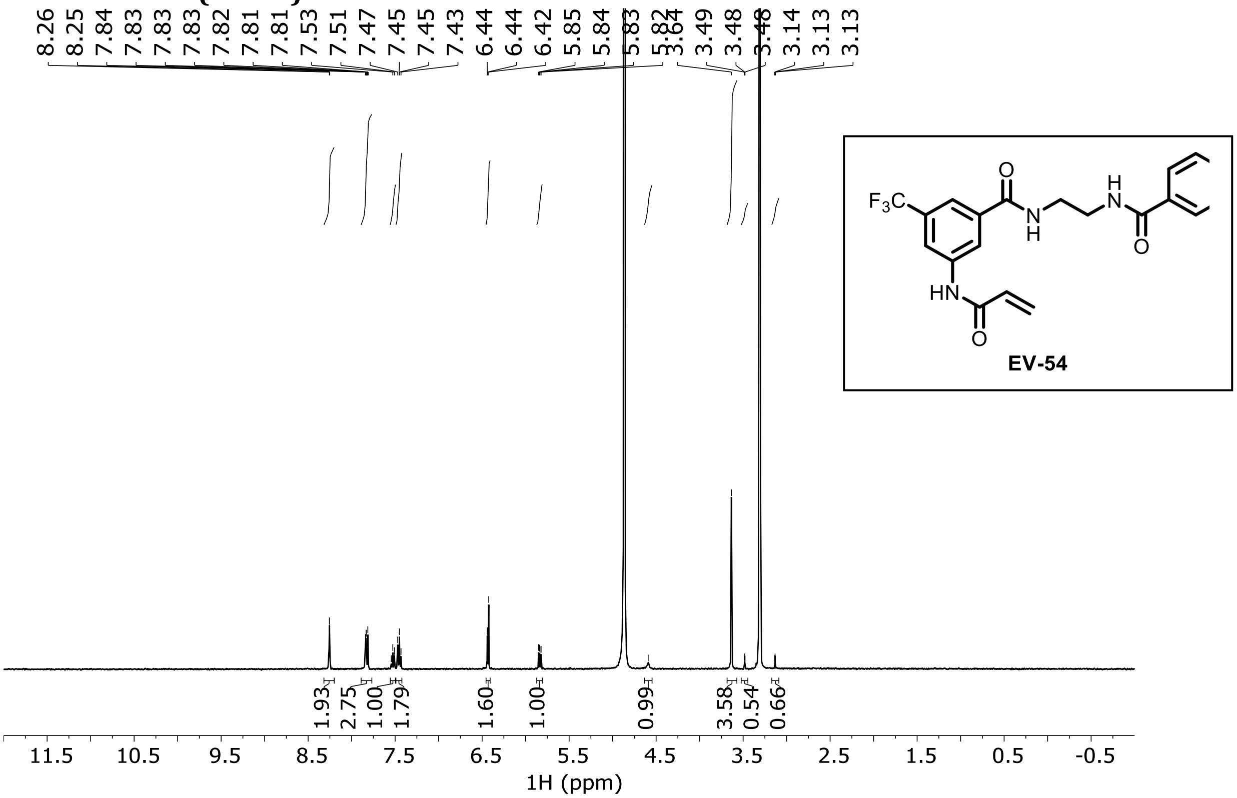

EV-55 ¹H NMR (MeOD)

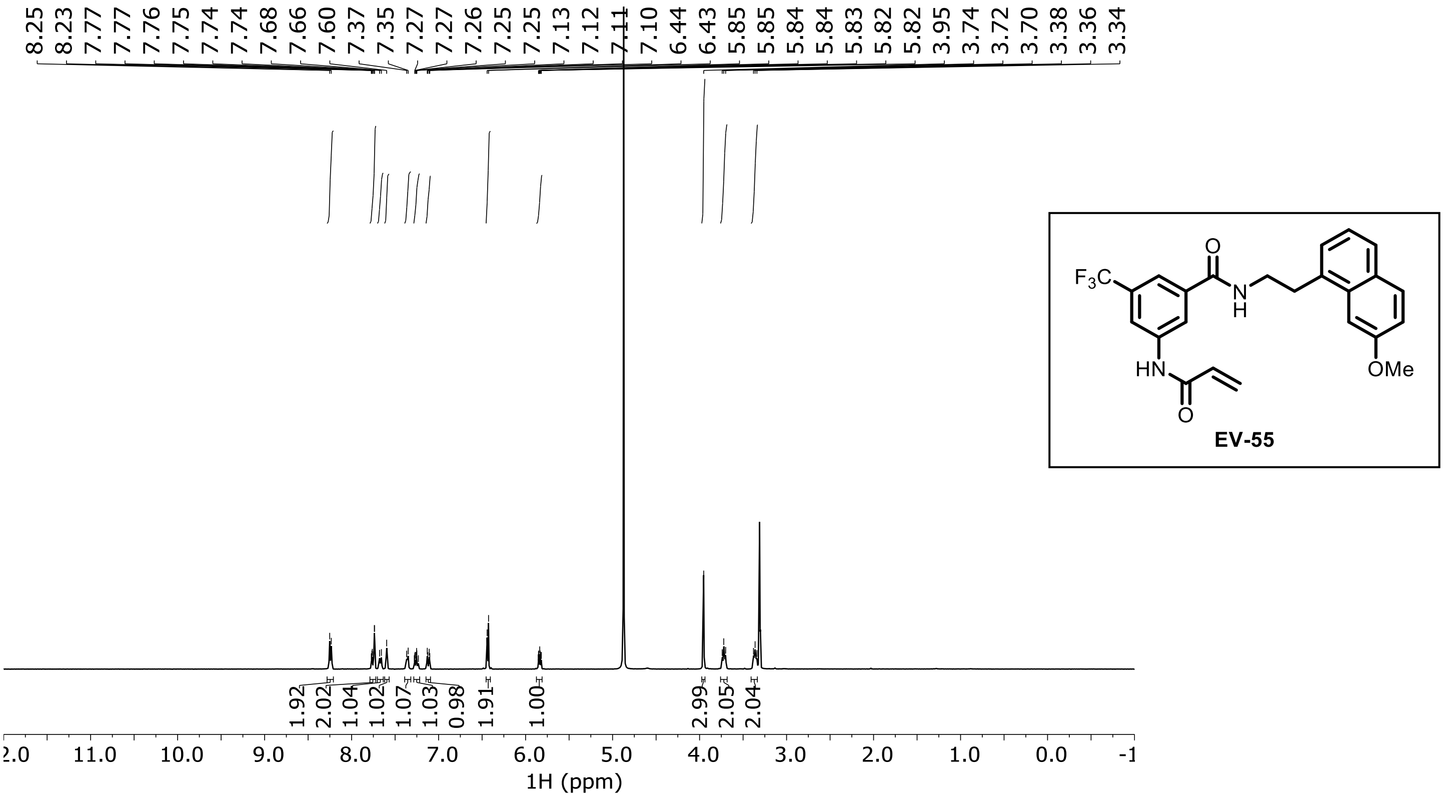

**EV-56 ¹H NMR ($CDCl_3$)**

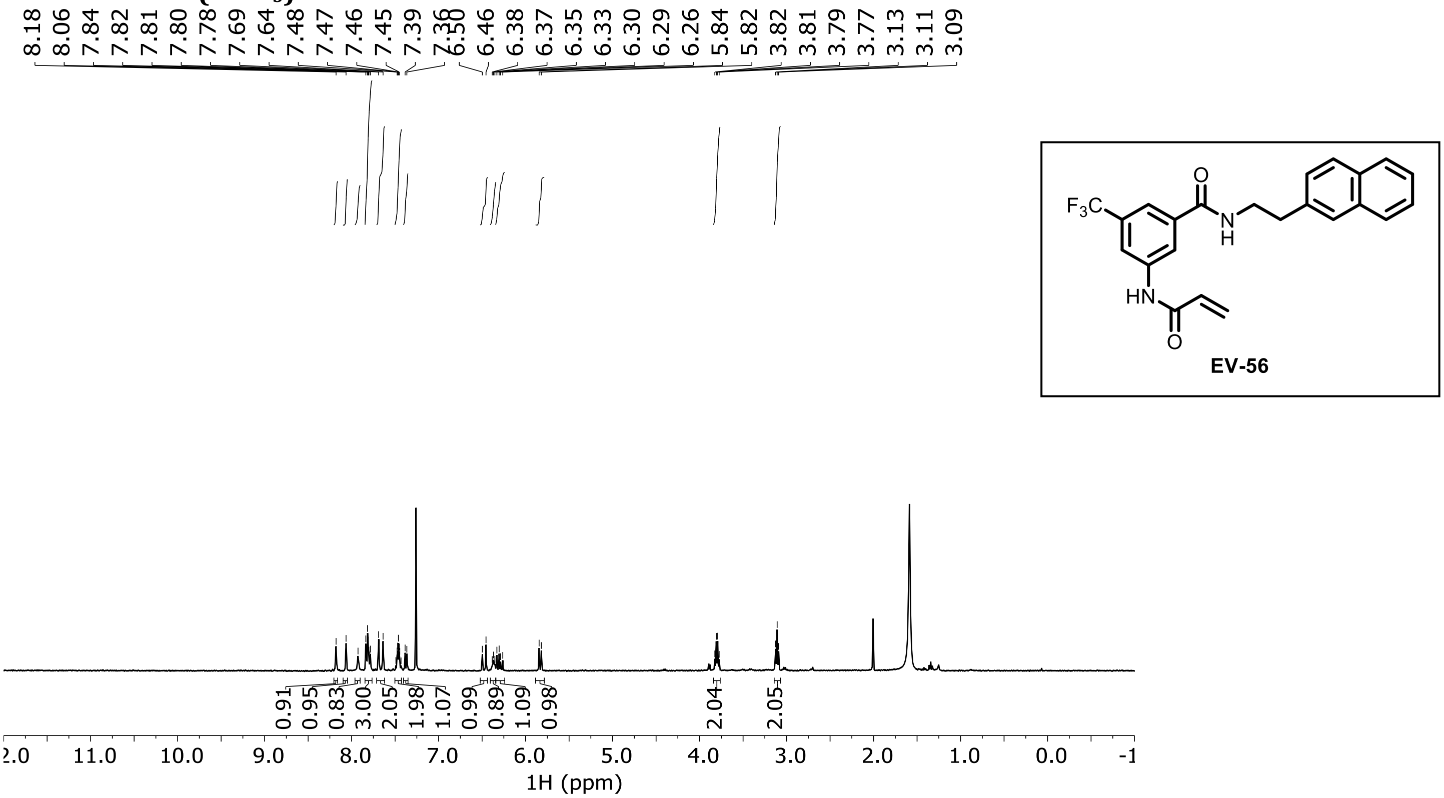

**EV-57 ¹H NMR (MeOD)**

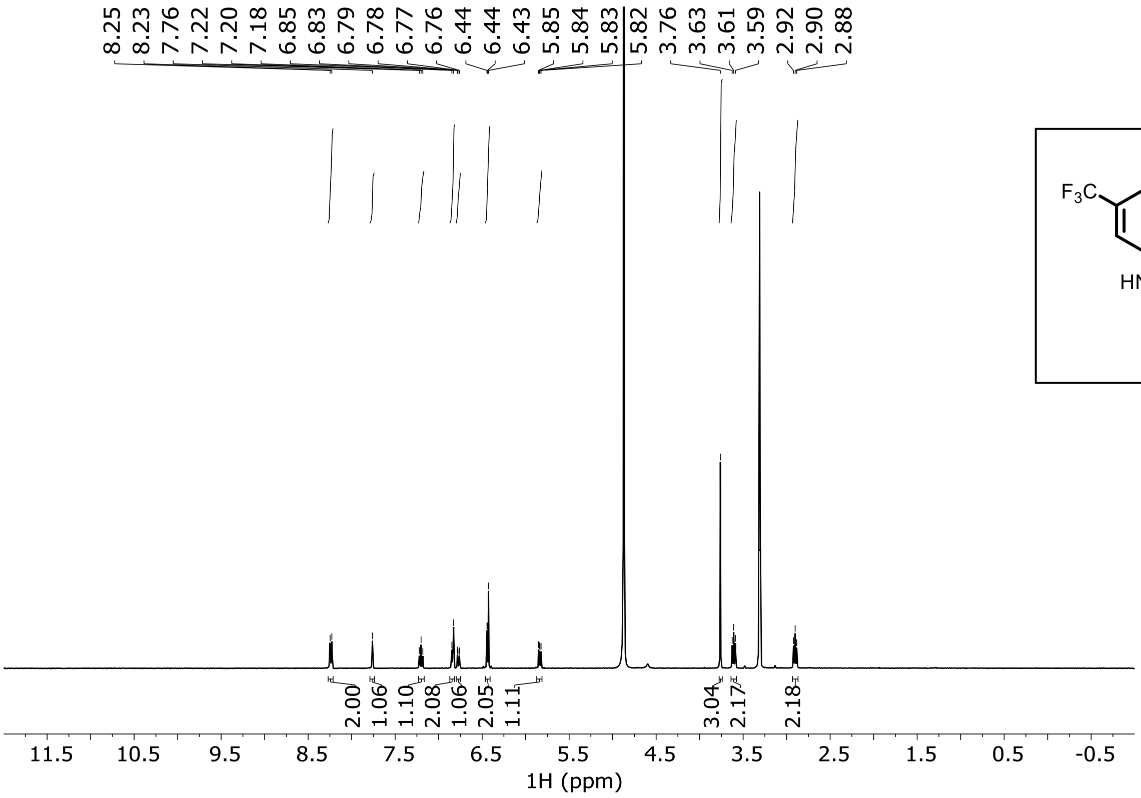

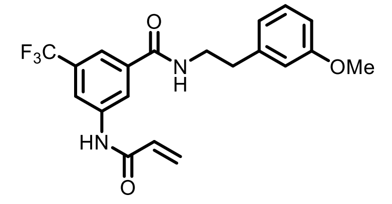

EV-57

**EV-58 $^1$H NMR (MeOD)**

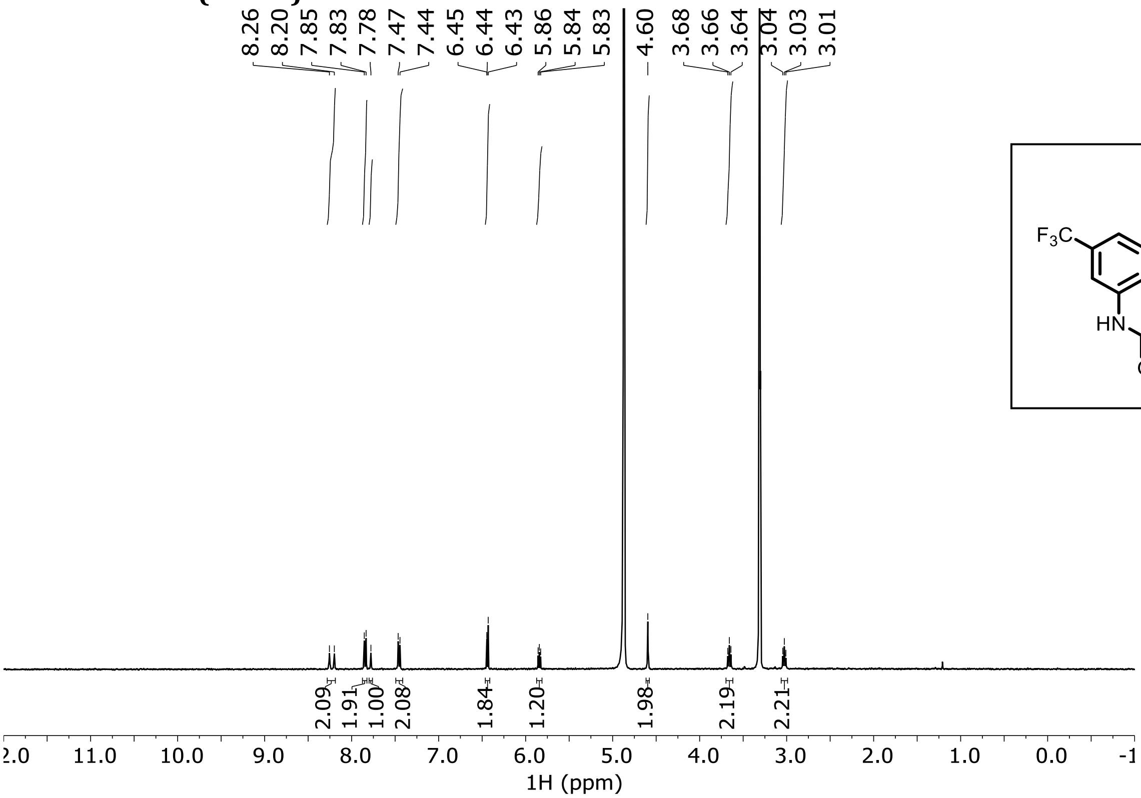

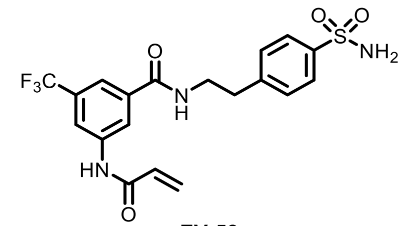

EV-58

**EV-59 $^1$H NMR (DMSO-$d_6$)**

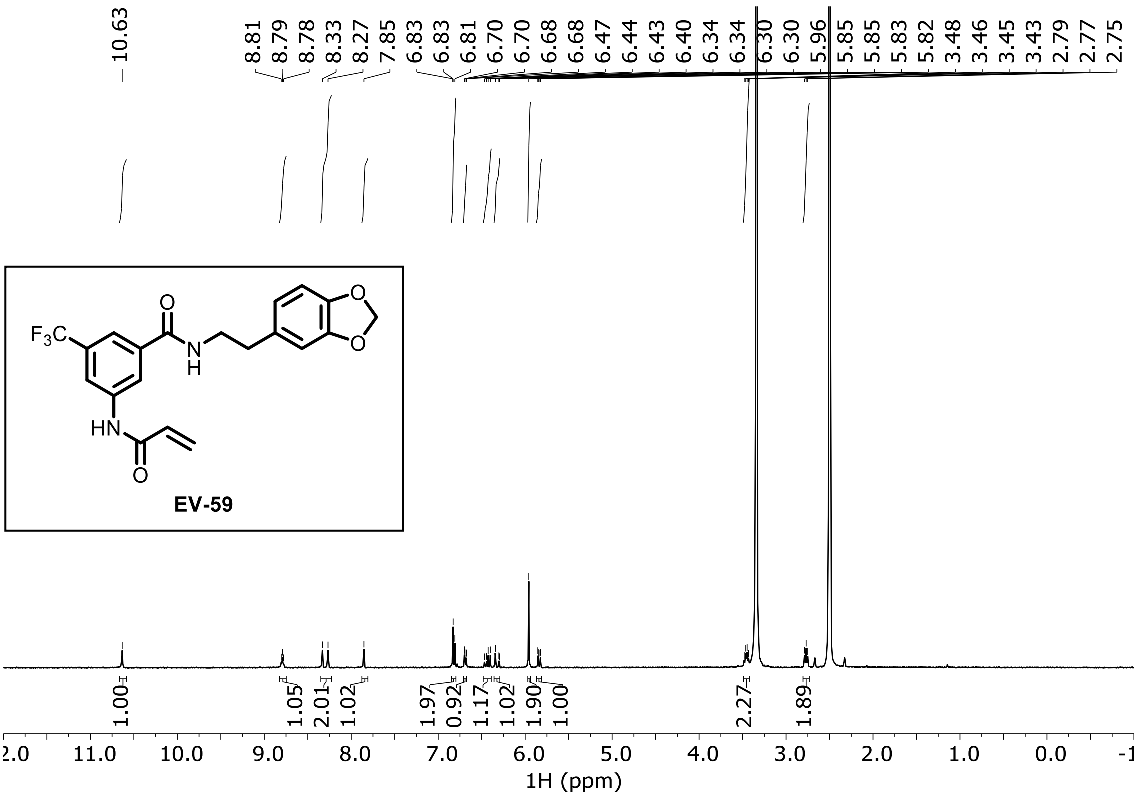

**EV-60 $^{1}H$ NMR (MeOD)**

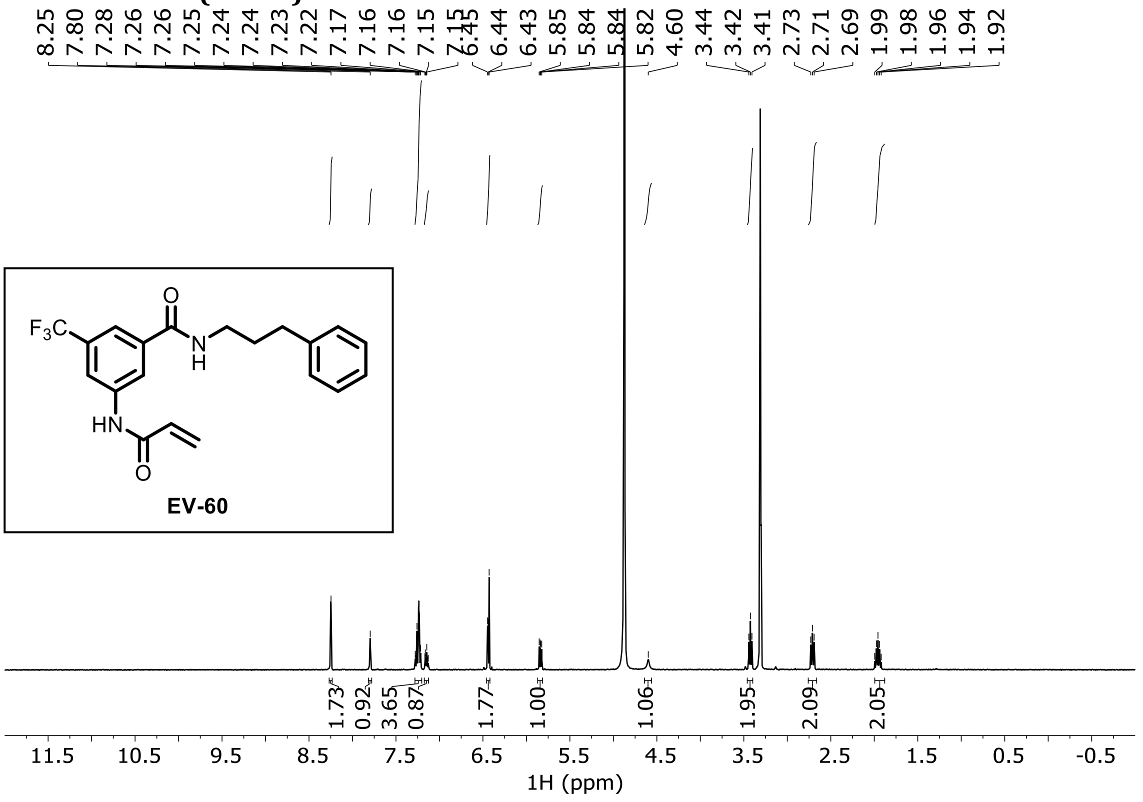

**EV-61 $^{1}H$ NMR ($CDCl_3$)**

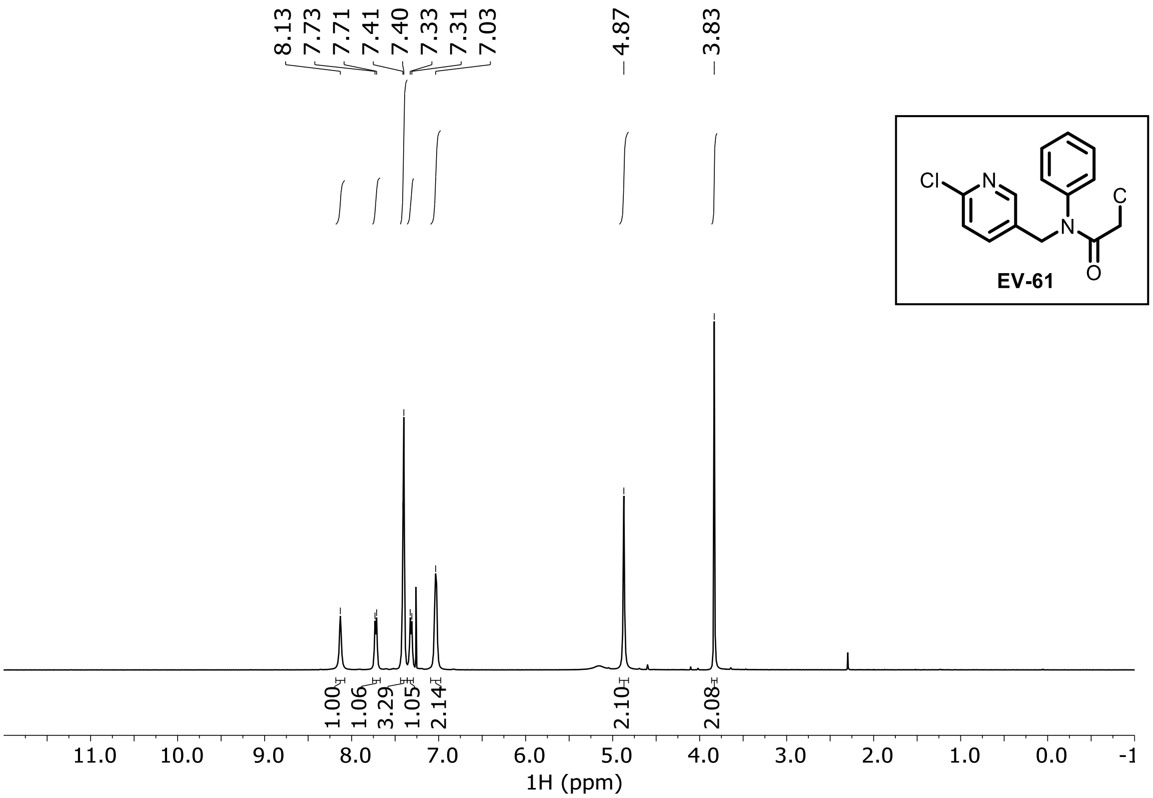

**EV-62 ¹H NMR ($CDCl_3$)**

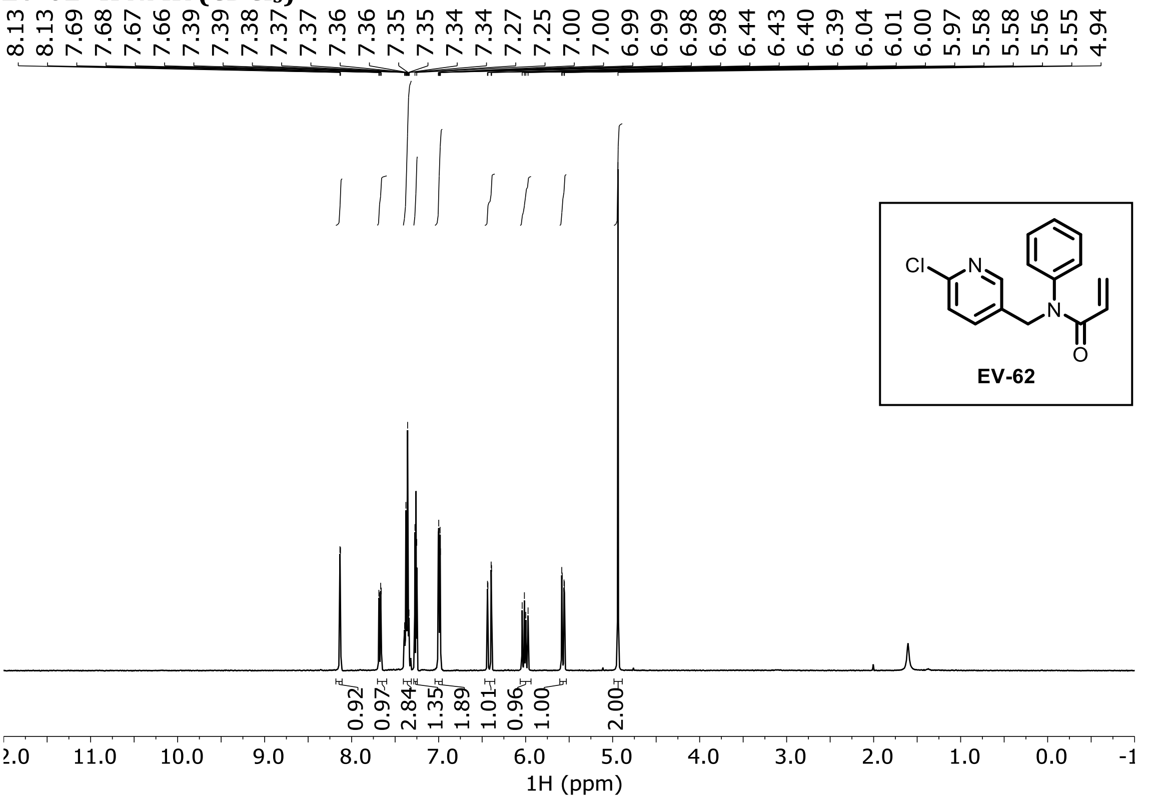

**EV-63 ¹H NMR (DMSO-$d_6$)**

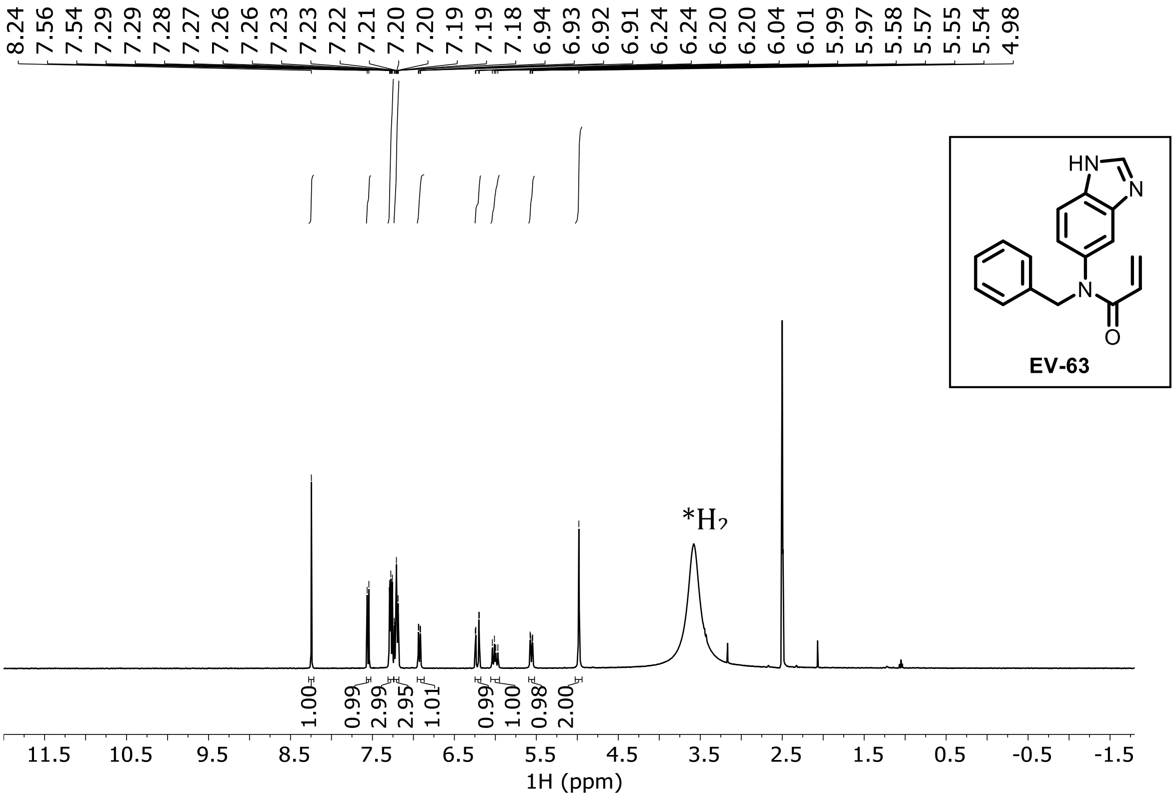

**EV-64 ¹H NMR ($CDCl_3$)**

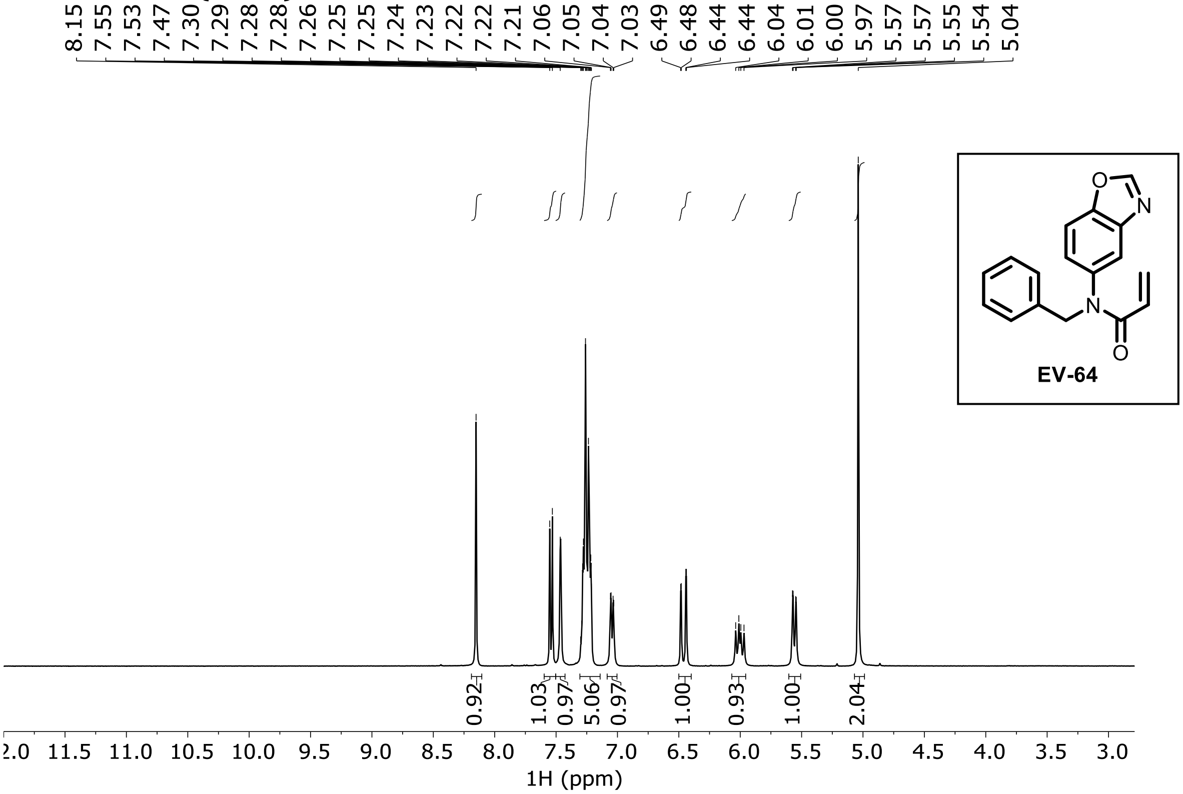

**EV-65 ¹H NMR (DMSO-$d_6$)**

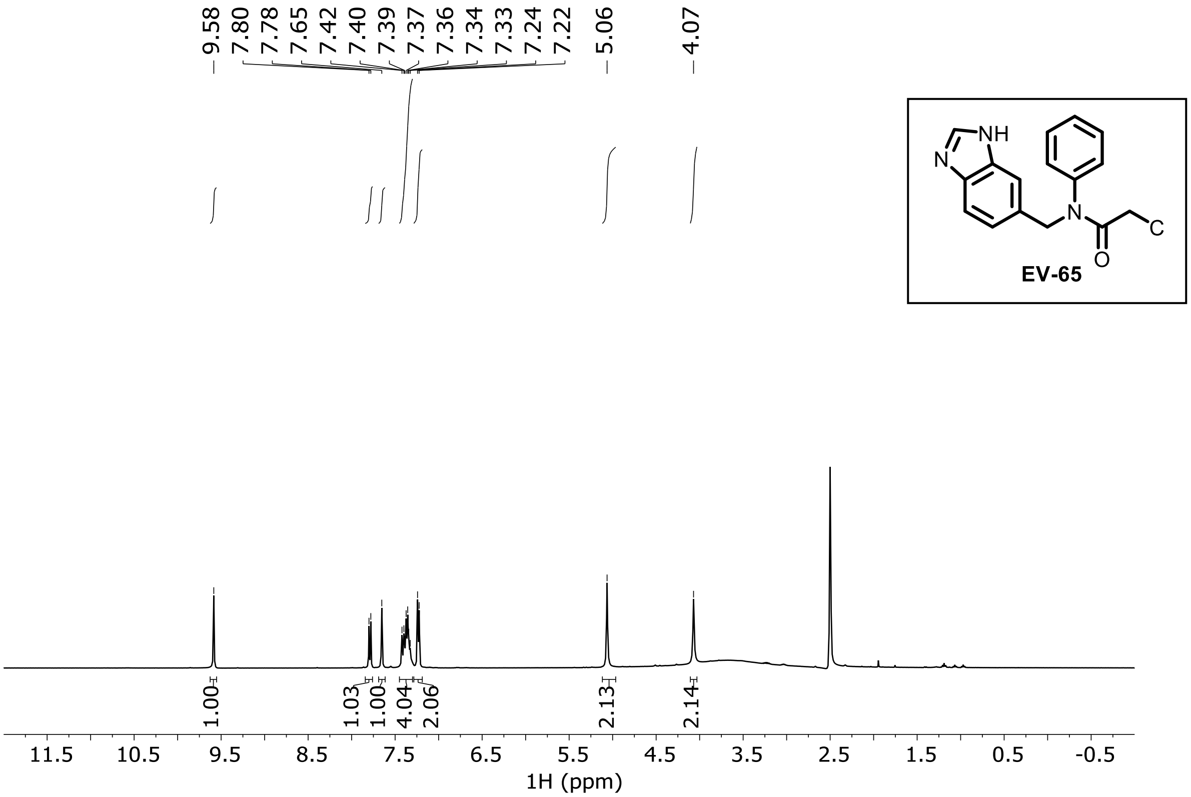

**EV-66 ¹H NMR (DMSO-$d_6$)**

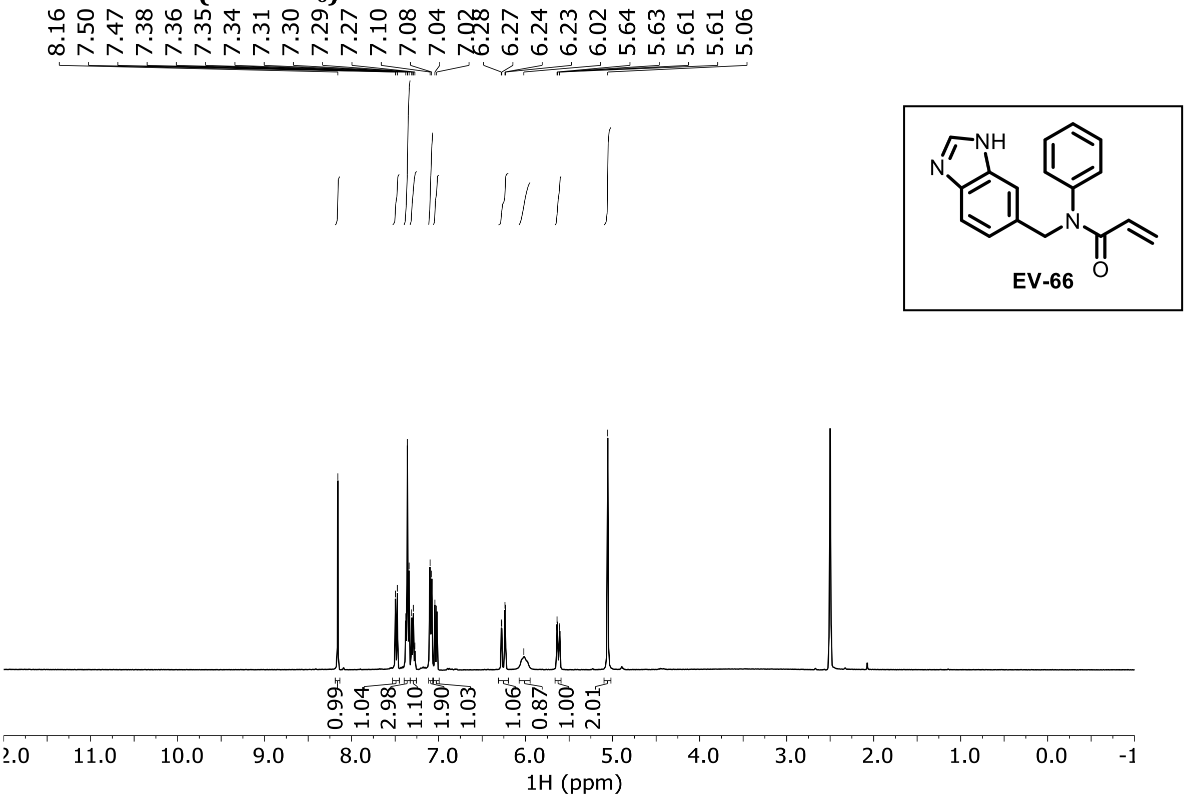

**EV-67 ¹H NMR (DMSO-$d_6$)**

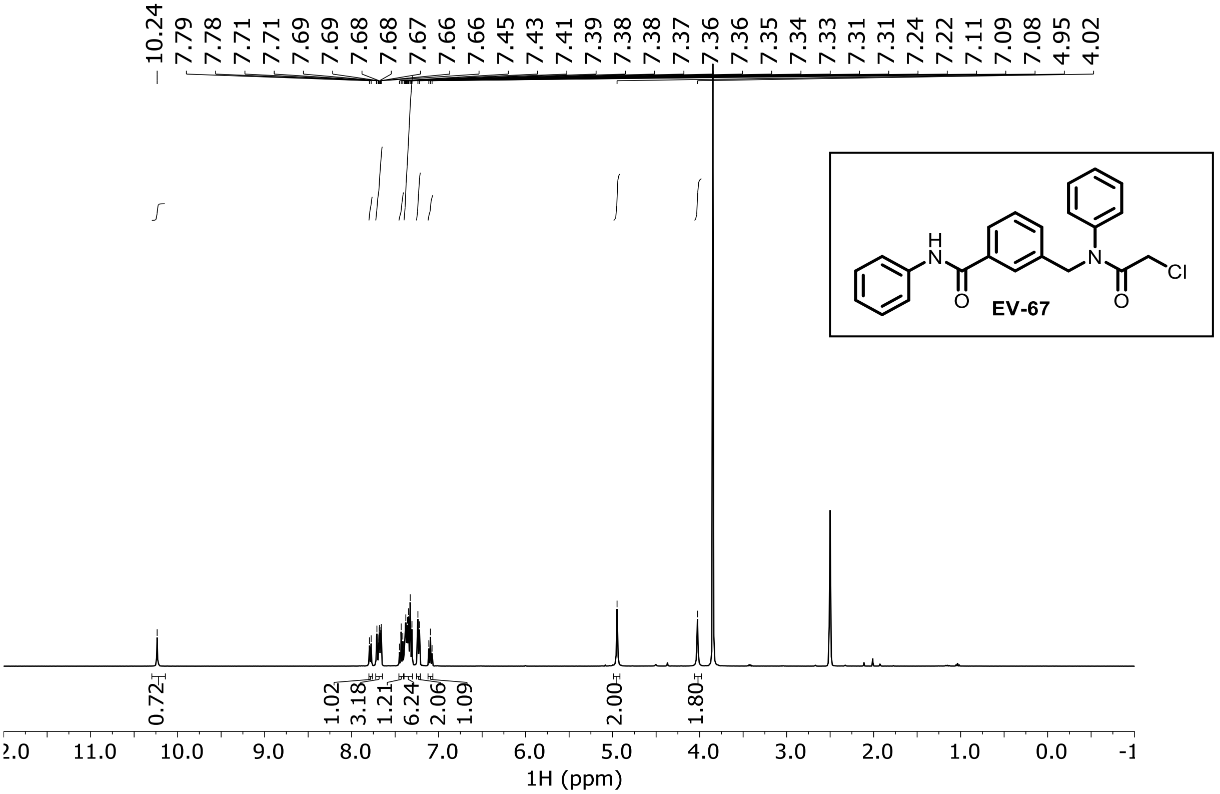

**EV-68 $^1$H NMR (DMSO-$d_6$)**

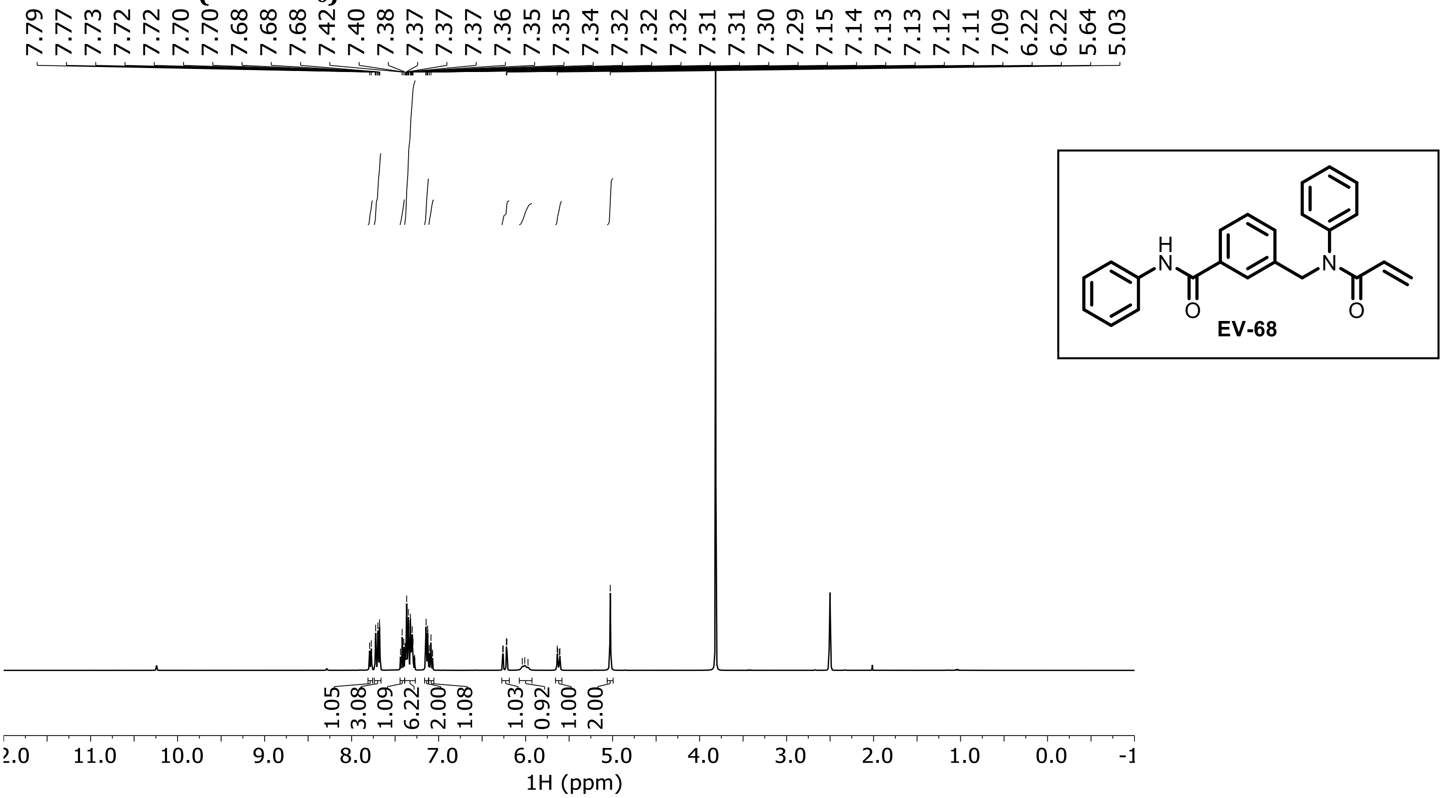

**EV-69 $^1$H NMR (DMSO-$d_6$)**

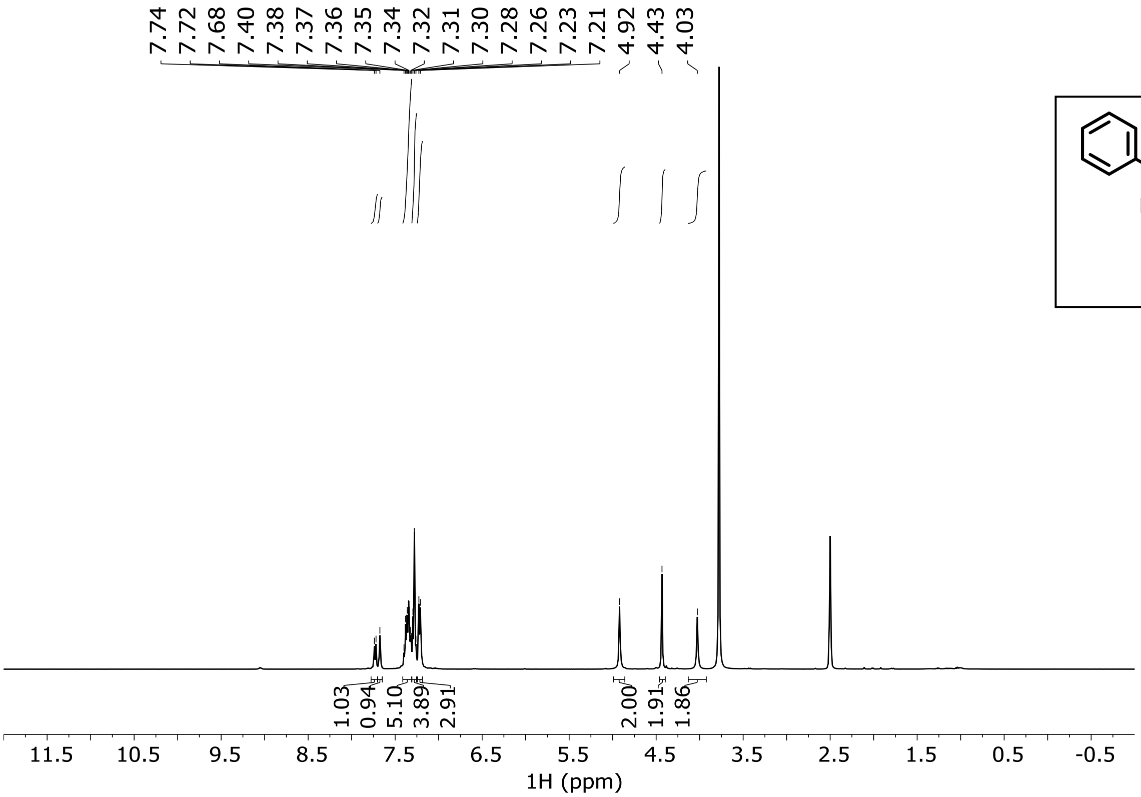

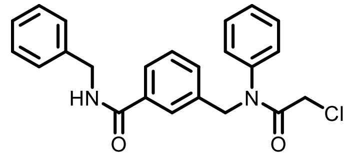

EV-69

**EV-70 $^1$H NMR (DMSO-$d_6$)**

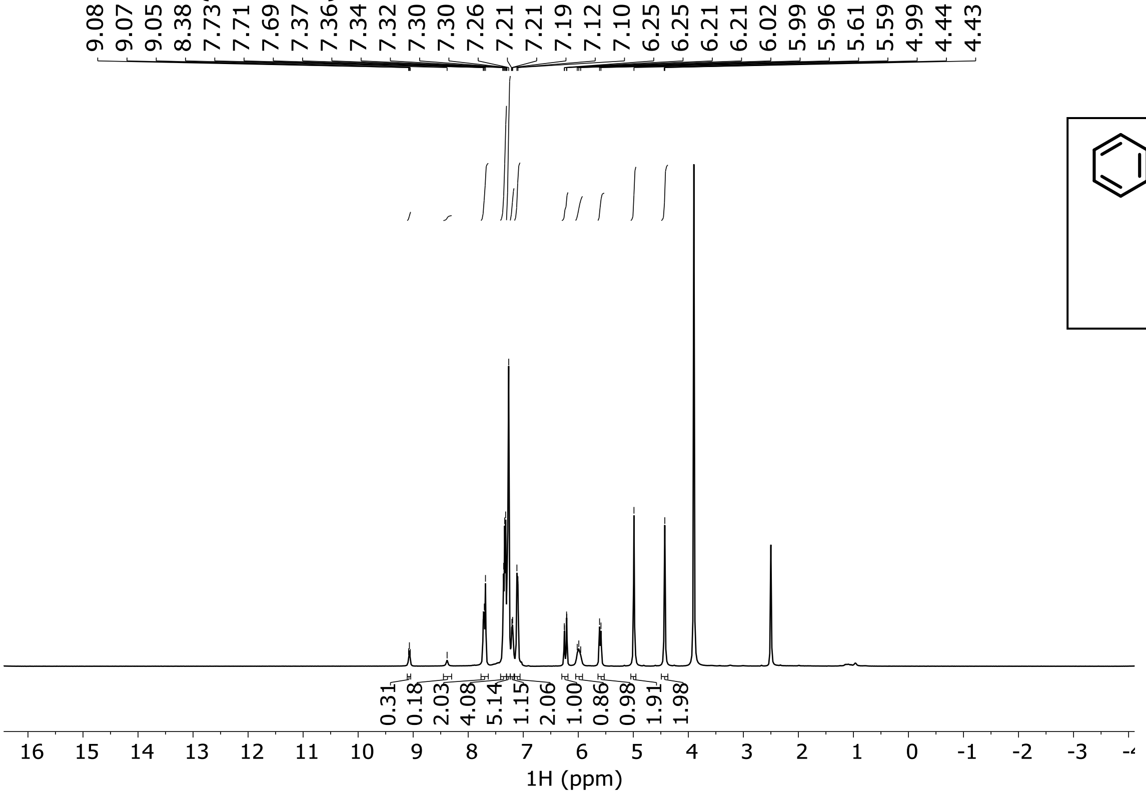

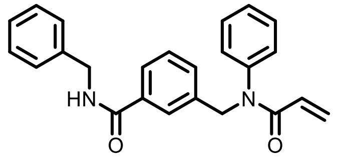

EV-70

**EV-71 $^1$H NMR (DMSO-$d_6$)**

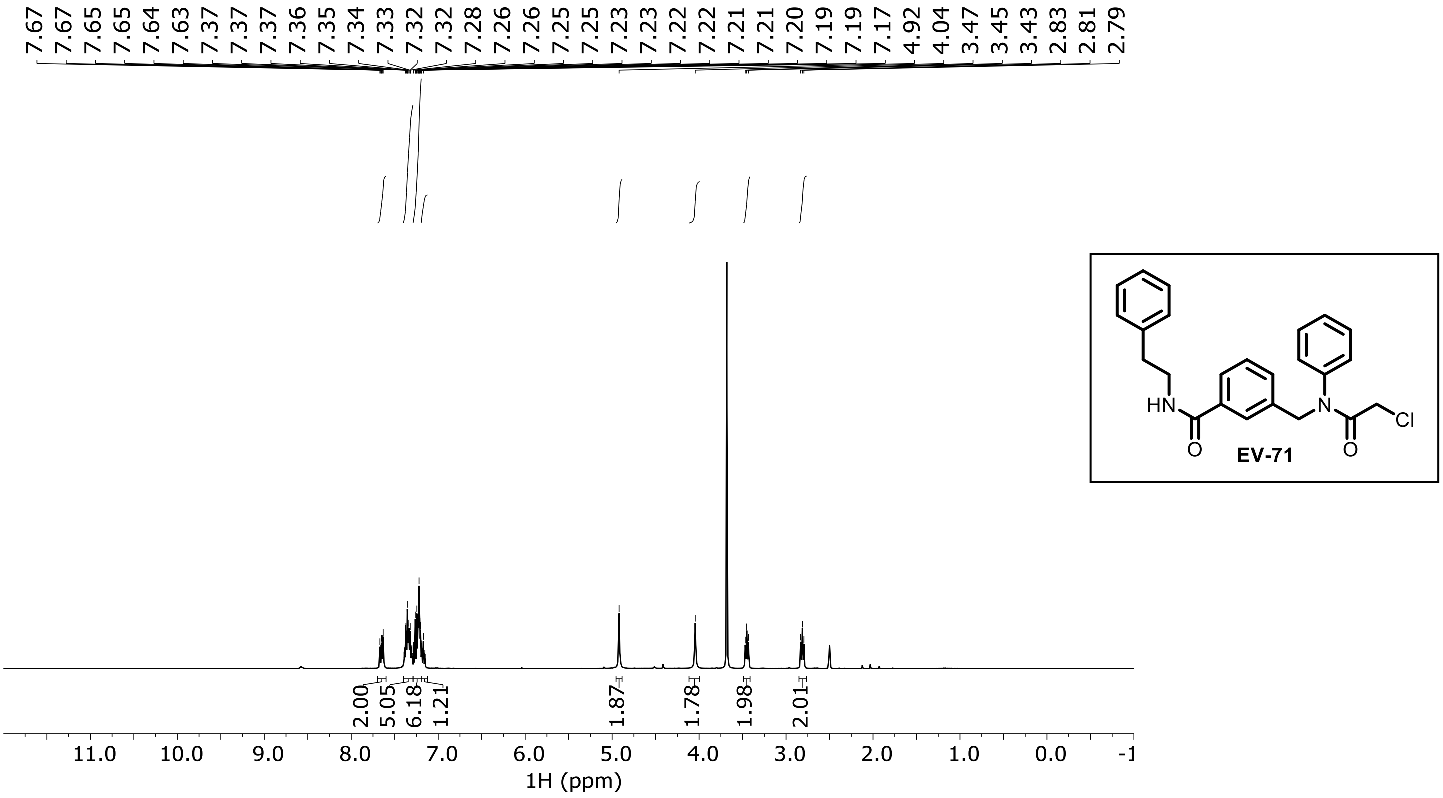

**EV-72 ¹H NMR (DMSO-$d_6$)**

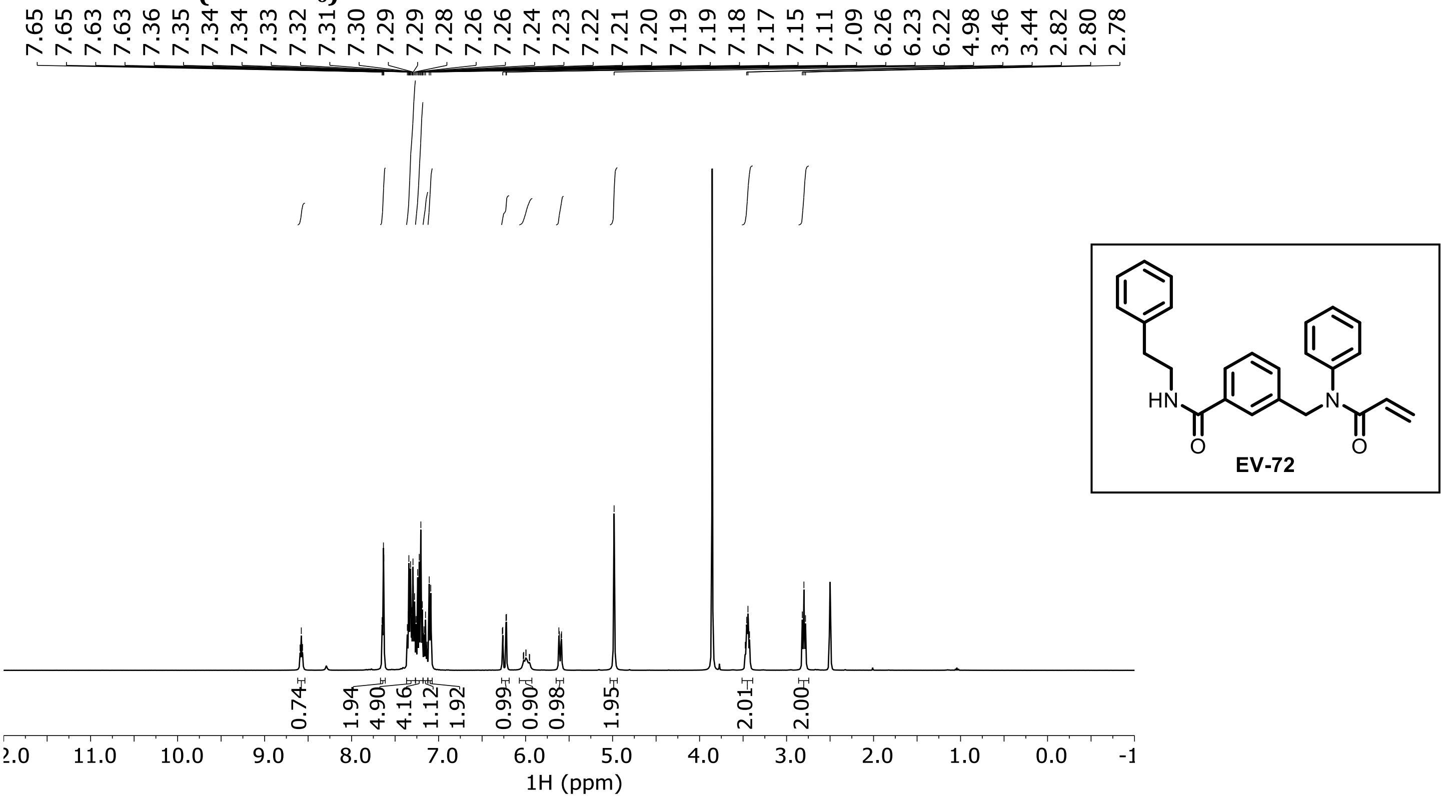

**EV-73 ¹H NMR (DMSO-$d_6$)**

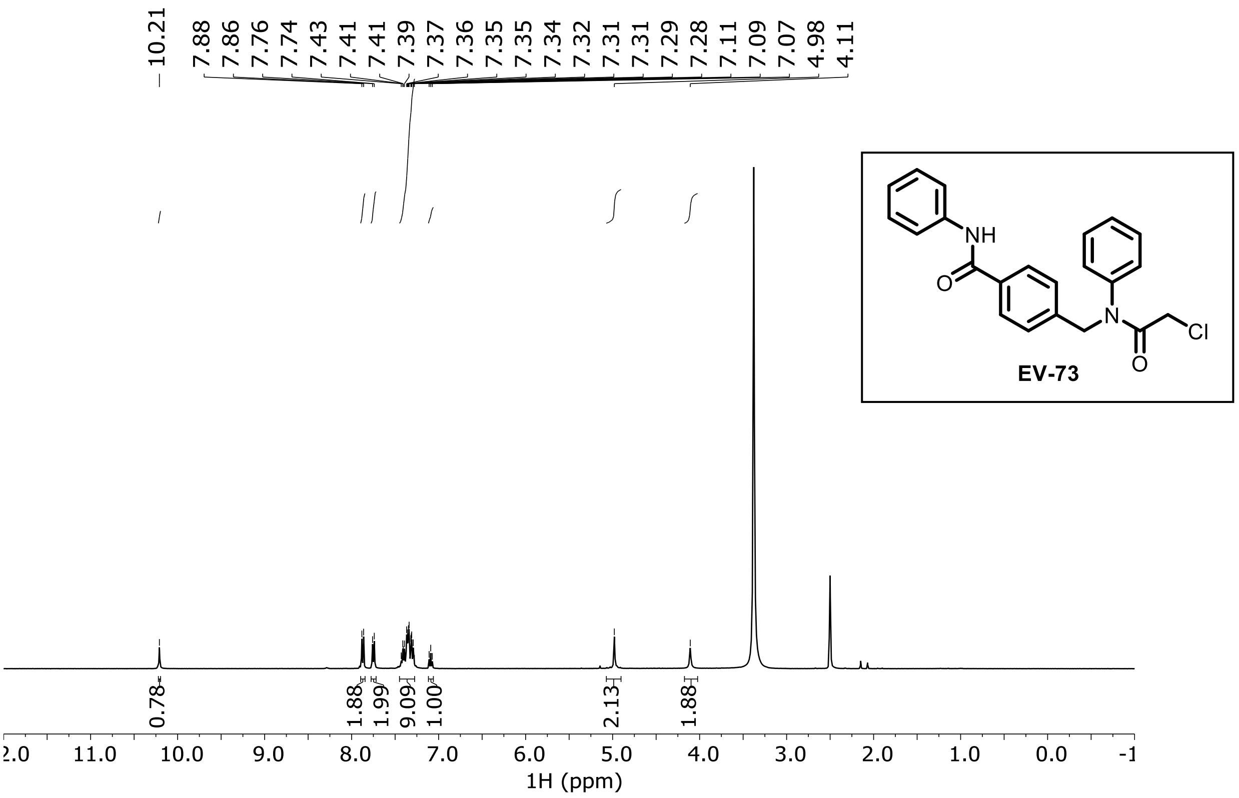

EV-74 1H NMR (DMSO-$d_6$)

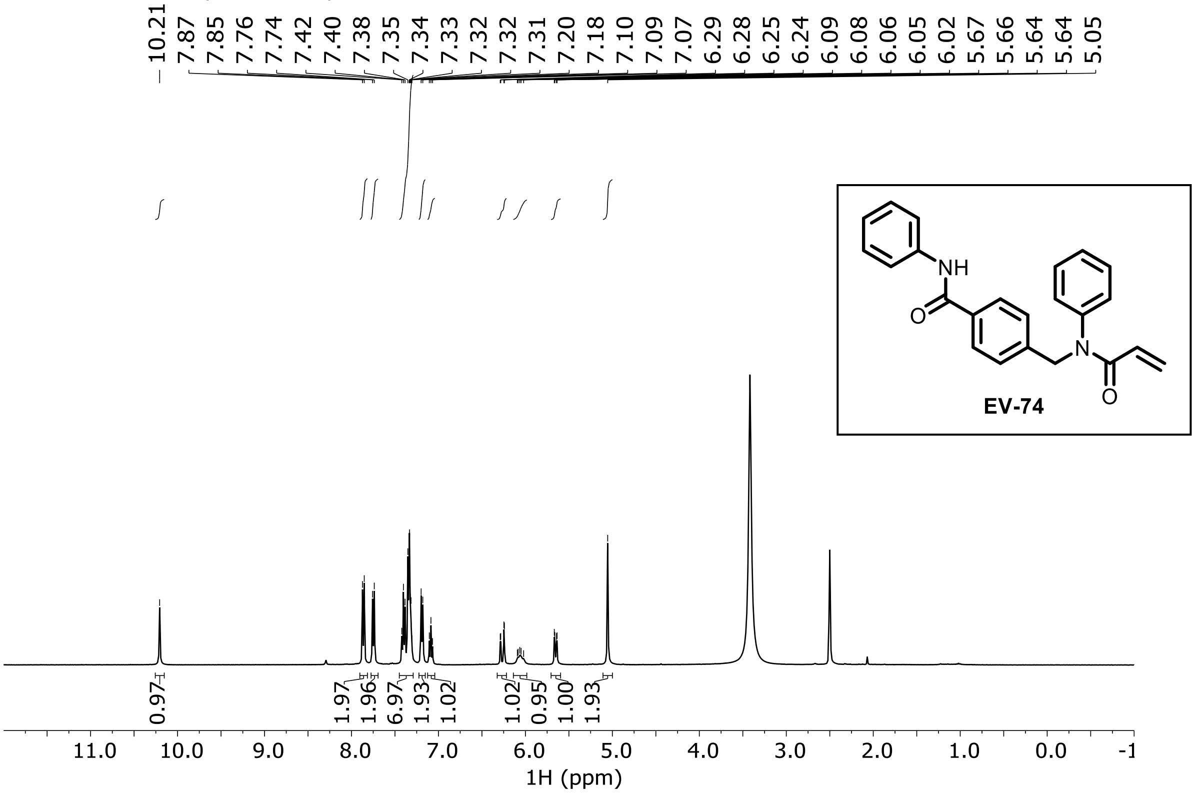

EV-75 1H NMR (DMSO-$d_6$)

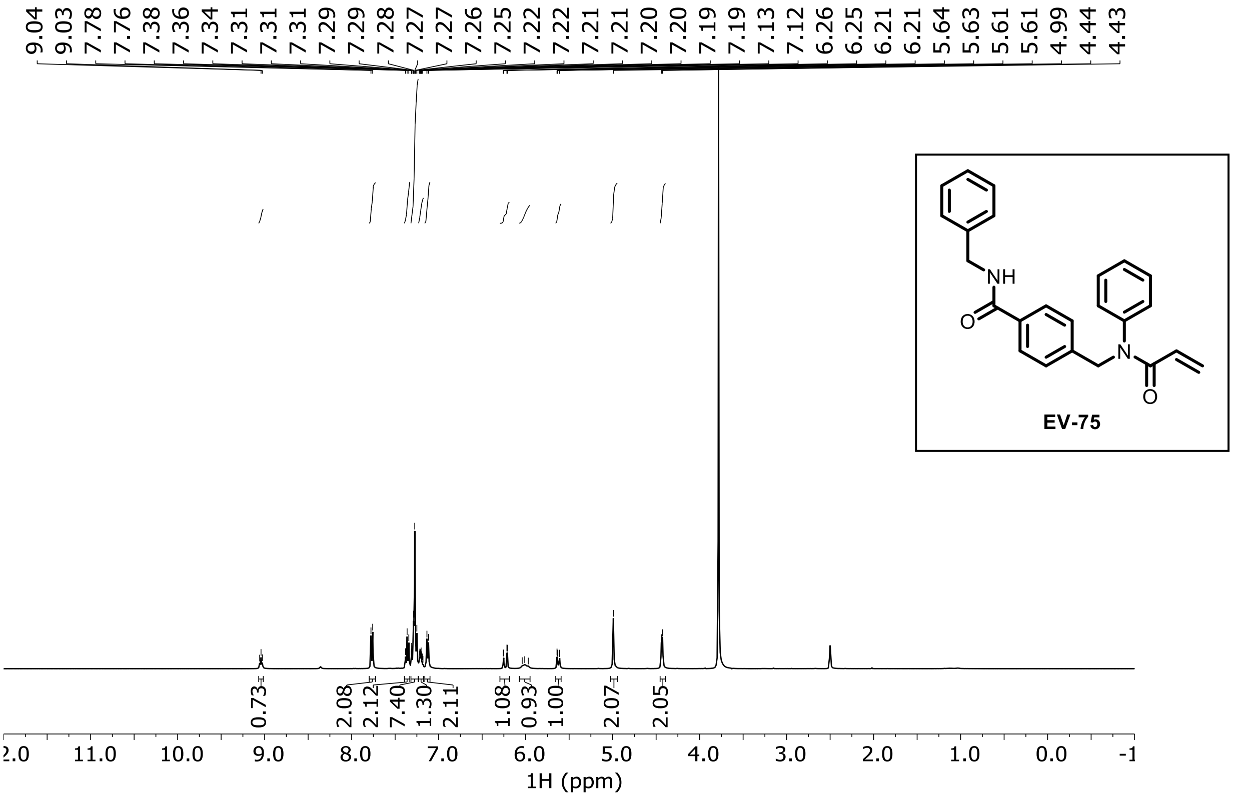

**EV-76 $^1$H NMR (DMSO-$d_6$)**

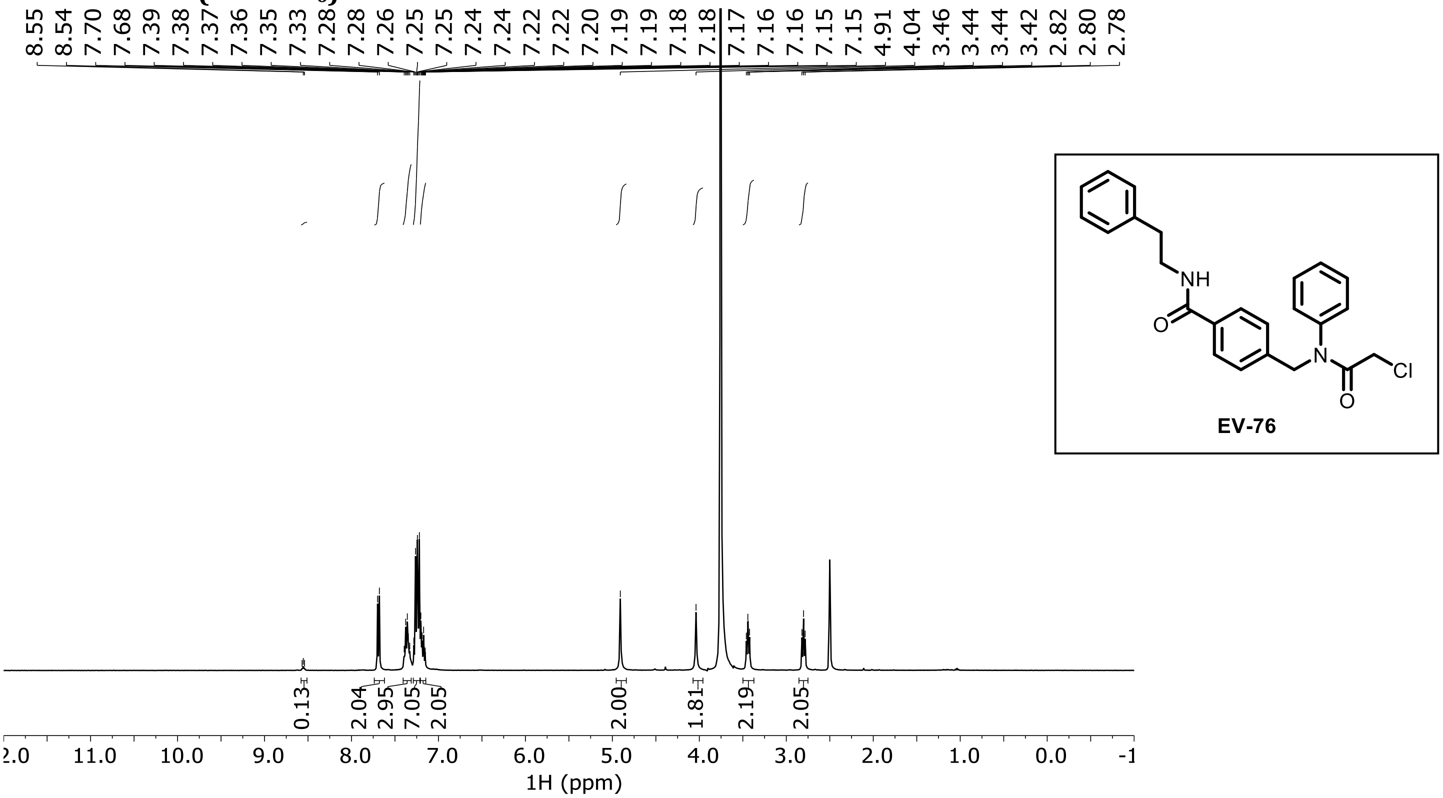

**EV-77 $^1$H NMR (DMSO-$d_6$)**

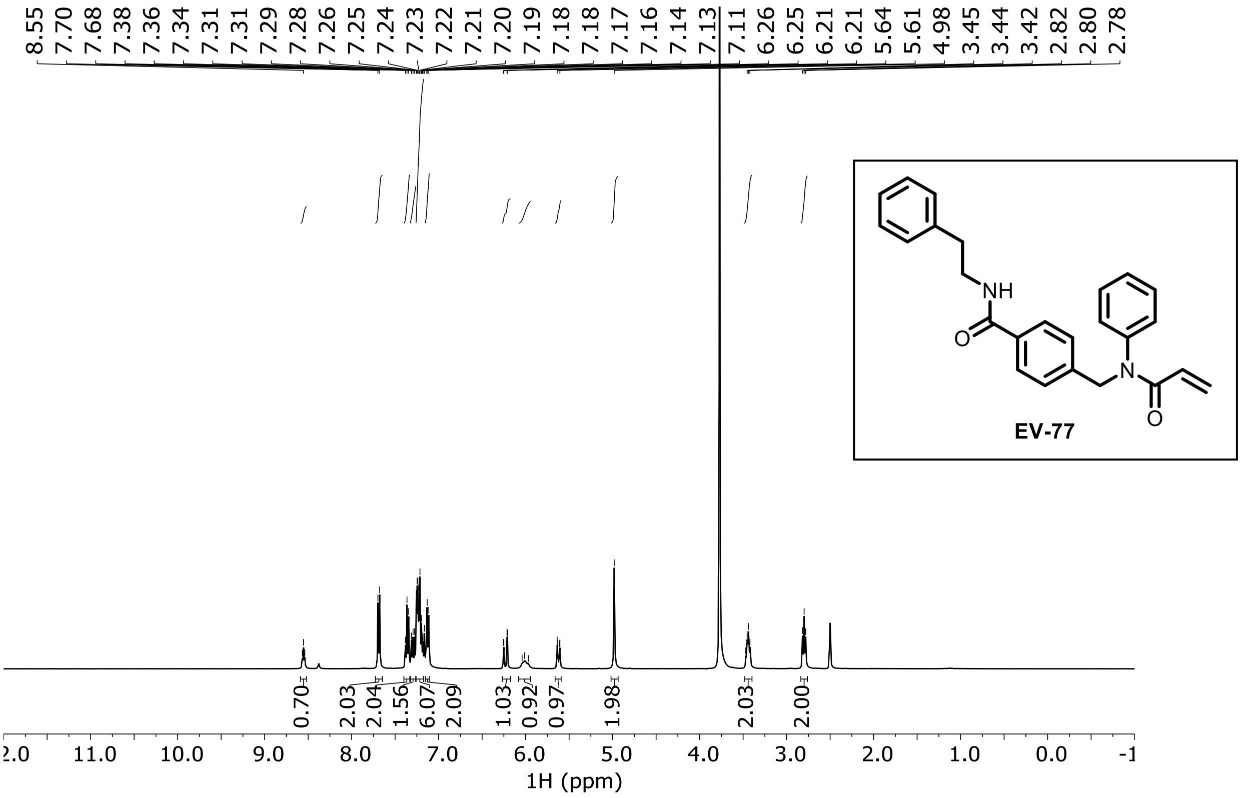

**EV-78 $^1$H NMR ($CDCl_3$)**

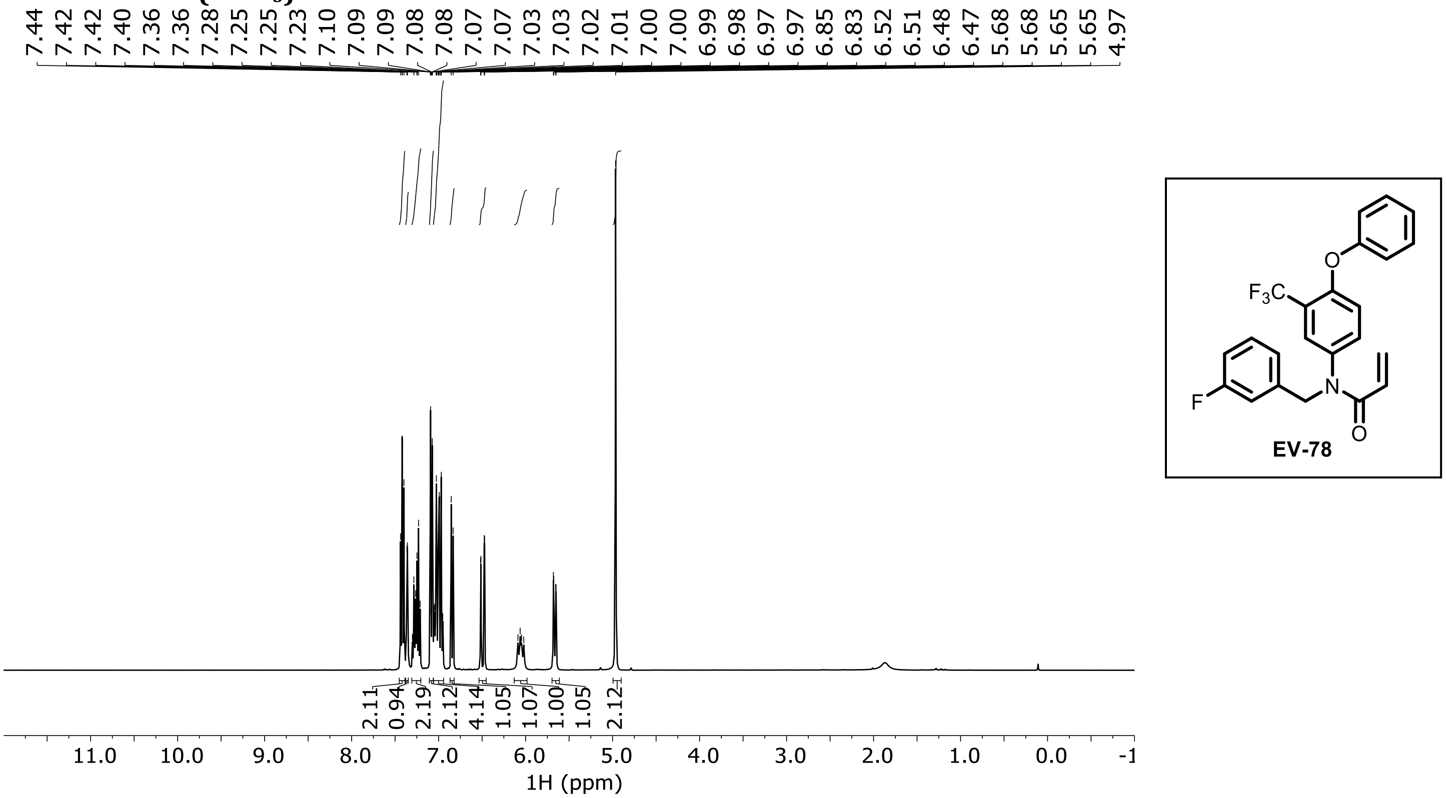

**EV-79 $^1$H NMR ($CDCl_3$)**

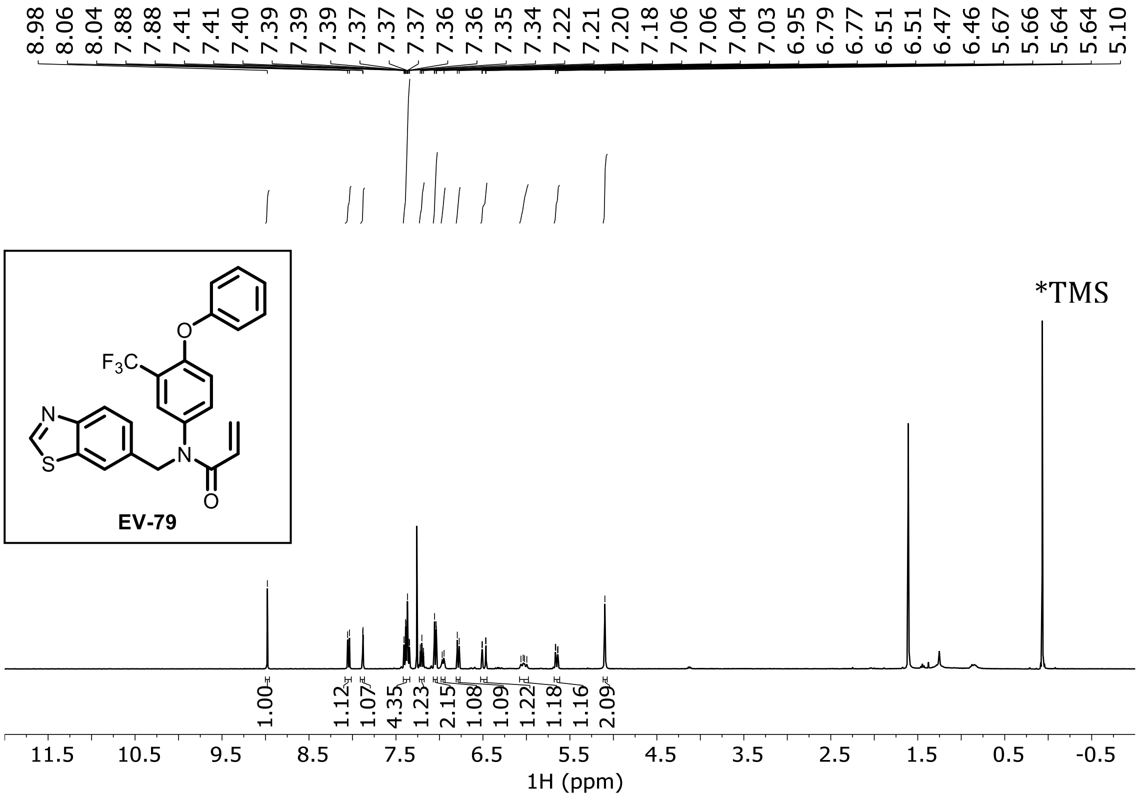

**EV-80 1H NMR ($CDCl_3$)**

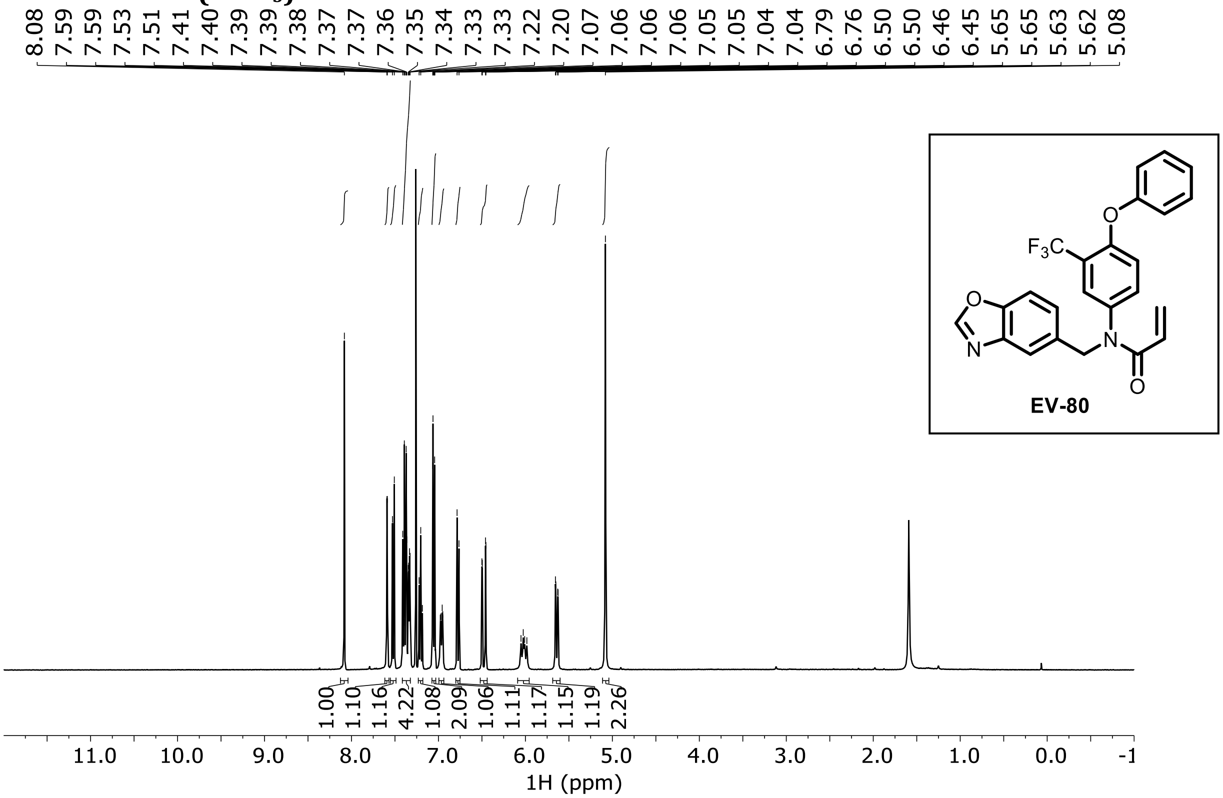

**EV-81 1H NMR ($CDCl_3$)**

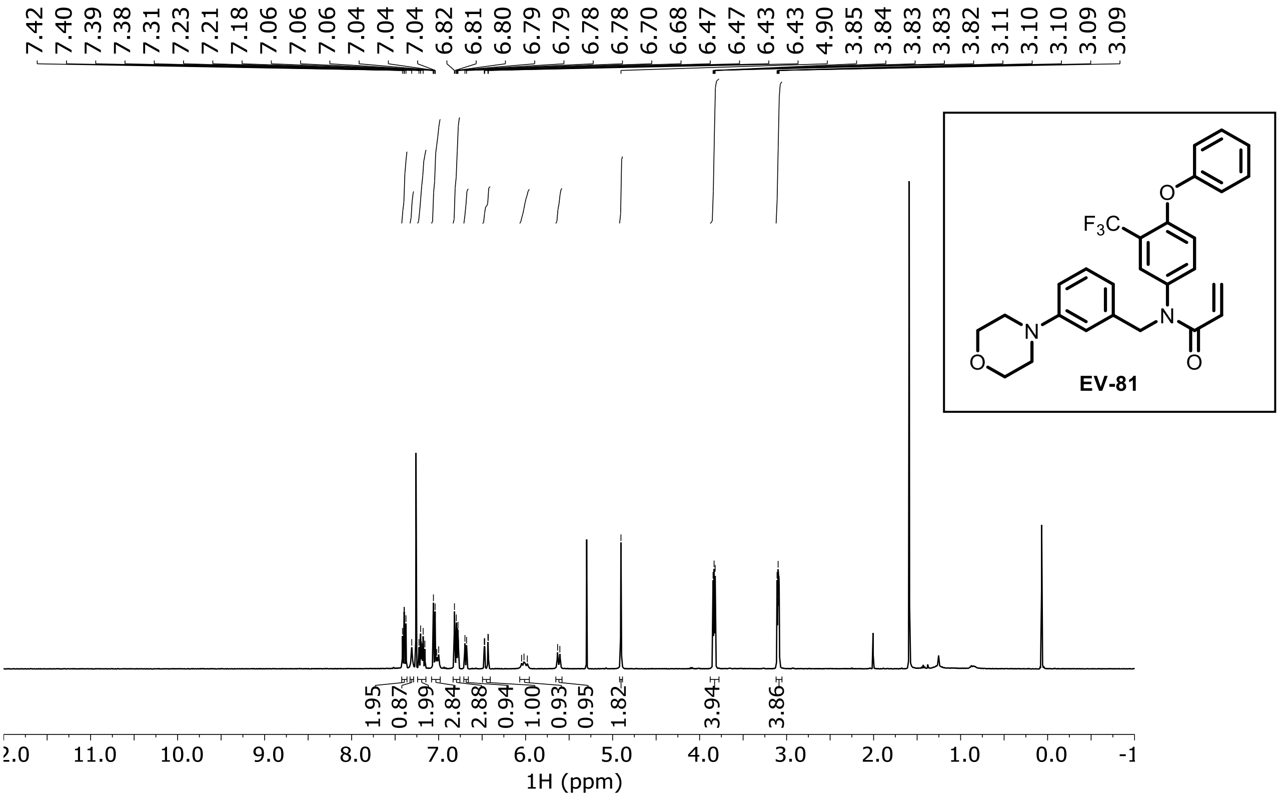

**EV-82 ¹H NMR (DMSO-$d_6$)**

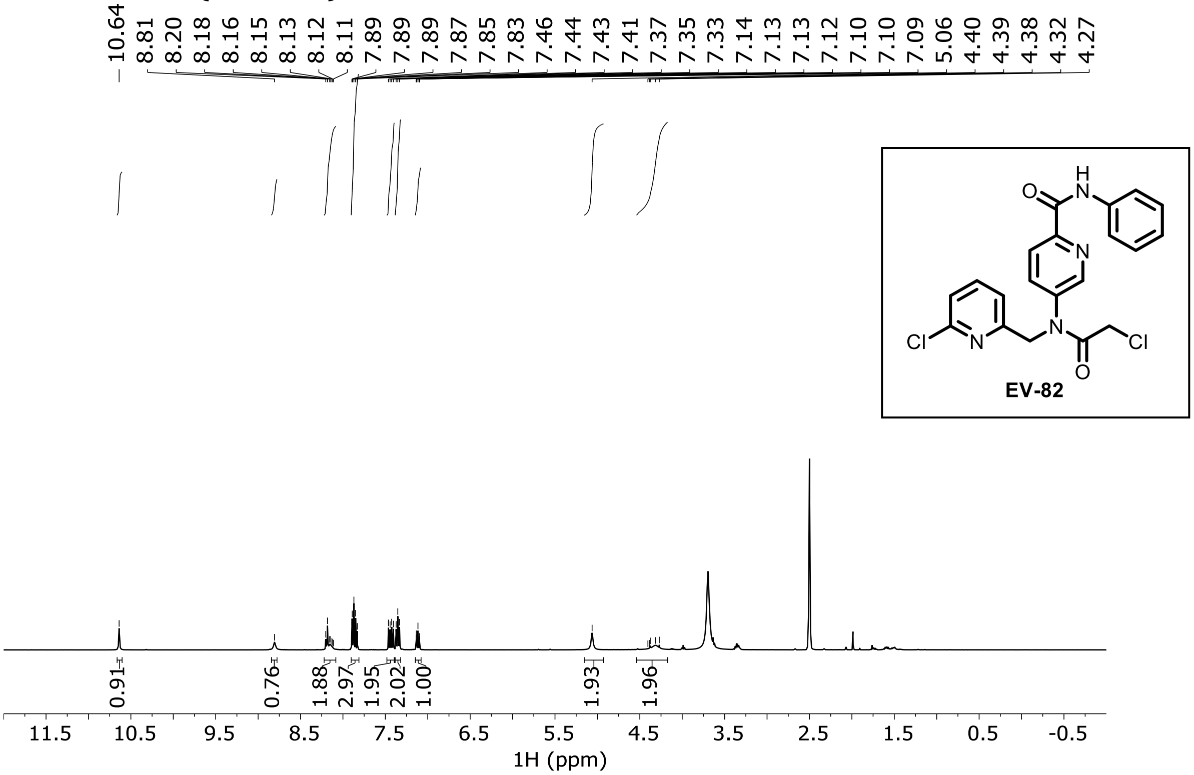

**EV-83 ¹H NMR (DMSO-$d_6$)**

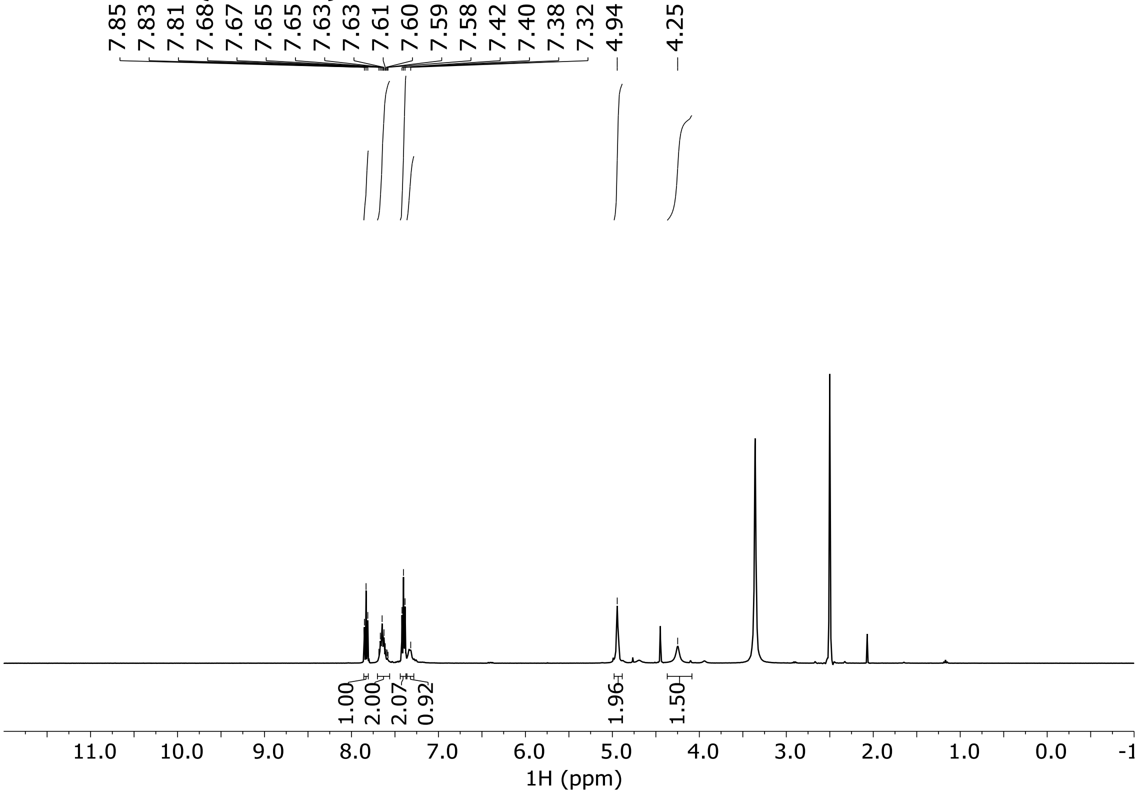

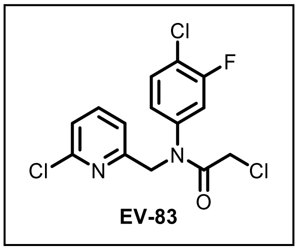

**EV-84 ¹H NMR ($CDCl_3$)**

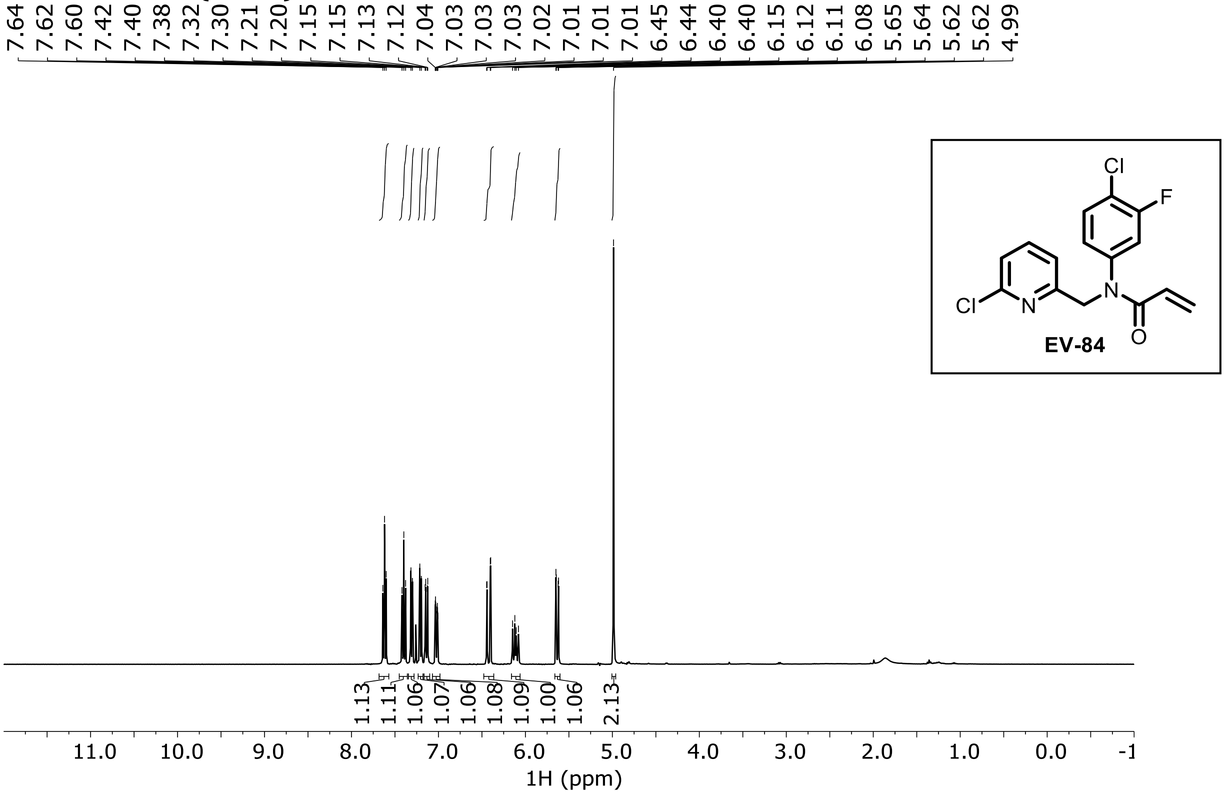

**EV-85 ¹H NMR ($CDCl_3$)**

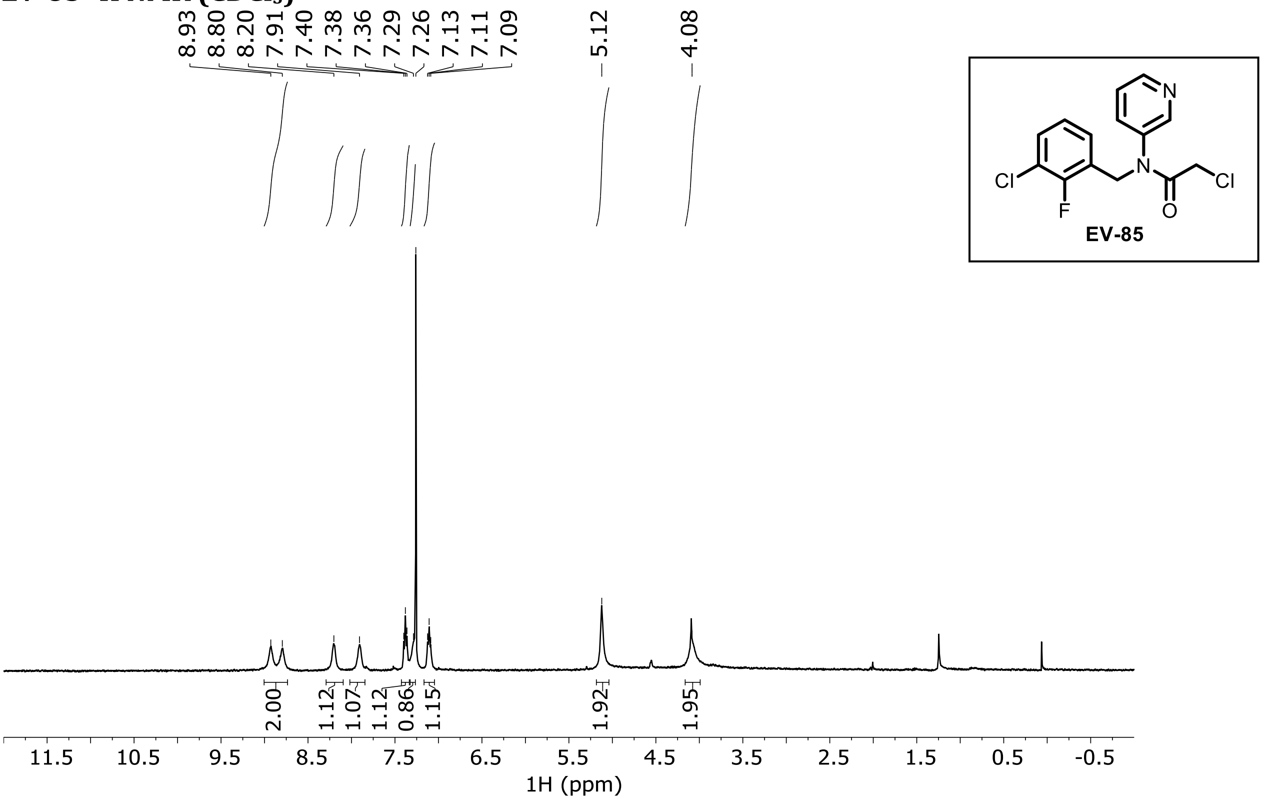

**EV-86 ¹H NMR (DMSO-$d_6$)**

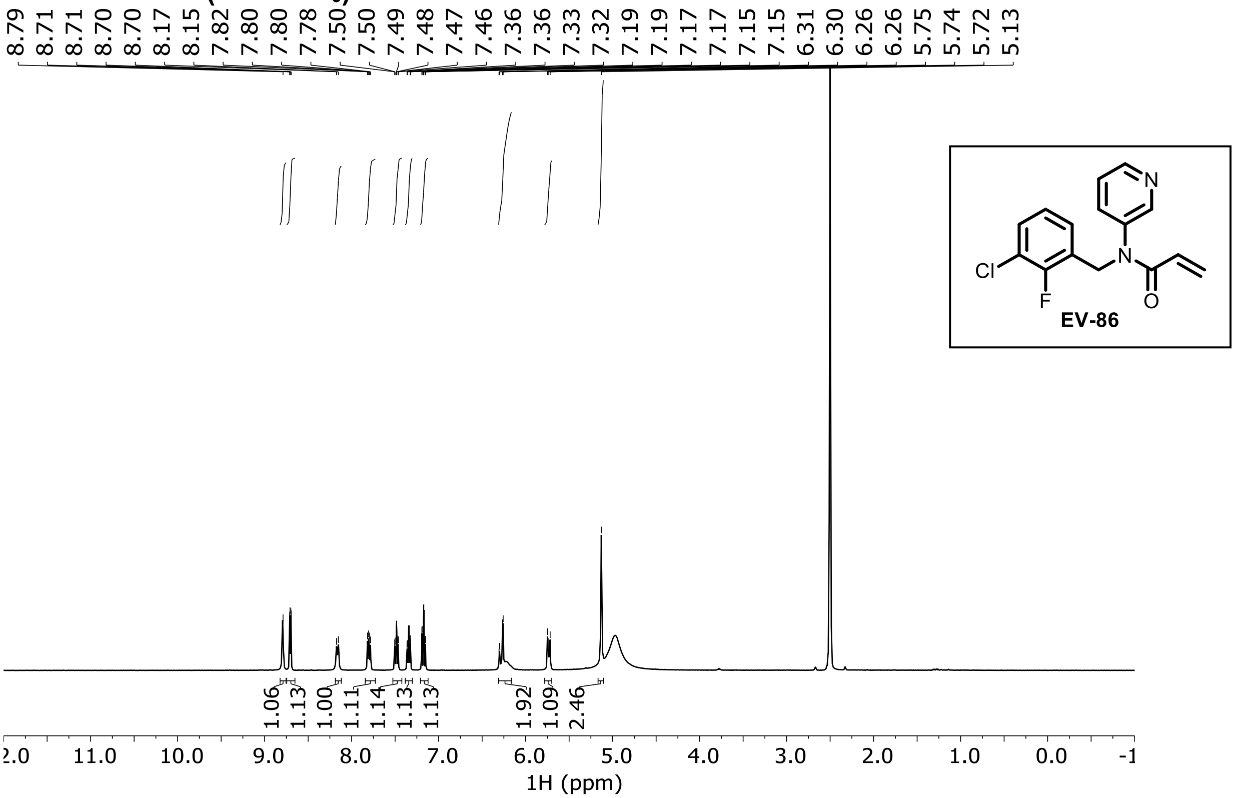

**EV-87 $^{1}$H NMR ($CDCl_3$)**

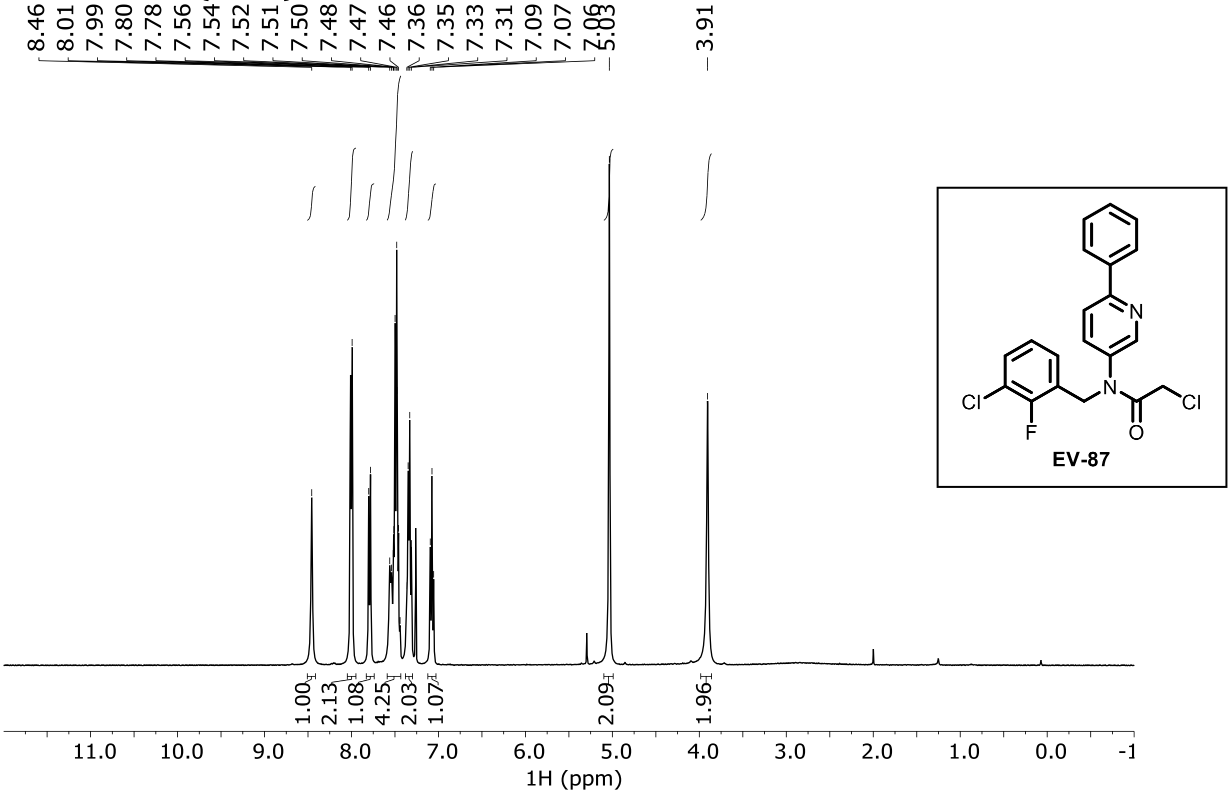

**EV-87**

**EV-88 $^{1}$H NMR ($CDCl_3$)**

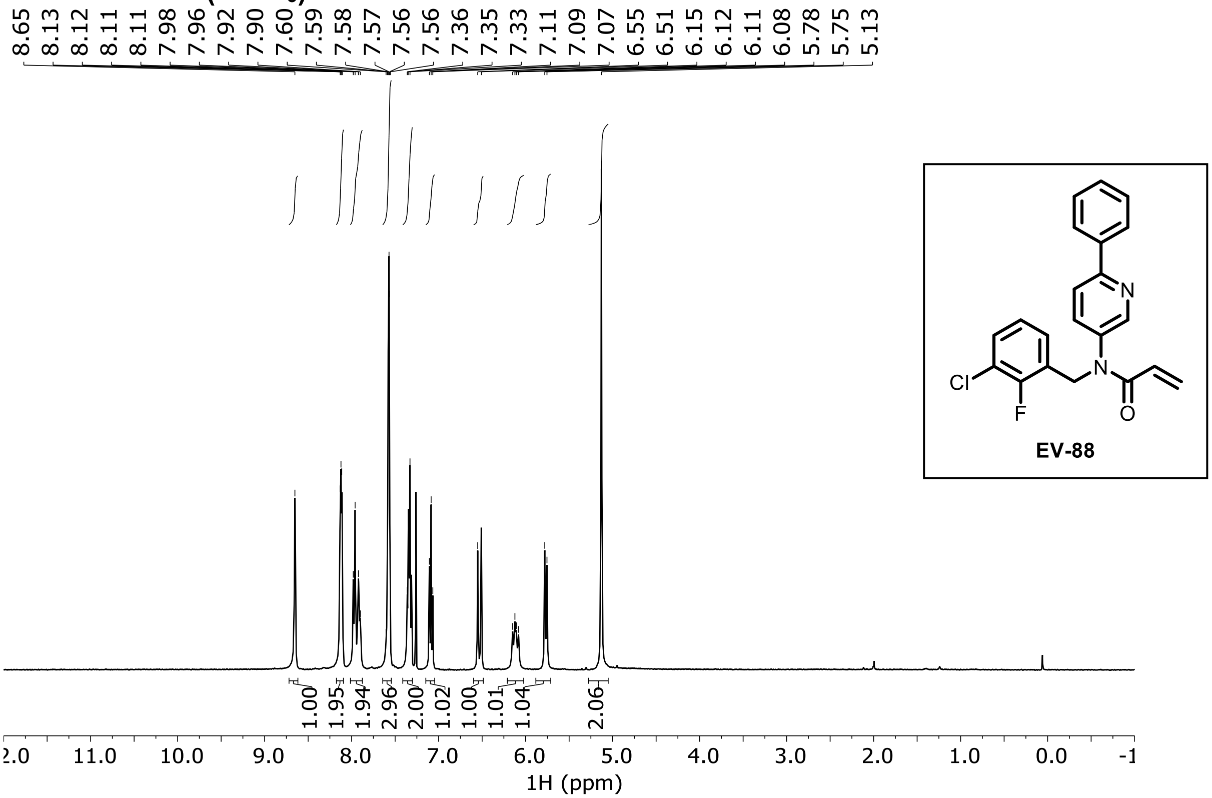

**EV-88**

**EV-89 $^1$H NMR ($CDCl_3$)**

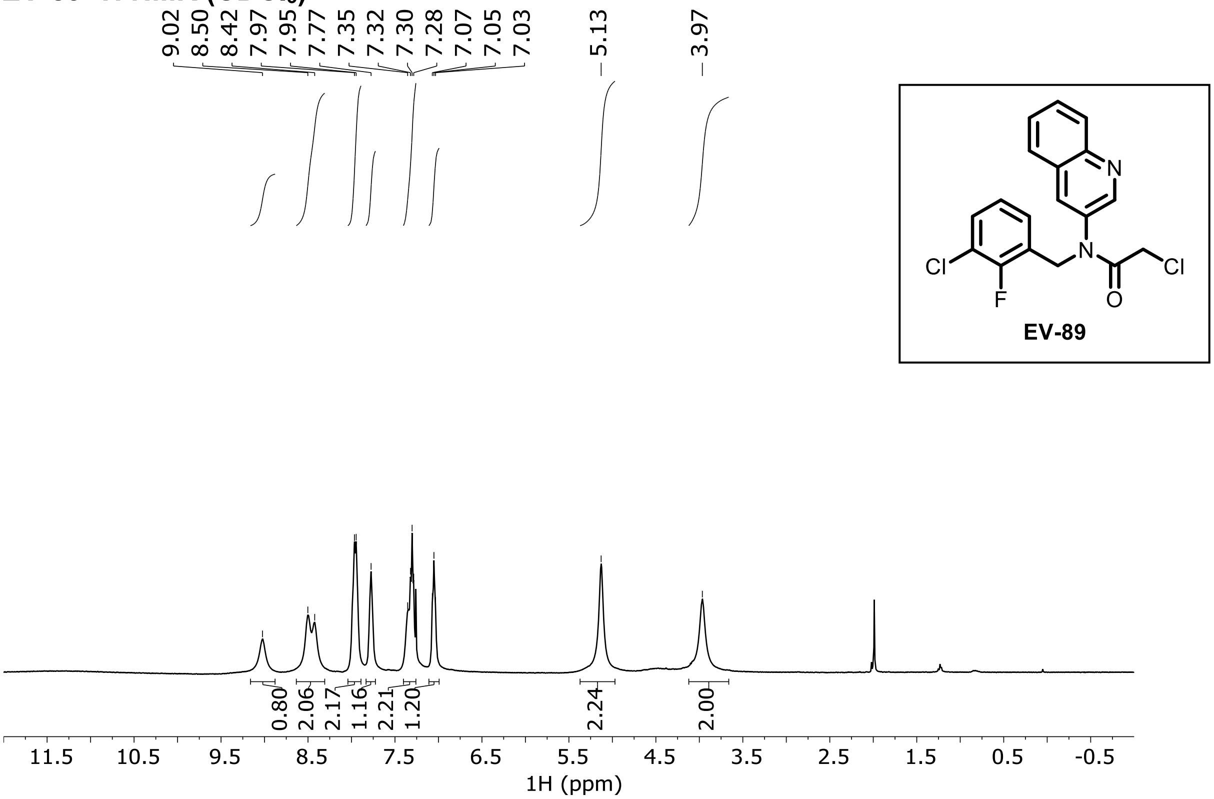

**EV-90 $^1$H NMR ($CDCl_3$)**

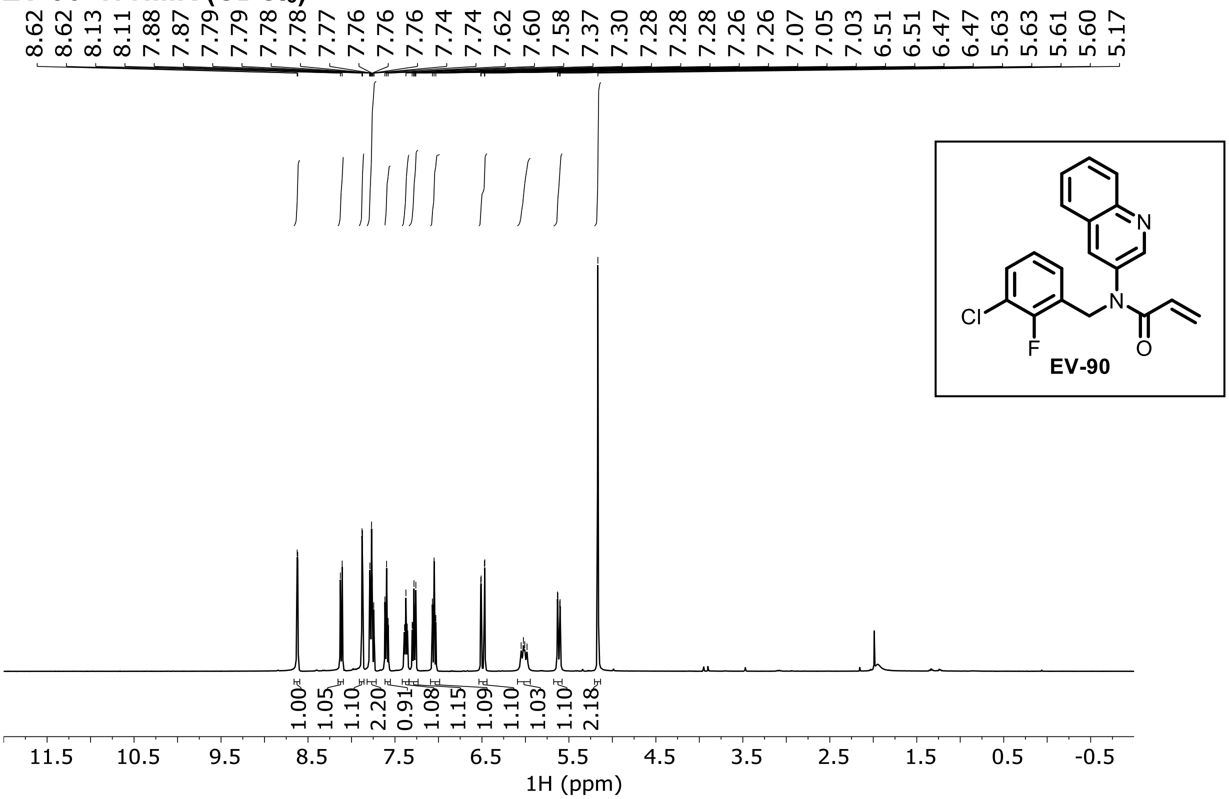

**EV-91 $^{1}$H NMR (DMSO-$d_6$)**

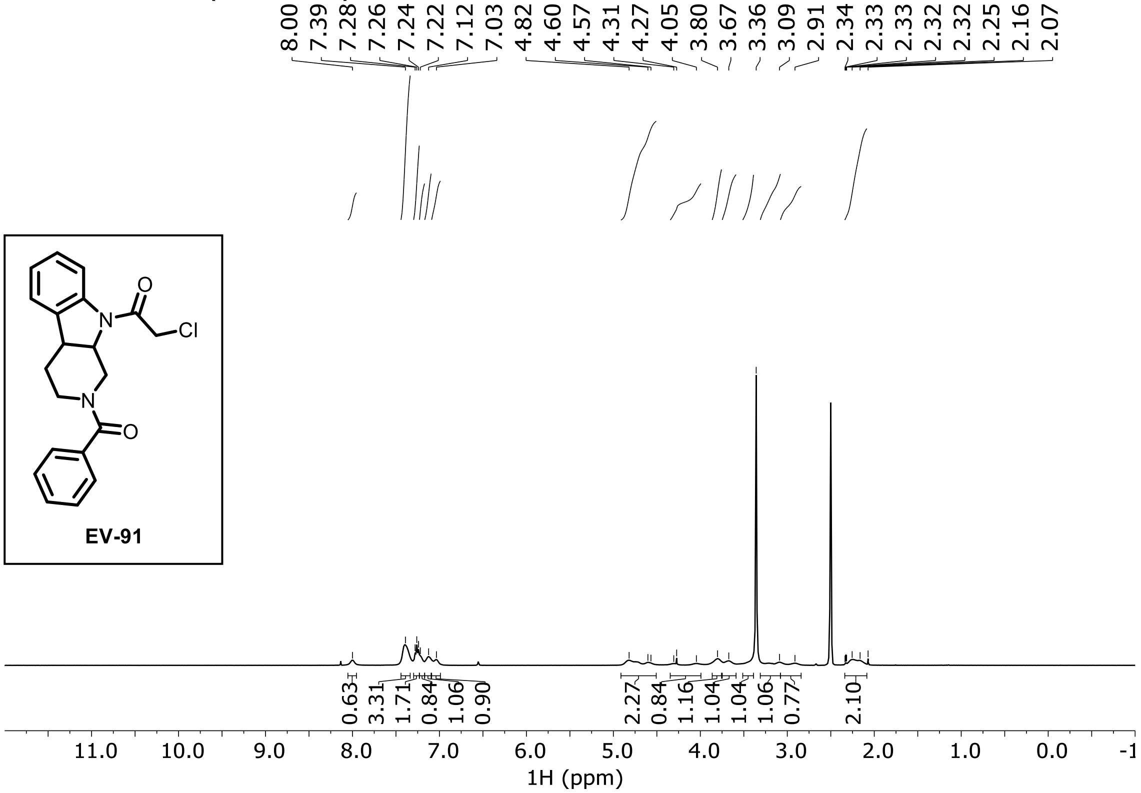

**EV-92 $^{1}$H NMR (CDCl$_3$)**

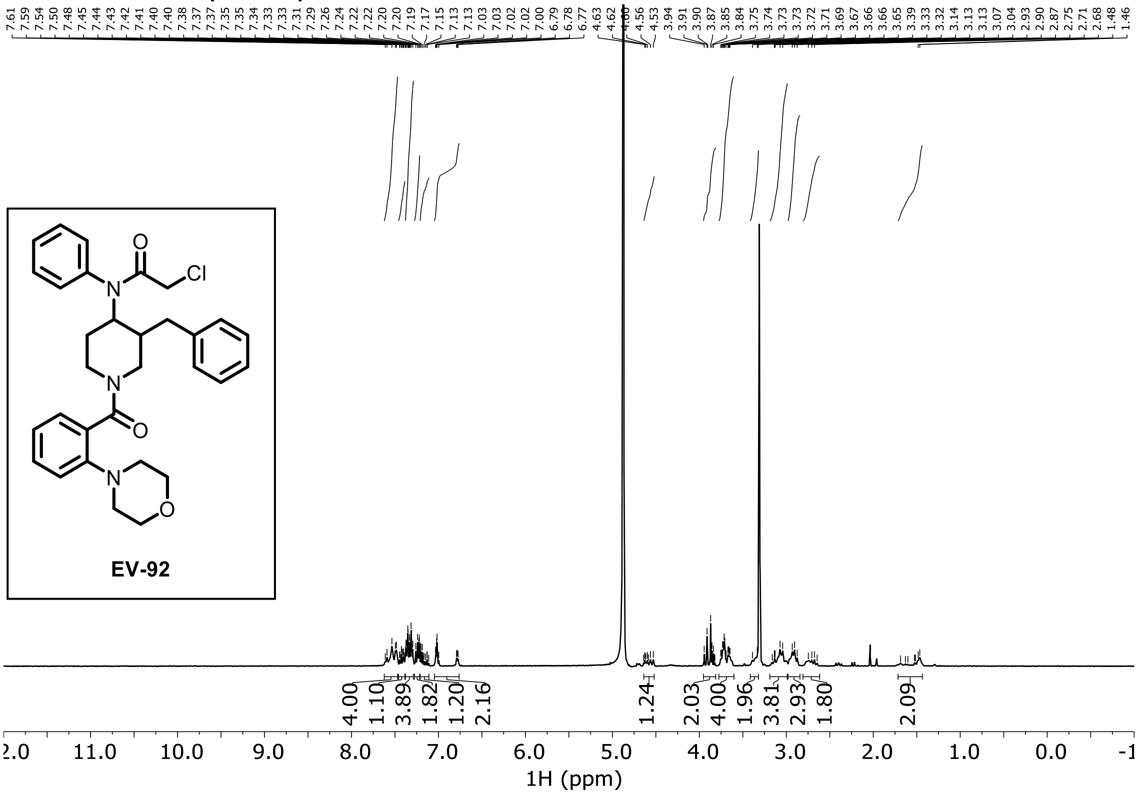

### EV-93 $^1$H NMR ($CDCl_3$)

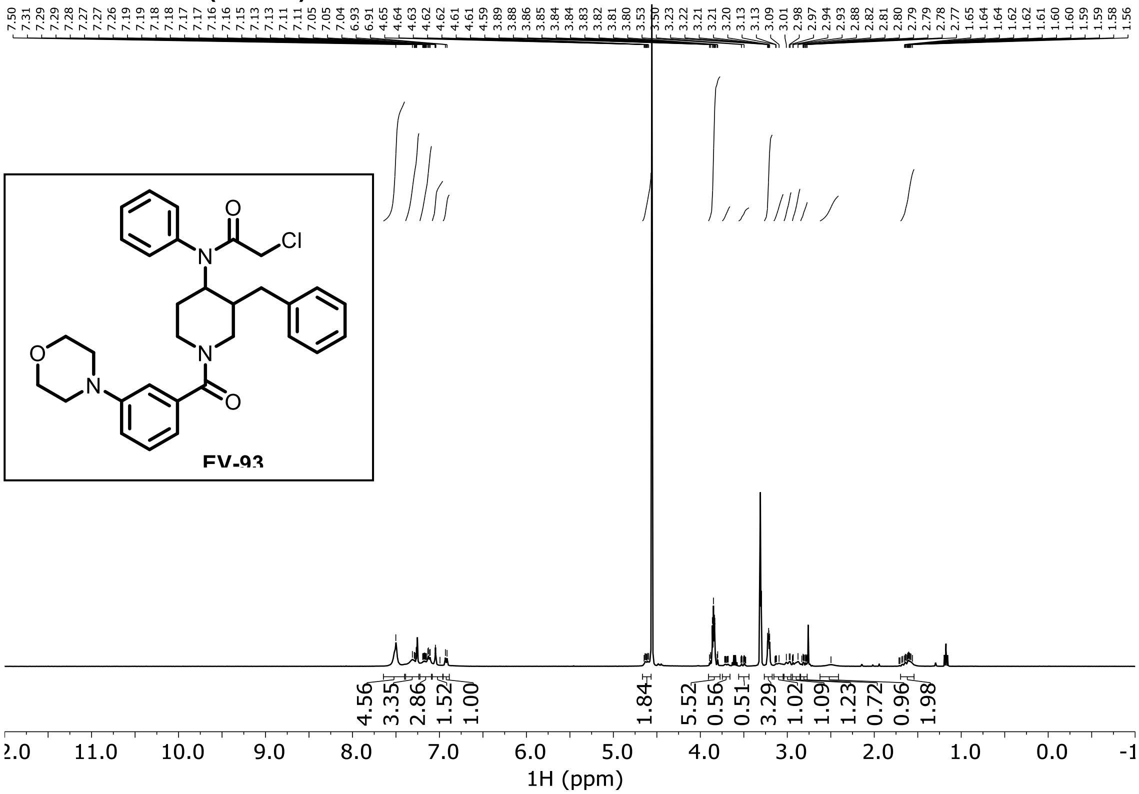

### EV-94 $^1$H NMR ($CDCl_3$)

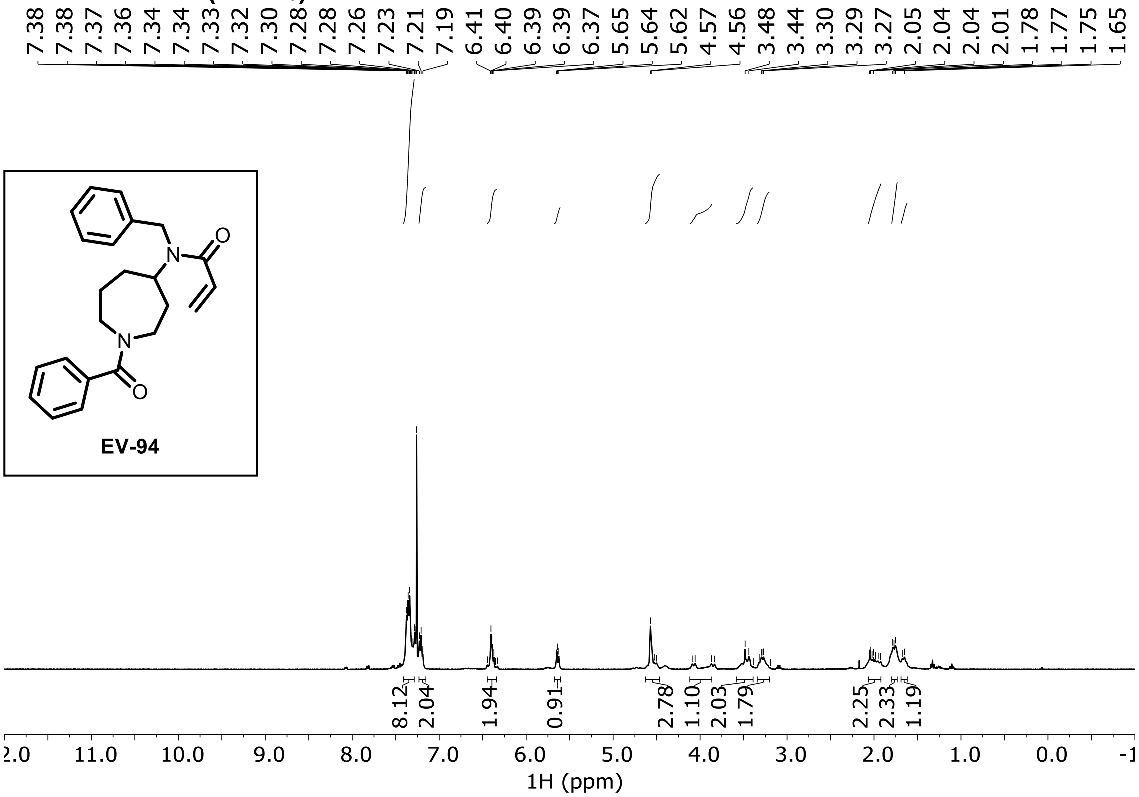

**EV-95 $^1$H NMR ($CDCl_3$)**

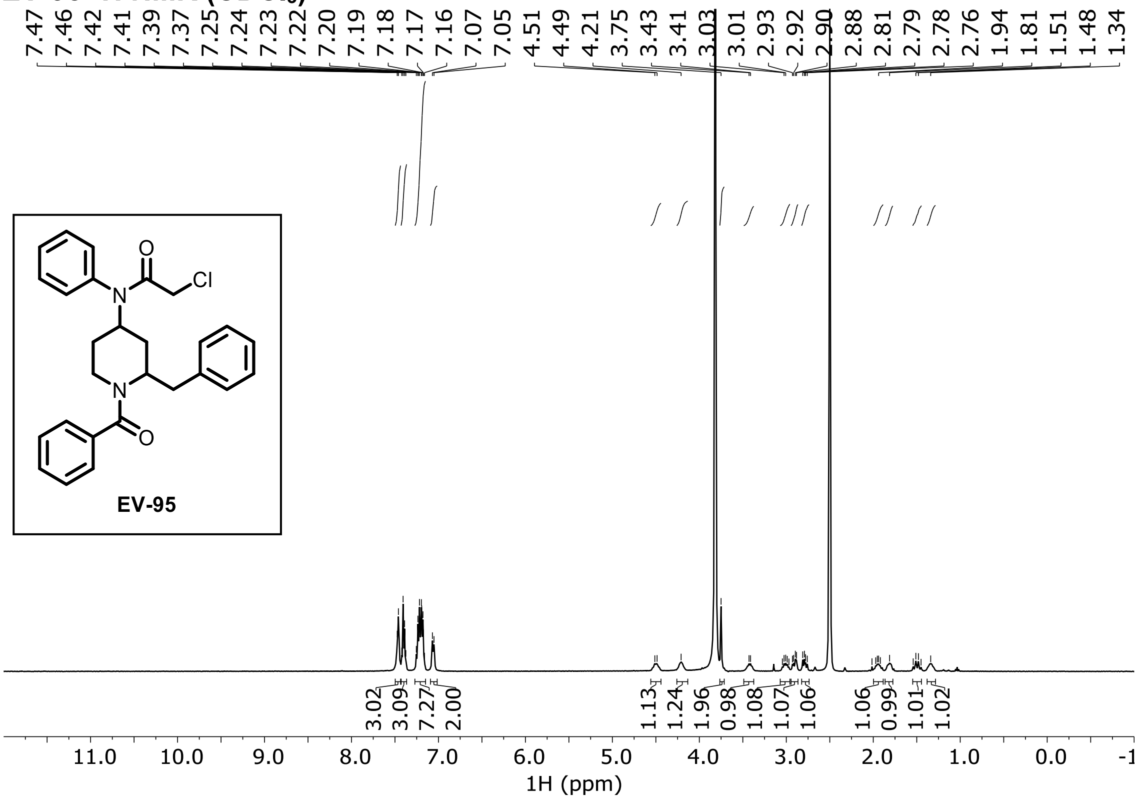
